# The effect of pre-analytical conditions on blood metabolomics in epidemiological studies

**DOI:** 10.1101/513903

**Authors:** Diana L Santos Ferreira, Hannah J Maple, Matt Goodwin, Judith S Brand, Vikki Yip, Josine L Min, Alix Groom, Debbie A Lawlor, Susan Ring

## Abstract

**Background:** Serum and plasma are commonly used biofluids for large-scale metabolomic-epidemiology studies. Their metabolomic profile is susceptible to changes due to variability in pre-analytical conditions and the impact of this is unclear.

**Methods:** Participant-matched EDTA-plasma and serum samples were collected from 37 non-fasting volunteers and profiled using a targeted nuclear magnetic resonance (NMR) metabolomics platform (*N*=151 traits). Metabolic concentrations were compared between reference (*pre*-storage: 4°C, 1.5h; *post*-storage: no sample preparation or NMR-analysis delays) and four, *pre*-storage, blood processing conditions, where samples were incubated at (i) 4°C, 24h; (ii) 4°C, 48h; (iii) 21°C, 24h; (iv) 21°C, 48h, before centrifugation; and two, *post*-storage, sample processing conditions in which samples (i) thawed overnight, then left for 24h before addition of sodium buffer followed by immediate NMR analysis; (ii) thawed overnight, addition of sodium buffer, then left for 24h before profiling. Linear regression models with random-intercepts were used to assess the impact of these six pre-analytical conditions on EDTA-plasma/serum metabolome.

**Results:** Fatty acids, beta-hydroxybutyrate, glycoprotein-acetyls and most lipid-related traits, in serum and plasma, were robust to the tested *pre* and *post*-storage conditions. *Pre*-storage conditions impacted concentrations of glycolysis metabolites, acetate, albumin and amino-acids by levels that could potentially bias research results (up to 1.4SD difference compared with reference). *Post*-storage conditions affected histidine, phenylalanine and LDL-particle-size, with differences up to 1.4SD.

**Conclusions:** Most metabolic traits are robust to the *pre*- and *post*-storage conditions tested here and that may commonly occur in large-scale cohorts. However, concentrations of glycolysis metabolites, and amino-acids may be compromised.

**Key messages:**

- In large scale epidemiological studies, blood processing delays, incubation at high temperature prior to long term storage, and NMR profiling delays after long term storage, may occur.
- Concentrations of fatty acids, beta-hydroxybutyrate, glycoprotein acetyls and most lipid-related traits are robust to variations in *pre*-storage temperature and duration of incubation (4°C or 21°C for up to 48h prior to centrifugation) and *post*-storage sample handling (24h delay in sample preparation or NMR profiling).
- Glycolytic metabolite concentrations are altered by *pre*-storage conditions and amino-acids, particularly histidine and phenylalanine, by both, *pre* and *post*-storage conditions.

## Introduction

The development of high throughput methods for quantifying multiple ‘omic traits in large-scale epidemiological studies, using stored biosamples, has the potential to rapidly advance our understanding into how human physiology and metabolism vary across the lifecourse. Applying these methods to existing samples from very densely pheno and geno-typed cohorts also has the potential to enhance our understanding of causal mechanisms for disease. To do this in a scientifically rigorous way, further information is required regarding the potential impact of differing pre-analytical conditions (e.g. variations in incubation duration and temperature before centrifugation) of stored samples that are likely to vary between cohorts and potentially within the same cohort over time. For example, since sample collection from longitudinal epidemiological cohorts are collected over many decades it may be impossible to ensure identical protocols are used throughout the life time of the study. In clinical cohorts and in pregnancy/birth cohorts, such as the Avon Longitudinal Study of Parents and Children (ALSPAC)^1, 2^ and Born in Bradford^3^ cohorts, some samples (e.g. during antenatal care or for cord-blood) are obtained during routine clinical practice where health care needs takes precedence over speedy sample processing and rapid storage. Yet these samples, collected early in life, are often the most valuable for research related to the long-term effects of childhood exposures. Other cohorts face different challenges, for example the Health Survey for England^4^ and Understanding Society^5^, collect biological samples in participants’ homes, which is likely to increase pre-analytical variability.

Additionally, we are increasingly interested in cohort collaborations (e.g. UCL-LSHTM-Edinburgh-Bristol^6^ consortium, COnsortium of METabolomics Studies^7^, Cohorts for Heart and Aging Research in Genomic Epidemiology^8^ consortium) and broader international comparisons, for example between determinants of health and wellbeing in low-, middle- and high-income countries and how these differ. Consequently, we often need to combine data from samples collected in very different circumstances. For all these reasons, it is important to recognise that biosample collection protocols are often a compromise between reducing any effects of pre-analytical variation with maximising the amount of data, that can be obtained from samples, within the financial and practical restraints of a given collection sweep.

In clinical chemistry, pre-analytics account for 60-80% of laboratory errors^9–12^ and current standard operating procedures (SOP) for blood handling in metabolomics, are generally based on best practices^13^ (not evidence based) for conventional biochemistry tests.^9, 14^ Whilst several studies have explored the impact of pre-analytical conditions on a small number of commonly assessed biomarkers in epidemiology^13, 15–18^, metabolomics – the simultaneous quantification of large numbers of metabolic traits – has particular challenges as different metabolites may have different susceptibilities to degradation. ^19–23^ The aim of this study was to determine the impact of different pre-analytical conditions, that reflect conditions potentially arising during sample collection and processing in large scale epidemiological studies, on serum and plasma metabolome.

## Methods

Samples from 37 healthy volunteers were collected. Exclusion criteria for the study were: clotting/bleeding disorders, anaemia, use of anti-coagulant medication or insulin treatment, and presence of blood borne viruses. Participants provided written informed consent and completed a questionnaire. Ethical approval for the study was obtained from South West Frenchay Proportionate Review Committee, Bristol, UK, reference 14/SW/0087.

An overview of the experimental design is given in sFigure 1. Ten blood tubes were drawn for each non-fasting participant (5 EDTA-plasma, 5 serum). Further details are provided in Supplementary methods.

### Reference samples

All samples were processed within 1.5h of blood withdrawal. Once centrifuged, serum and plasma samples were aliquoted into 1.5ml microtubes (STARLAB E1415-2240) and immediately frozen and stored at −80°C for one month prior to NMR analysis. Prior to profiling frozen samples were thawed in the refrigerator (4°C) overnight, prepared with sodium phosphate buffer^24–26^ and then immediately run through the NMR spectrometer.

### *pre*-storage handling: effect of pre-centrifugation delay and temperature

We compared metabolic trait concentrations, quantified by the NMR platform, between four *pre*- and two *post*-storage conditions and the reference samples. The four *pre*-storage conditions had the following combination of incubation temperature/duration before centrifugation (i) 4°C for 24h; (ii) 4°C for 48h; (iii) 21°C for 24h; (iv) 21°C for 48h. These conditions were chosen to reflect *pre*-storage variations likely to occur in epidemiological studies. Participant-matched serum and EDTA-plasma samples were processed according to these four conditions.

### *Post*-storage handling: sample preparation and sample NMR profiling delay

Two variations to the standard Nightingale^®^ NMR protocol^24–26^ were investigated to evaluate the impact of sample preparation and instrumental analysis delay. Matched serum and plasma aliquots of each participant previously prepared according to the reference *pre*-storage conditions, above described, were subjected to two *post*-storage conditions. Samples were either (i) thawed overnight (4°C), then left for 24 hours at 4°C in the dark before addition of sodium phosphate buffer followed by immediate NMR analysis (sample preparation delay); (ii) thawed overnight, addition of buffer before being left for 24 hours at 4°C in the dark, then NMR profiling (NMR analysis delay).

### Nuclear Magnetic Resonance metabolomics platform

A high-throughput NMR metabolomics platform, Nightingale Health^®^, at the University of Bristol, was used to quantify up to 151 lipoproteins, lipid and metabolites in serum and plasma. The platform applies a single experimental setup, providing the simultaneous quantification of routine lipids, 14 lipoprotein subclasses and lipids transported by these particles, various fatty acids (FA) and FA traits (e.g. chain length, degree of unsaturation), amino acids, ketone bodies, glycolysis and gluconeogenesis-related metabolites, fluid balance and one inflammation metabolite. Most of these are quantified in clinically meaningful concentrations (e.g. mmol/L), and particle size in nm. Details of this platform and its use in epidemiological studies have been described elsewhere^27–30^. Pyruvate, glycerol and glycine are not quantified in EDTA-plasma samples due to the interfering resonances of EDTA on their signals.

### Statistical Analysis

We used linear regression models with random intercepts to assess the impact of the different pre-analytical conditions on metabolic trait concentrations. Such models account for within individual clustering of observations as each individual participant contributes with multiple samples for analysis. All metabolic traits were scaled to standard deviation (SD) units (by subtracting the mean and dividing by the standard deviation); this was done separately for plasma and serum, and pre and post-storage conditions. Standardization allows comparison of metabolic traits with different units and/or different concentration ranges. Results in measured concentration units (e.g. mmol/l) are provided in Supplementary Data. All analyses were conducted using participant-matched serum and plasma samples.

Full specification of the models can be found in the Supplementary methods. First, we evaluated the impact of *pre*-storage duration, before centrifugation, by keeping temperature constant (at 4°C and 21°C). Incubation duration was entered as a continuous term (levels: 0=reference, 1=24h, 2=48h) in the model with betas representing the standardized mean difference in metabolite concentration per 24h increment in incubation time at 4°C and 21°C respectively. Next, we investigated the impact of *post*-storage conditions to estimate the standardized mean difference in metabolite concentration comparing delays in sample preparation and NMR profiling to the reference (levels: 0=reference, 1=sample preparation delay or NMR profiling delay). In addition, as the usefulness of measurements for epidemiological analysis often depends on the correct ranking of individuals (i.e. between-individual variations should be much bigger than within-individual variation due to pre-analytical handling/between days), we calculated the Spearman rank correlation coefficient between metabolic concentrations of reference samples and samples subjected to non-ideal conditions for *pre* and *post*-storage studies.

To better appreciate the effects of pre-storage temperature and duration of incubation on the overall metabolic profile,^31^ we conducted a principal component analysis (PCA)^32^. In PCA scores plots, the metabolic profile of each sample is shown as a single data-point, therefore each participant is represented by five data-points corresponding to each *pre*-storage condition. Samples (data-points) close to one another have similar metabolic composition in comparison to samples further apart.^33^ Therefore, if *pre*-storage conditions do not impact metabolic profiles, samples will not cluster by *pre*-storage conditions.

Statistical analyses were conducted using R version 3.0.1 (R Foundation for Statistical Computing, Vienna, Austria).

## Results

Characteristics of study participants are shown in Table 1, with metabolic trait distributions in sTable

**Table 1.** Characteristics of study participants. (N=37) who contributed to at least one pair of exposure-outcome analysis. Age was collected in categories.

| Characteristics | n | % |
| --- | --- | --- |
| **Female** | 29 | 78 |
| **Age** |  |  |
| [21-35] | 17 | 46 |
| [36-50] | 10 | 27 |
| [51-65] | 10 | 27 |
| **Ever smoked, No** | 15 | 41 |
| **Time since last alcohol consumption** |  |  |
| Never Drinks | 2 | 5 |
| <1 Week | 16 | 43 |
| 1-4 Weeks | 4 | 11 |
| 24h before blood collection | 15 | 41 |

1. Most participants were female (78%) and 46% were between 21-35 years old; 41% had drunk alcohol within the 24h period prior to blood sampling.

The following main conclusions can be drawn from our analyses of the effect of *pre*-analytical conditions on blood metabolic traits:

- Fatty acids, beta-hydroxybutyrate, glycoprotein acetyls and most lipids and lipoproteins, in serum and plasma, are robust to incubation temperatures of 4°C or 21°C for up to 48h prior to centrifugation (Figure 1, 2; sFigure 2; sTables 2, 3). They are also resilient to being left to thaw for up to 24h before buffer is added or having buffer added after overnight thaw and then left for a further 24h before NMR profiling (Figure 3, 4; sFigure 3; sTables 4, 5; sTables 6, 7).
- Labile traits include glycolysis related metabolites (glucose, lactate, pyruvate) and amino-acids which showed marked differences in concentration in relation to *pre*-storage conditions and *post*-storage conditions (amino-acids only) (Figure 2, 4; sFigure 4; sTables 2-5).
- Branched and aromatic amino-acids appear more robust to *pre*-storage condition in EDTA-plasma than in serum (Figure 2; sTables 2, 3).
- For pyruvate, *pre*-storage delay and temperature appeared to interact, with levels slightly decreasing/stable from reference levels per 24h at 4°C and increasing per 24 hours at 21°C (Figure 2; sTables 2, 3).
- As an illustration of the magnitude of effects in serum, glucose levels decreased by 0.91mmol/l (95%CI: 0.87 to 0.95) and 1.9mmol/l (95%CI: 1.7 to 2.1) for each 24h delay, compared with the reference sample, at 4°C and 21°C, respectively. Pyruvate levels decreased by 0.013mmol/l per 24h (95%CI: 0.008 to 0.018) at 4°C but increased by 0.73mmol/l per 24h (95%CI: 0.64, 0.82) at 21°C.
- *Post*-storage conditions affected histidine, phenylalanine and LDL-particle size, with changes up to 1.4SD from reference (Figure 3, 4; sTables 4, 5).
- 71.5% and 93% of metabolites across pre and post-storage conditions, respectively, had spearman correlations coefficients, between ideal and variant conditions, above 0.8 (sTables 6-9).
- Diacylglycerol (DAG) and histidine (range: 0.1-0.8); and DAG, histidine, acetate, pyruvate, lactate (range: 0.3-0.5), across pre and post-storage conditions, respectively, have the lowest correlations (sTables 6-9).

**Figure 1.**
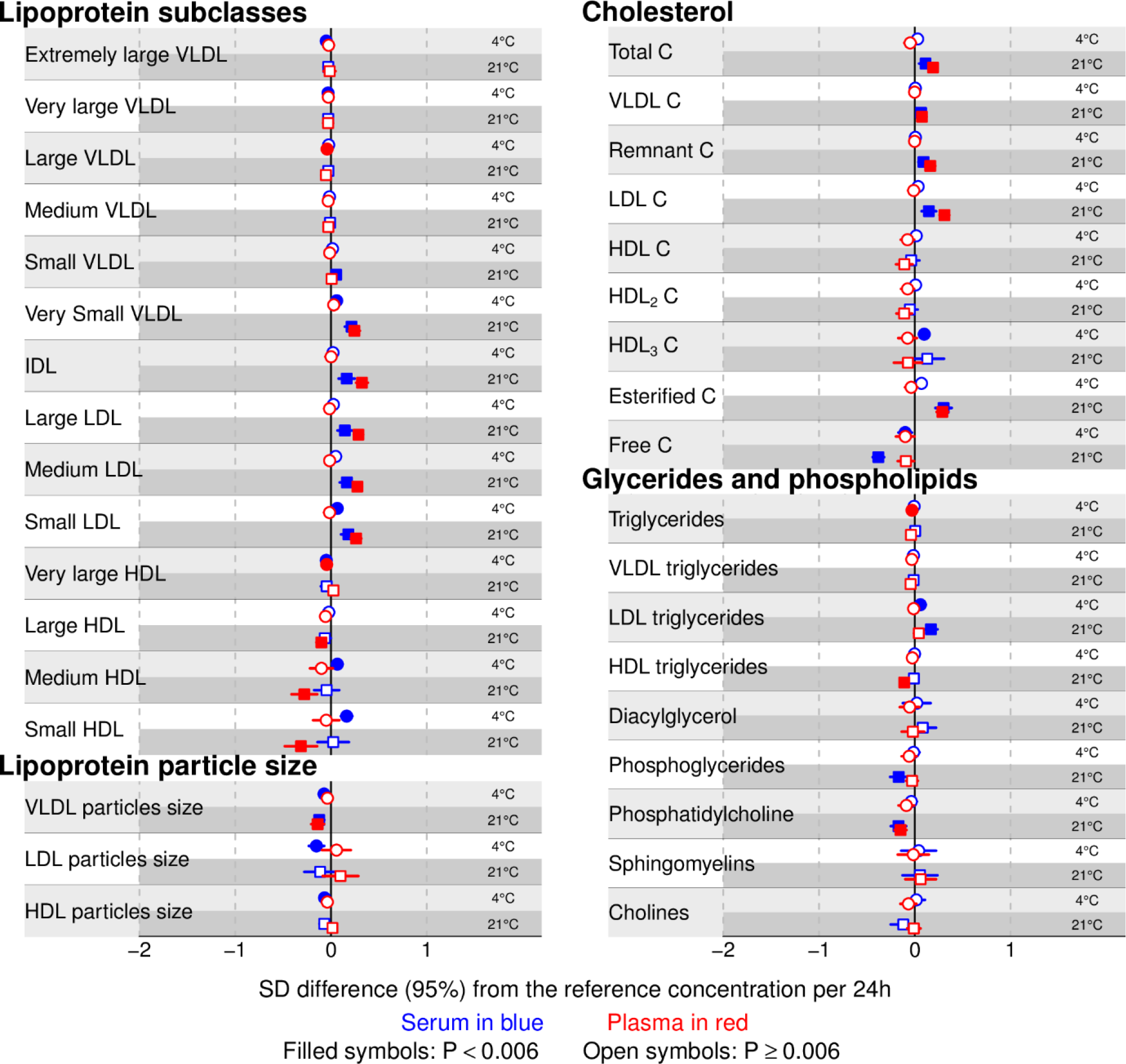
**Pre-storage handling effects:** standardized mean differences in metabolite concentrations (or trait value) per 24h increment in incubation duration at 4°C and 21°C. Standardized mean differences are given for serum and plasma traits (associations for detailed lipoprotein traits are given in sFigure 2). Mean differences in absolute units are listed in sTables 2 and 3. Pyruvate, glycerol and glycine are not quantified in Ethylenediaminetetraacetic acid (EDTA)-plasma samples due to the interfering resonances of EDTA on their signals. **Abbreviations**: **C**=cholesterol; **IDL**=intermediate-density lipoprotein; **LDL**=low-density lipoprotein; **HDL**=high-density lipoprotein; **VLDL**=very-low-density lipoprotein

**Figure 2.**
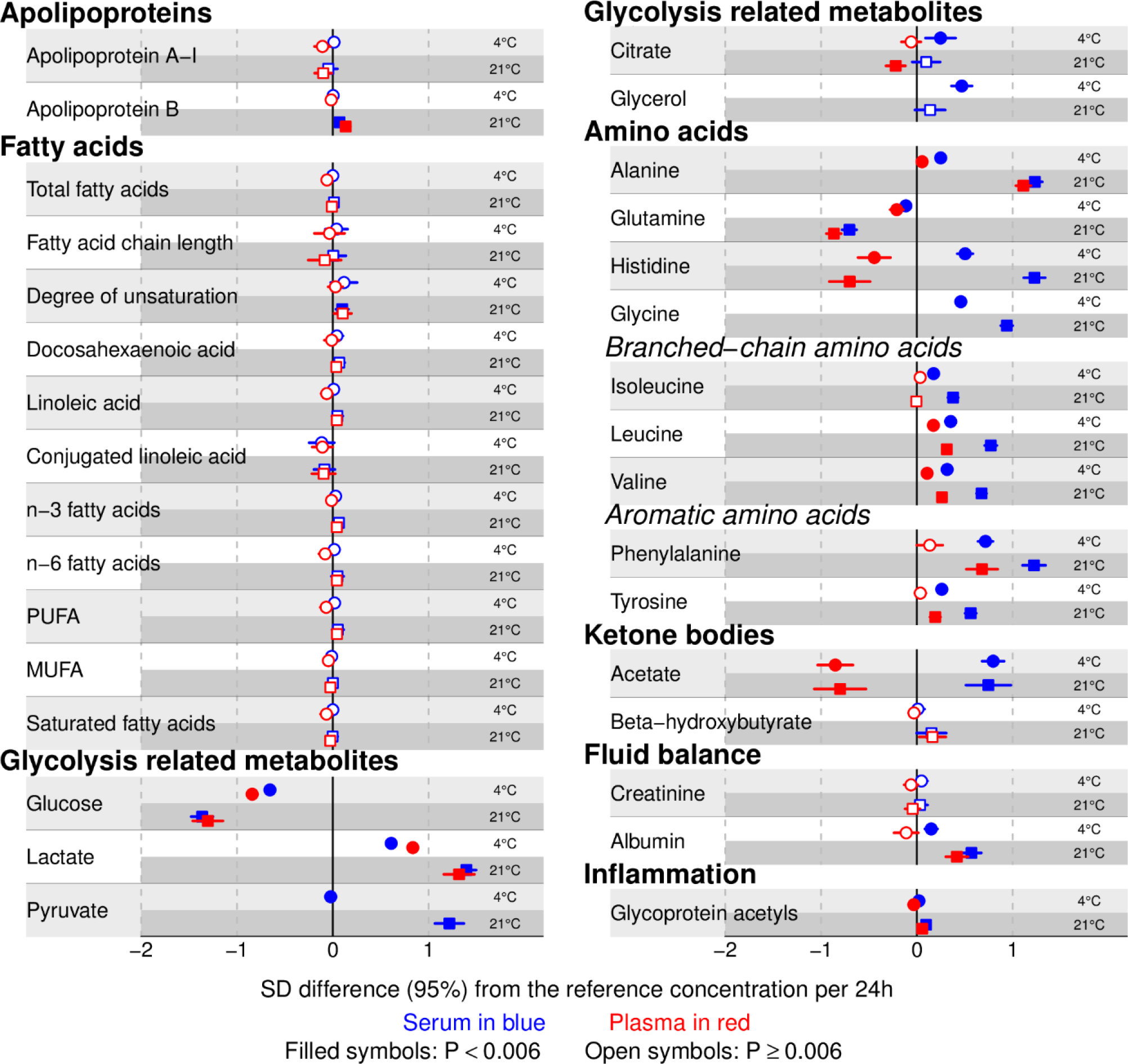
**Pre-storage handling effects (Figure 1 continued):** standardized mean differences in metabolite concentrations (or trait value) per 24h increment in incubation duration at 4°C and 21°C. Standardized mean differences are given for serum and plasma traits (associations for detailed lipoprotein traits are given in sFigure 2). Mean differences in absolute units are listed in sTables 2 and 3. Pyruvate, glycerol and glycine are not quantified in Ethylenediaminetetraacetic acid (EDTA)-plasma samples due to the interfering resonances of EDTA on their signals. **Abbreviations**: **MUFA**=monounsaturated fatty acids; **PUFA**=polyunsaturated fatty acids.

**Figure 3.**
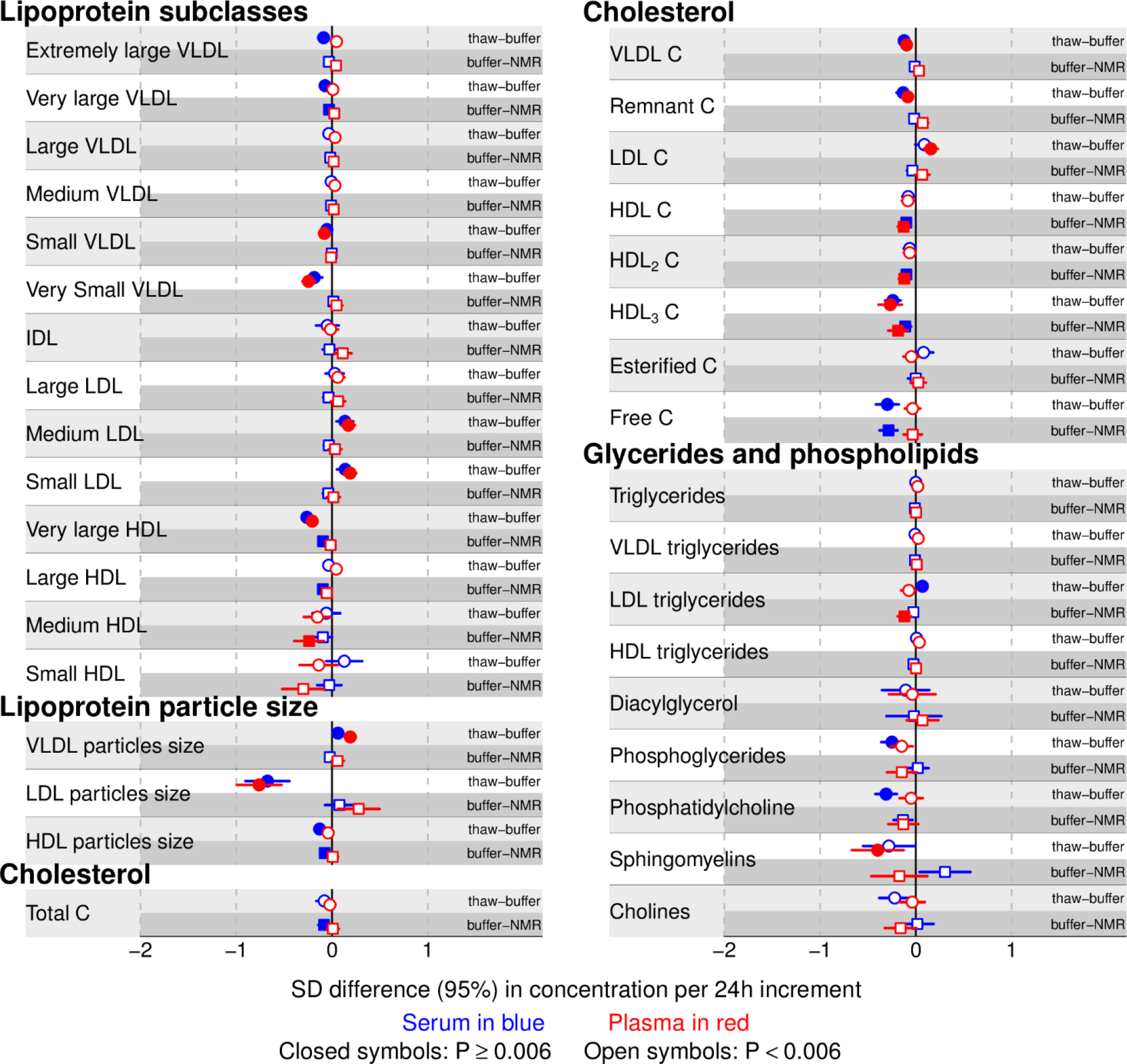
**Post-storage handling effects:** standardized mean differences in metabolite concentrations (or trait value) comparing delays in sample preparation (i.e. thaw to buffer addition delay) and NMR profiling (buffer addition to NMR profiling delay) to the reference, for serum and plasma samples (associations for detailed lipoprotein traits are given in sFigure 3). Mean differences in absolute units are listed in sTables 4 and 5. Pyruvate, glycerol and glycine are not quantified in Ethylenediaminetetraacetic acid (EDTA)-plasma samples due to the interfering resonances of EDTA on their signals. **Abbreviations**: **C**=cholesterol; **IDL**=intermediate-density lipoprotein; **LDL**=low-density lipoprotein; **HDL**=high-density lipoprotein; **VLDL**=very-low-density lipoprotein.

**Figure 4.**
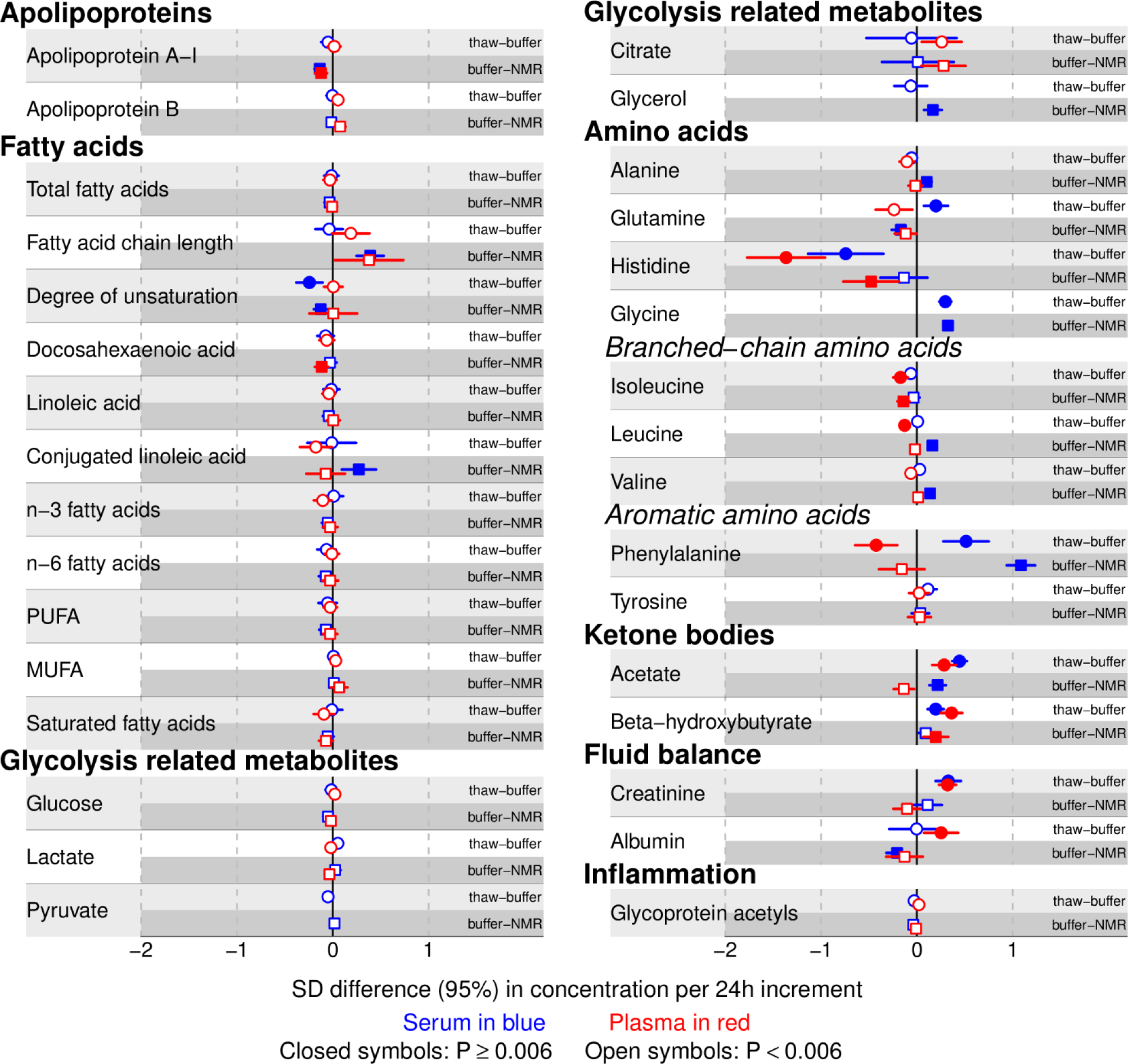
**Post-storage handling effects (Figure 3 continued):** standardized mean differences in metabolite concentrations (or trait value) comparing delays in sample preparation (i.e. thaw to buffer addition delay) and NMR profiling (buffer addition to NMR profiling delay) to the reference, for serum and plasma samples (associations for detailed lipoprotein traits are given in sFigure 3). Mean differences in absolute units are listed in sTables 4 and 5. Pyruvate, glycerol and glycine are not quantified in Ethylenediaminetetraacetic acid (EDTA)-plasma samples due to the interfering resonances of EDTA on their signals. **Abbreviations**: **MUFA**=monounsaturated fatty acids; **PUFA**=polyunsaturated fatty acids.

## Discussion

In this experiment we have shown that most metabolites, including lipids, lipoproteins and fatty acids are unaffected by different sample pre-analytical conditions that were designed to reflect plausible variation in large-scale epidemiological studies that have collected samples at different times and different populations. 91% and 97% of metabolites across *pre* and *post*-storage conditions, respectively, had SD differences from the reference below 0.5SD. We did however find effects on glucose and related glycolysis metabolites, as well as on amino acids (up to 1.4SD).

Previous studies have addressed this using clinical chemistry tests, mass spectrometry and NMR, with targeted and untargeted platforms. Where it is possible to make comparisons, these are summarized in sTable 10.

*Pre*-storage delay and incubation temperature of *uncentrifuged* samples, can cause changes to serum and plasma metabolomes because blood cells and enzymes are still metabolically active inside the sample tube, resulting in the uptake and release of metabolites. Concordant findings include the reported stability of triglycerides ^16–18^, high density lipoprotein-cholesterol (C) ^18^, low density lipoprotein-C (4°C) ^18^, total cholesterol (4°C) ^18^, apolipoprotein A-I and B ^18^, and creatinine ^18^. The observed changes in glucose, pyruvate, lactate and alanine, agree with previous studies ^17–19, 34, 35^, and are likely due to blood cell activity, primarily red blood cells (RBC).^23, 36^ The absence of RBC in *post*-storage samples explains why these metabolites are robust to the tested *post*-storage conditions.^17, 20^ Increases in concentrations of most amino-acids with the *pre*- and *post*-storage (non-ideal) conditions that we tested here are also consistent with previous studies.^14, 34^ One possible reason for this is protein degradation, and in the case of phenylalanine^14, 34^ accumulation, as it is an uncommon component of proteins and has few degradation pathways.^14^ Glutamine decreased,^14, 34^ converted into glutamate, and high levels of the latter prevent reliable quantification of pyruvate, since pyruvate only gives one peak in the ^1^H-NMR-spectrum, which can be overlapped by glutamate signals. Our results also show that amino-acids in serum are less robust than in EDTA-plasma, possibly due to the inhibition of metal-dependent proteases by EDTA in the latter, and activation of a range of proteases during coagulation, in the former. Histidine and acetate had opposite changes in serum and plasma likely due to the influence of the presence/absence of anti-coagulant. *Post*-storage effects were less pronounced than *pre*-storage ones indicating that most sample degradation are related to blood cells activities.^23, 37^ Nevertheless we found changes in histidine and phenylalanine and, although not directly comparable to the conditions we tested, others ^14, 19^ have report changes in some metabolites when delays occur *after* centrifugation. These are potentially caused by enzymes released due to cell damage during centrifugation and other proteins. ^14, 19, 37^

### Study strengths and limitations

We have explored the effects of plausible variation in sample processing conditions on over 151 metabolic traits, which is an important contribution to the literature given the increasing use of high throughput metabolomics in stored samples from epidemiological studies. Samples from just 37 participants were used but the narrow confidence intervals of our results suggest this was sufficient to provide enough sample comparisons for precise estimation of effects. Participants did not fast before providing a sample, but this would only affect our results if fasting were related to the different conditions we have tested (e.g. processing delay or incubation temperature); as samples from each individual were subjected to all experimental conditions and none of the participants were asked to fast, we think this is unlikely. Whilst this study highlights which metabolic traits are sensitive to variations in pre-analytical conditions this does not tell us whether this will bias association analyses. If measurements for all participants tend to change by a similar amount such analyses may not be affected (and this appears to be the case for most metabolites as given by the spearman correlation results). Therefore, we would suggest that where results are being pooled across studies that have had different pre-analytical processing conditions, between study heterogeneity of glycolysis, amino acid, acetate and DAG associations should be looked at in detail.

### Conclusion

Most serum and EDTA-plasma metabolic traits, quantified by Nightingale Health^®^ NMR platform (87% of measured traits are lipid-related), are stable to the *pre* and *post*-storage conditions tested with exception of glycolysis metabolites (labile to *pre*-storage conditions) and amino-acids (labile to both *pre* and *post*-storage conditions). In large collaborations and longitudinal studies, the possible impact of processing conditions on between metabolomic-data heterogeneity should be explored.

## Funding

This work was supported by ‘Cohorts and Longitudinal Studies Enhancement Resources’ (CLOSER), as part of a collaborative research program and is funded by the ESRC grant reference: ES/K000357/1. DLSF, MG, HM, JM, SR, AG, and DAL work in a Unit that receives funds from the University of Bristol and the UK Medical Research Council (MC_UU_00011/6). DAL is a UK National Institute of Health Research Senior Investigator (NF-SI-0166-10196). No funding bodies had any role in study design, data collection and analysis, decision to publish, or preparation of the manuscript.

## Acknowledgements

The authors are very grateful to all the participants who donated blood samples for this study. Sample processing and analysis was carried out at the Bristol Bioresource Laboratory and the NMR Metabolomics facility at the University of Bristol.

## Conflict of Interests

DAL has received support from several National and International government and charitable funders as well as Medtronic LTD and Roche Diagnostics research that is not related to the study presented in this paper. The other authors report no conflicts.

