## Supplementary material for "The effect of pre-analytical conditions on blood metabolomics in epidemiological studies"

### Supplementary methods

Order of blood collection was (i) 5 x 5ml serum SSTII (NHS supply chain KFK114), (ii) 1 x 6ml K2 EDTA (NHS supply chain KFK286) and (iii) 4 x 4ml K3 EDTA (NHS supply chain KFK042). Immediately after blood withdrawal, each serum and EDTA-plasma tubes were inverted 6 and 8-10 times, respectively. Serum and EDTA-plasma samples were stored on ice while transported to the laboratory.

#### Reference samples

Serum tubes were placed upright in a tube rack in the refrigerator (4°C) for 45 minutes to allow blood clotting, then centrifuged at speed 1500 g for 10 minutes at 18-25 °C (3K15, Sigma Laborzentrifugen, Germany); EDTA-plasma tubes were centrifuged as soon as possible after blood withdrawal at 2260g for 10 minutes at 4-5°C (Thermo Scientific Heraeus Megafuge 16R).

#### Pre-storage handling: effect of pre-storage delay and temperature

Serum and plasma tubes where covered with tin foil and kept upright in a tube rack, and afterwards stored on a roller mixer in the dark (Bibby scientific®) at 4°C or 21°C, for 24h or 48h prior to centrifugation. Centrifugation was performed as per the reference protocol, afterwards samples were aliquoted and immediately froze and stored at -80°C for 1 month until Nuclear Magnetic Resonance (NMR) analysis.

#### Nuclear Magnetic Resonance (NMR) metabolomics platform

Up to 230 quantified metabolomic measures were obtained per sample of serum or EDTA-plasma, using a 1D proton ($^1$H) NMR spectroscopy-based platform described previously.[1-3] Briefly, 260 µL plasma and 260 µL sodium phosphate buffer (75 mM $Na_2HPO_4$, 0.08% sodium 3-(trimethylsilyl)propionate-2,2,3,3-$d_4$, 0.04% sodium azide in 80%/20% $H_2O/D_2O$, pH 7.4) were mixed and transferred to NMR tubes using an 8-channel, Varispan Janus liquid handling robot (PerkinElmer). NMR spectra were acquired using a Bruker Avance III HD 500MHz spectrometer with a room temperature 5mm, inverse triple resonance TXI probe and a Bruker Avance III HD 600MHz spectrometer equipped with a nitrogen-cooled triple resonance probe (CryoProbe Prodigy TCI). Both spectrometers were equipped with SampleJet auto-samplers with cooled (6°C) sample storage. Spectra were acquired using standardized parameters using three NMR experiments or 'molecular windows' to characterize lipoproteins, low molecular weight metabolites and lipids. Lipid spectra were acquired after a standardised lipid extraction procedure performed on each sample using a VIAFLO 96 channel electronic pipette (Integra Biosciences). Data pre-processing and quantification were as previously described.[1-3]

#### Statistical Analysis

Specification of random intercept model for analysis of pre-storage and post-storage conditions:

$$y_{ij} = \beta_0 + \mu_{0j} + (\beta_1)x_{ij} + e_{ij}$$

where $y_{ij}$ is the standardized or un-standardized (for figures and tables, respectively) metabolite concentration at each condition ($x_{ij}$) for individual *j* and $\beta_0$ representing the average intercept of the model or the overall mean. Deviations from the average intercept for individual *j* is represented by $\mu_{0j}$. The $e_{ij}$ term describes the deviation of the *i*th measurement on the *j*th individual from the mean standardized (or mean) metabolite concentration. This is the residual error term. For our analysis, $\beta_1$ is the fixed parameter of interest, representing the standardized mean (or mean) difference in metabolite concentration for a 1-unit increase in $x_{ij}$. For analysis of *pre*-storage conditions, incubation duration was entered as continuous term in the model (levels: 0=1.5h [reference], 1=24 h, 2=48 h) with $\beta_1$ representing the standardized mean (or mean) difference in metabolite concentration for each 24h increment in *pre*-storage incubation duration. This *pre*-storage condition model was run separately for samples incubated at 4 and 21$^0$C and for serum and EDTA-plasma samples. For analysis of *post*-storage conditions, $\beta_1$ is estimated for each level of *post*-storage condition, representing the standardized mean (or mean) difference in metabolite concentration comparing delays in (a) post-storage sample preparation and (b) NMR profiling, to the reference (levels: 0=reference, 1= sample preparation delay or NMR profiling delay).

#### Supplementary figures

##### *sFigure 1. Experimental design diagram: pre and post-storage conditions.*

##samples subject to *pre*-storage conditions 2 to 5, where subjected to reference *post*-storage conditions.

**Abbreviations**: **NMR**=Nuclear Magnetic Resonance; **buffer***= sodium phosphate (see supplementary methods for more information).

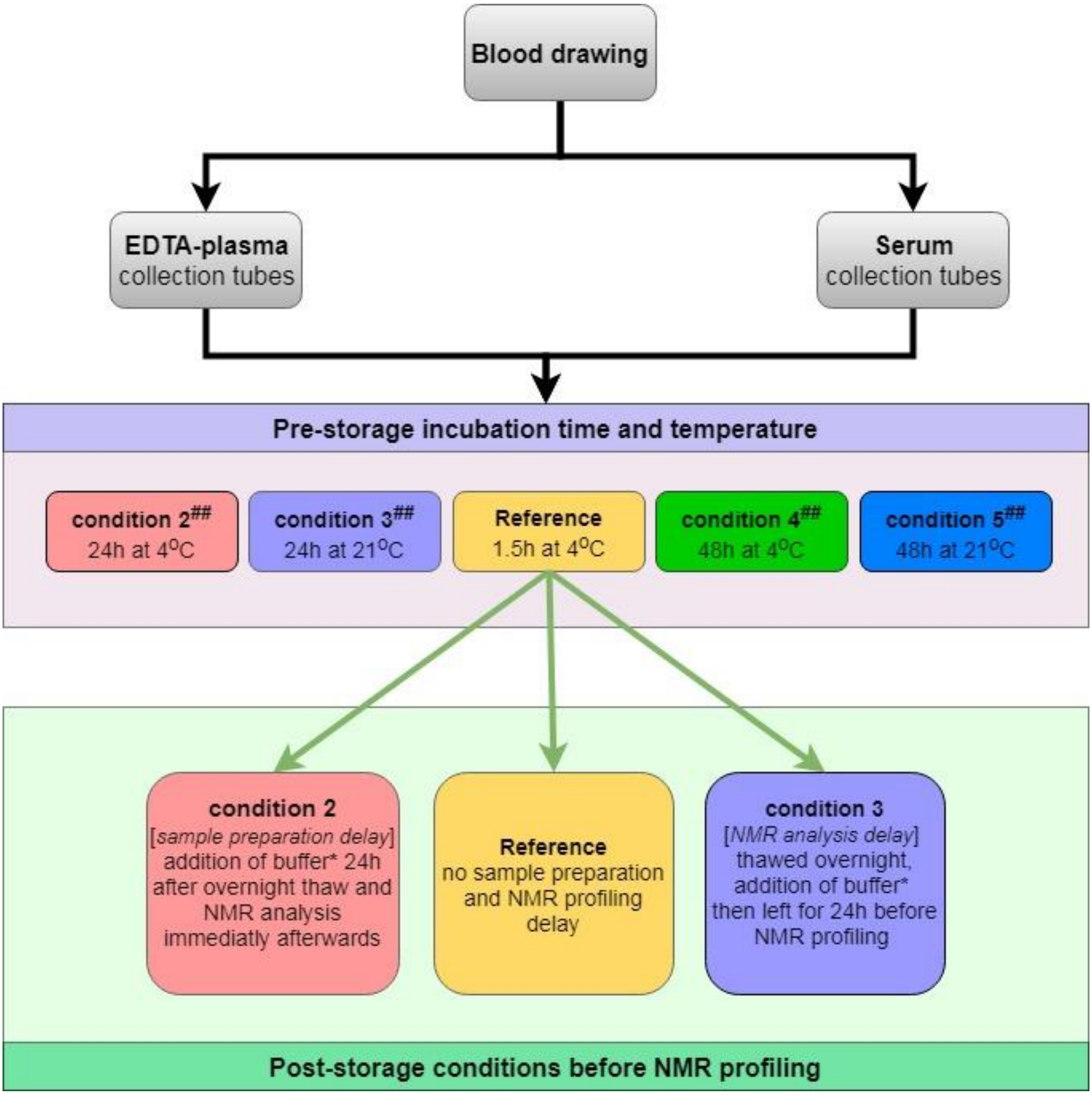

*sFigure 2. Pre-storage handling effects: standardized mean differences in lipoprotein particle and lipid concentration per 24h increment in incubation duration at 4°C and 21°C. Standardized mean differences are given for serum and plasma lipoprotein traits (associations for other metabolic traits are given in Figure 1 and 2). Mean differences in absolute units are listed in sTables 2 and 3.*

***Abbreviations**: **VLDL**=very-low-density lipoprotein.*

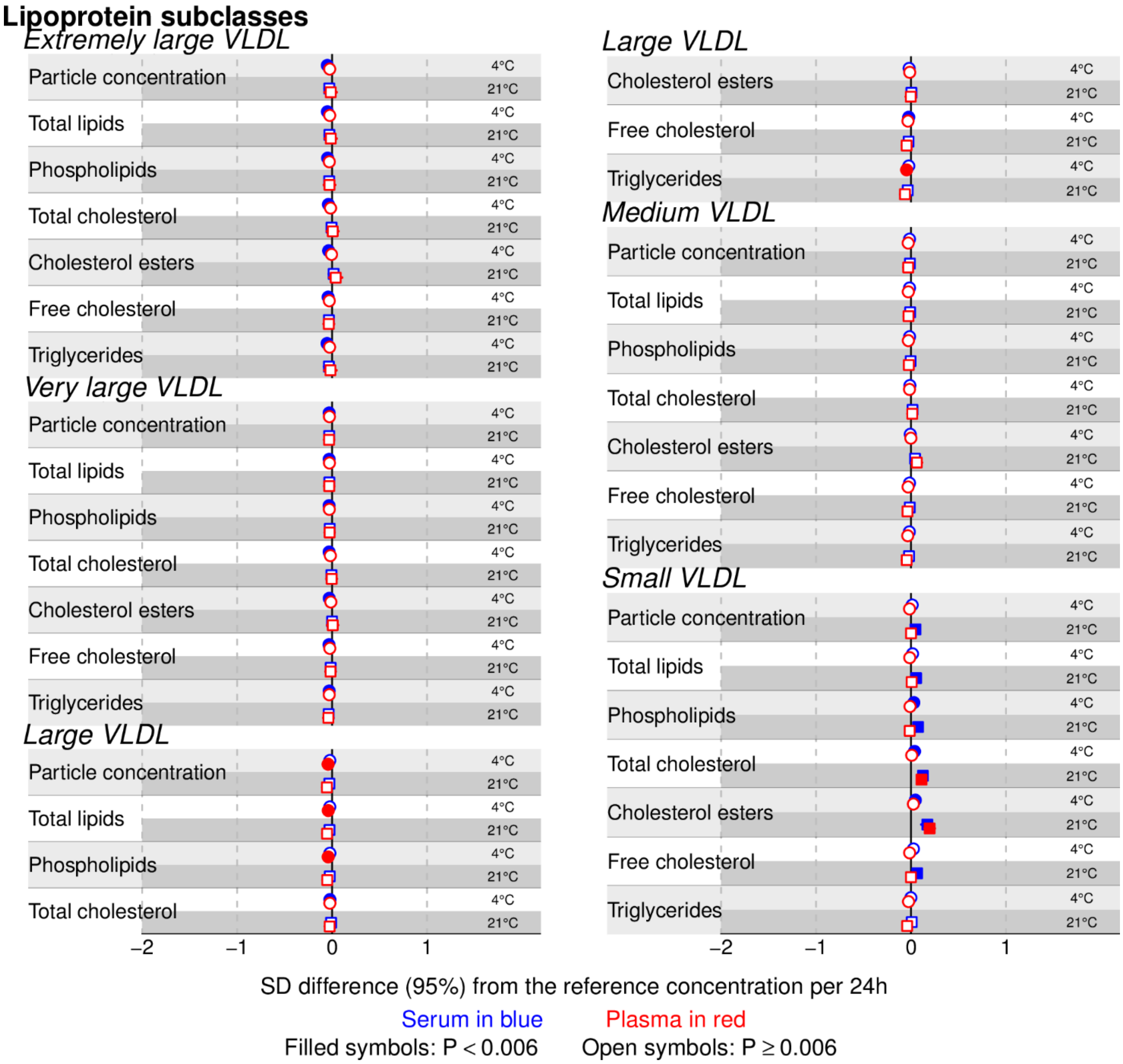

*sFigure 2 (continued). Pre-storage handling effects: standardized mean differences in lipoprotein particle and lipid concentration per 24h increment in incubation duration at 4°C and 21°C. Standardized mean differences are given for serum and plasma lipoprotein traits (associations for other metabolic traits are given in Figure 1 and 2). Mean differences in absolute units are listed in sTables 2 and 3.*

***Abbreviations**: **IDL**=intermediate-density lipoprotein; **LDL**=low-density lipoprotein; **VLDL**=very-low-density lipoprotein.*

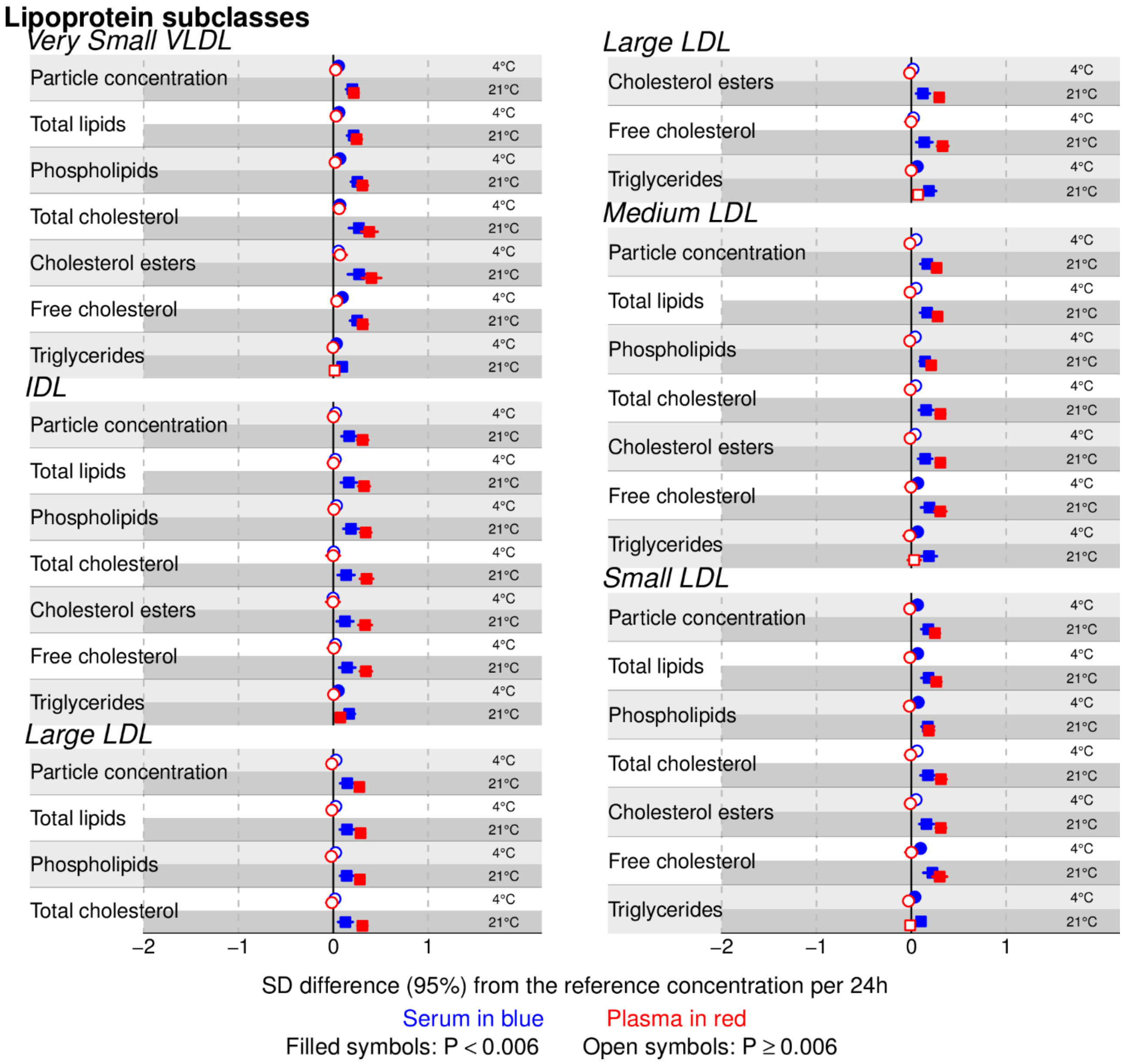

*sFigure 2 (continued). Pre-storage handling effects: standardized mean differences in lipoprotein particle and lipid concentration per 24h increment in incubation duration at 4°C and 21°C. Standardized mean differences are given for serum and plasma lipoprotein traits (associations for other metabolic traits are given in Figure 1 and 2). Mean differences in absolute units are listed in sTables 2 and 3.*

***Abbreviations**: **HDL**=high-density lipoprotein.*

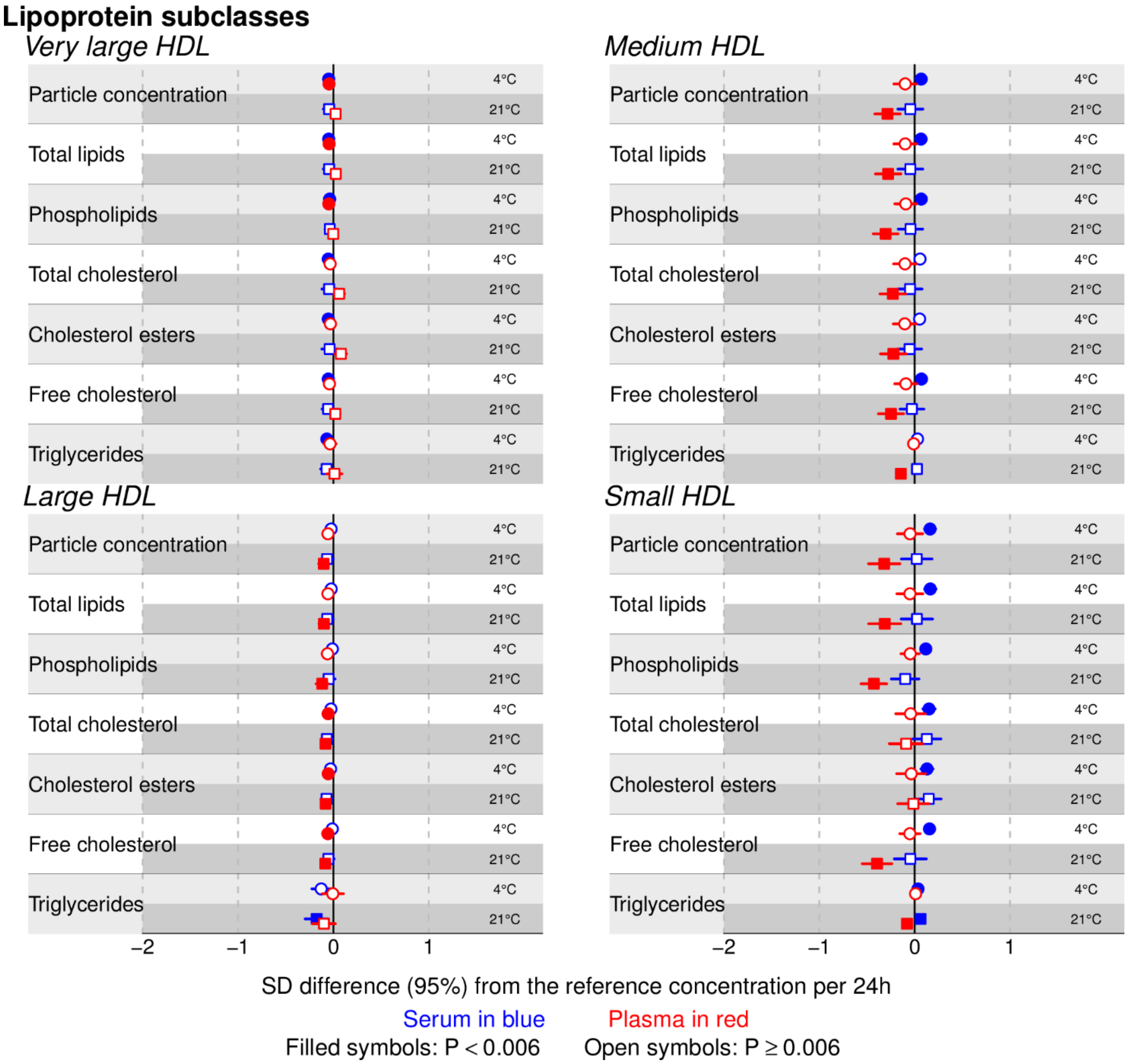

*sFigure 3. Post-storage handling effects: standardized mean differences in lipoprotein particle and lipid concentration comparing delays in sample preparation (i.e. thaw to buffer addition delay) and NMR profiling (buffer addition to NMR profiling delay) to the reference, for serum and plasma samples (associations for other metabolic traits are given in Figure 3 and 4). Mean differences in absolute units are listed in sTables 4 and 5.*
***Abbreviations**: **VLDL**=very-low-density lipoprotein.*

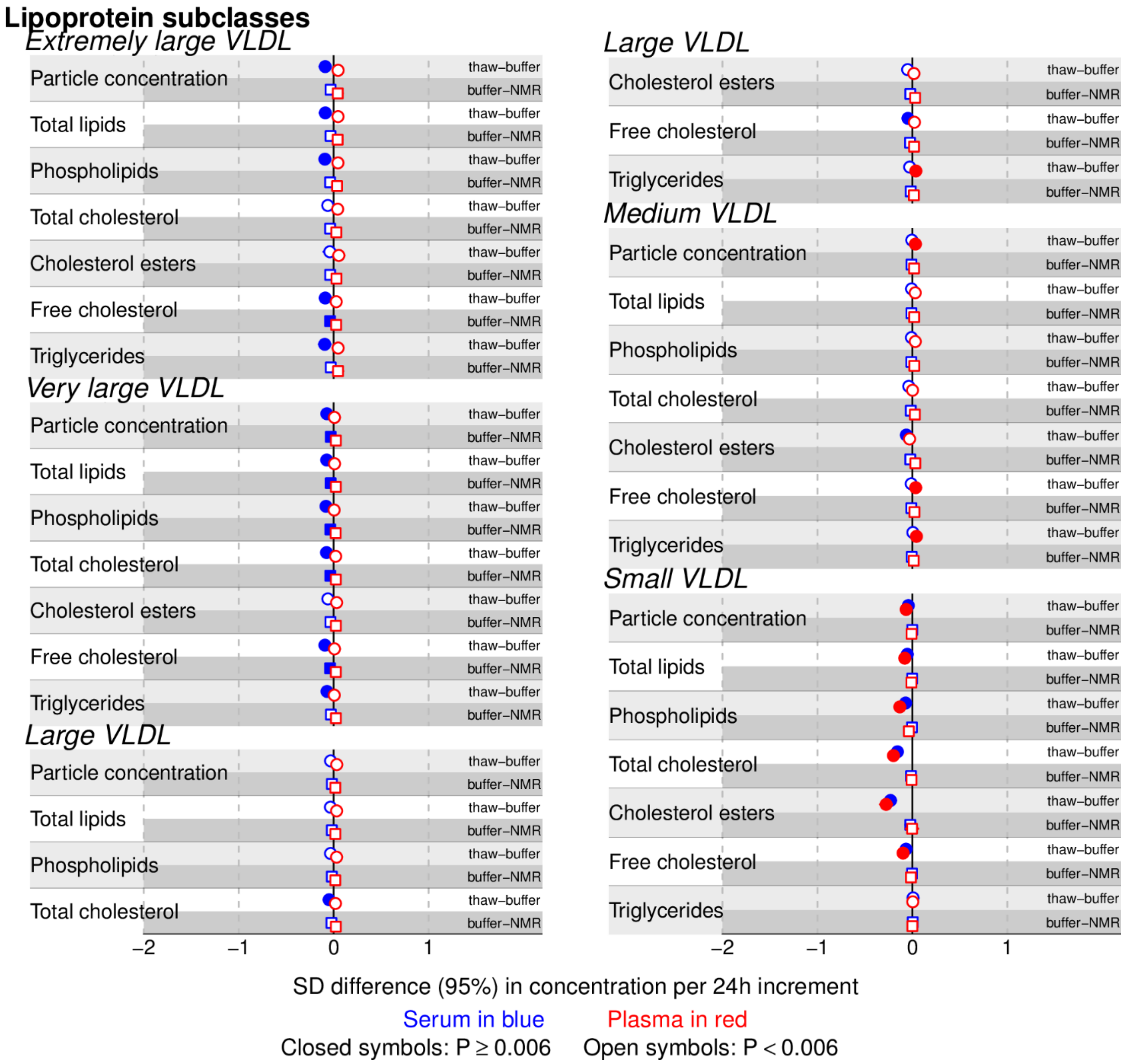

*sFigure 3 (continued). Post -storage handling effects: standardized mean differences in lipoprotein particle and lipid concentration comparing delays in sample preparation (i.e. thaw to buffer addition delay) and NMR profiling (buffer addition to NMR profiling delay) to the reference, for serum and plasma samples (associations for other metabolic traits are given in Figure 3 and 4). Mean differences in absolute units are listed in sTables 4 and 5.*

***Abbreviations**: **IDL**=intermediate-density lipoprotein; **LDL**=low-density lipoprotein; **VLDL**=very-low-density lipoprotein.*

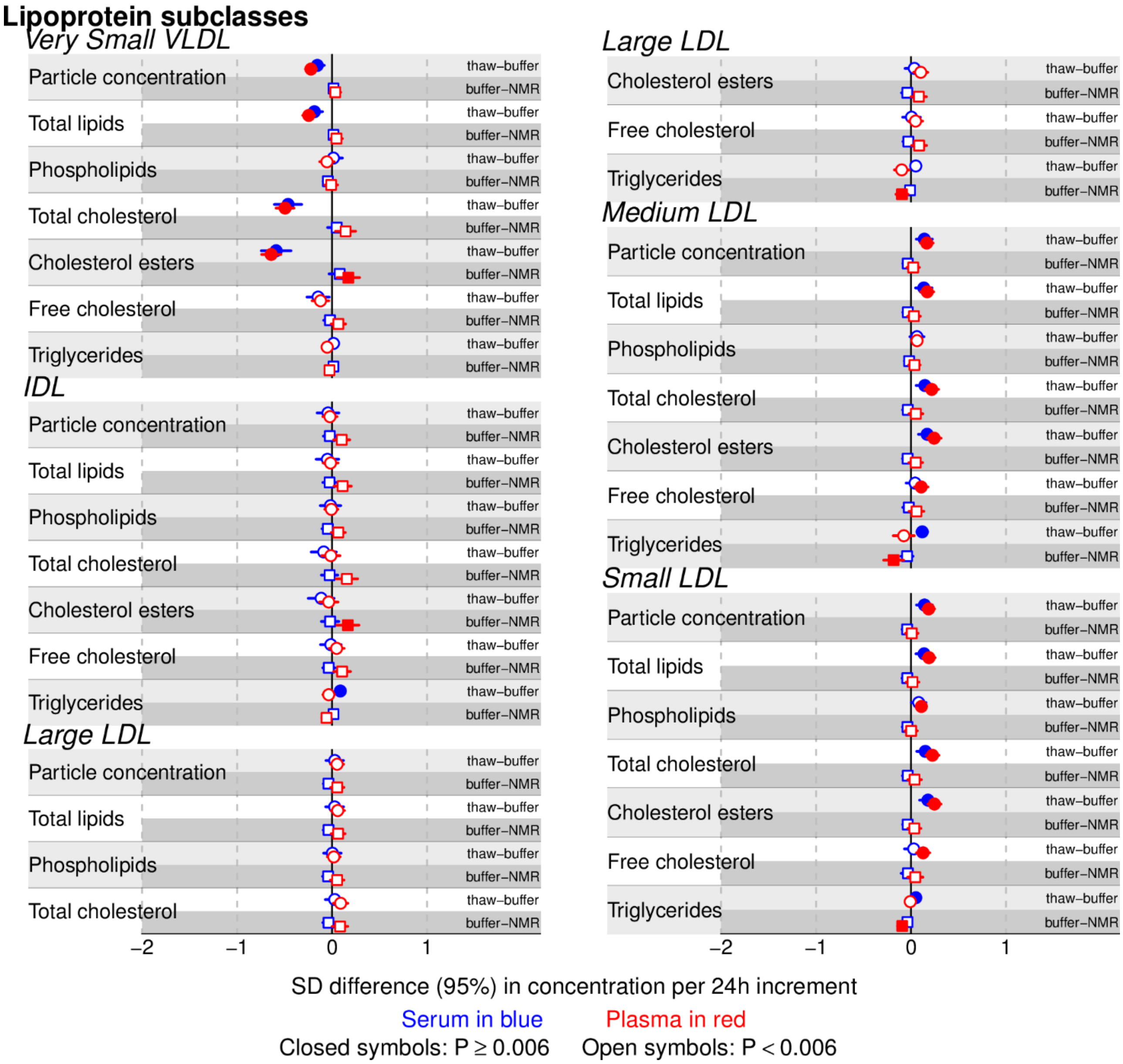

*sFigure 3 (continued). Post -storage handling effects: standardized mean differences in lipoprotein particle and lipid concentration comparing delays in sample preparation (i.e. thaw to buffer addition delay) and NMR profiling (buffer addition to NMR profiling delay) to the reference, for serum and plasma samples (associations for other metabolic traits are given in Figure 3 and 4). Mean differences in absolute units are listed in sTables 4 and 5.*

***Abbreviations**: **HDL**=high-density lipoprotein.*

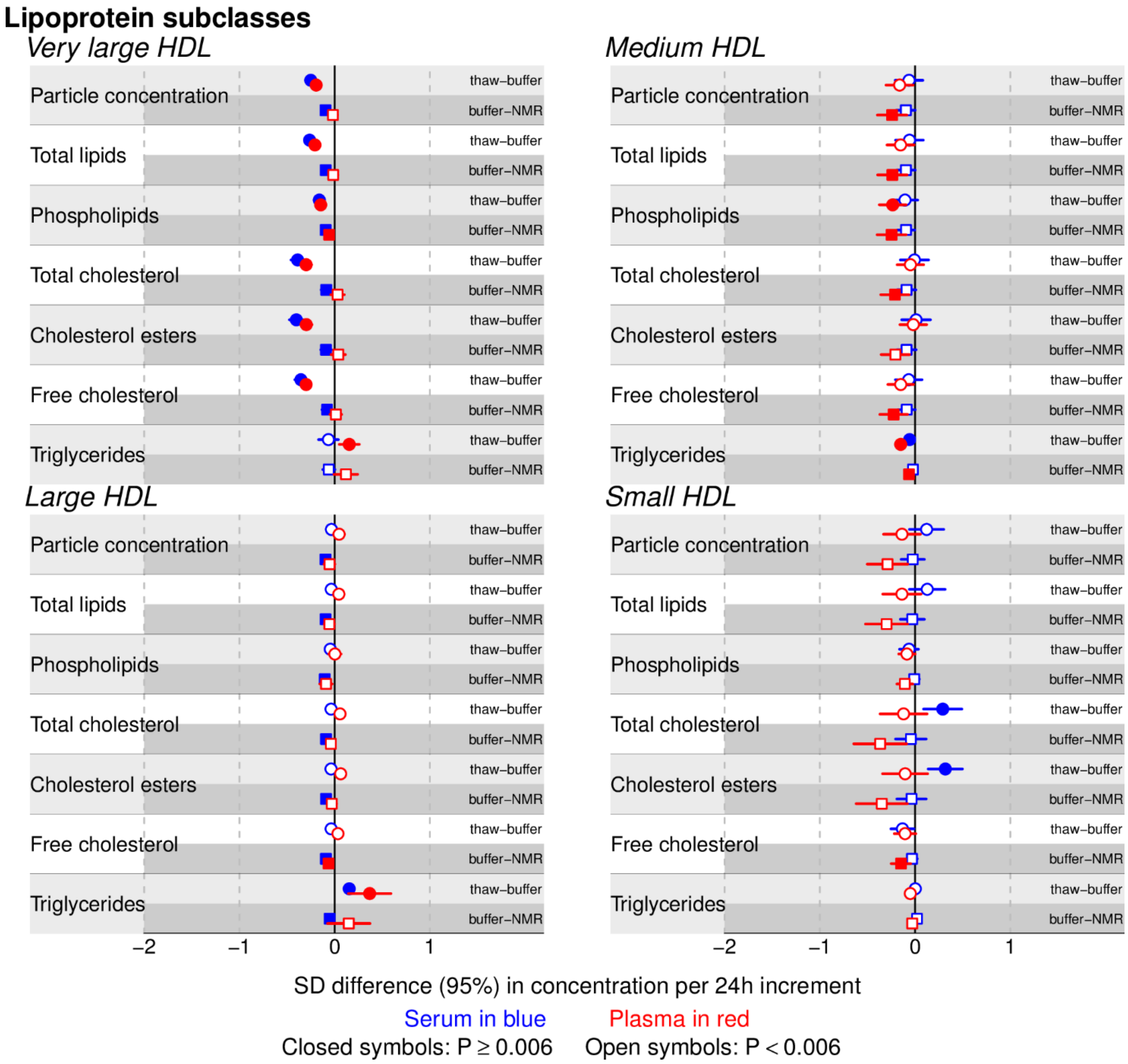

*sFigure 4-Principal Component Analysis (PCA) on serum non-lipid-related metabolic traits subjected to five different pre-storage conditions. PCA was done on subsetted metabolic profiles which only include the 19 (out of 151) non-lipid-related metabolic traits. Principal component (PC) 2 versus PC1. PCA is an exploratory method as it does not focus on finding differences between pre-storage conditions but to explain as much of the variation as possible of the metabolomic profiles with a few new variables (i.e. PCs).[4]* ***[A] Correlation circle****: each metabolic trait is shown as a vector. For the displayed PCs[4], angles between metabolite vectors indicate their degree of correlation. Positively correlated metabolic traits are grouped together (≈ 0° angle); negatively correlated ones are positioned on opposite sides of the plot origin (≈ 180° degrees); a 90° angle between two metabolites indicate that they are uncorrelated. Dashed grey circles represent 100%, 50% and 25% explained variance of a metabolic trait by a PC. Metabolic traits, whose variance is explained by less than 50% by a PC are usually considered nonrelevant for that PC.* ***[B] Scores plot with ellipses enclosing samples from the same participant:*** *each participant is shown as a number and is represented by 5 data-points corresponding to each pre-storage condition; each pre-storage condition is colour coded. Clustering of samples indicates their compositional similarity for the metabolic traits described by the PCs displayed.[4] Samples from the same participant are enclosed by dotted lines.* ***[C] Scores plot with ellipses enclosing samples subjected to the same pre-storage conditions:*** *same as [B] but with samples subjected by the same pre-storage handling conditions enclosed by dotted lines. Ellipses are colour coded by pre-storage conditions.*

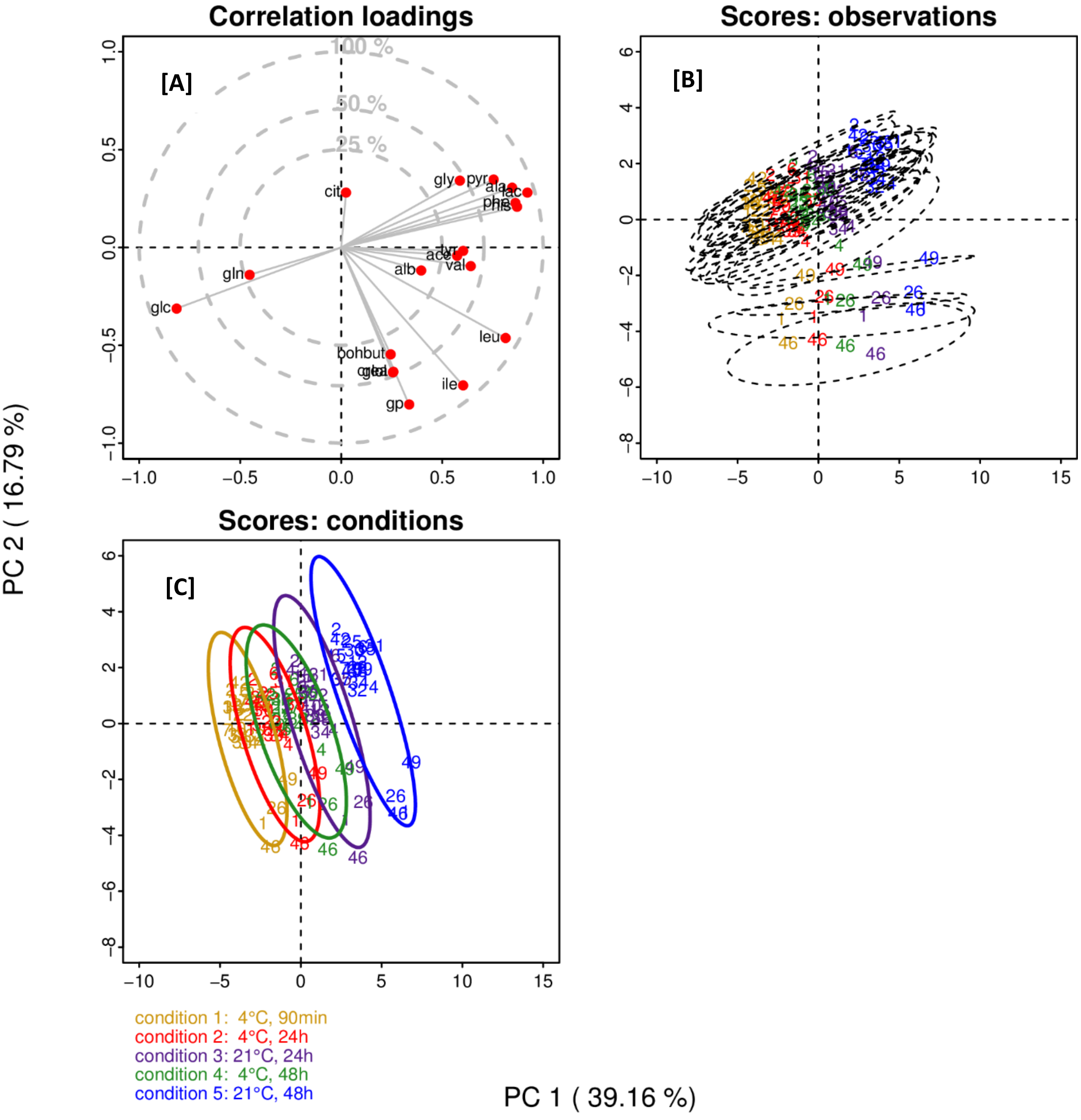

***PCA on non-lipid-related metabolic traits**: PC1 and PC2, which explains 55.95 % of the total variance of the non-lipid-related metabolic traits (19 of the 151 metabolic traits), contains information of both, pre-storage conditions (PC1) and inter-individual differences (PC2). On PC1, from left to right, samples cluster according to increase in incubation temperature first and duration in second (sample clustering order: reference condition ($4^0$C-1.5h); $4^0$C-24h; $4^0$C-48h; $21^0$C-24h; $21^0$C-48h) illustrating the overall sample degradation signature. Moreover, from left (reference condition) to right ($21^0$C-48h), samples have progressive decreasing concentrations of glucose and higher concentrations of lactate, pyruvate, alanine, histidine, and phenylalanine. Glucose is inversely correlated (≈$180^0$ angle) with lactate, pyruvate, histidine, phenylalanine and alanine. Moreover, the latter 4 metabolites are positively correlated amongst themselves (≈ $0^0$ angles). PC1 explains more than 50% of each of these 5 metabolites variance (i.e. correlation between PC1 scores and each metabolite is more than ± 0.71). No further clustering by pre-storage handling conditions at higher PC dimensions.*

*In plasma, the overall degradation fingerprint includes decrease in glucose and glutamine, alongside increased lactate and alanine (data not shown).*

*PCA on the **whole** serum metabolic profiles (i.e. the 151 lipid and non-lipid-related metabolic traits), did not differentiate samples by pre-storage conditions until PCs 4 and 5, which had strong contributions of glycolysis related metabolites and amino-acids (data not shown). In the scores plot of the first two PCs, samples clustered only according to participant, indicating that metabolic profile differences between participants were larger than metabolic concentration differences due to pre-storage conditions. Since Nightingale Health® NMR platform is mostly a lipidomic platform (only 13% of the metabolic traits are non-lipid-related) these traits are over represented in the metabolic profiles, and since these lipid-related traits are mostly robust to pre-storage conditions, the biggest variation in the whole metabolic profiles (and explained by the first few PCs) is associated with inter-individual differences and not differences induced by the pre-storage conditions (which only affects up to 9% of the metabolic traits).*

***Abbreviations**: **ala**=alanine; **alb**=albumin; **ace**=acetate; **acace**=acetoacetate; **bohbut**=beta-hydroxybutyrate; **cit**=citrate; **crea** =creatinine; **glc**=glucose; **lac**=lactate; **pyr**=pyruvate; **glol**=glycerol; **gln**=glutamine; **his**=histidine; **gly**=glycine; **ile**= isoleucine; **leu**= leucine; **val**= valine; **phe**=phenylalanine; **tyr**=tyrosine; **gp**=glycoprotein acetyls.*

#### Supplementary tables

*sTable 1. Characteristics of metabolic traits: metabolic traits concentration (or value) in serum and plasma samples, subjected to the reference pre and post-storage conditions, from individuals who contributed to at least one pair of exposure-outcome analysis (N=37). Pyruvate, glycerol and glycine are not quantified in Ethylenediaminetetraacetic acid (EDTA) - plasma samples due to the interfering resonances of EDTA on their signals.*

***Abbreviations**: **C**=cholesterol; **IDL**=intermediate-density lipoprotein; **iqr25**= 25th percentile; **iqr75**= 75th percentile; **LDL**=low-density lipoprotein; **HDL**=high-density lipoprotein; **MUFA**=monounsaturated fatty acids; **PUFA**=polyunsaturated fatty acids; **sd**= standard deviation; **VLDL**=very-low-density lipoprotein.*

| Metabolic traits | sample | mean | sd | median | iqr25 | iqr75 | min | max |
|---|---|---|---|---|---|---|---|---|
| **Lipoprotein subclasses** | | | | | | | | |
| *Extremely large VLDL* | | | | | | | | |
| Particle concentration (mol/l) | SERUM | 1.2e-10 | 1.9e-10 | 6.7e-11 | 4.4e-11 | 1.3e-10 | 0.0e+00 | 9.7e-10 |
| Particle concentration (mol/l) | PLASMA | 1.1e-10 | 2.1e-10 | 5.3e-11 | 0.0e+00 | 9.0e-11 | 0.0e+00 | 9.2e-10 |
| Total lipids (mmol/l) | SERUM | 2.5e-02 | 4.0e-02 | 1.4e-02 | 9.2e-03 | 2.8e-02 | 0.0e+00 | 2.1e-01 |
| Total lipids (mmol/l) | PLASMA | 2.4e-02 | 4.6e-02 | 1.1e-02 | 0.0e+00 | 1.9e-02 | 0.0e+00 | 2.0e-01 |
| Phospholipids (mmol/l) | SERUM | 3.1e-03 | 5.0e-03 | 1.6e-03 | 8.8e-04 | 3.4e-03 | 0.0e+00 | 2.6e-02 |
| Phospholipids (mmol/l) | PLASMA | 3.0e-03 | 5.7e-03 | 1.3e-03 | 0.0e+00 | 2.3e-03 | 0.0e+00 | 2.5e-02 |
| Total cholesterol (mmol/l) | SERUM | 4.3e-03 | 7.4e-03 | 2.3e-03 | 8.8e-04 | 4.6e-03 | 0.0e+00 | 3.9e-02 |
| Total cholesterol (mmol/l) | PLASMA | 4.2e-03 | 8.6e-03 | 1.3e-03 | 0.0e+00 | 3.3e-03 | 0.0e+00 | 3.7e-02 |
| Cholesterol esters (mmol/l) | SERUM | 2.4e-03 | 4.2e-03 | 1.1e-03 | 2.2e-04 | 2.3e-03 | 0.0e+00 | 2.2e-02 |
| Cholesterol esters (mmol/l) | PLASMA | 2.3e-03 | 4.8e-03 | 3.6e-04 | 0.0e+00 | 2.0e-03 | 0.0e+00 | 2.1e-02 |
| Free cholesterol (mmol/l) | SERUM | 1.9e-03 | 3.3e-03 | 9.6e-04 | 3.3e-04 | 2.1e-03 | 0.0e+00 | 1.7e-02 |
| Free cholesterol (mmol/l) | PLASMA | 1.9e-03 | 3.8e-03 | 6.6e-04 | 0.0e+00 | 1.3e-03 | 0.0e+00 | 1.6e-02 |
| Triglycerides (mmol/l) | SERUM | 1.8e-02 | 2.8e-02 | 1.0e-02 | 7.2e-03 | 2.0e-02 | 0.0e+00 | 1.4e-01 |
| Triglycerides (mmol/l) | PLASMA | 1.7e-02 | 3.2e-02 | 8.4e-03 | 0.0e+00 | 1.4e-02 | 0.0e+00 | 1.4e-01 |
| *Very large VLDL* | | | | | | | | |
| Particle concentration (mol/l) | SERUM | 6.4e-10 | 1.1e-09 | 3.1e-10 | 7.0e-11 | 7.7e-10 | 0.0e+00 | 5.6e-09 |
| Particle concentration (mol/l) | PLASMA | 6.8e-10 | 1.3e-09 | 2.8e-10 | 0.0e+00 | 6.4e-10 | 0.0e+00 | 5.5e-09 |
| Total lipids (mmol/l) | SERUM | 6.2e-02 | 1.1e-01 | 3.0e-02 | 6.6e-03 | 7.5e-02 | 0.0e+00 | 5.5e-01 |
| Total lipids (mmol/l) | PLASMA | 6.6e-02 | 1.3e-01 | 2.6e-02 | 0.0e+00 | 6.1e-02 | 0.0e+00 | 5.3e-01 |

| Metabolic traits | sample | mean | sd | median | iqr25 | iqr75 | min | max |
|---|---|---|---|---|---|---|---|---|
| Phospholipids (mmol/l) | SERUM | 1.0e-02 | 1.8e-02 | 4.8e-03 | 1.4e-03 | 1.2e-02 | 0.0e+00 | 9.2e-02 |
| Phospholipids (mmol/l) | PLASMA | 1.1e-02 | 2.1e-02 | 3.2e-03 | 0.0e+00 | 9.0e-03 | 0.0e+00 | 8.9e-02 |
| Total cholesterol (mmol/l) | SERUM | 1.2e-02 | 2.2e-02 | 5.4e-03 | 5.2e-04 | 1.5e-02 | 0.0e+00 | 1.1e-01 |
| Total cholesterol (mmol/l) | PLASMA | 1.2e-02 | 2.5e-02 | 3.5e-03 | 0.0e+00 | 8.6e-03 | 0.0e+00 | 1.1e-01 |
| Cholesterol esters (mmol/l) | SERUM | 6.6e-03 | 1.2e-02 | 3.1e-03 | 3.0e-06 | 7.6e-03 | 0.0e+00 | 6.2e-02 |
| Cholesterol esters (mmol/l) | PLASMA | 6.7e-03 | 1.4e-02 | 1.9e-03 | 0.0e+00 | 5.0e-03 | 0.0e+00 | 6.0e-02 |
| Free cholesterol (mmol/l) | SERUM | 5.4e-03 | 1.0e-02 | 2.1e-03 | 5.2e-04 | 6.3e-03 | 0.0e+00 | 5.2e-02 |
| Free cholesterol (mmol/l) | PLASMA | 5.6e-03 | 1.2e-02 | 1.7e-03 | 0.0e+00 | 4.2e-03 | 0.0e+00 | 5.0e-02 |
| Triglycerides (mmol/l) | SERUM | 4.0e-02 | 6.9e-02 | 2.0e-02 | 4.3e-03 | 4.8e-02 | 0.0e+00 | 3.5e-01 |
| Triglycerides (mmol/l) | PLASMA | 4.3e-02 | 7.9e-02 | 2.0e-02 | 0.0e+00 | 4.2e-02 | 0.0e+00 | 3.4e-01 |
| *Large VLDL* | | | | | | | | |
| Particle concentration (mol/l) | SERUM | 4.4e-09 | 6.3e-09 | 2.4e-09 | 1.3e-09 | 5.1e-09 | 0.0e+00 | 3.2e-08 |
| Particle concentration (mol/l) | PLASMA | 4.8e-09 | 7.3e-09 | 2.2e-09 | 1.6e-09 | 4.7e-09 | 0.0e+00 | 3.2e-08 |
| Total lipids (mmol/l) | SERUM | 2.5e-01 | 3.7e-01 | 1.4e-01 | 7.0e-02 | 3.0e-01 | 0.0e+00 | 1.9e+00 |
| Total lipids (mmol/l) | PLASMA | 2.8e-01 | 4.2e-01 | 1.2e-01 | 8.8e-02 | 2.7e-01 | 0.0e+00 | 1.8e+00 |
| Phospholipids (mmol/l) | SERUM | 4.6e-02 | 6.6e-02 | 2.6e-02 | 1.3e-02 | 5.5e-02 | 0.0e+00 | 3.4e-01 |
| Phospholipids (mmol/l) | PLASMA | 5.1e-02 | 7.6e-02 | 2.3e-02 | 1.6e-02 | 5.1e-02 | 0.0e+00 | 3.3e-01 |
| Total cholesterol (mmol/l) | SERUM | 5.5e-02 | 8.5e-02 | 2.7e-02 | 1.1e-02 | 6.6e-02 | 0.0e+00 | 4.4e-01 |
| Total cholesterol (mmol/l) | PLASMA | 5.8e-02 | 9.8e-02 | 2.4e-02 | 1.1e-02 | 5.3e-02 | 0.0e+00 | 4.2e-01 |
| Cholesterol esters (mmol/l) | SERUM | 2.9e-02 | 4.2e-02 | 1.6e-02 | 7.8e-03 | 3.2e-02 | 0.0e+00 | 2.2e-01 |
| Cholesterol esters (mmol/l) | PLASMA | 3.0e-02 | 4.9e-02 | 1.3e-02 | 7.2e-03 | 2.7e-02 | 0.0e+00 | 2.1e-01 |
| Free cholesterol (mmol/l) | SERUM | 2.6e-02 | 4.3e-02 | 1.3e-02 | 3.1e-03 | 3.1e-02 | 0.0e+00 | 2.2e-01 |
| Free cholesterol (mmol/l) | PLASMA | 2.8e-02 | 4.9e-02 | 1.1e-02 | 4.7e-03 | 2.7e-02 | 0.0e+00 | 2.1e-01 |
| Triglycerides (mmol/l) | SERUM | 1.5e-01 | 2.2e-01 | 8.6e-02 | 4.7e-02 | 1.7e-01 | 0.0e+00 | 1.1e+00 |
| Triglycerides (mmol/l) | PLASMA | 1.7e-01 | 2.5e-01 | 8.2e-02 | 5.6e-02 | 1.7e-01 | 0.0e+00 | 1.1e+00 |
| *Medium VLDL* | | | | | | | | |
| Particle concentration (mol/l) | SERUM | 1.5e-08 | 1.6e-08 | 9.5e-09 | 7.8e-09 | 1.6e-08 | 2.5e-09 | 8.3e-08 |
| Particle concentration (mol/l) | PLASMA | 1.6e-08 | 1.8e-08 | 9.5e-09 | 8.5e-09 | 1.5e-08 | 3.3e-09 | 8.3e-08 |
| Total lipids (mmol/l) | SERUM | 5.1e-01 | 5.2e-01 | 3.2e-01 | 2.6e-01 | 5.2e-01 | 8.6e-02 | 2.8e+00 |

| Metabolic traits | sample | mean | sd | median | iqr25 | iqr75 | min | max |
|---|---|---|---|---|---|---|---|---|
| Total lipids (mmol/l) | PLASMA | 5.5e-01 | 6.0e-01 | 3.1e-01 | 2.8e-01 | 5.1e-01 | 1.1e-01 | 2.7e+00 |
| Phospholipids (mmol/l) | SERUM | 1.0e-01 | 1.0e-01 | 6.7e-02 | 5.5e-02 | 1.1e-01 | 2.1e-02 | 5.3e-01 |
| Phospholipids (mmol/l) | PLASMA | 1.1e-01 | 1.1e-01 | 6.3e-02 | 5.9e-02 | 1.0e-01 | 2.5e-02 | 5.3e-01 |
| Total cholesterol (mmol/l) | SERUM | 1.4e-01 | 1.3e-01 | 9.3e-02 | 7.1e-02 | 1.4e-01 | 2.6e-02 | 7.1e-01 |
| Total cholesterol (mmol/l) | PLASMA | 1.4e-01 | 1.5e-01 | 8.1e-02 | 6.4e-02 | 1.4e-01 | 3.1e-02 | 7.0e-01 |
| Cholesterol esters (mmol/l) | SERUM | 7.8e-02 | 6.6e-02 | 5.7e-02 | 4.8e-02 | 8.5e-02 | 1.8e-02 | 3.7e-01 |
| Cholesterol esters (mmol/l) | PLASMA | 7.6e-02 | 7.7e-02 | 4.7e-02 | 3.7e-02 | 8.6e-02 | 2.1e-02 | 3.6e-01 |
| Free cholesterol (mmol/l) | SERUM | 5.8e-02 | 6.5e-02 | 3.8e-02 | 2.5e-02 | 6.7e-02 | 6.6e-03 | 3.4e-01 |
| Free cholesterol (mmol/l) | PLASMA | 6.2e-02 | 7.5e-02 | 3.4e-02 | 2.9e-02 | 6.0e-02 | 1.0e-02 | 3.4e-01 |
| Triglycerides (mmol/l) | SERUM | 2.7e-01 | 2.9e-01 | 1.6e-01 | 1.3e-01 | 2.9e-01 | 3.6e-02 | 1.5e+00 |
| Triglycerides (mmol/l) | PLASMA | 3.0e-01 | 3.4e-01 | 1.8e-01 | 1.4e-01 | 2.9e-01 | 5.3e-02 | 1.5e+00 |
| *Small VLDL* | | | | | | | | |
| Particle concentration (mol/l) | SERUM | 2.5e-08 | 1.5e-08 | 1.9e-08 | 1.7e-08 | 2.7e-08 | 9.5e-09 | 8.7e-08 |
| Particle concentration (mol/l) | PLASMA | 2.6e-08 | 1.7e-08 | 2.0e-08 | 1.8e-08 | 2.7e-08 | 1.1e-08 | 8.7e-08 |
| Total lipids (mmol/l) | SERUM | 4.8e-01 | 2.9e-01 | 3.8e-01 | 3.4e-01 | 5.3e-01 | 1.8e-01 | 1.7e+00 |
| Total lipids (mmol/l) | PLASMA | 5.1e-01 | 3.3e-01 | 4.0e-01 | 3.3e-01 | 5.3e-01 | 2.1e-01 | 1.7e+00 |
| Phospholipids (mmol/l) | SERUM | 1.2e-01 | 5.7e-02 | 9.7e-02 | 8.4e-02 | 1.3e-01 | 5.5e-02 | 3.5e-01 |
| Phospholipids (mmol/l) | PLASMA | 1.2e-01 | 6.5e-02 | 1.0e-01 | 8.7e-02 | 1.3e-01 | 6.2e-02 | 3.5e-01 |
| Total cholesterol (mmol/l) | SERUM | 1.6e-01 | 7.9e-02 | 1.4e-01 | 1.2e-01 | 1.8e-01 | 4.6e-02 | 4.5e-01 |
| Total cholesterol (mmol/l) | PLASMA | 1.6e-01 | 9.1e-02 | 1.3e-01 | 1.0e-01 | 1.7e-01 | 6.0e-02 | 4.5e-01 |
| Cholesterol esters (mmol/l) | SERUM | 9.6e-02 | 4.4e-02 | 8.3e-02 | 7.1e-02 | 1.1e-01 | 2.1e-02 | 2.4e-01 |
| Cholesterol esters (mmol/l) | PLASMA | 8.7e-02 | 5.0e-02 | 7.3e-02 | 5.5e-02 | 9.6e-02 | 2.8e-02 | 2.3e-01 |
| Free cholesterol (mmol/l) | SERUM | 6.7e-02 | 3.7e-02 | 5.4e-02 | 4.6e-02 | 7.5e-02 | 2.6e-02 | 2.1e-01 |
| Free cholesterol (mmol/l) | PLASMA | 7.1e-02 | 4.2e-02 | 5.7e-02 | 4.8e-02 | 7.5e-02 | 3.2e-02 | 2.1e-01 |
| Triglycerides (mmol/l) | SERUM | 2.0e-01 | 1.5e-01 | 1.5e-01 | 1.3e-01 | 2.1e-01 | 7.1e-02 | 8.5e-01 |
| Triglycerides (mmol/l) | PLASMA | 2.3e-01 | 1.7e-01 | 1.7e-01 | 1.4e-01 | 2.3e-01 | 8.9e-02 | 8.6e-01 |
| *Very Small VLDL* | | | | | | | | |
| Particle concentration (mol/l) | SERUM | 3.1e-08 | 8.4e-09 | 3.0e-08 | 2.7e-08 | 3.4e-08 | 1.6e-08 | 5.5e-08 |
| Particle concentration (mol/l) | PLASMA | 3.0e-08 | 9.4e-09 | 2.7e-08 | 2.5e-08 | 3.2e-08 | 1.7e-08 | 5.5e-08 |

| Metabolic traits | sample | mean | sd | median | iqr25 | iqr75 | min | max |
|---|---|---|---|---|---|---|---|---|
| Total lipids (mmol/l) | SERUM | 4.0e-01 | 1.0e-01 | 3.8e-01 | 3.4e-01 | 4.3e-01 | 2.0e-01 | 6.4e-01 |
| Total lipids (mmol/l) | PLASMA | 3.7e-01 | 1.1e-01 | 3.4e-01 | 3.1e-01 | 4.1e-01 | 2.2e-01 | 6.4e-01 |
| Phospholipids (mmol/l) | SERUM | 1.3e-01 | 2.9e-02 | 1.3e-01 | 1.1e-01 | 1.5e-01 | 7.1e-02 | 2.1e-01 |
| Phospholipids (mmol/l) | PLASMA | 1.2e-01 | 3.0e-02 | 1.2e-01 | 1.0e-01 | 1.4e-01 | 6.9e-02 | 2.0e-01 |
| Total cholesterol (mmol/l) | SERUM | 1.7e-01 | 4.4e-02 | 1.7e-01 | 1.5e-01 | 2.0e-01 | 8.3e-02 | 2.9e-01 |
| Total cholesterol (mmol/l) | PLASMA | 1.5e-01 | 4.5e-02 | 1.4e-01 | 1.2e-01 | 1.8e-01 | 9.4e-02 | 2.6e-01 |
| Cholesterol esters (mmol/l) | SERUM | 1.1e-01 | 3.1e-02 | 1.1e-01 | 9.0e-02 | 1.3e-01 | 4.6e-02 | 1.9e-01 |
| Cholesterol esters (mmol/l) | PLASMA | 9.5e-02 | 3.1e-02 | 8.6e-02 | 7.6e-02 | 1.1e-01 | 5.6e-02 | 1.7e-01 |
| Free cholesterol (mmol/l) | SERUM | 6.4e-02 | 1.4e-02 | 6.2e-02 | 5.6e-02 | 7.2e-02 | 3.3e-02 | 1.0e-01 |
| Free cholesterol (mmol/l) | PLASMA | 5.7e-02 | 1.4e-02 | 5.5e-02 | 4.6e-02 | 6.8e-02 | 3.6e-02 | 9.4e-02 |
| Triglycerides (mmol/l) | SERUM | 9.1e-02 | 4.2e-02 | 7.9e-02 | 6.8e-02 | 9.3e-02 | 4.6e-02 | 2.7e-01 |
| Triglycerides (mmol/l) | PLASMA | 9.8e-02 | 4.8e-02 | 8.1e-02 | 7.3e-02 | 1.0e-01 | 5.1e-02 | 2.7e-01 |
| *IDL* | | | | | | | | |
| Particle concentration (mol/l) | SERUM | 9.4e-08 | 1.9e-08 | 9.3e-08 | 8.0e-08 | 1.1e-07 | 5.2e-08 | 1.4e-07 |
| Particle concentration (mol/l) | PLASMA | 8.6e-08 | 2.0e-08 | 8.7e-08 | 7.1e-08 | 1.0e-07 | 5.5e-08 | 1.3e-07 |
| Total lipids (mmol/l) | SERUM | 9.5e-01 | 2.0e-01 | 9.5e-01 | 8.1e-01 | 1.1e+00 | 5.3e-01 | 1.4e+00 |
| Total lipids (mmol/l) | PLASMA | 8.6e-01 | 2.0e-01 | 8.7e-01 | 7.2e-01 | 1.0e+00 | 5.6e-01 | 1.3e+00 |
| Phospholipids (mmol/l) | SERUM | 2.6e-01 | 5.0e-02 | 2.6e-01 | 2.3e-01 | 3.0e-01 | 1.7e-01 | 3.8e-01 |
| Phospholipids (mmol/l) | PLASMA | 2.4e-01 | 5.0e-02 | 2.4e-01 | 2.1e-01 | 2.7e-01 | 1.6e-01 | 3.5e-01 |
| Total cholesterol (mmol/l) | SERUM | 5.9e-01 | 1.3e-01 | 5.9e-01 | 5.0e-01 | 6.6e-01 | 3.1e-01 | 9.0e-01 |
| Total cholesterol (mmol/l) | PLASMA | 5.2e-01 | 1.4e-01 | 5.1e-01 | 4.1e-01 | 6.3e-01 | 3.2e-01 | 7.8e-01 |
| Cholesterol esters (mmol/l) | SERUM | 4.2e-01 | 9.7e-02 | 4.2e-01 | 3.5e-01 | 4.8e-01 | 2.1e-01 | 6.4e-01 |
| Cholesterol esters (mmol/l) | PLASMA | 3.6e-01 | 1.1e-01 | 3.5e-01 | 2.8e-01 | 4.6e-01 | 2.2e-01 | 5.4e-01 |
| Free cholesterol (mmol/l) | SERUM | 1.7e-01 | 4.1e-02 | 1.7e-01 | 1.5e-01 | 2.0e-01 | 1.0e-01 | 2.7e-01 |
| Free cholesterol (mmol/l) | PLASMA | 1.6e-01 | 4.0e-02 | 1.5e-01 | 1.3e-01 | 1.7e-01 | 1.0e-01 | 2.4e-01 |
| Triglycerides (mmol/l) | SERUM | 9.7e-02 | 2.8e-02 | 9.3e-02 | 7.8e-02 | 1.1e-01 | 5.0e-02 | 2.0e-01 |
| Triglycerides (mmol/l) | PLASMA | 1.0e-01 | 3.2e-02 | 9.7e-02 | 8.2e-02 | 1.1e-01 | 5.2e-02 | 2.0e-01 |
| *Large LDL* | | | | | | | | |
| Particle concentration (mol/l) | SERUM | 1.6e-07 | 3.4e-08 | 1.5e-07 | 1.3e-07 | 1.8e-07 | 9.2e-08 | 2.3e-07 |

| Metabolic traits | sample | mean | sd | median | iqr25 | iqr75 | min | max |
|---|---|---|---|---|---|---|---|---|
| Particle concentration (mol/l) | PLASMA | 1.5e-07 | 3.5e-08 | 1.4e-07 | 1.2e-07 | 1.7e-07 | 9.0e-08 | 2.2e-07 |
| Total lipids (mmol/l) | SERUM | 1.1e+00 | 2.4e-01 | 1.1e+00 | 9.5e-01 | 1.3e+00 | 6.5e-01 | 1.7e+00 |
| Total lipids (mmol/l) | PLASMA | 1.1e+00 | 2.5e-01 | 1.0e+00 | 8.7e-01 | 1.2e+00 | 6.4e-01 | 1.5e+00 |
| Phospholipids (mmol/l) | SERUM | 2.9e-01 | 5.1e-02 | 2.8e-01 | 2.5e-01 | 3.2e-01 | 1.9e-01 | 4.1e-01 |
| Phospholipids (mmol/l) | PLASMA | 2.7e-01 | 5.2e-02 | 2.7e-01 | 2.3e-01 | 3.1e-01 | 1.8e-01 | 3.7e-01 |
| Total cholesterol (mmol/l) | SERUM | 7.5e-01 | 1.8e-01 | 7.2e-01 | 6.4e-01 | 8.8e-01 | 4.1e-01 | 1.2e+00 |
| Total cholesterol (mmol/l) | PLASMA | 6.9e-01 | 1.8e-01 | 6.6e-01 | 5.6e-01 | 8.2e-01 | 4.0e-01 | 1.1e+00 |
| Cholesterol esters (mmol/l) | SERUM | 5.4e-01 | 1.4e-01 | 5.1e-01 | 4.5e-01 | 6.2e-01 | 2.7e-01 | 8.6e-01 |
| Cholesterol esters (mmol/l) | PLASMA | 4.9e-01 | 1.4e-01 | 4.6e-01 | 3.7e-01 | 5.8e-01 | 2.6e-01 | 7.6e-01 |
| Free cholesterol (mmol/l) | SERUM | 2.2e-01 | 4.6e-02 | 2.1e-01 | 1.8e-01 | 2.5e-01 | 1.4e-01 | 3.2e-01 |
| Free cholesterol (mmol/l) | PLASMA | 2.0e-01 | 4.5e-02 | 2.0e-01 | 1.7e-01 | 2.1e-01 | 1.3e-01 | 2.9e-01 |
| Triglycerides (mmol/l) | SERUM | 8.7e-02 | 2.3e-02 | 9.0e-02 | 6.7e-02 | 1.0e-01 | 4.2e-02 | 1.5e-01 |
| Triglycerides (mmol/l) | PLASMA | 9.1e-02 | 2.6e-02 | 9.2e-02 | 6.9e-02 | 1.0e-01 | 4.2e-02 | 1.5e-01 |
| *Medium LDL* | | | | | | | | |
| Particle concentration (mol/l) | SERUM | 1.3e-07 | 3.0e-08 | 1.2e-07 | 1.1e-07 | 1.5e-07 | 6.9e-08 | 2.0e-07 |
| Particle concentration (mol/l) | PLASMA | 1.2e-07 | 3.0e-08 | 1.1e-07 | 9.8e-08 | 1.4e-07 | 6.6e-08 | 1.8e-07 |
| Total lipids (mmol/l) | SERUM | 6.5e-01 | 1.5e-01 | 6.2e-01 | 5.5e-01 | 7.5e-01 | 3.5e-01 | 1.0e+00 |
| Total lipids (mmol/l) | PLASMA | 6.1e-01 | 1.5e-01 | 5.8e-01 | 4.9e-01 | 7.2e-01 | 3.4e-01 | 9.0e-01 |
| Phospholipids (mmol/l) | SERUM | 1.8e-01 | 3.3e-02 | 1.7e-01 | 1.6e-01 | 2.0e-01 | 1.2e-01 | 2.4e-01 |
| Phospholipids (mmol/l) | PLASMA | 1.7e-01 | 3.6e-02 | 1.6e-01 | 1.4e-01 | 1.9e-01 | 1.1e-01 | 2.4e-01 |
| Total cholesterol (mmol/l) | SERUM | 4.3e-01 | 1.1e-01 | 4.0e-01 | 3.7e-01 | 5.0e-01 | 2.1e-01 | 7.1e-01 |
| Total cholesterol (mmol/l) | PLASMA | 3.9e-01 | 1.1e-01 | 3.5e-01 | 3.2e-01 | 4.7e-01 | 1.9e-01 | 6.3e-01 |
| Cholesterol esters (mmol/l) | SERUM | 3.1e-01 | 9.1e-02 | 2.7e-01 | 2.6e-01 | 3.6e-01 | 1.2e-01 | 5.3e-01 |
| Cholesterol esters (mmol/l) | PLASMA | 2.7e-01 | 9.3e-02 | 2.4e-01 | 2.2e-01 | 3.4e-01 | 1.2e-01 | 4.7e-01 |
| Free cholesterol (mmol/l) | SERUM | 1.3e-01 | 2.2e-02 | 1.2e-01 | 1.1e-01 | 1.4e-01 | 8.5e-02 | 1.8e-01 |
| Free cholesterol (mmol/l) | PLASMA | 1.2e-01 | 2.2e-02 | 1.1e-01 | 1.0e-01 | 1.3e-01 | 7.7e-02 | 1.6e-01 |
| Triglycerides (mmol/l) | SERUM | 4.5e-02 | 1.2e-02 | 4.6e-02 | 3.4e-02 | 5.2e-02 | 2.5e-02 | 7.6e-02 |
| Triglycerides (mmol/l) | PLASMA | 4.8e-02 | 1.3e-02 | 4.8e-02 | 3.8e-02 | 5.5e-02 | 2.2e-02 | 7.5e-02 |
| *Small LDL* | | | | | | | | |

| Metabolic traits | sample | mean | sd | median | iqr25 | iqr75 | min | max |
|---|---|---|---|---|---|---|---|---|
| Particle concentration (mol/l) | SERUM | 1.5e-07 | 3.4e-08 | 1.4e-07 | 1.3e-07 | 1.8e-07 | 8.6e-08 | 2.3e-07 |
| Particle concentration (mol/l) | PLASMA | 1.4e-07 | 3.5e-08 | 1.3e-07 | 1.1e-07 | 1.7e-07 | 7.6e-08 | 2.1e-07 |
| Total lipids (mmol/l) | SERUM | 4.3e-01 | 9.6e-02 | 4.0e-01 | 3.6e-01 | 4.8e-01 | 2.4e-01 | 6.4e-01 |
| Total lipids (mmol/l) | PLASMA | 3.9e-01 | 9.8e-02 | 3.7e-01 | 3.2e-01 | 4.7e-01 | 2.1e-01 | 5.8e-01 |
| Phospholipids (mmol/l) | SERUM | 1.3e-01 | 2.3e-02 | 1.3e-01 | 1.2e-01 | 1.5e-01 | 9.3e-02 | 1.8e-01 |
| Phospholipids (mmol/l) | PLASMA | 1.3e-01 | 2.5e-02 | 1.2e-01 | 1.1e-01 | 1.4e-01 | 8.1e-02 | 1.7e-01 |
| Total cholesterol (mmol/l) | SERUM | 2.6e-01 | 7.0e-02 | 2.4e-01 | 2.2e-01 | 3.2e-01 | 1.3e-01 | 4.4e-01 |
| Total cholesterol (mmol/l) | PLASMA | 2.4e-01 | 7.0e-02 | 2.2e-01 | 1.9e-01 | 2.9e-01 | 1.1e-01 | 3.8e-01 |
| Cholesterol esters (mmol/l) | SERUM | 1.9e-01 | 5.6e-02 | 1.7e-01 | 1.5e-01 | 2.2e-01 | 8.0e-02 | 3.3e-01 |
| Cholesterol esters (mmol/l) | PLASMA | 1.7e-01 | 5.7e-02 | 1.5e-01 | 1.4e-01 | 2.0e-01 | 6.9e-02 | 2.9e-01 |
| Free cholesterol (mmol/l) | SERUM | 7.7e-02 | 1.4e-02 | 7.4e-02 | 6.7e-02 | 8.6e-02 | 5.2e-02 | 1.1e-01 |
| Free cholesterol (mmol/l) | PLASMA | 6.9e-02 | 1.4e-02 | 6.7e-02 | 5.9e-02 | 8.0e-02 | 4.2e-02 | 9.9e-02 |
| Triglycerides (mmol/l) | SERUM | 2.9e-02 | 1.1e-02 | 2.7e-02 | 2.2e-02 | 3.2e-02 | 1.5e-02 | 7.3e-02 |
| Triglycerides (mmol/l) | PLASMA | 3.1e-02 | 1.2e-02 | 2.9e-02 | 2.4e-02 | 3.3e-02 | 1.3e-02 | 7.1e-02 |
| *Very large HDL* | | | | | | | | |
| Particle concentration (mol/l) | SERUM | 5.2e-07 | 2.5e-07 | 4.6e-07 | 3.2e-07 | 7.4e-07 | 1.4e-07 | 9.4e-07 |
| Particle concentration (mol/l) | PLASMA | 5.2e-07 | 2.5e-07 | 4.7e-07 | 3.2e-07 | 7.6e-07 | 1.3e-07 | 9.4e-07 |
| Total lipids (mmol/l) | SERUM | 5.2e-01 | 2.5e-01 | 4.6e-01 | 3.1e-01 | 7.4e-01 | 1.4e-01 | 9.5e-01 |
| Total lipids (mmol/l) | PLASMA | 5.2e-01 | 2.5e-01 | 4.7e-01 | 3.2e-01 | 7.6e-01 | 1.3e-01 | 9.4e-01 |
| Phospholipids (mmol/l) | SERUM | 2.8e-01 | 1.4e-01 | 2.5e-01 | 1.7e-01 | 4.1e-01 | 1.9e-02 | 5.2e-01 |
| Phospholipids (mmol/l) | PLASMA | 2.8e-01 | 1.5e-01 | 2.6e-01 | 2.0e-01 | 4.1e-01 | 1.6e-02 | 5.2e-01 |
| Total cholesterol (mmol/l) | SERUM | 2.3e-01 | 1.1e-01 | 2.0e-01 | 1.3e-01 | 3.1e-01 | 6.0e-02 | 4.2e-01 |
| Total cholesterol (mmol/l) | PLASMA | 2.3e-01 | 1.1e-01 | 2.0e-01 | 1.4e-01 | 3.2e-01 | 7.3e-02 | 4.0e-01 |
| Cholesterol esters (mmol/l) | SERUM | 1.7e-01 | 7.7e-02 | 1.5e-01 | 1.0e-01 | 2.2e-01 | 4.8e-02 | 3.0e-01 |
| Cholesterol esters (mmol/l) | PLASMA | 1.6e-01 | 7.6e-02 | 1.4e-01 | 1.0e-01 | 2.3e-01 | 6.1e-02 | 2.9e-01 |
| Free cholesterol (mmol/l) | SERUM | 6.1e-02 | 3.3e-02 | 5.1e-02 | 3.2e-02 | 8.8e-02 | 1.2e-02 | 1.2e-01 |
| Free cholesterol (mmol/l) | PLASMA | 6.1e-02 | 3.3e-02 | 5.4e-02 | 3.3e-02 | 9.2e-02 | 1.2e-02 | 1.1e-01 |
| Triglycerides (mmol/l) | SERUM | 1.8e-02 | 9.4e-03 | 1.7e-02 | 1.3e-02 | 2.4e-02 | 2.4e-03 | 5.2e-02 |
| Triglycerides (mmol/l) | PLASMA | 1.7e-02 | 1.1e-02 | 1.7e-02 | 8.5e-03 | 2.5e-02 | 7.2e-04 | 4.9e-02 |

| Metabolic traits | sample | mean | sd | median | iqr25 | iqr75 | min | max |
|---|---|---|---|---|---|---|---|---|
| *Large HDL* | | | | | | | | |
| Particle concentration (mol/l) | SERUM | 1.6e-06 | 6.5e-07 | 1.5e-06 | 1.1e-06 | 2.3e-06 | 3.7e-07 | 2.8e-06 |
| Particle concentration (mol/l) | PLASMA | 1.6e-06 | 7.0e-07 | 1.5e-06 | 1.3e-06 | 2.2e-06 | 0.0e+00 | 2.6e-06 |
| Total lipids (mmol/l) | SERUM | 1.0e+00 | 4.2e-01 | 9.5e-01 | 7.0e-01 | 1.4e+00 | 2.2e-01 | 1.8e+00 |
| Total lipids (mmol/l) | PLASMA | 1.0e+00 | 4.5e-01 | 9.2e-01 | 8.0e-01 | 1.4e+00 | 0.0e+00 | 1.7e+00 |
| Phospholipids (mmol/l) | SERUM | 4.9e-01 | 1.8e-01 | 4.6e-01 | 3.5e-01 | 6.5e-01 | 1.0e-01 | 8.3e-01 |
| Phospholipids (mmol/l) | PLASMA | 4.8e-01 | 2.0e-01 | 4.5e-01 | 4.2e-01 | 6.5e-01 | 0.0e+00 | 7.7e-01 |
| Total cholesterol (mmol/l) | SERUM | 5.0e-01 | 2.3e-01 | 4.6e-01 | 3.3e-01 | 7.2e-01 | 7.1e-02 | 8.7e-01 |
| Total cholesterol (mmol/l) | PLASMA | 4.8e-01 | 2.4e-01 | 4.5e-01 | 3.5e-01 | 7.0e-01 | 0.0e+00 | 8.3e-01 |
| Cholesterol esters (mmol/l) | SERUM | 3.8e-01 | 1.7e-01 | 3.6e-01 | 2.6e-01 | 5.6e-01 | 6.8e-02 | 6.7e-01 |
| Cholesterol esters (mmol/l) | PLASMA | 3.7e-01 | 1.8e-01 | 3.5e-01 | 2.7e-01 | 5.4e-01 | 0.0e+00 | 6.3e-01 |
| Free cholesterol (mmol/l) | SERUM | 1.1e-01 | 5.5e-02 | 1.0e-01 | 7.2e-02 | 1.7e-01 | 3.4e-03 | 2.0e-01 |
| Free cholesterol (mmol/l) | PLASMA | 1.1e-01 | 5.7e-02 | 1.0e-01 | 8.0e-02 | 1.6e-01 | 0.0e+00 | 1.9e-01 |
| Triglycerides (mmol/l) | SERUM | 3.6e-02 | 1.4e-02 | 3.2e-02 | 2.4e-02 | 5.1e-02 | 1.6e-02 | 5.7e-02 |
| Triglycerides (mmol/l) | PLASMA | 3.5e-02 | 1.6e-02 | 3.3e-02 | 2.6e-02 | 5.3e-02 | 0.0e+00 | 5.8e-02 |
| *Medium HDL* | | | | | | | | |
| Particle concentration (mol/l) | SERUM | 2.4e-06 | 3.7e-07 | 2.4e-06 | 2.1e-06 | 2.5e-06 | 1.8e-06 | 3.2e-06 |
| Particle concentration (mol/l) | PLASMA | 2.4e-06 | 3.8e-07 | 2.4e-06 | 2.1e-06 | 2.6e-06 | 1.8e-06 | 3.3e-06 |
| Total lipids (mmol/l) | SERUM | 1.0e+00 | 1.6e-01 | 1.0e+00 | 9.1e-01 | 1.1e+00 | 7.4e-01 | 1.4e+00 |
| Total lipids (mmol/l) | PLASMA | 1.0e+00 | 1.7e-01 | 1.0e+00 | 8.8e-01 | 1.1e+00 | 7.6e-01 | 1.4e+00 |
| Phospholipids (mmol/l) | SERUM | 4.6e-01 | 7.4e-02 | 4.5e-01 | 4.2e-01 | 4.9e-01 | 3.4e-01 | 6.3e-01 |
| Phospholipids (mmol/l) | PLASMA | 4.7e-01 | 7.6e-02 | 4.7e-01 | 4.1e-01 | 5.1e-01 | 3.6e-01 | 6.5e-01 |
| Total cholesterol (mmol/l) | SERUM | 5.1e-01 | 9.2e-02 | 5.2e-01 | 4.4e-01 | 5.5e-01 | 3.2e-01 | 7.1e-01 |
| Total cholesterol (mmol/l) | PLASMA | 5.0e-01 | 9.3e-02 | 4.9e-01 | 4.3e-01 | 5.7e-01 | 3.2e-01 | 6.6e-01 |
| Cholesterol esters (mmol/l) | SERUM | 4.1e-01 | 7.2e-02 | 4.2e-01 | 3.6e-01 | 4.5e-01 | 2.7e-01 | 5.7e-01 |
| Cholesterol esters (mmol/l) | PLASMA | 4.0e-01 | 7.2e-02 | 4.0e-01 | 3.5e-01 | 4.6e-01 | 2.7e-01 | 5.3e-01 |
| Free cholesterol (mmol/l) | SERUM | 9.4e-02 | 2.0e-02 | 9.4e-02 | 8.3e-02 | 1.0e-01 | 5.5e-02 | 1.4e-01 |
| Free cholesterol (mmol/l) | PLASMA | 9.4e-02 | 2.1e-02 | 9.4e-02 | 8.0e-02 | 1.1e-01 | 5.6e-02 | 1.4e-01 |
| Triglycerides (mmol/l) | SERUM | 4.4e-02 | 1.6e-02 | 4.1e-02 | 3.3e-02 | 5.1e-02 | 2.7e-02 | 1.0e-01 |

| Metabolic traits | sample | mean | sd | median | iqr25 | iqr75 | min | max |
|---|---|---|---|---|---|---|---|---|
| Triglycerides (mmol/l) | PLASMA | 4.9e-02 | 1.8e-02 | 4.3e-02 | 3.7e-02 | 5.1e-02 | 3.2e-02 | 1.0e-01 |
| *Small HDL* | | | | | | | | |
| Particle concentration (mol/l) | SERUM | 4.9e-06 | 5.0e-07 | 4.9e-06 | 4.6e-06 | 5.2e-06 | 3.9e-06 | 6.4e-06 |
| Particle concentration (mol/l) | PLASMA | 5.0e-06 | 5.7e-07 | 4.9e-06 | 4.7e-06 | 5.4e-06 | 4.1e-06 | 6.3e-06 |
| Total lipids (mmol/l) | SERUM | 1.1e+00 | 1.1e-01 | 1.1e+00 | 1.0e+00 | 1.2e+00 | 8.8e-01 | 1.4e+00 |
| Total lipids (mmol/l) | PLASMA | 1.1e+00 | 1.2e-01 | 1.1e+00 | 1.0e+00 | 1.2e+00 | 9.0e-01 | 1.4e+00 |
| Phospholipids (mmol/l) | SERUM | 6.0e-01 | 7.0e-02 | 6.0e-01 | 5.5e-01 | 6.4e-01 | 4.6e-01 | 7.6e-01 |
| Phospholipids (mmol/l) | PLASMA | 6.1e-01 | 7.5e-02 | 6.1e-01 | 5.6e-01 | 6.4e-01 | 4.9e-01 | 8.0e-01 |
| Total cholesterol (mmol/l) | SERUM | 4.4e-01 | 6.4e-02 | 4.4e-01 | 4.1e-01 | 4.6e-01 | 2.4e-01 | 6.0e-01 |
| Total cholesterol (mmol/l) | PLASMA | 4.4e-01 | 7.8e-02 | 4.3e-01 | 4.0e-01 | 5.1e-01 | 2.6e-01 | 5.9e-01 |
| Cholesterol esters (mmol/l) | SERUM | 3.3e-01 | 6.1e-02 | 3.3e-01 | 3.0e-01 | 3.5e-01 | 1.2e-01 | 4.7e-01 |
| Cholesterol esters (mmol/l) | PLASMA | 3.3e-01 | 7.4e-02 | 3.2e-01 | 2.9e-01 | 4.0e-01 | 1.4e-01 | 4.6e-01 |
| Free cholesterol (mmol/l) | SERUM | 1.1e-01 | 1.2e-02 | 1.1e-01 | 1.0e-01 | 1.2e-01 | 8.9e-02 | 1.4e-01 |
| Free cholesterol (mmol/l) | PLASMA | 1.1e-01 | 1.3e-02 | 1.1e-01 | 1.0e-01 | 1.2e-01 | 9.3e-02 | 1.5e-01 |
| Triglycerides (mmol/l) | SERUM | 4.5e-02 | 1.9e-02 | 3.8e-02 | 3.3e-02 | 4.7e-02 | 2.7e-02 | 1.2e-01 |
| Triglycerides (mmol/l) | PLASMA | 5.0e-02 | 2.1e-02 | 4.3e-02 | 3.8e-02 | 5.3e-02 | 3.1e-02 | 1.2e-01 |
| **Lipoprotein particle size** | | | | | | | | |
| VLDL particle size (nm) | SERUM | 3.7e+01 | 1.6e+00 | 3.6e+01 | 3.5e+01 | 3.7e+01 | 3.4e+01 | 4.1e+01 |
| VLDL particle size (nm) | PLASMA | 3.7e+01 | 1.6e+00 | 3.6e+01 | 3.6e+01 | 3.7e+01 | 3.4e+01 | 4.1e+01 |
| LDL particle size (nm) | SERUM | 2.4e+01 | 8.0e-02 | 2.4e+01 | 2.3e+01 | 2.4e+01 | 2.3e+01 | 2.4e+01 |
| LDL particle size (nm) | PLASMA | 2.4e+01 | 8.9e-02 | 2.4e+01 | 2.3e+01 | 2.4e+01 | 2.3e+01 | 2.4e+01 |
| HDL particle size (nm) | SERUM | 1.0e+01 | 2.8e-01 | 1.0e+01 | 9.9e+00 | 1.0e+01 | 9.6e+00 | 1.1e+01 |
| HDL particle size (nm) | PLASMA | 1.0e+01 | 3.0e-01 | 1.0e+01 | 9.9e+00 | 1.0e+01 | 9.6e+00 | 1.1e+01 |
| **Cholesterol** | | | | | | | | |
| Total cholesterol (mmol/l) | SERUM | 4.3e+00 | 7.4e-01 | 4.3e+00 | 3.7e+00 | 4.8e+00 | 2.9e+00 | 5.7e+00 |
| Total cholesterol (mmol/l) | PLASMA | 4.0e+00 | 7.5e-01 | 4.1e+00 | 3.4e+00 | 4.7e+00 | 2.7e+00 | 5.3e+00 |
| VLDL cholesterol (mmol/l) | SERUM | 5.4e-01 | 3.4e-01 | 4.5e-01 | 3.7e-01 | 5.9e-01 | 1.6e-01 | 2.0e+00 |
| VLDL cholesterol (mmol/l) | PLASMA | 5.2e-01 | 4.0e-01 | 3.6e-01 | 3.1e-01 | 5.8e-01 | 2.0e-01 | 1.9e+00 |
| Remnant cholesterol (mmol/l) | SERUM | 1.1e+00 | 4.0e-01 | 1.0e+00 | 9.1e-01 | 1.2e+00 | 5.0e-01 | 2.6e+00 |

| Metabolic traits | sample | mean | sd | median | iqr25 | iqr75 | min | max |
|---|---|---|---|---|---|---|---|---|
| Remnant cholesterol (mmol/l) | PLASMA | 1.0e+00 | 4.8e-01 | 8.7e-01 | 7.4e-01 | 1.1e+00 | 5.4e-01 | 2.5e+00 |
| LDL cholesterol (mmol/l) | SERUM | 1.4e+00 | 3.6e-01 | 1.4e+00 | 1.2e+00 | 1.7e+00 | 7.5e-01 | 2.3e+00 |
| LDL cholesterol (mmol/l) | PLASMA | 1.3e+00 | 3.7e-01 | 1.2e+00 | 1.1e+00 | 1.6e+00 | 7.0e-01 | 2.1e+00 |
| HDL cholesterol (mmol/l) | SERUM | 1.7e+00 | 4.1e-01 | 1.6e+00 | 1.4e+00 | 2.1e+00 | 8.5e-01 | 2.5e+00 |
| HDL cholesterol (mmol/l) | PLASMA | 1.6e+00 | 4.1e-01 | 1.6e+00 | 1.4e+00 | 2.0e+00 | 8.2e-01 | 2.3e+00 |
| HDL2 cholesterol (mmol/l) | SERUM | 1.2e+00 | 3.8e-01 | 1.1e+00 | 9.7e-01 | 1.6e+00 | 4.5e-01 | 1.9e+00 |
| HDL2 cholesterol (mmol/l) | PLASMA | 1.2e+00 | 3.7e-01 | 1.1e+00 | 9.9e-01 | 1.5e+00 | 4.3e-01 | 1.8e+00 |
| HDL3 cholesterol (mmol/l) | SERUM | 4.8e-01 | 3.6e-02 | 4.8e-01 | 4.6e-01 | 5.1e-01 | 4.0e-01 | 5.5e-01 |
| HDL3 cholesterol (mmol/l) | PLASMA | 4.8e-01 | 3.6e-02 | 4.8e-01 | 4.4e-01 | 5.0e-01 | 3.9e-01 | 5.3e-01 |
| Esterified cholesterol (mmol/l) | SERUM | 2.9e+00 | 5.2e-01 | 2.9e+00 | 2.6e+00 | 3.3e+00 | 1.9e+00 | 4.1e+00 |
| Esterified cholesterol (mmol/l) | PLASMA | 2.8e+00 | 5.4e-01 | 2.7e+00 | 2.4e+00 | 3.3e+00 | 1.9e+00 | 3.8e+00 |
| Free cholesterol (mmol/l) | SERUM | 1.3e+00 | 2.3e-01 | 1.3e+00 | 1.1e+00 | 1.5e+00 | 9.3e-01 | 1.8e+00 |
| Free cholesterol (mmol/l) | PLASMA | 1.2e+00 | 2.2e-01 | 1.2e+00 | 1.0e+00 | 1.3e+00 | 8.3e-01 | 1.6e+00 |
| **Glycerides and phospholipids** | | | | | | | | |
| Triglycerides (mmol/l) | SERUM | 1.2e+00 | 8.9e-01 | 8.9e-01 | 6.9e-01 | 1.3e+00 | 4.7e-01 | 5.0e+00 |
| Triglycerides (mmol/l) | PLASMA | 1.3e+00 | 1.0e+00 | 8.9e-01 | 7.8e-01 | 1.3e+00 | 5.3e-01 | 5.0e+00 |
| VLDL triglycerides (mmol/l) | SERUM | 7.8e-01 | 8.0e-01 | 5.3e-01 | 3.9e-01 | 8.6e-01 | 1.6e-01 | 4.2e+00 |
| VLDL triglycerides (mmol/l) | PLASMA | 8.6e-01 | 9.1e-01 | 5.3e-01 | 4.3e-01 | 8.3e-01 | 2.0e-01 | 4.2e+00 |
| LDL triglycerides (mmol/l) | SERUM | 1.6e-01 | 4.5e-02 | 1.6e-01 | 1.3e-01 | 1.9e-01 | 8.2e-02 | 3.0e-01 |
| LDL triglycerides (mmol/l) | PLASMA | 1.7e-01 | 5.0e-02 | 1.7e-01 | 1.3e-01 | 1.9e-01 | 7.7e-02 | 3.0e-01 |
| HDL triglycerides (mmol/l) | SERUM | 1.4e-01 | 4.3e-02 | 1.3e-01 | 1.1e-01 | 1.5e-01 | 8.6e-02 | 3.2e-01 |
| HDL triglycerides (mmol/l) | PLASMA | 1.5e-01 | 4.9e-02 | 1.4e-01 | 1.1e-01 | 1.7e-01 | 9.9e-02 | 3.1e-01 |
| Diacylglycerol (mmol/l) | SERUM | 1.6e-02 | 2.0e-02 | 1.3e-02 | 0.0e+00 | 2.2e-02 | 0.0e+00 | 9.3e-02 |
| Diacylglycerol (mmol/l) | PLASMA | 1.7e-02 | 2.5e-02 | 9.0e-03 | 2.3e-04 | 2.6e-02 | 0.0e+00 | 1.1e-01 |
| Phosphoglycerides (mmol/l) | SERUM | 2.0e+00 | 3.8e-01 | 1.9e+00 | 1.7e+00 | 2.3e+00 | 1.5e+00 | 2.8e+00 |
| Phosphoglycerides (mmol/l) | PLASMA | 1.9e+00 | 4.0e-01 | 1.9e+00 | 1.5e+00 | 2.2e+00 | 1.4e+00 | 2.7e+00 |
| Phosphatidylcholine + other cholines (mmol/l) | SERUM | 2.0e+00 | 3.6e-01 | 2.0e+00 | 1.7e+00 | 2.3e+00 | 1.5e+00 | 2.7e+00 |
| Phosphatidylcholine + other cholines (mmol/l) | PLASMA | 1.9e+00 | 3.7e-01 | 1.9e+00 | 1.6e+00 | 2.2e+00 | 1.5e+00 | 2.6e+00 |
| Sphingomyelins (mmol/l) | SERUM | 4.4e-01 | 7.3e-02 | 4.5e-01 | 3.8e-01 | 5.2e-01 | 3.2e-01 | 5.7e-01 |

| Metabolic traits | sample | mean | sd | median | iqr25 | iqr75 | min | max |
|---|---|---|---|---|---|---|---|---|
| Sphingomyelins (mmol/l) | PLASMA | 4.3e-01 | 8.6e-02 | 4.2e-01 | 3.7e-01 | 4.8e-01 | 2.9e-01 | 6.2e-01 |
| Cholines (mmol/l) | SERUM | 2.4e+00 | 4.2e-01 | 2.4e+00 | 2.0e+00 | 2.7e+00 | 1.3e+00 | 3.1e+00 |
| Cholines (mmol/l) | PLASMA | 2.3e+00 | 4.1e-01 | 2.3e+00 | 1.9e+00 | 2.6e+00 | 1.7e+00 | 3.0e+00 |
| **Apolipoproteins** | | | | | | | | |
| Apolipoprotein A-I (g/l) | SERUM | 1.7e+00 | 2.1e-01 | 1.6e+00 | 1.5e+00 | 1.8e+00 | 1.3e+00 | 2.1e+00 |
| Apolipoprotein A-I (g/l) | PLASMA | 1.6e+00 | 1.9e-01 | 1.6e+00 | 1.5e+00 | 1.8e+00 | 1.3e+00 | 2.0e+00 |
| Apolipoprotein B (g/l) | SERUM | 7.8e-01 | 2.1e-01 | 7.3e-01 | 6.7e-01 | 8.1e-01 | 4.6e-01 | 1.5e+00 |
| Apolipoprotein B (g/l) | PLASMA | 7.4e-01 | 2.6e-01 | 6.5e-01 | 5.8e-01 | 7.8e-01 | 5.0e-01 | 1.6e+00 |
| **Fatty acids** | | | | | | | | |
| Total fatty acids (mmol/l) | SERUM | 1.1e+01 | 2.7e+00 | 1.1e+01 | 8.8e+00 | 1.2e+01 | 7.2e+00 | 2.1e+01 |
| Total fatty acids (mmol/l) | PLASMA | 1.1e+01 | 3.4e+00 | 1.0e+01 | 8.3e+00 | 1.1e+01 | 6.8e+00 | 2.1e+01 |
| Fatty acid chain length | SERUM | 1.7e+01 | 2.6e-01 | 1.7e+01 | 1.7e+01 | 1.8e+01 | 1.7e+01 | 1.8e+01 |
| Fatty acid chain length | PLASMA | 1.7e+01 | 2.6e-01 | 1.7e+01 | 1.7e+01 | 1.8e+01 | 1.7e+01 | 1.8e+01 |
| Degree of unsaturation | SERUM | 1.2e+00 | 6.9e-02 | 1.2e+00 | 1.2e+00 | 1.3e+00 | 1.0e+00 | 1.3e+00 |
| Degree of unsaturation | PLASMA | 1.2e+00 | 7.3e-02 | 1.2e+00 | 1.2e+00 | 1.2e+00 | 1.1e+00 | 1.3e+00 |
| Docosahexaenoic acid (mmol/l) | SERUM | 1.3e-01 | 4.7e-02 | 1.2e-01 | 9.8e-02 | 1.5e-01 | 4.5e-02 | 2.3e-01 |
| Docosahexaenoic acid (mmol/l) | PLASMA | 1.3e-01 | 4.9e-02 | 1.2e-01 | 9.8e-02 | 1.7e-01 | 6.6e-02 | 2.4e-01 |
| Linoleic acid (mmol/l) | SERUM | 2.8e+00 | 5.4e-01 | 2.8e+00 | 2.4e+00 | 3.2e+00 | 1.9e+00 | 4.5e+00 |
| Linoleic acid (mmol/l) | PLASMA | 2.8e+00 | 6.2e-01 | 2.7e+00 | 2.2e+00 | 3.1e+00 | 1.9e+00 | 4.5e+00 |
| Conjugated linoleic acid (mmol/l) | SERUM | 2.7e-02 | 1.9e-02 | 2.5e-02 | 1.3e-02 | 3.3e-02 | 3.0e-03 | 8.8e-02 |
| Conjugated linoleic acid (mmol/l) | PLASMA | 3.1e-02 | 2.0e-02 | 2.7e-02 | 2.0e-02 | 3.5e-02 | 0.0e+00 | 9.5e-02 |
| n-3 fatty acids (mmol/l) | SERUM | 4.1e-01 | 1.2e-01 | 3.7e-01 | 3.3e-01 | 4.6e-01 | 2.4e-01 | 8.1e-01 |
| n-3 fatty acids (mmol/l) | PLASMA | 4.2e-01 | 1.5e-01 | 3.5e-01 | 3.1e-01 | 4.9e-01 | 2.3e-01 | 8.8e-01 |
| n-6 fatty acids (mmol/l) | SERUM | 3.6e+00 | 6.2e-01 | 3.5e+00 | 3.0e+00 | 4.0e+00 | 2.5e+00 | 5.4e+00 |
| n-6 fatty acids (mmol/l) | PLASMA | 3.4e+00 | 6.9e-01 | 3.6e+00 | 2.8e+00 | 3.8e+00 | 2.4e+00 | 5.3e+00 |
| PUFA (mmol/l) | SERUM | 4.0e+00 | 7.2e-01 | 3.8e+00 | 3.4e+00 | 4.5e+00 | 2.8e+00 | 6.2e+00 |
| PUFA (mmol/l) | PLASMA | 3.8e+00 | 8.3e-01 | 3.9e+00 | 3.1e+00 | 4.3e+00 | 2.7e+00 | 6.2e+00 |
| MUFA (mmol/l) | SERUM | 2.8e+00 | 1.1e+00 | 2.5e+00 | 2.0e+00 | 3.0e+00 | 1.4e+00 | 7.1e+00 |
| MUFA (mmol/l) | PLASMA | 2.7e+00 | 1.3e+00 | 2.3e+00 | 1.8e+00 | 2.8e+00 | 1.3e+00 | 6.9e+00 |

| Metabolic traits | sample | mean | sd | median | iqr25 | iqr75 | min | max |
|---|---|---|---|---|---|---|---|---|
| Saturated fatty acids (mmol/l) | SERUM | 4.2e+00 | 1.0e+00 | 4.0e+00 | 3.4e+00 | 4.4e+00 | 3.0e+00 | 7.5e+00 |
| Saturated fatty acids (mmol/l) | PLASMA | 4.1e+00 | 1.3e+00 | 3.9e+00 | 3.2e+00 | 4.3e+00 | 2.8e+00 | 8.0e+00 |
| **Glycolysis related metabolites** | | | | | | | | |
| Glucose (mmol/l) | SERUM | 4.2e+00 | 5.8e-01 | 4.1e+00 | 3.8e+00 | 4.6e+00 | 2.6e+00 | 5.5e+00 |
| Glucose (mmol/l) | PLASMA | 4.1e+00 | 5.2e-01 | 4.2e+00 | 4.0e+00 | 4.3e+00 | 2.6e+00 | 5.4e+00 |
| Lactate (mmol/l) | SERUM | 1.5e+00 | 5.5e-01 | 1.4e+00 | 1.2e+00 | 1.6e+00 | 8.3e-01 | 4.0e+00 |
| Lactate (mmol/l) | PLASMA | 1.2e+00 | 3.3e-01 | 1.1e+00 | 9.7e-01 | 1.4e+00 | 6.0e-01 | 2.1e+00 |
| Pyruvate (mmol/l) | SERUM | 8.5e-02 | 3.4e-02 | 8.2e-02 | 5.9e-02 | 1.1e-01 | 2.5e-02 | 1.7e-01 |
| Citrate (mmol/l) | SERUM | 1.0e-01 | 2.7e-02 | 9.6e-02 | 8.5e-02 | 1.1e-01 | 6.3e-02 | 2.0e-01 |
| Citrate (mmol/l) | PLASMA | 1.7e-01 | 3.7e-02 | 1.7e-01 | 1.4e-01 | 2.0e-01 | 1.1e-01 | 2.5e-01 |
| Glycerol (mmol/l) | SERUM | 5.5e-02 | 1.9e-02 | 5.2e-02 | 4.6e-02 | 6.6e-02 | 1.6e-02 | 1.1e-01 |
| **Amino acids** | | | | | | | | |
| Alanine (mmol/l) | SERUM | 4.3e-01 | 5.5e-02 | 4.3e-01 | 3.9e-01 | 4.6e-01 | 3.2e-01 | 5.9e-01 |
| Alanine (mmol/l) | PLASMA | 4.1e-01 | 5.3e-02 | 4.1e-01 | 3.8e-01 | 4.3e-01 | 3.5e-01 | 5.6e-01 |
| Glutamine (mmol/l) | SERUM | 5.2e-01 | 6.0e-02 | 5.2e-01 | 4.9e-01 | 5.6e-01 | 3.8e-01 | 6.5e-01 |
| Glutamine (mmol/l) | PLASMA | 5.1e-01 | 5.7e-02 | 5.1e-01 | 4.7e-01 | 5.4e-01 | 3.9e-01 | 6.2e-01 |
| Glycine (mmol/l) | SERUM | 2.6e-01 | 7.6e-02 | 2.3e-01 | 2.1e-01 | 2.7e-01 | 1.7e-01 | 4.8e-01 |
| Histidine (mmol/l) | SERUM | 6.4e-02 | 6.5e-03 | 6.4e-02 | 6.1e-02 | 6.9e-02 | 5.2e-02 | 8.1e-02 |
| Histidine (mmol/l) | PLASMA | 6.3e-02 | 8.4e-03 | 6.2e-02 | 5.8e-02 | 7.0e-02 | 4.8e-02 | 8.0e-02 |
| *Branched-chain amino acids* | | | | | | | | |
| Isoleucine (mmol/l) | SERUM | 5.6e-02 | 2.1e-02 | 4.8e-02 | 4.2e-02 | 6.4e-02 | 3.7e-02 | 1.3e-01 |
| Isoleucine (mmol/l) | PLASMA | 6.0e-02 | 2.5e-02 | 5.0e-02 | 4.5e-02 | 6.6e-02 | 3.0e-02 | 1.2e-01 |
| Leucine (mmol/l) | SERUM | 7.1e-02 | 1.9e-02 | 6.4e-02 | 5.8e-02 | 7.9e-02 | 4.2e-02 | 1.2e-01 |
| Leucine (mmol/l) | PLASMA | 7.1e-02 | 2.2e-02 | 6.5e-02 | 5.7e-02 | 7.4e-02 | 4.1e-02 | 1.2e-01 |
| Valine (mmol/l) | SERUM | 1.6e-01 | 3.2e-02 | 1.5e-01 | 1.3e-01 | 1.7e-01 | 1.1e-01 | 2.5e-01 |
| Valine (mmol/l) | PLASMA | 1.5e-01 | 3.6e-02 | 1.5e-01 | 1.3e-01 | 1.6e-01 | 1.0e-01 | 2.5e-01 |
| *Aromatic amino acids* | | | | | | | | |
| Phenylalanine (mmol/l) | SERUM | 6.1e-02 | 6.9e-03 | 5.9e-02 | 5.5e-02 | 6.5e-02 | 4.9e-02 | 7.8e-02 |
| Phenylalanine (mmol/l) | PLASMA | 5.7e-02 | 6.5e-03 | 5.6e-02 | 5.4e-02 | 5.9e-02 | 4.7e-02 | 7.2e-02 |

| Metabolic traits | sample | mean | sd | median | iqr25 | iqr75 | min | max |
|---|---|---|---|---|---|---|---|---|
| Tyrosine (mmol/l) | SERUM | 5.3e-02 | 1.3e-02 | 4.9e-02 | 4.5e-02 | 5.8e-02 | 3.6e-02 | 8.8e-02 |
| Tyrosine (mmol/l) | PLASMA | 5.5e-02 | 1.2e-02 | 5.2e-02 | 4.8e-02 | 6.2e-02 | 3.9e-02 | 9.2e-02 |
| **Ketone bodies** | | | | | | | | |
| Acetate (mmol/l) | SERUM | 3.7e-02 | 9.7e-03 | 3.4e-02 | 3.1e-02 | 3.8e-02 | 2.7e-02 | 7.2e-02 |
| Acetate (mmol/l) | PLASMA | 4.7e-02 | 1.0e-02 | 4.4e-02 | 4.1e-02 | 4.8e-02 | 3.7e-02 | 7.4e-02 |
| Beta-hydroxybutyrate (mmol/l) | SERUM | 8.4e-02 | 2.5e-02 | 8.4e-02 | 7.0e-02 | 9.3e-02 | 5.0e-02 | 2.0e-01 |
| Beta-hydroxybutyrate (mmol/l) | PLASMA | 8.8e-02 | 1.5e-02 | 9.0e-02 | 7.9e-02 | 9.9e-02 | 5.4e-02 | 1.2e-01 |
| **Fluid balance** | | | | | | | | |
| Creatinine (mmol/l) | SERUM | 5.6e-02 | 8.7e-03 | 5.5e-02 | 4.9e-02 | 6.2e-02 | 4.0e-02 | 7.1e-02 |
| Creatinine (mmol/l) | PLASMA | 5.7e-02 | 8.8e-03 | 5.5e-02 | 5.0e-02 | 6.5e-02 | 4.1e-02 | 7.2e-02 |
| Albumin (signal area) | SERUM | 9.3e-02 | 4.0e-03 | 9.3e-02 | 9.1e-02 | 9.5e-02 | 8.4e-02 | 1.0e-01 |
| Albumin (signal area) | PLASMA | 9.3e-02 | 3.6e-03 | 9.2e-02 | 9.1e-02 | 9.5e-02 | 8.6e-02 | 1.0e-01 |
| **Inflammation** | | | | | | | | |
| Glycoprotein acetyls (mmol/l) | SERUM | 1.3e+00 | 2.9e-01 | 1.3e+00 | 1.1e+00 | 1.4e+00 | 1.0e+00 | 2.6e+00 |
| Glycoprotein acetyls (mmol/l) | PLASMA | 1.3e+00 | 3.3e-01 | 1.3e+00 | 1.1e+00 | 1.3e+00 | 1.0e+00 | 2.5e+00 |

*sTable 2. Serum, pre-storage handling effects: mean differences in metabolite concentrations (or trait value) per 24h increment in incubation duration at 4°C and 21°C, for serum samples.*

*# Associations in sFigure2 and Figures 1 and 2 are presented in SD-units.* These SD point estimates can be obtained by dividing the point estimate (beta) in absolute (clinically meaningful) concentration by the metabolic trait standard deviation (SD), both provided in the below table.

***Abbreviations**: **C**=cholesterol; **IDL**=intermediate-density lipoprotein; **LCI**=lower confidence interval; **LDL**=low-density lipoprotein; **HDL**=high-density lipoprotein; **MUFA**=monounsaturated fatty acids; **N.obs**= number of observations (samples); **N.indiv**=number of individuals; **PUFA**=polyunsaturated fatty acids; **SD**=standard deviation; **UCI**= upper confidence interval; **VLDL**=very-low-density lipoprotein.*

| Metabolic traits | temperature | N.obs | N.indiv | Beta | LCI | UCI | Pvalue | SD |
|---|---|---|---|---|---|---|---|---|
| **Lipoprotein subclasses** | | | | | | | | |
| *Extremely large VLDL* | | | | | | | | |
| Particle concentration (mol/l) | 4°C | 69 | 23 | -1.1e-11 | -1.6e-11 | -6.1e-12 | 8.5e-06 | 2.2e-10 |
| Particle concentration (mol/l) | 21°C | 69 | 23 | -6.3e-12 | -1.7e-11 | 4.3e-12 | 2.4e-01 | 2.2e-10 |
| Total lipids (mmol/l) | 4°C | 69 | 23 | -2.3e-03 | -3.3e-03 | -1.3e-03 | 8.5e-06 | 4.7e-02 |
| Total lipids (mmol/l) | 21°C | 69 | 23 | -1.3e-03 | -3.5e-03 | 9.3e-04 | 2.5e-01 | 4.7e-02 |
| Phospholipids (mmol/l) | 4°C | 69 | 23 | -2.7e-04 | -4.0e-04 | -1.4e-04 | 3.6e-05 | 5.8e-03 |
| Phospholipids (mmol/l) | 21°C | 69 | 23 | -1.8e-04 | -4.4e-04 | 8.5e-05 | 1.9e-01 | 5.8e-03 |
| Total cholesterol (mmol/l) | 4°C | 69 | 23 | -3.4e-04 | -4.9e-04 | -1.8e-04 | 1.6e-05 | 8.8e-03 |
| Total cholesterol (mmol/l) | 21°C | 69 | 23 | -4.5e-05 | -4.2e-04 | 3.3e-04 | 8.2e-01 | 8.8e-03 |
| Cholesterol esters (mmol/l) | 4°C | 69 | 23 | -1.8e-04 | -2.7e-04 | -8.9e-05 | 1.1e-04 | 5.0e-03 |
| Cholesterol esters (mmol/l) | 21°C | 69 | 23 | 8.4e-05 | -1.7e-04 | 3.4e-04 | 5.1e-01 | 5.0e-03 |
| Free cholesterol (mmol/l) | 4°C | 69 | 23 | -1.6e-04 | -2.3e-04 | -8.3e-05 | 3.9e-05 | 3.8e-03 |
| Free cholesterol (mmol/l) | 21°C | 69 | 23 | -1.3e-04 | -2.7e-04 | 1.2e-05 | 7.3e-02 | 3.8e-03 |
| Triglycerides (mmol/l) | 4°C | 69 | 23 | -1.7e-03 | -2.5e-03 | -9.5e-04 | 8.9e-06 | 3.2e-02 |
| Triglycerides (mmol/l) | 21°C | 69 | 23 | -1.1e-03 | -2.7e-03 | 5.3e-04 | 1.9e-01 | 3.2e-02 |
| *Very large VLDL* | | | | | | | | |
| Particle concentration (mol/l) | 4°C | 69 | 23 | -4.0e-11 | -6.3e-11 | -1.7e-11 | 5.2e-04 | 1.3e-09 |
| Particle concentration (mol/l) | 21°C | 69 | 23 | -4.0e-11 | -8.2e-11 | 2.8e-12 | 6.7e-02 | 1.3e-09 |
| Total lipids (mmol/l) | 4°C | 69 | 23 | -3.9e-03 | -6.1e-03 | -1.7e-03 | 4.2e-04 | 1.3e-01 |
| Total lipids (mmol/l) | 21°C | 69 | 23 | -3.6e-03 | -7.8e-03 | 5.3e-04 | 8.7e-02 | 1.3e-01 |

| Metabolic traits | temperature | N.obs | N.indiv | Beta | LCI | UCI | Pvalue | SD |
|---|---|---|---|---|---|---|---|---|
| Phospholipids (mmol/l) | 4°C | 69 | 23 | -6.7e-04 | -1.0e-03 | -3.1e-04 | 2.5e-04 | 2.1e-02 |
| Phospholipids (mmol/l) | 21°C | 69 | 23 | -5.2e-04 | -1.2e-03 | 1.9e-04 | 1.5e-01 | 2.1e-02 |
| Total cholesterol (mmol/l) | 4°C | 69 | 23 | -8.2e-04 | -1.2e-03 | -4.0e-04 | 1.0e-04 | 2.6e-02 |
| Total cholesterol (mmol/l) | 21°C | 69 | 23 | -1.7e-04 | -1.1e-03 | 7.7e-04 | 7.2e-01 | 2.6e-02 |
| Cholesterol esters (mmol/l) | 4°C | 69 | 23 | -4.3e-04 | -6.5e-04 | -2.0e-04 | 2.3e-04 | 1.4e-02 |
| Cholesterol esters (mmol/l) | 21°C | 69 | 23 | 6.3e-06 | -5.3e-04 | 5.4e-04 | 9.8e-01 | 1.4e-02 |
| Free cholesterol (mmol/l) | 4°C | 69 | 23 | -3.9e-04 | -5.8e-04 | -2.0e-04 | 5.2e-05 | 1.2e-02 |
| Free cholesterol (mmol/l) | 21°C | 69 | 23 | -1.8e-04 | -6.0e-04 | 2.3e-04 | 3.9e-01 | 1.2e-02 |
| Triglycerides (mmol/l) | 4°C | 69 | 23 | -2.4e-03 | -3.9e-03 | -1.0e-03 | 9.1e-04 | 8.1e-02 |
| Triglycerides (mmol/l) | 21°C | 69 | 23 | -2.9e-03 | -5.5e-03 | -3.2e-04 | 2.8e-02 | 8.1e-02 |
| *Large VLDL* | | | | | | | | |
| Particle concentration (mol/l) | 4°C | 69 | 23 | -1.8e-10 | -3.2e-10 | -3.8e-11 | 1.3e-02 | 7.5e-09 |
| Particle concentration (mol/l) | 21°C | 69 | 23 | -2.1e-10 | -4.3e-10 | 8.7e-12 | 6.0e-02 | 7.5e-09 |
| Total lipids (mmol/l) | 4°C | 69 | 23 | -1.0e-02 | -1.8e-02 | -2.3e-03 | 1.2e-02 | 4.4e-01 |
| Total lipids (mmol/l) | 21°C | 69 | 23 | -1.2e-02 | -2.5e-02 | 9.7e-04 | 7.0e-02 | 4.4e-01 |
| Phospholipids (mmol/l) | 4°C | 69 | 23 | -1.8e-03 | -3.3e-03 | -3.7e-04 | 1.4e-02 | 7.9e-02 |
| Phospholipids (mmol/l) | 21°C | 69 | 23 | -1.9e-03 | -4.2e-03 | 4.3e-04 | 1.1e-01 | 7.9e-02 |
| Total cholesterol (mmol/l) | 4°C | 69 | 23 | -2.3e-03 | -3.9e-03 | -6.8e-04 | 5.2e-03 | 1.0e-01 |
| Total cholesterol (mmol/l) | 21°C | 69 | 23 | -1.1e-03 | -4.1e-03 | 1.9e-03 | 4.6e-01 | 1.0e-01 |
| Cholesterol esters (mmol/l) | 4°C | 69 | 23 | -1.0e-03 | -1.9e-03 | -2.1e-04 | 1.4e-02 | 5.1e-02 |
| Cholesterol esters (mmol/l) | 21°C | 69 | 23 | 1.9e-04 | -1.6e-03 | 1.9e-03 | 8.3e-01 | 5.1e-02 |
| Free cholesterol (mmol/l) | 4°C | 69 | 23 | -1.2e-03 | -2.1e-03 | -4.2e-04 | 2.9e-03 | 5.0e-02 |
| Free cholesterol (mmol/l) | 21°C | 69 | 23 | -1.3e-03 | -2.6e-03 | 1.4e-05 | 5.3e-02 | 5.0e-02 |
| Triglycerides (mmol/l) | 4°C | 69 | 23 | -6.2e-03 | -1.1e-02 | -1.2e-03 | 1.6e-02 | 2.6e-01 |
| Triglycerides (mmol/l) | 21°C | 69 | 23 | -8.8e-03 | -1.7e-02 | -1.1e-03 | 2.5e-02 | 2.6e-01 |
| *Medium VLDL* | | | | | | | | |
| Particle concentration (mol/l) | 4°C | 69 | 23 | -3.1e-10 | -6.4e-10 | 2.4e-11 | 6.9e-02 | 1.9e-08 |

| Metabolic traits | temperature | N.obs | N.indiv | Beta | LCI | UCI | Pvalue | SD |
|---|---|---|---|---|---|---|---|---|
| Particle concentration (mol/l) | 21°C | 69 | 23 | -2.1e-10 | -7.7e-10 | 3.5e-10 | 4.6e-01 | 1.9e-08 |
| Total lipids (mmol/l) | 4°C | 69 | 23 | -1.0e-02 | -2.1e-02 | 9.1e-04 | 7.2e-02 | 6.3e-01 |
| Total lipids (mmol/l) | 21°C | 69 | 23 | -5.8e-03 | -2.4e-02 | 1.3e-02 | 5.4e-01 | 6.3e-01 |
| Phospholipids (mmol/l) | 4°C | 69 | 23 | -1.8e-03 | -3.9e-03 | 2.4e-04 | 8.4e-02 | 1.2e-01 |
| Phospholipids (mmol/l) | 21°C | 69 | 23 | -5.7e-04 | -4.1e-03 | 3.0e-03 | 7.6e-01 | 1.2e-01 |
| Total cholesterol (mmol/l) | 4°C | 69 | 23 | -1.9e-03 | -4.5e-03 | 7.0e-04 | 1.5e-01 | 1.6e-01 |
| Total cholesterol (mmol/l) | 21°C | 69 | 23 | 2.4e-03 | -2.5e-03 | 7.4e-03 | 3.4e-01 | 1.6e-01 |
| Cholesterol esters (mmol/l) | 4°C | 69 | 23 | -6.7e-04 | -2.2e-03 | 8.3e-04 | 3.8e-01 | 8.2e-02 |
| Cholesterol esters (mmol/l) | 21°C | 69 | 23 | 3.5e-03 | 2.0e-04 | 6.7e-03 | 3.8e-02 | 8.2e-02 |
| Free cholesterol (mmol/l) | 4°C | 69 | 23 | -1.2e-03 | -2.5e-03 | 3.5e-07 | 5.0e-02 | 7.7e-02 |
| Free cholesterol (mmol/l) | 21°C | 69 | 23 | -1.0e-03 | -3.0e-03 | 9.1e-04 | 3.0e-01 | 7.7e-02 |
| Triglycerides (mmol/l) | 4°C | 69 | 23 | -6.3e-03 | -1.3e-02 | 4.6e-05 | 5.2e-02 | 3.5e-01 |
| Triglycerides (mmol/l) | 21°C | 69 | 23 | -7.6e-03 | -1.8e-02 | 2.8e-03 | 1.5e-01 | 3.5e-01 |
| *Small VLDL* | | | | | | | | |
| Particle concentration (mol/l) | 4°C | 69 | 23 | 2.3e-10 | -1.3e-10 | 5.8e-10 | 2.1e-01 | 1.9e-08 |
| Particle concentration (mol/l) | 21°C | 69 | 23 | 8.7e-10 | 3.4e-10 | 1.4e-09 | 1.4e-03 | 1.9e-08 |
| Total lipids (mmol/l) | 4°C | 69 | 23 | 5.4e-03 | -1.4e-03 | 1.2e-02 | 1.2e-01 | 3.5e-01 |
| Total lipids (mmol/l) | 21°C | 69 | 23 | 1.9e-02 | 9.0e-03 | 3.0e-02 | 2.4e-04 | 3.5e-01 |
| Phospholipids (mmol/l) | 4°C | 69 | 23 | 2.1e-03 | 7.5e-04 | 3.5e-03 | 2.5e-03 | 7.1e-02 |
| Phospholipids (mmol/l) | 21°C | 69 | 23 | 5.3e-03 | 3.5e-03 | 7.0e-03 | 5.6e-09 | 7.1e-02 |
| Total cholesterol (mmol/l) | 4°C | 69 | 23 | 3.7e-03 | 1.2e-03 | 6.2e-03 | 3.5e-03 | 1.0e-01 |
| Total cholesterol (mmol/l) | 21°C | 69 | 23 | 1.3e-02 | 8.0e-03 | 1.7e-02 | 5.7e-08 | 1.0e-01 |
| Cholesterol esters (mmol/l) | 4°C | 69 | 23 | 2.5e-03 | 7.7e-04 | 4.3e-03 | 4.9e-03 | 5.6e-02 |
| Cholesterol esters (mmol/l) | 21°C | 69 | 23 | 9.6e-03 | 6.1e-03 | 1.3e-02 | 6.7e-08 | 5.6e-02 |
| Free cholesterol (mmol/l) | 4°C | 69 | 23 | 1.2e-03 | 3.1e-04 | 2.0e-03 | 7.7e-03 | 4.6e-02 |
| Free cholesterol (mmol/l) | 21°C | 69 | 23 | 2.9e-03 | 1.8e-03 | 4.1e-03 | 8.3e-07 | 4.6e-02 |
| Triglycerides (mmol/l) | 4°C | 69 | 23 | -4.2e-04 | -3.8e-03 | 3.0e-03 | 8.1e-01 | 1.9e-01 |

| Metabolic traits | temperature | N.obs | N.indiv | Beta | LCI | UCI | Pvalue | SD |
|---|---|---|---|---|---|---|---|---|
| Triglycerides (mmol/l) | 21°C | 69 | 23 | 1.4e-03 | -3.5e-03 | 6.3e-03 | 5.7e-01 | 1.9e-01 |
| *Very Small VLDL* | | | | | | | | |
| Particle concentration (mol/l) | 4°C | 69 | 23 | 5.9e-10 | 3.0e-10 | 8.8e-10 | 6.7e-05 | 1.1e-08 |
| Particle concentration (mol/l) | 21°C | 69 | 23 | 2.1e-09 | 1.4e-09 | 2.8e-09 | 1.4e-09 | 1.1e-08 |
| Total lipids (mmol/l) | 4°C | 69 | 23 | 7.5e-03 | 3.7e-03 | 1.1e-02 | 1.1e-04 | 1.3e-01 |
| Total lipids (mmol/l) | 21°C | 69 | 23 | 2.7e-02 | 1.8e-02 | 3.6e-02 | 2.1e-09 | 1.3e-01 |
| Phospholipids (mmol/l) | 4°C | 69 | 23 | 2.3e-03 | 9.8e-04 | 3.5e-03 | 5.6e-04 | 3.3e-02 |
| Phospholipids (mmol/l) | 21°C | 69 | 23 | 8.3e-03 | 6.0e-03 | 1.1e-02 | 7.7e-13 | 3.3e-02 |
| Total cholesterol (mmol/l) | 4°C | 69 | 23 | 3.5e-03 | 1.1e-03 | 5.8e-03 | 4.0e-03 | 5.1e-02 |
| Total cholesterol (mmol/l) | 21°C | 69 | 23 | 1.4e-02 | 8.3e-03 | 1.9e-02 | 5.3e-07 | 5.1e-02 |
| Cholesterol esters (mmol/l) | 4°C | 69 | 23 | 2.0e-03 | 1.7e-04 | 3.8e-03 | 3.2e-02 | 3.6e-02 |
| Cholesterol esters (mmol/l) | 21°C | 69 | 23 | 9.7e-03 | 5.5e-03 | 1.4e-02 | 7.1e-06 | 3.6e-02 |
| Free cholesterol (mmol/l) | 4°C | 69 | 23 | 1.5e-03 | 8.0e-04 | 2.2e-03 | 1.8e-05 | 1.6e-02 |
| Free cholesterol (mmol/l) | 21°C | 69 | 23 | 3.9e-03 | 2.7e-03 | 5.1e-03 | 1.1e-10 | 1.6e-02 |
| Triglycerides (mmol/l) | 4°C | 69 | 23 | 1.8e-03 | 9.3e-04 | 2.7e-03 | 4.8e-05 | 5.4e-02 |
| Triglycerides (mmol/l) | 21°C | 69 | 23 | 5.1e-03 | 3.3e-03 | 6.8e-03 | 1.1e-08 | 5.4e-02 |
| *IDL* | | | | | | | | |
| Particle concentration (mol/l) | 4°C | 69 | 23 | 4.9e-10 | -3.5e-10 | 1.3e-09 | 2.6e-01 | 2.1e-08 |
| Particle concentration (mol/l) | 21°C | 69 | 23 | 3.5e-09 | 1.8e-09 | 5.2e-09 | 6.8e-05 | 2.1e-08 |
| Total lipids (mmol/l) | 4°C | 69 | 23 | 4.3e-03 | -4.5e-03 | 1.3e-02 | 3.4e-01 | 2.1e-01 |
| Total lipids (mmol/l) | 21°C | 69 | 23 | 3.4e-02 | 1.6e-02 | 5.1e-02 | 1.5e-04 | 2.1e-01 |
| Phospholipids (mmol/l) | 4°C | 69 | 23 | 1.6e-03 | -5.3e-04 | 3.8e-03 | 1.4e-01 | 4.9e-02 |
| Phospholipids (mmol/l) | 21°C | 69 | 23 | 9.2e-03 | 5.2e-03 | 1.3e-02 | 7.7e-06 | 4.9e-02 |
| Total cholesterol (mmol/l) | 4°C | 69 | 23 | 6.1e-04 | -5.7e-03 | 6.9e-03 | 8.5e-01 | 1.3e-01 |
| Total cholesterol (mmol/l) | 21°C | 69 | 23 | 1.8e-02 | 6.1e-03 | 3.0e-02 | 3.0e-03 | 1.3e-01 |
| Cholesterol esters (mmol/l) | 4°C | 69 | 23 | -2.3e-04 | -4.9e-03 | 4.4e-03 | 9.2e-01 | 1.0e-01 |
| Cholesterol esters (mmol/l) | 21°C | 69 | 23 | 1.2e-02 | 3.6e-03 | 2.1e-02 | 5.8e-03 | 1.0e-01 |

| Metabolic traits | temperature | N.obs | N.indiv | Beta | LCI | UCI | Pvalue | SD |
|---|---|---|---|---|---|---|---|---|
| Free cholesterol (mmol/l) | 4°C | 69 | 23 | 8.5e-04 | -8.7e-04 | 2.6e-03 | 3.3e-01 | 3.8e-02 |
| Free cholesterol (mmol/l) | 21°C | 69 | 23 | 5.5e-03 | 2.3e-03 | 8.8e-03 | 8.7e-04 | 3.8e-02 |
| Triglycerides (mmol/l) | 4°C | 69 | 23 | 2.0e-03 | 1.3e-03 | 2.8e-03 | 4.1e-08 | 3.9e-02 |
| Triglycerides (mmol/l) | 21°C | 69 | 23 | 6.5e-03 | 4.2e-03 | 8.9e-03 | 4.1e-08 | 3.9e-02 |
| *Large LDL* | | | | | | | | |
| Particle concentration (mol/l) | 4°C | 69 | 23 | 9.6e-10 | -4.0e-10 | 2.3e-09 | 1.7e-01 | 3.5e-08 |
| Particle concentration (mol/l) | 21°C | 69 | 23 | 5.1e-09 | 2.5e-09 | 7.7e-09 | 1.2e-04 | 3.5e-08 |
| Total lipids (mmol/l) | 4°C | 69 | 23 | 6.2e-03 | -3.5e-03 | 1.6e-02 | 2.1e-01 | 2.4e-01 |
| Total lipids (mmol/l) | 21°C | 69 | 23 | 3.5e-02 | 1.7e-02 | 5.4e-02 | 2.0e-04 | 2.4e-01 |
| Phospholipids (mmol/l) | 4°C | 69 | 23 | 1.2e-03 | -8.2e-04 | 3.2e-03 | 2.4e-01 | 5.1e-02 |
| Phospholipids (mmol/l) | 21°C | 69 | 23 | 7.1e-03 | 3.6e-03 | 1.1e-02 | 7.3e-05 | 5.1e-02 |
| Total cholesterol (mmol/l) | 4°C | 69 | 23 | 3.0e-03 | -4.3e-03 | 1.0e-02 | 4.2e-01 | 1.7e-01 |
| Total cholesterol (mmol/l) | 21°C | 69 | 23 | 2.2e-02 | 8.9e-03 | 3.6e-02 | 1.1e-03 | 1.7e-01 |
| Cholesterol esters (mmol/l) | 4°C | 69 | 23 | 2.1e-03 | -3.4e-03 | 7.6e-03 | 4.5e-01 | 1.4e-01 |
| Cholesterol esters (mmol/l) | 21°C | 69 | 23 | 1.7e-02 | 6.7e-03 | 2.6e-02 | 9.9e-04 | 1.4e-01 |
| Free cholesterol (mmol/l) | 4°C | 69 | 23 | 9.2e-04 | -9.0e-04 | 2.7e-03 | 3.2e-01 | 4.2e-02 |
| Free cholesterol (mmol/l) | 21°C | 69 | 23 | 5.8e-03 | 2.1e-03 | 9.4e-03 | 1.8e-03 | 4.2e-02 |
| Triglycerides (mmol/l) | 4°C | 69 | 23 | 2.0e-03 | 1.2e-03 | 2.7e-03 | 9.4e-08 | 3.1e-02 |
| Triglycerides (mmol/l) | 21°C | 69 | 23 | 5.8e-03 | 3.4e-03 | 8.2e-03 | 1.8e-06 | 3.1e-02 |
| *Medium LDL* | | | | | | | | |
| Particle concentration (mol/l) | 4°C | 69 | 23 | 1.5e-09 | 2.8e-10 | 2.7e-09 | 1.6e-02 | 3.0e-08 |
| Particle concentration (mol/l) | 21°C | 69 | 23 | 5.1e-09 | 2.8e-09 | 7.4e-09 | 1.0e-05 | 3.0e-08 |
| Total lipids (mmol/l) | 4°C | 69 | 23 | 7.5e-03 | 1.2e-03 | 1.4e-02 | 1.9e-02 | 1.5e-01 |
| Total lipids (mmol/l) | 21°C | 69 | 23 | 2.5e-02 | 1.4e-02 | 3.7e-02 | 1.4e-05 | 1.5e-01 |
| Phospholipids (mmol/l) | 4°C | 69 | 23 | 1.5e-03 | 2.8e-04 | 2.7e-03 | 1.6e-02 | 3.7e-02 |
| Phospholipids (mmol/l) | 21°C | 69 | 23 | 5.3e-03 | 3.2e-03 | 7.4e-03 | 7.5e-07 | 3.7e-02 |
| Total cholesterol (mmol/l) | 4°C | 69 | 23 | 4.9e-03 | 1.0e-04 | 9.8e-03 | 4.5e-02 | 1.1e-01 |

| Metabolic traits | temperature | N.obs | N.indiv | Beta | LCI | UCI | Pvalue | SD |
|---|---|---|---|---|---|---|---|---|
| Total cholesterol (mmol/l) | 21°C | 69 | 23 | 1.7e-02 | 8.6e-03 | 2.6e-02 | 7.4e-05 | 1.1e-01 |
| Cholesterol esters (mmol/l) | 4°C | 69 | 23 | 3.5e-03 | -4.2e-04 | 7.3e-03 | 8.0e-02 | 8.9e-02 |
| Cholesterol esters (mmol/l) | 21°C | 69 | 23 | 1.3e-02 | 6.3e-03 | 2.0e-02 | 1.2e-04 | 8.9e-02 |
| Free cholesterol (mmol/l) | 4°C | 69 | 23 | 1.5e-03 | 5.1e-04 | 2.5e-03 | 2.9e-03 | 2.2e-02 |
| Free cholesterol (mmol/l) | 21°C | 69 | 23 | 4.2e-03 | 2.2e-03 | 6.1e-03 | 2.7e-05 | 2.2e-02 |
| Triglycerides (mmol/l) | 4°C | 69 | 23 | 1.0e-03 | 6.0e-04 | 1.5e-03 | 3.0e-06 | 1.6e-02 |
| Triglycerides (mmol/l) | 21°C | 69 | 23 | 2.9e-03 | 1.6e-03 | 4.3e-03 | 1.6e-05 | 1.6e-02 |
| *Small LDL* | | | | | | | | |
| Particle concentration (mol/l) | 4°C | 69 | 23 | 2.3e-09 | 8.3e-10 | 3.7e-09 | 2.1e-03 | 3.5e-08 |
| Particle concentration (mol/l) | 21°C | 69 | 23 | 6.3e-09 | 3.7e-09 | 8.9e-09 | 2.4e-06 | 3.5e-08 |
| Total lipids (mmol/l) | 4°C | 69 | 23 | 6.5e-03 | 2.3e-03 | 1.1e-02 | 2.2e-03 | 9.8e-02 |
| Total lipids (mmol/l) | 21°C | 69 | 23 | 1.8e-02 | 1.0e-02 | 2.5e-02 | 3.2e-06 | 9.8e-02 |
| Phospholipids (mmol/l) | 4°C | 69 | 23 | 1.9e-03 | 9.1e-04 | 2.8e-03 | 1.3e-04 | 2.6e-02 |
| Phospholipids (mmol/l) | 21°C | 69 | 23 | 4.4e-03 | 2.7e-03 | 6.1e-03 | 2.2e-07 | 2.6e-02 |
| Total cholesterol (mmol/l) | 4°C | 69 | 23 | 4.1e-03 | 1.0e-03 | 7.1e-03 | 9.4e-03 | 6.8e-02 |
| Total cholesterol (mmol/l) | 21°C | 69 | 23 | 1.2e-02 | 6.5e-03 | 1.7e-02 | 1.8e-05 | 6.8e-02 |
| Cholesterol esters (mmol/l) | 4°C | 69 | 23 | 2.7e-03 | 2.8e-04 | 5.1e-03 | 2.8e-02 | 5.4e-02 |
| Cholesterol esters (mmol/l) | 21°C | 69 | 23 | 8.7e-03 | 4.5e-03 | 1.3e-02 | 3.9e-05 | 5.4e-02 |
| Free cholesterol (mmol/l) | 4°C | 69 | 23 | 1.4e-03 | 6.9e-04 | 2.1e-03 | 9.2e-05 | 1.4e-02 |
| Free cholesterol (mmol/l) | 21°C | 69 | 23 | 3.2e-03 | 1.8e-03 | 4.6e-03 | 7.0e-06 | 1.4e-02 |
| Triglycerides (mmol/l) | 4°C | 69 | 23 | 5.4e-04 | 2.7e-04 | 8.1e-04 | 1.0e-04 | 1.4e-02 |
| Triglycerides (mmol/l) | 21°C | 69 | 23 | 1.5e-03 | 8.8e-04 | 2.0e-03 | 9.1e-07 | 1.4e-02 |
| *Very large HDL* | | | | | | | | |
| Particle concentration (mol/l) | 4°C | 69 | 23 | -1.1e-08 | -1.6e-08 | -6.7e-09 | 1.3e-06 | 2.3e-07 |
| Particle concentration (mol/l) | 21°C | 69 | 23 | -1.0e-08 | -2.4e-08 | 4.0e-09 | 1.6e-01 | 2.3e-07 |
| Total lipids (mmol/l) | 4°C | 69 | 23 | -1.1e-02 | -1.6e-02 | -6.7e-03 | 1.5e-06 | 2.3e-01 |
| Total lipids (mmol/l) | 21°C | 69 | 23 | -1.0e-02 | -2.4e-02 | 4.1e-03 | 1.6e-01 | 2.3e-01 |

| Metabolic traits | temperature | N.obs | N.indiv | Beta | LCI | UCI | Pvalue | SD |
|---|---|---|---|---|---|---|---|---|
| Phospholipids (mmol/l) | 4°C | 69 | 23 | -5.3e-03 | -7.6e-03 | -3.0e-03 | 6.3e-06 | 1.4e-01 |
| Phospholipids (mmol/l) | 21°C | 69 | 23 | -5.1e-03 | -1.2e-02 | 1.8e-03 | 1.5e-01 | 1.4e-01 |
| Total cholesterol (mmol/l) | 4°C | 69 | 23 | -5.2e-03 | -7.5e-03 | -2.9e-03 | 1.2e-05 | 9.7e-02 |
| Total cholesterol (mmol/l) | 21°C | 69 | 23 | -4.3e-03 | -1.2e-02 | 3.1e-03 | 2.5e-01 | 9.7e-02 |
| Cholesterol esters (mmol/l) | 4°C | 69 | 23 | -3.6e-03 | -5.3e-03 | -1.9e-03 | 2.8e-05 | 6.8e-02 |
| Cholesterol esters (mmol/l) | 21°C | 69 | 23 | -2.7e-03 | -8.2e-03 | 2.9e-03 | 3.5e-01 | 6.8e-02 |
| Free cholesterol (mmol/l) | 4°C | 69 | 23 | -1.6e-03 | -2.3e-03 | -9.2e-04 | 3.6e-06 | 3.0e-02 |
| Free cholesterol (mmol/l) | 21°C | 69 | 23 | -1.7e-03 | -3.6e-03 | 2.3e-04 | 8.5e-02 | 3.0e-02 |
| Triglycerides (mmol/l) | 4°C | 69 | 23 | -7.6e-04 | -1.1e-03 | -4.5e-04 | 1.1e-06 | 1.1e-02 |
| Triglycerides (mmol/l) | 21°C | 69 | 23 | -7.8e-04 | -1.5e-03 | -8.7e-05 | 2.7e-02 | 1.1e-02 |
| *Large HDL* | | | | | | | | |
| Particle concentration (mol/l) | 4°C | 69 | 23 | -1.6e-08 | -3.4e-08 | 2.3e-09 | 8.6e-02 | 6.7e-07 |
| Particle concentration (mol/l) | 21°C | 69 | 23 | -4.4e-08 | -8.7e-08 | -9.4e-10 | 4.5e-02 | 6.7e-07 |
| Total lipids (mmol/l) | 4°C | 69 | 23 | -9.8e-03 | -2.1e-02 | 1.4e-03 | 8.7e-02 | 4.3e-01 |
| Total lipids (mmol/l) | 21°C | 69 | 23 | -2.7e-02 | -5.5e-02 | -3.8e-04 | 4.7e-02 | 4.3e-01 |
| Phospholipids (mmol/l) | 4°C | 69 | 23 | -2.1e-03 | -7.6e-03 | 3.4e-03 | 4.5e-01 | 1.9e-01 |
| Phospholipids (mmol/l) | 21°C | 69 | 23 | -9.7e-03 | -2.3e-02 | 3.5e-03 | 1.5e-01 | 1.9e-01 |
| Total cholesterol (mmol/l) | 4°C | 69 | 23 | -5.8e-03 | -1.1e-02 | -7.6e-04 | 2.4e-02 | 2.3e-01 |
| Total cholesterol (mmol/l) | 21°C | 69 | 23 | -1.5e-02 | -2.8e-02 | -2.1e-03 | 2.3e-02 | 2.3e-01 |
| Cholesterol esters (mmol/l) | 4°C | 69 | 23 | -5.1e-03 | -9.2e-03 | -1.1e-03 | 1.3e-02 | 1.7e-01 |
| Cholesterol esters (mmol/l) | 21°C | 69 | 23 | -1.2e-02 | -2.2e-02 | -2.5e-03 | 1.4e-02 | 1.7e-01 |
| Free cholesterol (mmol/l) | 4°C | 69 | 23 | -6.4e-04 | -1.7e-03 | 4.5e-04 | 2.5e-01 | 5.4e-02 |
| Free cholesterol (mmol/l) | 21°C | 69 | 23 | -2.8e-03 | -6.2e-03 | 4.7e-04 | 9.3e-02 | 5.4e-02 |
| Triglycerides (mmol/l) | 4°C | 69 | 23 | -1.8e-03 | -3.2e-03 | -4.3e-04 | 1.0e-02 | 1.4e-02 |
| Triglycerides (mmol/l) | 21°C | 69 | 23 | -2.5e-03 | -4.2e-03 | -7.6e-04 | 4.9e-03 | 1.4e-02 |
| *Medium HDL* | | | | | | | | |
| Particle concentration (mol/l) | 4°C | 69 | 23 | 3.0e-08 | 9.6e-09 | 5.0e-08 | 3.8e-03 | 4.4e-07 |

| Metabolic traits | temperature | N.obs | N.indiv | Beta | LCI | UCI | Pvalue | SD |
|---|---|---|---|---|---|---|---|---|
| Particle concentration (mol/l) | 21°C | 69 | 23 | -1.9e-08 | -7.7e-08 | 3.8e-08 | 5.1e-01 | 4.4e-07 |
| Total lipids (mmol/l) | 4°C | 69 | 23 | 1.3e-02 | 3.9e-03 | 2.2e-02 | 4.6e-03 | 1.9e-01 |
| Total lipids (mmol/l) | 21°C | 69 | 23 | -8.6e-03 | -3.4e-02 | 1.6e-02 | 5.0e-01 | 1.9e-01 |
| Phospholipids (mmol/l) | 4°C | 69 | 23 | 6.0e-03 | 2.1e-03 | 9.8e-03 | 2.4e-03 | 8.6e-02 |
| Phospholipids (mmol/l) | 21°C | 69 | 23 | -3.8e-03 | -1.5e-02 | 7.4e-03 | 5.1e-01 | 8.6e-02 |
| Total cholesterol (mmol/l) | 4°C | 69 | 23 | 6.3e-03 | 1.3e-03 | 1.1e-02 | 1.3e-02 | 1.1e-01 |
| Total cholesterol (mmol/l) | 21°C | 69 | 23 | -5.3e-03 | -1.9e-02 | 8.7e-03 | 4.6e-01 | 1.1e-01 |
| Cholesterol esters (mmol/l) | 4°C | 69 | 23 | 4.6e-03 | 6.6e-04 | 8.5e-03 | 2.2e-02 | 8.8e-02 |
| Cholesterol esters (mmol/l) | 21°C | 69 | 23 | -4.6e-03 | -1.6e-02 | 6.4e-03 | 4.1e-01 | 8.8e-02 |
| Free cholesterol (mmol/l) | 4°C | 69 | 23 | 1.7e-03 | 6.6e-04 | 2.8e-03 | 1.5e-03 | 2.4e-02 |
| Free cholesterol (mmol/l) | 21°C | 69 | 23 | -6.8e-04 | -3.7e-03 | 2.3e-03 | 6.6e-01 | 2.4e-02 |
| Triglycerides (mmol/l) | 4°C | 69 | 23 | 5.9e-04 | 1.6e-04 | 1.0e-03 | 7.0e-03 | 2.0e-02 |
| Triglycerides (mmol/l) | 21°C | 69 | 23 | 4.7e-04 | -1.0e-04 | 1.0e-03 | 1.1e-01 | 2.0e-02 |
| *Small HDL* | | | | | | | | |
| Particle concentration (mol/l) | 4°C | 69 | 23 | 9.0e-08 | 5.7e-08 | 1.2e-07 | 7.1e-08 | 5.6e-07 |
| Particle concentration (mol/l) | 21°C | 69 | 23 | 1.2e-08 | -7.7e-08 | 1.0e-07 | 7.9e-01 | 5.6e-07 |
| Total lipids (mmol/l) | 4°C | 69 | 23 | 2.0e-02 | 1.3e-02 | 2.8e-02 | 9.0e-08 | 1.2e-01 |
| Total lipids (mmol/l) | 21°C | 69 | 23 | 2.9e-03 | -1.7e-02 | 2.3e-02 | 7.8e-01 | 1.2e-01 |
| Phospholipids (mmol/l) | 4°C | 69 | 23 | 8.9e-03 | 5.3e-03 | 1.2e-02 | 1.2e-06 | 7.6e-02 |
| Phospholipids (mmol/l) | 21°C | 69 | 23 | -7.5e-03 | -1.9e-02 | 3.5e-03 | 1.8e-01 | 7.6e-02 |
| Total cholesterol (mmol/l) | 4°C | 69 | 23 | 1.1e-02 | 6.0e-03 | 1.5e-02 | 6.5e-06 | 7.0e-02 |
| Total cholesterol (mmol/l) | 21°C | 69 | 23 | 8.9e-03 | -1.4e-03 | 1.9e-02 | 9.1e-02 | 7.0e-02 |
| Cholesterol esters (mmol/l) | 4°C | 69 | 23 | 8.4e-03 | 4.3e-03 | 1.3e-02 | 6.7e-05 | 6.5e-02 |
| Cholesterol esters (mmol/l) | 21°C | 69 | 23 | 9.6e-03 | 1.2e-03 | 1.8e-02 | 2.6e-02 | 6.5e-02 |
| Free cholesterol (mmol/l) | 4°C | 69 | 23 | 2.1e-03 | 1.4e-03 | 2.9e-03 | 3.7e-08 | 1.4e-02 |
| Free cholesterol (mmol/l) | 21°C | 69 | 23 | -6.2e-04 | -2.9e-03 | 1.7e-03 | 6.0e-01 | 1.4e-02 |
| Triglycerides (mmol/l) | 4°C | 69 | 23 | 8.5e-04 | 4.6e-04 | 1.2e-03 | 2.1e-05 | 2.4e-02 |

| Metabolic traits | temperature | N.obs | N.indiv | Beta | LCI | UCI | Pvalue | SD |
|---|---|---|---|---|---|---|---|---|
| Triglycerides (mmol/l) | 21°C | 69 | 23 | 1.5e-03 | 8.5e-04 | 2.2e-03 | 6.4e-06 | 2.4e-02 |
| **Lipoprotein particle size** | | | | | | | | |
| VLDL particle size (nm) | 4°C | 69 | 23 | -1.2e-01 | -1.6e-01 | -7.6e-02 | 8.5e-09 | 1.6e+00 |
| VLDL particle size (nm) | 21°C | 69 | 23 | -2.0e-01 | -2.8e-01 | -1.1e-01 | 8.0e-06 | 1.6e+00 |
| LDL particle size (nm) | 4°C | 69 | 23 | -1.3e-02 | -2.0e-02 | -6.0e-03 | 2.5e-04 | 8.5e-02 |
| LDL particle size (nm) | 21°C | 69 | 23 | -9.6e-03 | -2.4e-02 | 4.4e-03 | 1.8e-01 | 8.5e-02 |
| HDL particle size (nm) | 4°C | 69 | 23 | -1.9e-02 | -2.4e-02 | -1.3e-02 | 4.9e-11 | 2.8e-01 |
| HDL particle size (nm) | 21°C | 69 | 23 | -1.9e-02 | -3.4e-02 | -5.1e-03 | 7.9e-03 | 2.8e-01 |
| **Cholesterol** | | | | | | | | |
| Total cholesterol (mmol/l) | 4°C | 69 | 23 | 2.1e-02 | -7.4e-03 | 4.9e-02 | 1.5e-01 | 7.2e-01 |
| Total cholesterol (mmol/l) | 21°C | 69 | 23 | 8.1e-02 | 3.2e-02 | 1.3e-01 | 1.2e-03 | 7.2e-01 |
| VLDL cholesterol (mmol/l) | 4°C | 69 | 23 | 1.9e-03 | -5.6e-03 | 9.4e-03 | 6.1e-01 | 4.3e-01 |
| VLDL cholesterol (mmol/l) | 21°C | 69 | 23 | 2.8e-02 | 1.1e-02 | 4.4e-02 | 1.3e-03 | 4.3e-01 |
| Remnant cholesterol (mmol/l) | 4°C | 69 | 23 | 2.6e-03 | -9.4e-03 | 1.5e-02 | 6.8e-01 | 5.0e-01 |
| Remnant cholesterol (mmol/l) | 21°C | 69 | 23 | 4.5e-02 | 1.9e-02 | 7.2e-02 | 8.6e-04 | 5.0e-01 |
| LDL cholesterol (mmol/l) | 4°C | 69 | 23 | 1.2e-02 | -3.1e-03 | 2.7e-02 | 1.2e-01 | 3.5e-01 |
| LDL cholesterol (mmol/l) | 21°C | 69 | 23 | 5.1e-02 | 2.4e-02 | 7.8e-02 | 1.8e-04 | 3.5e-01 |
| HDL cholesterol (mmol/l) | 4°C | 69 | 23 | 6.2e-03 | -5.0e-03 | 1.7e-02 | 2.8e-01 | 4.2e-01 |
| HDL cholesterol (mmol/l) | 21°C | 69 | 23 | -1.6e-02 | -5.2e-02 | 2.0e-02 | 3.9e-01 | 4.2e-01 |
| HDL2 cholesterol (mmol/l) | 4°C | 69 | 23 | 2.8e-03 | -7.2e-03 | 1.3e-02 | 5.9e-01 | 3.9e-01 |
| HDL2 cholesterol (mmol/l) | 21°C | 69 | 23 | -2.0e-02 | -5.2e-02 | 1.2e-02 | 2.1e-01 | 3.9e-01 |
| HDL3 cholesterol (mmol/l) | 4°C | 69 | 23 | 3.3e-03 | 1.6e-03 | 5.1e-03 | 2.3e-04 | 3.4e-02 |
| HDL3 cholesterol (mmol/l) | 21°C | 69 | 23 | 4.4e-03 | -1.6e-03 | 1.0e-02 | 1.5e-01 | 3.4e-02 |
| Esterified cholesterol (mmol/l) | 4°C | 60 | 20 | 3.5e-02 | 8.9e-03 | 6.0e-02 | 8.2e-03 | 5.0e-01 |
| Esterified cholesterol (mmol/l) | 21°C | 60 | 20 | 1.5e-01 | 1.1e-01 | 1.9e-01 | 8.6e-12 | 5.0e-01 |
| Free cholesterol (mmol/l) | 4°C | 60 | 20 | -2.2e-02 | -3.7e-02 | -6.6e-03 | 5.1e-03 | 2.2e-01 |
| Free cholesterol (mmol/l) | 21°C | 60 | 20 | -8.3e-02 | -9.6e-02 | -7.0e-02 | 0.0e+00 | 2.2e-01 |

| Metabolic traits | temperature | N.obs | N.indiv | Beta | LCI | UCI | Pvalue | SD |
|---|---|---|---|---|---|---|---|---|
| **Glycerides and phospholipids** | | | | | | | | |
| Triglycerides (mmol/l) | 4°C | 69 | 23 | -8.9e-03 | -2.7e-02 | 8.9e-03 | 3.3e-01 | 1.1e+00 |
| Triglycerides (mmol/l) | 21°C | 69 | 23 | 4.1e-03 | -2.0e-02 | 2.8e-02 | 7.4e-01 | 1.1e+00 |
| VLDL triglycerides (mmol/l) | 4°C | 69 | 23 | -1.4e-02 | -3.1e-02 | 2.3e-03 | 9.2e-02 | 9.5e-01 |
| VLDL triglycerides (mmol/l) | 21°C | 69 | 23 | -1.2e-02 | -3.8e-02 | 1.4e-02 | 3.7e-01 | 9.5e-01 |
| LDL triglycerides (mmol/l) | 4°C | 69 | 23 | 3.5e-03 | 2.2e-03 | 4.9e-03 | 5.0e-07 | 6.1e-02 |
| LDL triglycerides (mmol/l) | 21°C | 69 | 23 | 1.0e-02 | 5.9e-03 | 1.4e-02 | 2.8e-06 | 6.1e-02 |
| HDL triglycerides (mmol/l) | 4°C | 69 | 23 | -1.8e-04 | -1.1e-03 | 7.3e-04 | 7.0e-01 | 5.7e-02 |
| HDL triglycerides (mmol/l) | 21°C | 69 | 23 | -5.9e-04 | -2.6e-03 | 1.4e-03 | 5.7e-01 | 5.7e-02 |
| Diacylglycerol (mmol/l) | 4°C | 57 | 19 | 5.9e-04 | -3.7e-03 | 4.9e-03 | 7.9e-01 | 3.0e-02 |
| Diacylglycerol (mmol/l) | 21°C | 57 | 19 | 2.4e-03 | -1.7e-03 | 6.6e-03 | 2.5e-01 | 3.0e-02 |
| Phosphoglycerides (mmol/l) | 4°C | 60 | 20 | -5.1e-03 | -2.9e-02 | 1.9e-02 | 6.7e-01 | 3.7e-01 |
| Phosphoglycerides (mmol/l) | 21°C | 60 | 20 | -6.3e-02 | -9.6e-02 | -2.9e-02 | 2.4e-04 | 3.7e-01 |
| Phosphatidylcholine + other cholines (mmol/l) | 4°C | 60 | 20 | -1.3e-02 | -3.3e-02 | 7.0e-03 | 2.0e-01 | 3.5e-01 |
| Phosphatidylcholine + other cholines (mmol/l) | 21°C | 60 | 20 | -6.0e-02 | -8.8e-02 | -3.2e-02 | 3.2e-05 | 3.5e-01 |
| Sphingomyelins (mmol/l) | 4°C | 60 | 20 | 2.4e-03 | -8.8e-03 | 1.4e-02 | 6.7e-01 | 6.2e-02 |
| Sphingomyelins (mmol/l) | 21°C | 60 | 20 | 3.3e-03 | -7.9e-03 | 1.5e-02 | 5.6e-01 | 6.2e-02 |
| Cholines (mmol/l) | 4°C | 60 | 20 | 5.4e-03 | -2.8e-02 | 3.9e-02 | 7.5e-01 | 3.7e-01 |
| Cholines (mmol/l) | 21°C | 60 | 20 | -4.4e-02 | -9.3e-02 | 3.7e-03 | 7.0e-02 | 3.7e-01 |
| **Apolipoproteins** | | | | | | | | |
| Apolipoprotein A-I (g/l) | 4°C | 69 | 23 | 2.5e-03 | -4.3e-03 | 9.4e-03 | 4.6e-01 | 2.0e-01 |
| Apolipoprotein A-I (g/l) | 21°C | 69 | 23 | -9.3e-03 | -2.8e-02 | 9.8e-03 | 3.4e-01 | 2.0e-01 |
| Apolipoprotein B (g/l) | 4°C | 69 | 23 | 9.5e-04 | -5.4e-03 | 7.3e-03 | 7.7e-01 | 2.6e-01 |
| Apolipoprotein B (g/l) | 21°C | 69 | 23 | 1.9e-02 | 6.0e-03 | 3.1e-02 | 3.9e-03 | 2.6e-01 |
| **Fatty acids** | | | | | | | | |
| Total fatty acids (mmol/l) | 4°C | 60 | 20 | -1.4e-03 | -1.2e-01 | 1.2e-01 | 9.8e-01 | 3.6e+00 |

| Metabolic traits | temperature | N.obs | N.indiv | Beta | LCI | UCI | Pvalue | SD |
|---|---|---|---|---|---|---|---|---|
| Total fatty acids (mmol/l) | 21°C | 60 | 20 | 4.5e-02 | -1.2e-01 | 2.1e-01 | 5.9e-01 | 3.6e+00 |
| Fatty acid chain length | 4°C | 60 | 20 | 1.1e-02 | -2.2e-02 | 4.5e-02 | 5.1e-01 | 2.9e-01 |
| Fatty acid chain length | 21°C | 60 | 20 | 1.5e-03 | -3.7e-02 | 4.0e-02 | 9.4e-01 | 2.9e-01 |
| Degree of unsaturation | 4°C | 60 | 20 | 1.0e-02 | -1.8e-03 | 2.2e-02 | 9.6e-02 | 8.7e-02 |
| Degree of unsaturation | 21°C | 60 | 20 | 8.6e-03 | 2.9e-03 | 1.4e-02 | 3.0e-03 | 8.7e-02 |
| Docosahexaenoic acid (mmol/l) | 4°C | 60 | 20 | 2.1e-03 | -1.1e-03 | 5.2e-03 | 1.9e-01 | 4.8e-02 |
| Docosahexaenoic acid (mmol/l) | 21°C | 60 | 20 | 3.3e-03 | 5.6e-04 | 6.0e-03 | 1.8e-02 | 4.8e-02 |
| Linoleic acid (mmol/l) | 4°C | 60 | 20 | 5.9e-03 | -1.9e-02 | 3.1e-02 | 6.4e-01 | 6.4e-01 |
| Linoleic acid (mmol/l) | 21°C | 60 | 20 | 3.1e-02 | -6.0e-03 | 6.8e-02 | 1.0e-01 | 6.4e-01 |
| Conjugated linoleic acid (mmol/l) | 4°C | 60 | 20 | -2.8e-03 | -6.0e-03 | 3.9e-04 | 8.5e-02 | 2.4e-02 |
| Conjugated linoleic acid (mmol/l) | 21°C | 60 | 20 | -2.1e-03 | -4.8e-03 | 5.6e-04 | 1.2e-01 | 2.4e-02 |
| n-3 fatty acids (mmol/l) | 4°C | 60 | 20 | 4.5e-03 | -3.5e-03 | 1.2e-02 | 2.7e-01 | 1.5e-01 |
| n-3 fatty acids (mmol/l) | 21°C | 60 | 20 | 9.8e-03 | 2.6e-03 | 1.7e-02 | 7.8e-03 | 1.5e-01 |
| n-6 fatty acids (mmol/l) | 4°C | 60 | 20 | 1.2e-02 | -2.2e-02 | 4.5e-02 | 5.0e-01 | 7.4e-01 |
| n-6 fatty acids (mmol/l) | 21°C | 60 | 20 | 3.7e-02 | -1.1e-02 | 8.4e-02 | 1.3e-01 | 7.4e-01 |
| PUFA (mmol/l) | 4°C | 60 | 20 | 1.6e-02 | -2.3e-02 | 5.5e-02 | 4.2e-01 | 8.7e-01 |
| PUFA (mmol/l) | 21°C | 60 | 20 | 4.6e-02 | -7.0e-03 | 1.0e-01 | 8.9e-02 | 8.7e-01 |
| MUFA (mmol/l) | 4°C | 60 | 20 | -1.8e-02 | -5.2e-02 | 1.7e-02 | 3.3e-01 | 1.5e+00 |
| MUFA (mmol/l) | 21°C | 60 | 20 | 1.1e-03 | -4.8e-02 | 5.0e-02 | 9.6e-01 | 1.5e+00 |
| Saturated fatty acids (mmol/l) | 4°C | 60 | 20 | -7.5e-05 | -5.2e-02 | 5.2e-02 | 1.0e+00 | 1.3e+00 |
| Saturated fatty acids (mmol/l) | 21°C | 60 | 20 | -2.0e-03 | -7.2e-02 | 6.8e-02 | 9.6e-01 | 1.3e+00 |
| **Glycolysis related metabolites** | | | | | | | | |
| Glucose (mmol/l) | 4°C | 69 | 23 | -9.1e-01 | -9.5e-01 | -8.7e-01 | 0.0e+00 | 1.4e+00 |
| Glucose (mmol/l) | 21°C | 69 | 23 | -1.9e+00 | -2.1e+00 | -1.7e+00 | 0.0e+00 | 1.4e+00 |
| Lactate (mmol/l) | 4°C | 69 | 23 | 1.6e+00 | 1.5e+00 | 1.7e+00 | 0.0e+00 | 2.6e+00 |
| Lactate (mmol/l) | 21°C | 69 | 23 | 3.7e+00 | 3.4e+00 | 3.9e+00 | 0.0e+00 | 2.6e+00 |

| Metabolic traits | temperature | N.obs | N.indiv | Beta | LCI | UCI | Pvalue | SD |
|---|---|---|---|---|---|---|---|---|
| Pyruvate (mmol/l) | 4°C | 60 | 20 | -1.3e-02 | -1.8e-02 | -7.9e-03 | 6.0e-07 | 6.0e-01 |
| Pyruvate (mmol/l) | 21°C | 60 | 20 | 7.3e-01 | 6.4e-01 | 8.2e-01 | 0.0e+00 | 6.0e-01 |
| Citrate (mmol/l) | 4°C | 69 | 23 | 4.9e-03 | 1.8e-03 | 8.0e-03 | 2.2e-03 | 2.0e-02 |
| Citrate (mmol/l) | 21°C | 69 | 23 | 1.9e-03 | -9.8e-04 | 4.8e-03 | 1.9e-01 | 2.0e-02 |
| Glycerol (mmol/l) | 4°C | 69 | 23 | 9.1e-03 | 7.0e-03 | 1.1e-02 | 0.0e+00 | 1.9e-02 |
| Glycerol (mmol/l) | 21°C | 69 | 23 | 2.7e-03 | -4.3e-04 | 5.7e-03 | 9.1e-02 | 1.9e-02 |
| **Amino acids** | | | | | | | | |
| Alanine (mmol/l) | 4°C | 69 | 23 | 3.1e-02 | 2.8e-02 | 3.3e-02 | 0.0e+00 | 1.2e-01 |
| Alanine (mmol/l) | 21°C | 69 | 23 | 1.5e-01 | 1.4e-01 | 1.6e-01 | 0.0e+00 | 1.2e-01 |
| Glutamine (mmol/l) | 4°C | 69 | 23 | -7.5e-03 | -1.0e-02 | -4.7e-03 | 1.0e-07 | 6.5e-02 |
| Glutamine (mmol/l) | 21°C | 69 | 23 | -4.6e-02 | -5.1e-02 | -4.1e-02 | 0.0e+00 | 6.5e-02 |
| Histidine (mmol/l) | 4°C | 69 | 23 | 6.8e-03 | 5.7e-03 | 7.9e-03 | 0.0e+00 | 1.4e-02 |
| Histidine (mmol/l) | 21°C | 69 | 23 | 1.7e-02 | 1.5e-02 | 1.8e-02 | 0.0e+00 | 1.4e-02 |
| Glycine (mmol/l) | 4°C | 69 | 23 | 4.0e-02 | 3.7e-02 | 4.2e-02 | 0.0e+00 | 8.6e-02 |
| Glycine (mmol/l) | 21°C | 69 | 23 | 8.1e-02 | 7.5e-02 | 8.6e-02 | 0.0e+00 | 8.6e-02 |
| *Branched-chain amino acids* | | | | | | | | |
| Isoleucine (mmol/l) | 4°C | 69 | 23 | 5.0e-03 | 4.0e-03 | 6.0e-03 | 0.0e+00 | 2.9e-02 |
| Isoleucine (mmol/l) | 21°C | 69 | 23 | 1.1e-02 | 9.3e-03 | 1.3e-02 | 0.0e+00 | 2.9e-02 |
| Leucine (mmol/l) | 4°C | 69 | 23 | 9.4e-03 | 8.7e-03 | 1.0e-02 | 0.0e+00 | 2.7e-02 |
| Leucine (mmol/l) | 21°C | 69 | 23 | 2.0e-02 | 1.9e-02 | 2.2e-02 | 0.0e+00 | 2.7e-02 |
| Valine (mmol/l) | 4°C | 69 | 23 | 1.3e-02 | 1.2e-02 | 1.4e-02 | 0.0e+00 | 4.1e-02 |
| Valine (mmol/l) | 21°C | 69 | 23 | 2.8e-02 | 2.6e-02 | 3.0e-02 | 0.0e+00 | 4.1e-02 |
| *Aromatic amino acids* | | | | | | | | |
| Phenylalanine (mmol/l) | 4°C | 69 | 23 | 1.1e-02 | 9.4e-03 | 1.2e-02 | 0.0e+00 | 1.5e-02 |
| Phenylalanine (mmol/l) | 21°C | 69 | 23 | 1.8e-02 | 1.6e-02 | 2.0e-02 | 0.0e+00 | 1.5e-02 |
| Tyrosine (mmol/l) | 4°C | 69 | 23 | 3.4e-03 | 2.8e-03 | 4.0e-03 | 0.0e+00 | 1.3e-02 |
| Tyrosine (mmol/l) | 21°C | 69 | 23 | 7.3e-03 | 6.5e-03 | 8.1e-03 | 0.0e+00 | 1.3e-02 |

| Metabolic traits | temperature | N.obs | N.indiv | Beta | LCI | UCI | Pvalue | SD |
|---|---|---|---|---|---|---|---|---|
| **Ketone bodies** | | | | | | | | |
| Acetate (mmol/l) | 4°C | 69 | 23 | 9.3e-03 | 8.0e-03 | 1.1e-02 | 0.0e+00 | 1.2e-02 |
| Acetate (mmol/l) | 21°C | 69 | 23 | 8.7e-03 | 6.0e-03 | 1.1e-02 | 4.8e-10 | 1.2e-02 |
| Beta-hydroxybutyrate (mmol/l) | 4°C | 69 | 23 | 1.9e-04 | -1.2e-03 | 1.6e-03 | 7.8e-01 | 2.0e-02 |
| Beta-hydroxybutyrate (mmol/l) | 21°C | 69 | 23 | 3.0e-03 | -7.9e-05 | 6.0e-03 | 5.6e-02 | 2.0e-02 |
| **Fluid balance** | | | | | | | | |
| Creatinine (mmol/l) | 4°C | 69 | 23 | 4.9e-04 | -1.7e-04 | 1.1e-03 | 1.4e-01 | 1.0e-02 |
| Creatinine (mmol/l) | 21°C | 69 | 23 | 3.3e-04 | -4.9e-04 | 1.1e-03 | 4.3e-01 | 1.0e-02 |
| Albumin (signal area) | 4°C | 69 | 23 | 6.8e-04 | 3.8e-04 | 9.9e-04 | 1.2e-05 | 4.6e-03 |
| Albumin (signal area) | 21°C | 69 | 23 | 2.6e-03 | 2.1e-03 | 3.1e-03 | 0.0e+00 | 4.6e-03 |
| **Inflammation** | | | | | | | | |
| Glycoprotein acetyls (mmol/l) | 4°C | 69 | 23 | 7.1e-03 | 2.4e-03 | 1.2e-02 | 3.2e-03 | 3.4e-01 |
| Glycoprotein acetyls (mmol/l) | 21°C | 69 | 23 | 3.2e-02 | 2.6e-02 | 3.9e-02 | 0.0e+00 | 3.4e-01 |

*sTable 3. EDTA-Plasma, pre-storage handling effects: mean differences in metabolite concentrations (or trait value) per 24h increment in incubation duration at 4°C and 21°C, for EDTA-samples samples. Pyruvate, glycerol and glycine are not quantified in EDTA-plasma samples due to the interfering resonances of EDTA on their signals.*

*# Associations in sFigure2 and Figures 1 and 2 are presented in SD-units.* These SD point estimate can be obtained by dividing the point estimate in absolute (clinically meaningful) concentration by the metabolic trait standard deviation (SD), both provided in the below table.

***Abbreviations**: **C**=cholesterol; **IDL**=intermediate-density lipoprotein; **LCI**=lower confidence interval; **LDL**=low-density lipoprotein; **HDL**=high-density lipoprotein; **MUFA**=monounsaturated fatty acids; **N.obs**= number of observations (samples); **N.indiv**=number of individuals; **PUFA**=polyunsaturated fatty acids; **SD**=standard deviation; **UCI**= upper confidence interval; **VLDL**=very-low-density lipoprotein.*

| Metabolic traits | temperature | N.obs | N.indiv | Beta | LCI | UCI | Pvalue | SD |
|---|---|---|---|---|---|---|---|---|
| **Lipoprotein subclasses** | | | | | | | | |
| *Extremely large VLDL* | | | | | | | | |
| Particle concentration (mol/l) | 4°C | 69 | 23 | -4.8e-12 | -1.3e-11 | 2.8e-12 | 2.2e-01 | 2.0e-10 |
| Particle concentration (mol/l) | 21°C | 69 | 23 | -2.8e-12 | -1.5e-11 | 9.5e-12 | 6.5e-01 | 2.0e-10 |
| Total lipids (mmol/l) | 4°C | 69 | 23 | -1.0e-03 | -2.7e-03 | 6.0e-04 | 2.1e-01 | 4.2e-02 |
| Total lipids (mmol/l) | 21°C | 69 | 23 | -6.0e-04 | -3.2e-03 | 2.0e-03 | 6.5e-01 | 4.2e-02 |
| Phospholipids (mmol/l) | 4°C | 69 | 23 | -1.5e-04 | -3.4e-04 | 3.9e-05 | 1.2e-01 | 5.3e-03 |
| Phospholipids (mmol/l) | 21°C | 69 | 23 | -1.4e-04 | -4.5e-04 | 1.7e-04 | 3.7e-01 | 5.3e-03 |
| Total cholesterol (mmol/l) | 4°C | 69 | 23 | -1.3e-04 | -4.1e-04 | 1.6e-04 | 3.9e-01 | 7.9e-03 |
| Total cholesterol (mmol/l) | 21°C | 69 | 23 | 5.1e-05 | -4.2e-04 | 5.3e-04 | 8.3e-01 | 7.9e-03 |
| Cholesterol esters (mmol/l) | 4°C | 69 | 23 | -2.4e-05 | -2.0e-04 | 1.5e-04 | 7.9e-01 | 4.5e-03 |
| Cholesterol esters (mmol/l) | 21°C | 69 | 23 | 1.7e-04 | -1.2e-04 | 4.7e-04 | 2.6e-01 | 4.5e-03 |
| Free cholesterol (mmol/l) | 4°C | 69 | 23 | -1.0e-04 | -2.1e-04 | 1.2e-05 | 8.1e-02 | 3.5e-03 |
| Free cholesterol (mmol/l) | 21°C | 69 | 23 | -1.2e-04 | -3.1e-04 | 6.9e-05 | 2.1e-01 | 3.5e-03 |
| Triglycerides (mmol/l) | 4°C | 69 | 23 | -7.5e-04 | -1.9e-03 | 4.1e-04 | 2.0e-01 | 2.9e-02 |
| Triglycerides (mmol/l) | 21°C | 69 | 23 | -5.1e-04 | -2.4e-03 | 1.3e-03 | 5.9e-01 | 2.9e-02 |
| *Very large VLDL* | | | | | | | | |
| Particle concentration (mol/l) | 4°C | 69 | 23 | -3.3e-11 | -6.7e-11 | 1.6e-12 | 6.2e-02 | 1.2e-09 |
| Particle concentration (mol/l) | 21°C | 69 | 23 | -3.7e-11 | -9.9e-11 | 2.4e-11 | 2.3e-01 | 1.2e-09 |
| Total lipids (mmol/l) | 4°C | 69 | 23 | -3.1e-03 | -6.5e-03 | 2.4e-04 | 6.8e-02 | 1.1e-01 |

| Metabolic traits | temperature | N.obs | N.indiv | Beta | LCI | UCI | Pvalue | SD |
|---|---|---|---|---|---|---|---|---|
| Total lipids (mmol/l) | 21°C | 69 | 23 | -3.4e-03 | -9.4e-03 | 2.6e-03 | 2.7e-01 | 1.1e-01 |
| Phospholipids (mmol/l) | 4°C | 69 | 23 | -5.5e-04 | -1.1e-03 | -9.3e-06 | 4.6e-02 | 1.9e-02 |
| Phospholipids (mmol/l) | 21°C | 69 | 23 | -4.8e-04 | -1.5e-03 | 5.0e-04 | 3.4e-01 | 1.9e-02 |
| Total cholesterol (mmol/l) | 4°C | 69 | 23 | -4.2e-04 | -1.2e-03 | 3.4e-04 | 2.8e-01 | 2.3e-02 |
| Total cholesterol (mmol/l) | 21°C | 69 | 23 | -8.9e-05 | -1.4e-03 | 1.2e-03 | 8.9e-01 | 2.3e-02 |
| Cholesterol esters (mmol/l) | 4°C | 69 | 23 | -1.7e-04 | -5.9e-04 | 2.6e-04 | 4.5e-01 | 1.3e-02 |
| Cholesterol esters (mmol/l) | 21°C | 69 | 23 | 8.4e-05 | -6.3e-04 | 8.0e-04 | 8.2e-01 | 1.3e-02 |
| Free cholesterol (mmol/l) | 4°C | 69 | 23 | -2.5e-04 | -5.8e-04 | 7.8e-05 | 1.3e-01 | 1.1e-02 |
| Free cholesterol (mmol/l) | 21°C | 69 | 23 | -1.7e-04 | -7.3e-04 | 3.9e-04 | 5.5e-01 | 1.1e-02 |
| Triglycerides (mmol/l) | 4°C | 69 | 23 | -2.2e-03 | -4.3e-03 | -1.5e-05 | 4.8e-02 | 7.2e-02 |
| Triglycerides (mmol/l) | 21°C | 69 | 23 | -2.8e-03 | -6.6e-03 | 9.8e-04 | 1.5e-01 | 7.2e-02 |
| *Large VLDL* | | | | | | | | |
| Particle concentration (mol/l) | 4°C | 69 | 23 | -2.8e-10 | -4.6e-10 | -9.8e-11 | 2.6e-03 | 6.7e-09 |
| Particle concentration (mol/l) | 21°C | 69 | 23 | -3.7e-10 | -7.0e-10 | -4.1e-11 | 2.8e-02 | 6.7e-09 |
| Total lipids (mmol/l) | 4°C | 69 | 23 | -1.6e-02 | -2.6e-02 | -5.2e-03 | 3.4e-03 | 3.9e-01 |
| Total lipids (mmol/l) | 21°C | 69 | 23 | -2.1e-02 | -4.0e-02 | -1.5e-03 | 3.5e-02 | 3.9e-01 |
| Phospholipids (mmol/l) | 4°C | 69 | 23 | -2.9e-03 | -4.7e-03 | -1.0e-03 | 2.2e-03 | 7.0e-02 |
| Phospholipids (mmol/l) | 21°C | 69 | 23 | -3.6e-03 | -7.0e-03 | -1.7e-04 | 4.0e-02 | 7.0e-02 |
| Total cholesterol (mmol/l) | 4°C | 69 | 23 | -2.1e-03 | -4.5e-03 | 3.4e-04 | 9.2e-02 | 9.1e-02 |
| Total cholesterol (mmol/l) | 21°C | 69 | 23 | -2.3e-03 | -6.8e-03 | 2.2e-03 | 3.2e-01 | 9.1e-02 |
| Cholesterol esters (mmol/l) | 4°C | 69 | 23 | -6.3e-04 | -1.9e-03 | 6.9e-04 | 3.5e-01 | 4.6e-02 |
| Cholesterol esters (mmol/l) | 21°C | 69 | 23 | -1.5e-04 | -2.5e-03 | 2.2e-03 | 9.0e-01 | 4.6e-02 |
| Free cholesterol (mmol/l) | 4°C | 69 | 23 | -1.5e-03 | -2.6e-03 | -3.0e-04 | 1.4e-02 | 4.5e-02 |
| Free cholesterol (mmol/l) | 21°C | 69 | 23 | -2.1e-03 | -4.3e-03 | 4.4e-05 | 5.5e-02 | 4.5e-02 |
| Triglycerides (mmol/l) | 4°C | 69 | 23 | -1.1e-02 | -1.7e-02 | -4.4e-03 | 9.6e-04 | 2.3e-01 |
| Triglycerides (mmol/l) | 21°C | 69 | 23 | -1.5e-02 | -2.6e-02 | -3.4e-03 | 1.1e-02 | 2.3e-01 |
| *Medium VLDL* | | | | | | | | |

| Metabolic traits | temperature | N.obs | N.indiv | Beta | LCI | UCI | Pvalue | SD |
|---|---|---|---|---|---|---|---|---|
| Particle concentration (mol/l) | 4°C | 69 | 23 | -4.9e-10 | -9.2e-10 | -6.7e-11 | 2.3e-02 | 1.7e-08 |
| Particle concentration (mol/l) | 21°C | 69 | 23 | -4.9e-10 | -1.3e-09 | 3.2e-10 | 2.4e-01 | 1.7e-08 |
| Total lipids (mmol/l) | 4°C | 69 | 23 | -1.6e-02 | -3.0e-02 | -1.8e-03 | 2.7e-02 | 5.5e-01 |
| Total lipids (mmol/l) | 21°C | 69 | 23 | -1.5e-02 | -4.2e-02 | 1.2e-02 | 2.8e-01 | 5.5e-01 |
| Phospholipids (mmol/l) | 4°C | 69 | 23 | -3.0e-03 | -5.6e-03 | -3.5e-04 | 2.6e-02 | 1.1e-01 |
| Phospholipids (mmol/l) | 21°C | 69 | 23 | -2.5e-03 | -7.6e-03 | 2.6e-03 | 3.3e-01 | 1.1e-01 |
| Total cholesterol (mmol/l) | 4°C | 69 | 23 | -2.3e-03 | -5.8e-03 | 1.2e-03 | 2.0e-01 | 1.4e-01 |
| Total cholesterol (mmol/l) | 21°C | 69 | 23 | 1.9e-03 | -5.1e-03 | 8.8e-03 | 6.0e-01 | 1.4e-01 |
| Cholesterol esters (mmol/l) | 4°C | 69 | 23 | -1.6e-04 | -2.1e-03 | 1.8e-03 | 8.7e-01 | 7.3e-02 |
| Cholesterol esters (mmol/l) | 21°C | 69 | 23 | 4.4e-03 | 5.1e-04 | 8.3e-03 | 2.7e-02 | 7.3e-02 |
| Free cholesterol (mmol/l) | 4°C | 69 | 23 | -2.1e-03 | -3.8e-03 | -4.7e-04 | 1.2e-02 | 6.9e-02 |
| Free cholesterol (mmol/l) | 21°C | 69 | 23 | -2.6e-03 | -5.7e-03 | 6.0e-04 | 1.1e-01 | 6.9e-02 |
| Triglycerides (mmol/l) | 4°C | 69 | 23 | -1.1e-02 | -1.9e-02 | -2.5e-03 | 1.0e-02 | 3.1e-01 |
| Triglycerides (mmol/l) | 21°C | 69 | 23 | -1.4e-02 | -2.9e-02 | 1.0e-03 | 6.8e-02 | 3.1e-01 |
| *Small VLDL* | | | | | | | | |
| Particle concentration (mol/l) | 4°C | 69 | 23 | -2.6e-10 | -5.9e-10 | 6.6e-11 | 1.2e-01 | 1.6e-08 |
| Particle concentration (mol/l) | 21°C | 69 | 23 | -1.4e-11 | -6.6e-10 | 6.3e-10 | 9.7e-01 | 1.6e-08 |
| Total lipids (mmol/l) | 4°C | 69 | 23 | -4.3e-03 | -1.0e-02 | 1.8e-03 | 1.6e-01 | 3.1e-01 |
| Total lipids (mmol/l) | 21°C | 69 | 23 | 2.3e-03 | -9.7e-03 | 1.4e-02 | 7.1e-01 | 3.1e-01 |
| Phospholipids (mmol/l) | 4°C | 69 | 23 | -6.5e-04 | -1.9e-03 | 6.4e-04 | 3.2e-01 | 6.2e-02 |
| Phospholipids (mmol/l) | 21°C | 69 | 23 | -8.7e-04 | -3.0e-03 | 1.3e-03 | 4.2e-01 | 6.2e-02 |
| Total cholesterol (mmol/l) | 4°C | 69 | 23 | 6.5e-04 | -1.5e-03 | 2.8e-03 | 5.6e-01 | 9.0e-02 |
| Total cholesterol (mmol/l) | 21°C | 69 | 23 | 9.9e-03 | 6.1e-03 | 1.4e-02 | 2.6e-07 | 9.0e-02 |
| Cholesterol esters (mmol/l) | 4°C | 69 | 23 | 1.2e-03 | -4.6e-04 | 2.9e-03 | 1.5e-01 | 5.1e-02 |
| Cholesterol esters (mmol/l) | 21°C | 69 | 23 | 9.9e-03 | 7.2e-03 | 1.3e-02 | 1.3e-12 | 5.1e-02 |
| Free cholesterol (mmol/l) | 4°C | 69 | 23 | -5.9e-04 | -1.3e-03 | 1.1e-04 | 1.0e-01 | 4.0e-02 |
| Free cholesterol (mmol/l) | 21°C | 69 | 23 | -5.6e-05 | -1.3e-03 | 1.2e-03 | 9.3e-01 | 4.0e-02 |

| Metabolic traits | temperature | N.obs | N.indiv | Beta | LCI | UCI | Pvalue | SD |
|---|---|---|---|---|---|---|---|---|
| Triglycerides (mmol/l) | 4°C | 69 | 23 | -4.3e-03 | -8.0e-03 | -6.1e-04 | 2.2e-02 | 1.6e-01 |
| Triglycerides (mmol/l) | 21°C | 69 | 23 | -6.7e-03 | -1.4e-02 | 1.7e-04 | 5.6e-02 | 1.6e-01 |
| *Very Small VLDL* | | | | | | | | |
| Particle concentration (mol/l) | 4°C | 69 | 23 | 2.4e-10 | -6.9e-11 | 5.5e-10 | 1.3e-01 | 9.9e-09 |
| Particle concentration (mol/l) | 21°C | 69 | 23 | 2.1e-09 | 1.6e-09 | 2.7e-09 | 1.1e-15 | 9.9e-09 |
| Total lipids (mmol/l) | 4°C | 69 | 23 | 3.5e-03 | -7.0e-04 | 7.6e-03 | 1.0e-01 | 1.2e-01 |
| Total lipids (mmol/l) | 21°C | 69 | 23 | 2.9e-02 | 2.2e-02 | 3.6e-02 | 0.0e+00 | 1.2e-01 |
| Phospholipids (mmol/l) | 4°C | 69 | 23 | 6.2e-04 | -7.4e-04 | 2.0e-03 | 3.7e-01 | 3.2e-02 |
| Phospholipids (mmol/l) | 21°C | 69 | 23 | 9.6e-03 | 7.8e-03 | 1.2e-02 | 0.0e+00 | 3.2e-02 |
| Total cholesterol (mmol/l) | 4°C | 69 | 23 | 3.0e-03 | 3.8e-06 | 6.0e-03 | 5.0e-02 | 4.9e-02 |
| Total cholesterol (mmol/l) | 21°C | 69 | 23 | 1.9e-02 | 1.4e-02 | 2.3e-02 | 0.0e+00 | 4.9e-02 |
| Cholesterol esters (mmol/l) | 4°C | 69 | 23 | 2.5e-03 | 2.1e-05 | 4.9e-03 | 4.8e-02 | 3.5e-02 |
| Cholesterol esters (mmol/l) | 21°C | 69 | 23 | 1.4e-02 | 1.1e-02 | 1.8e-02 | 6.2e-15 | 3.5e-02 |
| Free cholesterol (mmol/l) | 4°C | 69 | 23 | 5.4e-04 | -1.4e-04 | 1.2e-03 | 1.2e-01 | 1.5e-02 |
| Free cholesterol (mmol/l) | 21°C | 69 | 23 | 4.6e-03 | 3.8e-03 | 5.5e-03 | 0.0e+00 | 1.5e-02 |
| Triglycerides (mmol/l) | 4°C | 69 | 23 | -1.7e-04 | -1.1e-03 | 7.4e-04 | 7.1e-01 | 4.8e-02 |
| Triglycerides (mmol/l) | 21°C | 69 | 23 | 6.3e-04 | -7.2e-04 | 2.0e-03 | 3.6e-01 | 4.8e-02 |
| *IDL* | | | | | | | | |
| Particle concentration (mol/l) | 4°C | 69 | 23 | 2.2e-11 | -1.1e-09 | 1.1e-09 | 9.7e-01 | 2.0e-08 |
| Particle concentration (mol/l) | 21°C | 69 | 23 | 6.3e-09 | 5.0e-09 | 7.5e-09 | 0.0e+00 | 2.0e-08 |
| Total lipids (mmol/l) | 4°C | 69 | 23 | 1.9e-04 | -1.2e-02 | 1.2e-02 | 9.7e-01 | 2.0e-01 |
| Total lipids (mmol/l) | 21°C | 69 | 23 | 6.5e-02 | 5.3e-02 | 7.8e-02 | 0.0e+00 | 2.0e-01 |
| Phospholipids (mmol/l) | 4°C | 69 | 23 | 2.7e-04 | -2.3e-03 | 2.9e-03 | 8.4e-01 | 4.8e-02 |
| Phospholipids (mmol/l) | 21°C | 69 | 23 | 1.6e-02 | 1.4e-02 | 1.9e-02 | 0.0e+00 | 4.8e-02 |
| Total cholesterol (mmol/l) | 4°C | 69 | 23 | -2.1e-04 | -9.6e-03 | 9.2e-03 | 9.7e-01 | 1.3e-01 |
| Total cholesterol (mmol/l) | 21°C | 69 | 23 | 4.6e-02 | 3.7e-02 | 5.6e-02 | 0.0e+00 | 1.3e-01 |
| Cholesterol esters (mmol/l) | 4°C | 69 | 23 | -3.9e-04 | -7.6e-03 | 6.8e-03 | 9.2e-01 | 1.0e-01 |

| Metabolic traits | temperature | N.obs | N.indiv | Beta | LCI | UCI | Pvalue | SD |
|---|---|---|---|---|---|---|---|---|
| Cholesterol esters (mmol/l) | 21°C | 69 | 23 | 3.4e-02 | 2.7e-02 | 4.1e-02 | 0.0e+00 | 1.0e-01 |
| Free cholesterol (mmol/l) | 4°C | 69 | 23 | 1.7e-04 | -2.1e-03 | 2.5e-03 | 8.8e-01 | 3.6e-02 |
| Free cholesterol (mmol/l) | 21°C | 69 | 23 | 1.2e-02 | 1.0e-02 | 1.5e-02 | 0.0e+00 | 3.6e-02 |
| Triglycerides (mmol/l) | 4°C | 69 | 23 | 1.3e-04 | -1.0e-03 | 1.3e-03 | 8.2e-01 | 3.5e-02 |
| Triglycerides (mmol/l) | 21°C | 69 | 23 | 2.5e-03 | 9.8e-04 | 4.1e-03 | 1.4e-03 | 3.5e-02 |
| *Large LDL* | | | | | | | | |
| Particle concentration (mol/l) | 4°C | 69 | 23 | -5.4e-10 | -2.3e-09 | 1.2e-09 | 5.5e-01 | 3.4e-08 |
| Particle concentration (mol/l) | 21°C | 69 | 23 | 9.3e-09 | 7.7e-09 | 1.1e-08 | 0.0e+00 | 3.4e-08 |
| Total lipids (mmol/l) | 4°C | 69 | 23 | -3.6e-03 | -1.7e-02 | 9.4e-03 | 5.9e-01 | 2.4e-01 |
| Total lipids (mmol/l) | 21°C | 69 | 23 | 6.8e-02 | 5.7e-02 | 8.0e-02 | 0.0e+00 | 2.4e-01 |
| Phospholipids (mmol/l) | 4°C | 69 | 23 | -1.0e-03 | -3.9e-03 | 1.8e-03 | 4.9e-01 | 5.0e-02 |
| Phospholipids (mmol/l) | 21°C | 69 | 23 | 1.4e-02 | 1.2e-02 | 1.6e-02 | 0.0e+00 | 5.0e-02 |
| Total cholesterol (mmol/l) | 4°C | 69 | 23 | -2.6e-03 | -1.3e-02 | 7.4e-03 | 6.1e-01 | 1.7e-01 |
| Total cholesterol (mmol/l) | 21°C | 69 | 23 | 5.2e-02 | 4.3e-02 | 6.1e-02 | 0.0e+00 | 1.7e-01 |
| Cholesterol esters (mmol/l) | 4°C | 69 | 23 | -2.4e-03 | -9.9e-03 | 5.1e-03 | 5.3e-01 | 1.3e-01 |
| Cholesterol esters (mmol/l) | 21°C | 69 | 23 | 3.9e-02 | 3.2e-02 | 4.5e-02 | 0.0e+00 | 1.3e-01 |
| Free cholesterol (mmol/l) | 4°C | 69 | 23 | -1.3e-04 | -2.6e-03 | 2.4e-03 | 9.2e-01 | 4.1e-02 |
| Free cholesterol (mmol/l) | 21°C | 69 | 23 | 1.3e-02 | 1.1e-02 | 1.6e-02 | 0.0e+00 | 4.1e-02 |
| Triglycerides (mmol/l) | 4°C | 69 | 23 | -5.7e-05 | -1.3e-03 | 1.2e-03 | 9.3e-01 | 2.8e-02 |
| Triglycerides (mmol/l) | 21°C | 69 | 23 | 2.0e-03 | 4.6e-04 | 3.5e-03 | 1.1e-02 | 2.8e-02 |
| *Medium LDL* | | | | | | | | |
| Particle concentration (mol/l) | 4°C | 69 | 23 | -4.5e-10 | -2.0e-09 | 1.1e-09 | 5.7e-01 | 2.9e-08 |
| Particle concentration (mol/l) | 21°C | 69 | 23 | 7.8e-09 | 6.4e-09 | 9.2e-09 | 0.0e+00 | 2.9e-08 |
| Total lipids (mmol/l) | 4°C | 69 | 23 | -2.1e-03 | -9.9e-03 | 5.8e-03 | 6.1e-01 | 1.5e-01 |
| Total lipids (mmol/l) | 21°C | 69 | 23 | 4.1e-02 | 3.3e-02 | 4.8e-02 | 0.0e+00 | 1.5e-01 |
| Phospholipids (mmol/l) | 4°C | 69 | 23 | -6.1e-04 | -2.3e-03 | 1.1e-03 | 4.8e-01 | 3.6e-02 |
| Phospholipids (mmol/l) | 21°C | 69 | 23 | 7.4e-03 | 6.1e-03 | 8.8e-03 | 0.0e+00 | 3.6e-02 |

| Metabolic traits | temperature | N.obs | N.indiv | Beta | LCI | UCI | Pvalue | SD |
|---|---|---|---|---|---|---|---|---|
| Total cholesterol (mmol/l) | 4°C | 69 | 23 | -1.2e-03 | -7.2e-03 | 4.8e-03 | 6.9e-01 | 1.1e-01 |
| Total cholesterol (mmol/l) | 21°C | 69 | 23 | 3.3e-02 | 2.7e-02 | 3.8e-02 | 0.0e+00 | 1.1e-01 |
| Cholesterol esters (mmol/l) | 4°C | 69 | 23 | -1.1e-03 | -5.8e-03 | 3.6e-03 | 6.4e-01 | 8.6e-02 |
| Cholesterol esters (mmol/l) | 21°C | 69 | 23 | 2.6e-02 | 2.2e-02 | 3.0e-02 | 0.0e+00 | 8.6e-02 |
| Free cholesterol (mmol/l) | 4°C | 69 | 23 | -9.2e-05 | -1.4e-03 | 1.2e-03 | 8.9e-01 | 2.1e-02 |
| Free cholesterol (mmol/l) | 21°C | 69 | 23 | 6.5e-03 | 5.1e-03 | 7.9e-03 | 0.0e+00 | 2.1e-02 |
| Triglycerides (mmol/l) | 4°C | 69 | 23 | -2.6e-04 | -1.2e-03 | 6.5e-04 | 5.8e-01 | 1.4e-02 |
| Triglycerides (mmol/l) | 21°C | 69 | 23 | 4.3e-04 | -5.4e-04 | 1.4e-03 | 3.9e-01 | 1.4e-02 |
| *Small LDL* | | | | | | | | |
| Particle concentration (mol/l) | 4°C | 69 | 23 | -6.0e-10 | -2.5e-09 | 1.3e-09 | 5.4e-01 | 3.4e-08 |
| Particle concentration (mol/l) | 21°C | 69 | 23 | 8.5e-09 | 6.7e-09 | 1.0e-08 | 0.0e+00 | 3.4e-08 |
| Total lipids (mmol/l) | 4°C | 69 | 23 | -1.5e-03 | -6.8e-03 | 3.9e-03 | 6.0e-01 | 9.5e-02 |
| Total lipids (mmol/l) | 21°C | 69 | 23 | 2.5e-02 | 2.0e-02 | 3.0e-02 | 0.0e+00 | 9.5e-02 |
| Phospholipids (mmol/l) | 4°C | 69 | 23 | -5.2e-04 | -2.0e-03 | 9.2e-04 | 4.8e-01 | 2.5e-02 |
| Phospholipids (mmol/l) | 21°C | 69 | 23 | 4.7e-03 | 3.3e-03 | 6.1e-03 | 6.2e-11 | 2.5e-02 |
| Total cholesterol (mmol/l) | 4°C | 69 | 23 | -5.6e-04 | -4.4e-03 | 3.3e-03 | 7.8e-01 | 6.6e-02 |
| Total cholesterol (mmol/l) | 21°C | 69 | 23 | 2.0e-02 | 1.7e-02 | 2.4e-02 | 0.0e+00 | 6.6e-02 |
| Cholesterol esters (mmol/l) | 4°C | 69 | 23 | -5.5e-04 | -3.5e-03 | 2.4e-03 | 7.2e-01 | 5.2e-02 |
| Cholesterol esters (mmol/l) | 21°C | 69 | 23 | 1.6e-02 | 1.3e-02 | 1.9e-02 | 0.0e+00 | 5.2e-02 |
| Free cholesterol (mmol/l) | 4°C | 69 | 23 | -7.4e-06 | -9.0e-04 | 8.8e-04 | 9.9e-01 | 1.4e-02 |
| Free cholesterol (mmol/l) | 21°C | 69 | 23 | 4.2e-03 | 3.1e-03 | 5.3e-03 | 7.1e-14 | 1.4e-02 |
| Triglycerides (mmol/l) | 4°C | 69 | 23 | -3.8e-04 | -8.0e-04 | 3.8e-05 | 7.5e-02 | 1.3e-02 |
| Triglycerides (mmol/l) | 21°C | 69 | 23 | -1.6e-04 | -5.3e-04 | 2.2e-04 | 4.2e-01 | 1.3e-02 |
| *Very large HDL* | | | | | | | | |
| Particle concentration (mol/l) | 4°C | 69 | 23 | -9.8e-09 | -1.6e-08 | -3.9e-09 | 1.2e-03 | 2.2e-07 |
| Particle concentration (mol/l) | 21°C | 69 | 23 | 5.2e-09 | -4.5e-09 | 1.5e-08 | 2.9e-01 | 2.2e-07 |
| Total lipids (mmol/l) | 4°C | 69 | 23 | -9.8e-03 | -1.6e-02 | -3.7e-03 | 1.7e-03 | 2.2e-01 |

| Metabolic traits | temperature | N.obs | N.indiv | Beta | LCI | UCI | Pvalue | SD |
|---|---|---|---|---|---|---|---|---|
| Total lipids (mmol/l) | 21°C | 69 | 23 | 5.6e-03 | -4.3e-03 | 1.6e-02 | 2.7e-01 | 2.2e-01 |
| Phospholipids (mmol/l) | 4°C | 69 | 23 | -6.2e-03 | -9.3e-03 | -3.1e-03 | 1.1e-04 | 1.3e-01 |
| Phospholipids (mmol/l) | 21°C | 69 | 23 | -1.6e-04 | -5.2e-03 | 4.9e-03 | 9.5e-01 | 1.3e-01 |
| Total cholesterol (mmol/l) | 4°C | 69 | 23 | -3.2e-03 | -7.0e-03 | 5.9e-04 | 9.8e-02 | 9.3e-02 |
| Total cholesterol (mmol/l) | 21°C | 69 | 23 | 5.7e-03 | 3.8e-04 | 1.1e-02 | 3.5e-02 | 9.3e-02 |
| Cholesterol esters (mmol/l) | 4°C | 69 | 23 | -2.0e-03 | -4.9e-03 | 8.5e-04 | 1.7e-01 | 6.5e-02 |
| Cholesterol esters (mmol/l) | 21°C | 69 | 23 | 5.1e-03 | 1.2e-03 | 9.0e-03 | 1.1e-02 | 6.5e-02 |
| Free cholesterol (mmol/l) | 4°C | 69 | 23 | -1.2e-03 | -2.1e-03 | -2.0e-04 | 1.8e-02 | 2.8e-02 |
| Free cholesterol (mmol/l) | 21°C | 69 | 23 | 5.9e-04 | -8.2e-04 | 2.0e-03 | 4.1e-01 | 2.8e-02 |
| Triglycerides (mmol/l) | 4°C | 69 | 23 | -3.7e-04 | -1.0e-03 | 3.0e-04 | 2.8e-01 | 1.0e-02 |
| Triglycerides (mmol/l) | 21°C | 69 | 23 | 1.1e-04 | -7.1e-04 | 9.3e-04 | 7.9e-01 | 1.0e-02 |
| *Large HDL* | | | | | | | | |
| Particle concentration (mol/l) | 4°C | 69 | 23 | -3.6e-08 | -6.5e-08 | -6.0e-09 | 1.8e-02 | 6.3e-07 |
| Particle concentration (mol/l) | 21°C | 69 | 23 | -6.4e-08 | -9.7e-08 | -3.1e-08 | 1.7e-04 | 6.3e-07 |
| Total lipids (mmol/l) | 4°C | 69 | 23 | -2.3e-02 | -4.1e-02 | -4.6e-03 | 1.4e-02 | 4.0e-01 |
| Total lipids (mmol/l) | 21°C | 69 | 23 | -4.0e-02 | -6.1e-02 | -1.9e-02 | 1.9e-04 | 4.0e-01 |
| Phospholipids (mmol/l) | 4°C | 69 | 23 | -1.1e-02 | -2.1e-02 | -7.5e-04 | 3.6e-02 | 1.8e-01 |
| Phospholipids (mmol/l) | 21°C | 69 | 23 | -2.1e-02 | -3.3e-02 | -9.8e-03 | 2.7e-04 | 1.8e-01 |
| Total cholesterol (mmol/l) | 4°C | 69 | 23 | -1.2e-02 | -2.0e-02 | -4.0e-03 | 3.1e-03 | 2.1e-01 |
| Total cholesterol (mmol/l) | 21°C | 69 | 23 | -1.8e-02 | -2.7e-02 | -7.9e-03 | 3.6e-04 | 2.1e-01 |
| Cholesterol esters (mmol/l) | 4°C | 69 | 23 | -8.8e-03 | -1.5e-02 | -2.7e-03 | 4.4e-03 | 1.6e-01 |
| Cholesterol esters (mmol/l) | 21°C | 69 | 23 | -1.3e-02 | -2.1e-02 | -6.0e-03 | 3.5e-04 | 1.6e-01 |
| Free cholesterol (mmol/l) | 4°C | 69 | 23 | -3.0e-03 | -4.9e-03 | -1.1e-03 | 1.5e-03 | 5.2e-02 |
| Free cholesterol (mmol/l) | 21°C | 69 | 23 | -4.4e-03 | -6.9e-03 | -1.9e-03 | 6.5e-04 | 5.2e-02 |
| Triglycerides (mmol/l) | 4°C | 69 | 23 | -8.7e-05 | -1.6e-03 | 1.5e-03 | 9.1e-01 | 1.4e-02 |
| Triglycerides (mmol/l) | 21°C | 69 | 23 | -1.4e-03 | -3.0e-03 | 2.9e-04 | 1.1e-01 | 1.4e-02 |
| *Medium HDL* | | | | | | | | |

| Metabolic traits | temperature | N.obs | N.indiv | Beta | LCI | UCI | Pvalue | SD |
|---|---|---|---|---|---|---|---|---|
| Particle concentration (mol/l) | 4°C | 69 | 23 | -4.4e-08 | -9.7e-08 | 9.1e-09 | 1.0e-01 | 4.4e-07 |
| Particle concentration (mol/l) | 21°C | 69 | 23 | -1.3e-07 | -1.8e-07 | -6.7e-08 | 3.0e-05 | 4.4e-07 |
| Total lipids (mmol/l) | 4°C | 69 | 23 | -1.9e-02 | -4.2e-02 | 4.0e-03 | 1.0e-01 | 1.9e-01 |
| Total lipids (mmol/l) | 21°C | 69 | 23 | -5.3e-02 | -7.9e-02 | -2.8e-02 | 5.0e-05 | 1.9e-01 |
| Phospholipids (mmol/l) | 4°C | 69 | 23 | -8.2e-03 | -1.8e-02 | 1.9e-03 | 1.1e-01 | 8.8e-02 |
| Phospholipids (mmol/l) | 21°C | 69 | 23 | -2.7e-02 | -3.8e-02 | -1.5e-02 | 5.0e-06 | 8.8e-02 |
| Total cholesterol (mmol/l) | 4°C | 69 | 23 | -1.1e-02 | -2.3e-02 | 2.0e-03 | 9.9e-02 | 1.1e-01 |
| Total cholesterol (mmol/l) | 21°C | 69 | 23 | -2.4e-02 | -3.8e-02 | -9.9e-03 | 9.3e-04 | 1.1e-01 |
| Cholesterol esters (mmol/l) | 4°C | 69 | 23 | -8.6e-03 | -1.9e-02 | 1.4e-03 | 9.4e-02 | 8.2e-02 |
| Cholesterol esters (mmol/l) | 21°C | 69 | 23 | -1.8e-02 | -3.0e-02 | -7.2e-03 | 1.3e-03 | 8.2e-02 |
| Free cholesterol (mmol/l) | 4°C | 69 | 23 | -2.2e-03 | -4.9e-03 | 5.8e-04 | 1.2e-01 | 2.3e-02 |
| Free cholesterol (mmol/l) | 21°C | 69 | 23 | -5.8e-03 | -8.9e-03 | -2.7e-03 | 2.6e-04 | 2.3e-02 |
| Triglycerides (mmol/l) | 4°C | 69 | 23 | -1.8e-04 | -6.1e-04 | 2.5e-04 | 4.2e-01 | 1.7e-02 |
| Triglycerides (mmol/l) | 21°C | 69 | 23 | -2.5e-03 | -3.4e-03 | -1.7e-03 | 1.9e-09 | 1.7e-02 |
| *Small HDL* | | | | | | | | |
| Particle concentration (mol/l) | 4°C | 69 | 23 | -3.0e-08 | -1.1e-07 | 5.3e-08 | 4.8e-01 | 6.2e-07 |
| Particle concentration (mol/l) | 21°C | 69 | 23 | -2.0e-07 | -3.0e-07 | -9.5e-08 | 1.5e-04 | 6.2e-07 |
| Total lipids (mmol/l) | 4°C | 69 | 23 | -6.8e-03 | -2.6e-02 | 1.2e-02 | 4.8e-01 | 1.4e-01 |
| Total lipids (mmol/l) | 21°C | 69 | 23 | -4.3e-02 | -6.6e-02 | -2.0e-02 | 2.9e-04 | 1.4e-01 |
| Phospholipids (mmol/l) | 4°C | 69 | 23 | -3.8e-03 | -1.2e-02 | 4.0e-03 | 3.4e-01 | 8.1e-02 |
| Phospholipids (mmol/l) | 21°C | 69 | 23 | -3.5e-02 | -4.6e-02 | -2.4e-02 | 6.5e-10 | 8.1e-02 |
| Total cholesterol (mmol/l) | 4°C | 69 | 23 | -3.3e-03 | -1.5e-02 | 8.6e-03 | 5.9e-01 | 7.6e-02 |
| Total cholesterol (mmol/l) | 21°C | 69 | 23 | -6.8e-03 | -2.0e-02 | 6.7e-03 | 3.2e-01 | 7.6e-02 |
| Cholesterol esters (mmol/l) | 4°C | 69 | 23 | -2.5e-03 | -1.3e-02 | 8.0e-03 | 6.4e-01 | 7.0e-02 |
| Cholesterol esters (mmol/l) | 21°C | 69 | 23 | -9.3e-04 | -1.2e-02 | 1.1e-02 | 8.7e-01 | 7.0e-02 |
| Free cholesterol (mmol/l) | 4°C | 69 | 23 | -7.4e-04 | -2.3e-03 | 8.4e-04 | 3.6e-01 | 1.5e-02 |
| Free cholesterol (mmol/l) | 21°C | 69 | 23 | -5.9e-03 | -8.2e-03 | -3.6e-03 | 6.2e-07 | 1.5e-02 |

| Metabolic traits | temperature | N.obs | N.indiv | Beta | LCI | UCI | Pvalue | SD |
|---|---|---|---|---|---|---|---|---|
| Triglycerides (mmol/l) | 4°C | 69 | 23 | 2.0e-04 | -2.8e-04 | 6.7e-04 | 4.2e-01 | 2.1e-02 |
| Triglycerides (mmol/l) | 21°C | 69 | 23 | -1.7e-03 | -2.2e-03 | -1.2e-03 | 3.7e-11 | 2.1e-02 |
| **Lipoprotein particle size** | | | | | | | | |
| VLDL particle size (nm) | 4°C | 69 | 23 | -5.5e-02 | -1.1e-01 | 1.5e-03 | 5.7e-02 | 1.5e+00 |
| VLDL particle size (nm) | 21°C | 69 | 23 | -2.0e-01 | -3.0e-01 | -1.0e-01 | 5.6e-05 | 1.5e+00 |
| LDL particle size (nm) | 4°C | 69 | 23 | 5.7e-03 | -9.1e-03 | 2.0e-02 | 4.5e-01 | 9.8e-02 |
| LDL particle size (nm) | 21°C | 69 | 23 | 9.8e-03 | -8.5e-03 | 2.8e-02 | 3.0e-01 | 9.8e-02 |
| HDL particle size (nm) | 4°C | 69 | 23 | -1.0e-02 | -1.8e-02 | -2.5e-03 | 9.8e-03 | 2.7e-01 |
| HDL particle size (nm) | 21°C | 69 | 23 | 4.3e-03 | -5.4e-03 | 1.4e-02 | 3.9e-01 | 2.7e-01 |
| **Cholesterol** | | | | | | | | |
| Total cholesterol (mmol/l) | 4°C | 69 | 23 | -3.5e-02 | -8.0e-02 | 9.2e-03 | 1.2e-01 | 7.3e-01 |
| Total cholesterol (mmol/l) | 21°C | 69 | 23 | 1.4e-01 | 1.0e-01 | 1.7e-01 | 8.9e-16 | 7.3e-01 |
| VLDL cholesterol (mmol/l) | 4°C | 69 | 23 | -1.0e-03 | -1.0e-02 | 8.2e-03 | 8.3e-01 | 3.8e-01 |
| VLDL cholesterol (mmol/l) | 21°C | 69 | 23 | 2.8e-02 | 1.0e-02 | 4.7e-02 | 2.5e-03 | 3.8e-01 |
| Remnant cholesterol (mmol/l) | 4°C | 69 | 23 | -1.2e-03 | -1.7e-02 | 1.5e-02 | 8.8e-01 | 4.7e-01 |
| Remnant cholesterol (mmol/l) | 21°C | 69 | 23 | 7.5e-02 | 5.0e-02 | 1.0e-01 | 3.6e-09 | 4.7e-01 |
| LDL cholesterol (mmol/l) | 4°C | 69 | 23 | -4.2e-03 | -2.4e-02 | 1.5e-02 | 6.7e-01 | 3.4e-01 |
| LDL cholesterol (mmol/l) | 21°C | 69 | 23 | 1.1e-01 | 8.8e-02 | 1.2e-01 | 0.0e+00 | 3.4e-01 |
| HDL cholesterol (mmol/l) | 4°C | 69 | 23 | -3.0e-02 | -5.8e-02 | -2.3e-03 | 3.4e-02 | 3.9e-01 |
| HDL cholesterol (mmol/l) | 21°C | 69 | 23 | -4.3e-02 | -7.6e-02 | -8.8e-03 | 1.4e-02 | 3.9e-01 |
| HDL2 cholesterol (mmol/l) | 4°C | 69 | 23 | -2.7e-02 | -5.2e-02 | -2.8e-03 | 2.9e-02 | 3.6e-01 |
| HDL2 cholesterol (mmol/l) | 21°C | 69 | 23 | -4.0e-02 | -6.9e-02 | -1.1e-02 | 7.1e-03 | 3.6e-01 |
| HDL3 cholesterol (mmol/l) | 4°C | 69 | 23 | -2.6e-03 | -5.9e-03 | 7.2e-04 | 1.2e-01 | 3.4e-02 |
| HDL3 cholesterol (mmol/l) | 21°C | 69 | 23 | -2.5e-03 | -7.5e-03 | 2.6e-03 | 3.4e-01 | 3.4e-02 |
| Esterified cholesterol (mmol/l) | 4°C | 60 | 20 | -2.0e-02 | -5.5e-02 | 1.4e-02 | 2.4e-01 | 5.3e-01 |
| Esterified cholesterol (mmol/l) | 21°C | 60 | 20 | 1.5e-01 | 1.2e-01 | 1.9e-01 | 0.0e+00 | 5.3e-01 |
| Free cholesterol (mmol/l) | 4°C | 60 | 20 | -2.0e-02 | -4.0e-02 | -5.4e-04 | 4.4e-02 | 2.0e-01 |

| Metabolic traits | temperature | N.obs | N.indiv | Beta | LCI | UCI | Pvalue | SD |
|---|---|---|---|---|---|---|---|---|
| Free cholesterol (mmol/l) | 21°C | 60 | 20 | -1.9e-02 | -3.6e-02 | -1.6e-03 | 3.2e-02 | 2.0e-01 |
| **Glycerides and phospholipids** | | | | | | | | |
| Triglycerides (mmol/l) | 4°C | 69 | 23 | -2.9e-02 | -4.8e-02 | -9.4e-03 | 3.6e-03 | 9.5e-01 |
| Triglycerides (mmol/l) | 21°C | 69 | 23 | -3.8e-02 | -7.7e-02 | 3.9e-04 | 5.2e-02 | 9.5e-01 |
| VLDL triglycerides (mmol/l) | 4°C | 69 | 23 | -2.7e-02 | -4.7e-02 | -6.4e-03 | 1.0e-02 | 8.4e-01 |
| VLDL triglycerides (mmol/l) | 21°C | 69 | 23 | -3.8e-02 | -7.7e-02 | 1.4e-03 | 5.8e-02 | 8.4e-01 |
| LDL triglycerides (mmol/l) | 4°C | 69 | 23 | -6.9e-04 | -3.2e-03 | 1.8e-03 | 5.9e-01 | 5.5e-02 |
| LDL triglycerides (mmol/l) | 21°C | 69 | 23 | 2.3e-03 | -4.9e-04 | 5.0e-03 | 1.1e-01 | 5.5e-02 |
| HDL triglycerides (mmol/l) | 4°C | 69 | 23 | -1.3e-03 | -2.3e-03 | -3.2e-04 | 9.0e-03 | 4.9e-02 |
| HDL triglycerides (mmol/l) | 21°C | 69 | 23 | -5.4e-03 | -7.1e-03 | -3.7e-03 | 6.3e-10 | 4.9e-02 |
| Diacylglycerol (mmol/l) | 4°C | 57 | 19 | -1.4e-03 | -3.9e-03 | 1.2e-03 | 3.0e-01 | 2.5e-02 |
| Diacylglycerol (mmol/l) | 21°C | 57 | 19 | -5.0e-04 | -3.4e-03 | 2.4e-03 | 7.4e-01 | 2.5e-02 |
| Phosphoglycerides (mmol/l) | 4°C | 60 | 20 | -2.2e-02 | -5.2e-02 | 7.3e-03 | 1.4e-01 | 3.8e-01 |
| Phosphoglycerides (mmol/l) | 21°C | 60 | 20 | -1.1e-02 | -3.4e-02 | 1.2e-02 | 3.6e-01 | 3.8e-01 |
| Phosphatidylcholine + other cholines (mmol/l) | 4°C | 60 | 20 | -3.1e-02 | -6.0e-02 | -2.3e-03 | 3.4e-02 | 3.5e-01 |
| Phosphatidylcholine + other cholines (mmol/l) | 21°C | 60 | 20 | -5.2e-02 | -7.4e-02 | -3.0e-02 | 2.7e-06 | 3.5e-01 |
| Sphingomyelins (mmol/l) | 4°C | 60 | 20 | -1.1e-03 | -1.3e-02 | 1.1e-02 | 8.5e-01 | 7.1e-02 |
| Sphingomyelins (mmol/l) | 21°C | 60 | 20 | 4.3e-03 | -7.2e-03 | 1.6e-02 | 4.6e-01 | 7.1e-02 |
| Cholines (mmol/l) | 4°C | 60 | 20 | -2.5e-02 | -5.7e-02 | 6.6e-03 | 1.2e-01 | 3.7e-01 |
| Cholines (mmol/l) | 21°C | 60 | 20 | -4.0e-03 | -3.0e-02 | 2.2e-02 | 7.7e-01 | 3.7e-01 |
| **Apolipoproteins** | | | | | | | | |
| Apolipoprotein A-I (g/l) | 4°C | 69 | 23 | -2.1e-02 | -3.8e-02 | -4.8e-03 | 1.1e-02 | 2.0e-01 |
| Apolipoprotein A-I (g/l) | 21°C | 69 | 23 | -2.0e-02 | -3.7e-02 | -1.9e-03 | 3.0e-02 | 2.0e-01 |
| Apolipoprotein B (g/l) | 4°C | 69 | 23 | -4.1e-03 | -1.3e-02 | 4.5e-03 | 3.5e-01 | 2.4e-01 |
| Apolipoprotein B (g/l) | 21°C | 69 | 23 | 3.3e-02 | 2.1e-02 | 4.5e-02 | 1.2e-07 | 2.4e-01 |
| **Fatty acids** | | | | | | | | |

| Metabolic traits | temperature | N.obs | N.indiv | Beta | LCI | UCI | Pvalue | SD |
|---|---|---|---|---|---|---|---|---|
| Total fatty acids (mmol/l) | 4°C | 60 | 20 | -2.1e-01 | -3.7e-01 | -3.8e-02 | 1.6e-02 | 3.4e+00 |
| Total fatty acids (mmol/l) | 21°C | 60 | 20 | -2.9e-02 | -1.7e-01 | 1.1e-01 | 6.8e-01 | 3.4e+00 |
| Fatty acid chain length | 4°C | 60 | 20 | -9.0e-03 | -5.1e-02 | 3.3e-02 | 6.7e-01 | 2.6e-01 |
| Fatty acid chain length | 21°C | 60 | 20 | -2.2e-02 | -6.8e-02 | 2.3e-02 | 3.4e-01 | 2.6e-01 |
| Degree of unsaturation | 4°C | 60 | 20 | 2.1e-03 | -3.6e-03 | 7.7e-03 | 4.7e-01 | 7.4e-02 |
| Degree of unsaturation | 21°C | 60 | 20 | 7.7e-03 | 9.9e-04 | 1.4e-02 | 2.5e-02 | 7.4e-02 |
| Docosahexaenoic acid (mmol/l) | 4°C | 60 | 20 | -5.2e-04 | -4.3e-03 | 3.2e-03 | 7.8e-01 | 4.8e-02 |
| Docosahexaenoic acid (mmol/l) | 21°C | 60 | 20 | 1.8e-03 | -5.1e-04 | 4.0e-03 | 1.3e-01 | 4.8e-02 |
| Linoleic acid (mmol/l) | 4°C | 60 | 20 | -3.9e-02 | -7.7e-02 | -1.4e-03 | 4.2e-02 | 6.3e-01 |
| Linoleic acid (mmol/l) | 21°C | 60 | 20 | 2.7e-02 | -2.2e-03 | 5.5e-02 | 7.1e-02 | 6.3e-01 |
| Conjugated linoleic acid (mmol/l) | 4°C | 60 | 20 | -2.3e-03 | -4.7e-03 | -1.4e-05 | 4.9e-02 | 2.2e-02 |
| Conjugated linoleic acid (mmol/l) | 21°C | 60 | 20 | -2.1e-03 | -4.7e-03 | 6.0e-04 | 1.3e-01 | 2.2e-02 |
| n-3 fatty acids (mmol/l) | 4°C | 60 | 20 | -2.0e-03 | -1.1e-02 | 6.7e-03 | 6.5e-01 | 1.6e-01 |
| n-3 fatty acids (mmol/l) | 21°C | 60 | 20 | 6.7e-03 | 1.7e-03 | 1.2e-02 | 8.9e-03 | 1.6e-01 |
| n-6 fatty acids (mmol/l) | 4°C | 60 | 20 | -5.6e-02 | -1.0e-01 | -9.5e-03 | 1.8e-02 | 7.1e-01 |
| n-6 fatty acids (mmol/l) | 21°C | 60 | 20 | 3.1e-02 | -4.2e-03 | 6.6e-02 | 8.5e-02 | 7.1e-01 |
| PUFA (mmol/l) | 4°C | 60 | 20 | -5.8e-02 | -1.1e-01 | -5.7e-03 | 3.0e-02 | 8.5e-01 |
| PUFA (mmol/l) | 21°C | 60 | 20 | 3.7e-02 | -1.6e-03 | 7.6e-02 | 6.0e-02 | 8.5e-01 |
| MUFA (mmol/l) | 4°C | 60 | 20 | -6.0e-02 | -1.0e-01 | -1.5e-02 | 8.4e-03 | 1.3e+00 |
| MUFA (mmol/l) | 21°C | 60 | 20 | -3.2e-02 | -8.1e-02 | 1.6e-02 | 1.9e-01 | 1.3e+00 |
| Saturated fatty acids (mmol/l) | 4°C | 60 | 20 | -8.8e-02 | -1.7e-01 | -4.9e-03 | 3.8e-02 | 1.3e+00 |
| Saturated fatty acids (mmol/l) | 21°C | 60 | 20 | -3.4e-02 | -9.9e-02 | 3.0e-02 | 3.0e-01 | 1.3e+00 |
| **Glycolysis related metabolites** | | | | | | | | |
| Glucose (mmol/l) | 4°C | 69 | 23 | -1.2e+00 | -1.3e+00 | -1.2e+00 | 0.0e+00 | 1.5e+00 |
| Glucose (mmol/l) | 21°C | 69 | 23 | -1.9e+00 | -2.2e+00 | -1.7e+00 | 0.0e+00 | 1.5e+00 |
| Lactate (mmol/l) | 4°C | 69 | 23 | 2.1e+00 | 2.0e+00 | 2.2e+00 | 0.0e+00 | 2.5e+00 |

| Metabolic traits | temperature | N.obs | N.indiv | Beta | LCI | UCI | Pvalue | SD |
|---|---|---|---|---|---|---|---|---|
| Lactate (mmol/l) | 21°C | 69 | 23 | 3.3e+00 | 2.9e+00 | 3.7e+00 | 0.0e+00 | 2.5e+00 |
| Pyruvate (mmol/l) | 4°C | | | NA | NA | NA | NA | NA |
| Pyruvate (mmol/l) | 21°C | | | NA | NA | NA | NA | NA |
| Citrate (mmol/l) | 4°C | 69 | 23 | -2.0e-03 | -5.5e-03 | 1.4e-03 | 2.4e-01 | 3.4e-02 |
| Citrate (mmol/l) | 21°C | 69 | 23 | -7.5e-03 | -1.1e-02 | -4.0e-03 | 2.1e-05 | 3.4e-02 |
| Glycerol (mmol/l) | 4°C | | | NA | NA | NA | NA | NA |
| Glycerol (mmol/l) | 21°C | | | NA | NA | NA | NA | NA |
| **Amino acids** | | | | | | | | |
| Alanine (mmol/l) | 4°C | 69 | 23 | 6.4e-03 | 3.3e-03 | 9.4e-03 | 3.6e-05 | 1.2e-01 |
| Alanine (mmol/l) | 21°C | 69 | 23 | 1.3e-01 | 1.2e-01 | 1.4e-01 | 0.0e+00 | 1.2e-01 |
| Glutamine (mmol/l) | 4°C | 69 | 23 | -1.4e-02 | -1.9e-02 | -9.5e-03 | 2.5e-09 | 6.7e-02 |
| Glutamine (mmol/l) | 21°C | 69 | 23 | -5.8e-02 | -6.4e-02 | -5.3e-02 | 0.0e+00 | 6.7e-02 |
| Histidine (mmol/l) | 4°C | 69 | 23 | -3.5e-03 | -4.9e-03 | -2.2e-03 | 3.0e-07 | 7.9e-03 |
| Histidine (mmol/l) | 21°C | 69 | 23 | -5.6e-03 | -7.3e-03 | -3.9e-03 | 1.2e-10 | 7.9e-03 |
| Glycine (mmol/l) | 4°C | | | NA | NA | NA | NA | NA |
| Glycine (mmol/l) | 21°C | | | NA | NA | NA | NA | NA |
| *Branched-chain amino acids* | | | | | | | | |
| Isoleucine (mmol/l) | 4°C | 69 | 23 | 8.2e-04 | -1.9e-04 | 1.8e-03 | 1.1e-01 | 2.4e-02 |
| Isoleucine (mmol/l) | 21°C | 69 | 23 | -1.4e-04 | -1.3e-03 | 1.0e-03 | 8.1e-01 | 2.4e-02 |
| Leucine (mmol/l) | 4°C | 69 | 23 | 3.8e-03 | 3.3e-03 | 4.3e-03 | 0.0e+00 | 2.2e-02 |
| Leucine (mmol/l) | 21°C | 69 | 23 | 6.9e-03 | 5.9e-03 | 7.9e-03 | 0.0e+00 | 2.2e-02 |
| Valine (mmol/l) | 4°C | 69 | 23 | 4.0e-03 | 2.9e-03 | 5.1e-03 | 5.7e-13 | 3.7e-02 |
| Valine (mmol/l) | 21°C | 69 | 23 | 9.8e-03 | 8.4e-03 | 1.1e-02 | 0.0e+00 | 3.7e-02 |
| *Aromatic amino acids* | | | | | | | | |
| Phenylalanine (mmol/l) | 4°C | 69 | 23 | 8.3e-04 | -1.4e-05 | 1.7e-03 | 5.4e-02 | 6.2e-03 |
| Phenylalanine (mmol/l) | 21°C | 69 | 23 | 4.2e-03 | 3.2e-03 | 5.2e-03 | 4.4e-16 | 6.2e-03 |
| Tyrosine (mmol/l) | 4°C | 69 | 23 | 4.4e-04 | -2.8e-04 | 1.2e-03 | 2.3e-01 | 1.2e-02 |

| Metabolic traits | temperature | N.obs | N.indiv | Beta | LCI | UCI | Pvalue | SD |
|---|---|---|---|---|---|---|---|---|
| Tyrosine (mmol/l) | 21°C | 69 | 23 | 2.3e-03 | 1.7e-03 | 3.0e-03 | 1.3e-11 | 1.2e-02 |
| **Ketone bodies** | | | | | | | | |
| Acetate (mmol/l) | 4°C | 69 | 23 | -8.0e-03 | -9.7e-03 | -6.2e-03 | 0.0e+00 | 9.4e-03 |
| Acetate (mmol/l) | 21°C | 69 | 23 | -7.5e-03 | -1.0e-02 | -5.0e-03 | 6.5e-09 | 9.4e-03 |
| Beta-hydroxybutyrate (mmol/l) | 4°C | 69 | 23 | -5.4e-04 | -1.5e-03 | 4.3e-04 | 2.7e-01 | 1.8e-02 |
| Beta-hydroxybutyrate (mmol/l) | 21°C | 69 | 23 | 2.9e-03 | 4.2e-04 | 5.3e-03 | 2.2e-02 | 1.8e-02 |
| **Fluid balance** | | | | | | | | |
| Creatinine (mmol/l) | 4°C | 69 | 23 | -5.6e-04 | -1.2e-03 | 9.9e-05 | 9.6e-02 | 9.1e-03 |
| Creatinine (mmol/l) | 21°C | 69 | 23 | -4.1e-04 | -1.1e-03 | 3.1e-04 | 2.6e-01 | 9.1e-03 |
| Albumin (signal area) | 4°C | 69 | 23 | -4.8e-04 | -1.0e-03 | 7.2e-05 | 8.8e-02 | 4.3e-03 |
| Albumin (signal area) | 21°C | 69 | 23 | 1.8e-03 | 1.3e-03 | 2.3e-03 | 2.3e-12 | 4.3e-03 |
| **Inflammation** | | | | | | | | |
| Glycoprotein acetyls (mmol/l) | 4°C | 69 | 23 | -1.0e-02 | -1.8e-02 | -3.0e-03 | 5.8e-03 | 3.2e-01 |
| Glycoprotein acetyls (mmol/l) | 21°C | 69 | 23 | 1.8e-02 | 7.5e-03 | 2.8e-02 | 7.4e-04 | 3.2e-01 |

*sTable 4. Serum, post-storage handling effects: mean differences in metabolite concentrations (or trait value) comparing delays in sample preparation (i.e. thaw to buffer addition delay) and NMR profiling (buffer addition to NMR profiling delay) to the reference, for serum samples.*

*# Associations in sFigure3 and Figures 3 and 4 are presented in SD-units.* These SD point estimate can be obtained by dividing the point estimate (beta) in absolute (clinically meaningful) concentration by the metabolic trait standard deviation (SD), both provided in the below table.

***Abbreviations**: **C**=cholesterol; **IDL**=intermediate-density lipoprotein; **LCI**=lower confidence interval; **LDL**=low-density lipoprotein; **HDL**=high-density lipoprotein; **MUFA**=monounsaturated fatty acids; **N.obs**= number of observations (samples); **N.indiv**=number of individuals; **PUFA**=polyunsaturated fatty acids; **SD**=standard deviation; **UCI**= upper confidence interval; **VLDL**=very-low-density lipoprotein.*

| Metabolic traits | delay | N.obs | N.indiv | Beta | LCI | UCI | Pvalue | SD |
|---|---|---|---|---|---|---|---|---|
| **Lipoprotein subclasses** | | | | | | | | |
| *Extremely large VLDL* | | | | | | | | |
| Particle concentration (mol/l) | thaw-buffer | 50 | 25 | -1.9e-11 | -3.1e-11 | -6.8e-12 | 2.2e-03 | 2.1e-10 |
| Particle concentration (mol/l) | buffer-NMR | 50 | 25 | -6.8e-12 | -1.3e-11 | -1.0e-13 | 4.7e-02 | 2.1e-10 |
| Total lipids (mmol/l) | thaw-buffer | 50 | 25 | -4.0e-03 | -6.6e-03 | -1.4e-03 | 2.3e-03 | 4.6e-02 |
| Total lipids (mmol/l) | buffer-NMR | 50 | 25 | -1.5e-03 | -2.9e-03 | -5.7e-05 | 4.1e-02 | 4.6e-02 |
| Phospholipids (mmol/l) | thaw-buffer | 50 | 25 | -5.3e-04 | -8.3e-04 | -2.3e-04 | 5.7e-04 | 5.7e-03 |
| Phospholipids (mmol/l) | buffer-NMR | 50 | 25 | -2.1e-04 | -3.8e-04 | -4.0e-05 | 1.6e-02 | 5.7e-03 |
| Total cholesterol (mmol/l) | thaw-buffer | 50 | 25 | -5.2e-04 | -9.8e-04 | -6.4e-05 | 2.6e-02 | 8.5e-03 |
| Total cholesterol (mmol/l) | buffer-NMR | 50 | 25 | -3.1e-04 | -5.7e-04 | -5.5e-05 | 1.7e-02 | 8.5e-03 |
| Cholesterol esters (mmol/l) | thaw-buffer | 50 | 25 | -1.9e-04 | -4.9e-04 | 1.1e-04 | 2.1e-01 | 4.7e-03 |
| Cholesterol esters (mmol/l) | buffer-NMR | 50 | 25 | -1.7e-04 | -3.4e-04 | -6.0e-06 | 4.2e-02 | 4.7e-03 |
| Free cholesterol (mmol/l) | thaw-buffer | 50 | 25 | -3.3e-04 | -5.1e-04 | -1.6e-04 | 1.6e-04 | 3.8e-03 |
| Free cholesterol (mmol/l) | buffer-NMR | 50 | 25 | -1.4e-04 | -2.4e-04 | -4.1e-05 | 5.7e-03 | 3.8e-03 |
| Triglycerides (mmol/l) | thaw-buffer | 50 | 25 | -3.0e-03 | -4.8e-03 | -1.1e-03 | 1.6e-03 | 3.1e-02 |
| Triglycerides (mmol/l) | buffer-NMR | 50 | 25 | -9.6e-04 | -2.0e-03 | 5.0e-05 | 6.3e-02 | 3.1e-02 |
| *Very large VLDL* | | | | | | | | |
| Particle concentration (mol/l) | thaw-buffer | 50 | 25 | -9.1e-11 | -1.4e-10 | -4.2e-11 | 2.8e-04 | 1.3e-09 |
| Particle concentration (mol/l) | buffer-NMR | 50 | 25 | -3.9e-11 | -6.6e-11 | -1.2e-11 | 4.9e-03 | 1.3e-09 |
| Total lipids (mmol/l) | thaw-buffer | 50 | 25 | -9.1e-03 | -1.4e-02 | -4.2e-03 | 2.5e-04 | 1.2e-01 |

| Metabolic traits | delay | N.obs | N.indiv | Beta | LCI | UCI | Pvalue | SD |
|---|---|---|---|---|---|---|---|---|
| Total lipids (mmol/l) | buffer-NMR | 50 | 25 | -3.9e-03 | -6.5e-03 | -1.2e-03 | 4.5e-03 | 1.2e-01 |
| Phospholipids (mmol/l) | thaw-buffer | 50 | 25 | -1.7e-03 | -2.5e-03 | -8.5e-04 | 5.9e-05 | 2.1e-02 |
| Phospholipids (mmol/l) | buffer-NMR | 50 | 25 | -7.1e-04 | -1.2e-03 | -2.4e-04 | 3.4e-03 | 2.1e-02 |
| Total cholesterol (mmol/l) | thaw-buffer | 50 | 25 | -1.9e-03 | -3.0e-03 | -7.4e-04 | 1.3e-03 | 2.5e-02 |
| Total cholesterol (mmol/l) | buffer-NMR | 50 | 25 | -8.9e-04 | -1.5e-03 | -2.6e-04 | 5.3e-03 | 2.5e-02 |
| Cholesterol esters (mmol/l) | thaw-buffer | 50 | 25 | -8.3e-04 | -1.5e-03 | -2.0e-04 | 9.8e-03 | 1.4e-02 |
| Cholesterol esters (mmol/l) | buffer-NMR | 50 | 25 | -4.6e-04 | -8.0e-04 | -1.1e-04 | 8.9e-03 | 1.4e-02 |
| Free cholesterol (mmol/l) | thaw-buffer | 50 | 25 | -1.1e-03 | -1.6e-03 | -5.2e-04 | 1.0e-04 | 1.1e-02 |
| Free cholesterol (mmol/l) | buffer-NMR | 50 | 25 | -4.3e-04 | -7.2e-04 | -1.4e-04 | 3.4e-03 | 1.1e-02 |
| Triglycerides (mmol/l) | thaw-buffer | 50 | 25 | -5.5e-03 | -8.5e-03 | -2.5e-03 | 3.4e-04 | 7.8e-02 |
| Triglycerides (mmol/l) | buffer-NMR | 50 | 25 | -2.3e-03 | -3.9e-03 | -6.5e-04 | 6.1e-03 | 7.8e-02 |
| *Large VLDL* | | | | | | | | |
| Particle concentration (mol/l) | thaw-buffer | 50 | 25 | -2.3e-10 | -4.5e-10 | -1.7e-11 | 3.4e-02 | 7.2e-09 |
| Particle concentration (mol/l) | buffer-NMR | 50 | 25 | -1.3e-10 | -2.7e-10 | 2.8e-12 | 5.5e-02 | 7.2e-09 |
| Total lipids (mmol/l) | thaw-buffer | 50 | 25 | -1.4e-02 | -2.6e-02 | -1.8e-03 | 2.5e-02 | 4.2e-01 |
| Total lipids (mmol/l) | buffer-NMR | 50 | 25 | -8.1e-03 | -1.6e-02 | -6.4e-05 | 4.8e-02 | 4.2e-01 |
| Phospholipids (mmol/l) | thaw-buffer | 50 | 25 | -2.5e-03 | -4.7e-03 | -2.1e-04 | 3.2e-02 | 7.6e-02 |
| Phospholipids (mmol/l) | buffer-NMR | 50 | 25 | -1.5e-03 | -2.9e-03 | -5.5e-06 | 4.9e-02 | 7.6e-02 |
| Total cholesterol (mmol/l) | thaw-buffer | 50 | 25 | -4.7e-03 | -7.8e-03 | -1.5e-03 | 3.5e-03 | 9.7e-02 |
| Total cholesterol (mmol/l) | buffer-NMR | 50 | 25 | -2.3e-03 | -4.3e-03 | -3.4e-04 | 2.2e-02 | 9.7e-02 |
| Cholesterol esters (mmol/l) | thaw-buffer | 50 | 25 | -2.4e-03 | -4.2e-03 | -5.6e-04 | 1.1e-02 | 4.8e-02 |
| Cholesterol esters (mmol/l) | buffer-NMR | 50 | 25 | -1.1e-03 | -2.1e-03 | -7.1e-05 | 3.6e-02 | 4.8e-02 |
| Free cholesterol (mmol/l) | thaw-buffer | 50 | 25 | -2.3e-03 | -3.6e-03 | -8.9e-04 | 1.2e-03 | 4.9e-02 |
| Free cholesterol (mmol/l) | buffer-NMR | 50 | 25 | -1.2e-03 | -2.2e-03 | -2.4e-04 | 1.5e-02 | 4.9e-02 |
| Triglycerides (mmol/l) | thaw-buffer | 50 | 25 | -7.0e-03 | -1.4e-02 | 3.6e-04 | 6.2e-02 | 2.5e-01 |
| Triglycerides (mmol/l) | buffer-NMR | 50 | 25 | -4.3e-03 | -9.0e-03 | 3.6e-04 | 7.1e-02 | 2.5e-01 |
| *Medium VLDL* | | | | | | | | |

| Metabolic traits | delay | N.obs | N.indiv | Beta | LCI | UCI | Pvalue | SD |
|---|---|---|---|---|---|---|---|---|
| Particle concentration (mol/l) | thaw-buffer | 50 | 25 | -1.0e-10 | -5.4e-10 | 3.3e-10 | 6.4e-01 | 1.8e-08 |
| Particle concentration (mol/l) | buffer-NMR | 50 | 25 | -1.7e-10 | -5.2e-10 | 1.7e-10 | 3.3e-01 | 1.8e-08 |
| Total lipids (mmol/l) | thaw-buffer | 50 | 25 | -5.0e-03 | -2.0e-02 | 9.7e-03 | 5.0e-01 | 5.9e-01 |
| Total lipids (mmol/l) | buffer-NMR | 50 | 25 | -6.1e-03 | -1.7e-02 | 5.4e-03 | 3.0e-01 | 5.9e-01 |
| Phospholipids (mmol/l) | thaw-buffer | 50 | 25 | -1.1e-03 | -3.9e-03 | 1.7e-03 | 4.5e-01 | 1.1e-01 |
| Phospholipids (mmol/l) | buffer-NMR | 50 | 25 | -1.3e-03 | -3.5e-03 | 9.1e-04 | 2.5e-01 | 1.1e-01 |
| Total cholesterol (mmol/l) | thaw-buffer | 50 | 25 | -5.6e-03 | -1.0e-02 | -1.1e-03 | 1.4e-02 | 1.5e-01 |
| Total cholesterol (mmol/l) | buffer-NMR | 50 | 25 | -2.5e-03 | -5.5e-03 | 5.8e-04 | 1.1e-01 | 1.5e-01 |
| Cholesterol esters (mmol/l) | thaw-buffer | 50 | 25 | -4.7e-03 | -7.5e-03 | -1.9e-03 | 9.8e-04 | 7.4e-02 |
| Cholesterol esters (mmol/l) | buffer-NMR | 50 | 25 | -1.7e-03 | -3.5e-03 | 1.5e-04 | 7.3e-02 | 7.4e-02 |
| Free cholesterol (mmol/l) | thaw-buffer | 50 | 25 | -8.9e-04 | -2.7e-03 | 9.1e-04 | 3.3e-01 | 7.3e-02 |
| Free cholesterol (mmol/l) | buffer-NMR | 50 | 25 | -7.8e-04 | -2.2e-03 | 6.4e-04 | 2.8e-01 | 7.3e-02 |
| Triglycerides (mmol/l) | thaw-buffer | 50 | 25 | 1.7e-03 | -6.1e-03 | 9.5e-03 | 6.7e-01 | 3.3e-01 |
| Triglycerides (mmol/l) | buffer-NMR | 50 | 25 | -2.4e-03 | -8.9e-03 | 4.2e-03 | 4.8e-01 | 3.3e-01 |
| *Small VLDL* | | | | | | | | |
| Particle concentration (mol/l) | thaw-buffer | 50 | 25 | -7.3e-10 | -1.1e-09 | -3.7e-10 | 7.2e-05 | 1.7e-08 |
| Particle concentration (mol/l) | buffer-NMR | 50 | 25 | -3.2e-11 | -3.6e-10 | 3.0e-10 | 8.5e-01 | 1.7e-08 |
| Total lipids (mmol/l) | thaw-buffer | 50 | 25 | -1.7e-02 | -2.4e-02 | -1.0e-02 | 1.4e-06 | 3.2e-01 |
| Total lipids (mmol/l) | buffer-NMR | 50 | 25 | -8.8e-04 | -7.3e-03 | 5.5e-03 | 7.9e-01 | 3.2e-01 |
| Phospholipids (mmol/l) | thaw-buffer | 50 | 25 | -4.7e-03 | -6.2e-03 | -3.1e-03 | 4.5e-09 | 6.5e-02 |
| Phospholipids (mmol/l) | buffer-NMR | 50 | 25 | -3.0e-04 | -1.9e-03 | 1.3e-03 | 7.2e-01 | 6.5e-02 |
| Total cholesterol (mmol/l) | thaw-buffer | 50 | 25 | -1.4e-02 | -1.7e-02 | -1.0e-02 | 2.0e-15 | 8.7e-02 |
| Total cholesterol (mmol/l) | buffer-NMR | 50 | 25 | -1.2e-03 | -4.1e-03 | 1.7e-03 | 4.1e-01 | 8.7e-02 |
| Cholesterol esters (mmol/l) | thaw-buffer | 50 | 25 | -1.1e-02 | -1.4e-02 | -8.2e-03 | 4.4e-15 | 4.7e-02 |
| Cholesterol esters (mmol/l) | buffer-NMR | 50 | 25 | -1.1e-03 | -3.4e-03 | 1.3e-03 | 3.8e-01 | 4.7e-02 |
| Free cholesterol (mmol/l) | thaw-buffer | 50 | 25 | -2.8e-03 | -3.6e-03 | -1.9e-03 | 1.2e-09 | 4.1e-02 |
| Free cholesterol (mmol/l) | buffer-NMR | 50 | 25 | -1.8e-04 | -1.0e-03 | 6.6e-04 | 6.8e-01 | 4.1e-02 |

| Metabolic traits | delay | N.obs | N.indiv | Beta | LCI | UCI | Pvalue | SD |
|---|---|---|---|---|---|---|---|---|
| Triglycerides (mmol/l) | thaw-buffer | 50 | 25 | 1.2e-03 | -2.2e-03 | 4.6e-03 | 4.9e-01 | 1.7e-01 |
| Triglycerides (mmol/l) | buffer-NMR | 50 | 25 | 5.8e-04 | -2.6e-03 | 3.8e-03 | 7.2e-01 | 1.7e-01 |
| *Very Small VLDL* | | | | | | | | |
| Particle concentration (mol/l) | thaw-buffer | 50 | 25 | -1.4e-09 | -2.1e-09 | -7.4e-10 | 3.7e-05 | 9.1e-09 |
| Particle concentration (mol/l) | buffer-NMR | 50 | 25 | 1.4e-10 | -2.3e-10 | 5.0e-10 | 4.6e-01 | 9.1e-09 |
| Total lipids (mmol/l) | thaw-buffer | 50 | 25 | -2.0e-02 | -2.9e-02 | -1.1e-02 | 2.2e-05 | 1.1e-01 |
| Total lipids (mmol/l) | buffer-NMR | 50 | 25 | 1.6e-03 | -3.6e-03 | 6.7e-03 | 5.5e-01 | 1.1e-01 |
| Phospholipids (mmol/l) | thaw-buffer | 50 | 25 | 4.2e-04 | -2.5e-03 | 3.3e-03 | 7.8e-01 | 3.0e-02 |
| Phospholipids (mmol/l) | buffer-NMR | 50 | 25 | -1.4e-03 | -2.8e-03 | 1.3e-04 | 7.4e-02 | 3.0e-02 |
| Total cholesterol (mmol/l) | thaw-buffer | 50 | 25 | -2.1e-02 | -2.8e-02 | -1.5e-02 | 3.2e-10 | 4.6e-02 |
| Total cholesterol (mmol/l) | buffer-NMR | 50 | 25 | 2.3e-03 | -2.2e-03 | 6.7e-03 | 3.2e-01 | 4.6e-02 |
| Cholesterol esters (mmol/l) | thaw-buffer | 50 | 25 | -1.9e-02 | -2.4e-02 | -1.4e-02 | 2.0e-13 | 3.2e-02 |
| Cholesterol esters (mmol/l) | buffer-NMR | 50 | 25 | 2.6e-03 | -1.0e-03 | 6.2e-03 | 1.6e-01 | 3.2e-02 |
| Free cholesterol (mmol/l) | thaw-buffer | 50 | 25 | -2.1e-03 | -3.7e-03 | -4.1e-04 | 1.5e-02 | 1.4e-02 |
| Free cholesterol (mmol/l) | buffer-NMR | 50 | 25 | -3.2e-04 | -1.3e-03 | 6.3e-04 | 5.1e-01 | 1.4e-02 |
| Triglycerides (mmol/l) | thaw-buffer | 50 | 25 | 7.0e-04 | -2.1e-04 | 1.6e-03 | 1.3e-01 | 4.8e-02 |
| Triglycerides (mmol/l) | buffer-NMR | 50 | 25 | 6.8e-04 | -2.1e-04 | 1.6e-03 | 1.3e-01 | 4.8e-02 |
| *IDL* | | | | | | | | |
| Particle concentration (mol/l) | thaw-buffer | 50 | 25 | -8.4e-10 | -3.1e-09 | 1.4e-09 | 4.6e-01 | 2.0e-08 |
| Particle concentration (mol/l) | buffer-NMR | 50 | 25 | -4.8e-10 | -1.8e-09 | 8.6e-10 | 4.9e-01 | 2.0e-08 |
| Total lipids (mmol/l) | thaw-buffer | 50 | 25 | -9.7e-03 | -3.3e-02 | 1.4e-02 | 4.2e-01 | 2.0e-01 |
| Total lipids (mmol/l) | buffer-NMR | 50 | 25 | -5.3e-03 | -2.0e-02 | 9.1e-03 | 4.7e-01 | 2.0e-01 |
| Phospholipids (mmol/l) | thaw-buffer | 50 | 25 | -8.4e-04 | -6.3e-03 | 4.6e-03 | 7.6e-01 | 5.1e-02 |
| Phospholipids (mmol/l) | buffer-NMR | 50 | 25 | -2.3e-03 | -5.3e-03 | 7.5e-04 | 1.4e-01 | 5.1e-02 |
| Total cholesterol (mmol/l) | thaw-buffer | 50 | 25 | -1.2e-02 | -3.0e-02 | 6.0e-03 | 1.9e-01 | 1.3e-01 |
| Total cholesterol (mmol/l) | buffer-NMR | 50 | 25 | -3.5e-03 | -1.5e-02 | 8.1e-03 | 5.6e-01 | 1.3e-01 |
| Cholesterol esters (mmol/l) | thaw-buffer | 50 | 25 | -1.1e-02 | -2.5e-02 | 2.1e-03 | 1.0e-01 | 9.8e-02 |

| Metabolic traits | delay | N.obs | N.indiv | Beta | LCI | UCI | Pvalue | SD |
|---|---|---|---|---|---|---|---|---|
| Cholesterol esters (mmol/l) | buffer-NMR | 50 | 25 | -2.0e-03 | -1.1e-02 | 7.0e-03 | 6.6e-01 | 9.8e-02 |
| Free cholesterol (mmol/l) | thaw-buffer | 50 | 25 | -5.4e-04 | -5.1e-03 | 4.0e-03 | 8.1e-01 | 4.1e-02 |
| Free cholesterol (mmol/l) | buffer-NMR | 50 | 25 | -1.4e-03 | -4.1e-03 | 1.2e-03 | 3.0e-01 | 4.1e-02 |
| Triglycerides (mmol/l) | thaw-buffer | 50 | 25 | 2.8e-03 | 1.7e-03 | 4.0e-03 | 1.6e-06 | 3.2e-02 |
| Triglycerides (mmol/l) | buffer-NMR | 50 | 25 | 4.8e-04 | -3.7e-04 | 1.3e-03 | 2.7e-01 | 3.2e-02 |
| *Large LDL* | | | | | | | | |
| Particle concentration (mol/l) | thaw-buffer | 50 | 25 | 9.8e-10 | -2.2e-09 | 4.2e-09 | 5.5e-01 | 3.4e-08 |
| Particle concentration (mol/l) | buffer-NMR | 50 | 25 | -1.3e-09 | -3.2e-09 | 6.1e-10 | 1.8e-01 | 3.4e-08 |
| Total lipids (mmol/l) | thaw-buffer | 50 | 25 | 6.7e-03 | -1.7e-02 | 3.0e-02 | 5.7e-01 | 2.5e-01 |
| Total lipids (mmol/l) | buffer-NMR | 50 | 25 | -9.1e-03 | -2.3e-02 | 5.0e-03 | 2.1e-01 | 2.5e-01 |
| Phospholipids (mmol/l) | thaw-buffer | 50 | 25 | 1.6e-04 | -4.6e-03 | 4.9e-03 | 9.5e-01 | 5.1e-02 |
| Phospholipids (mmol/l) | buffer-NMR | 50 | 25 | -2.2e-03 | -5.2e-03 | 8.5e-04 | 1.6e-01 | 5.1e-02 |
| Total cholesterol (mmol/l) | thaw-buffer | 50 | 25 | 5.0e-03 | -1.3e-02 | 2.3e-02 | 5.8e-01 | 1.8e-01 |
| Total cholesterol (mmol/l) | buffer-NMR | 50 | 25 | -6.7e-03 | -1.8e-02 | 4.4e-03 | 2.3e-01 | 1.8e-01 |
| Cholesterol esters (mmol/l) | thaw-buffer | 50 | 25 | 4.8e-03 | -8.7e-03 | 1.8e-02 | 4.9e-01 | 1.4e-01 |
| Cholesterol esters (mmol/l) | buffer-NMR | 50 | 25 | -5.4e-03 | -1.4e-02 | 3.0e-03 | 2.1e-01 | 1.4e-01 |
| Free cholesterol (mmol/l) | thaw-buffer | 50 | 25 | 2.5e-04 | -4.2e-03 | 4.7e-03 | 9.1e-01 | 4.7e-02 |
| Free cholesterol (mmol/l) | buffer-NMR | 50 | 25 | -1.3e-03 | -4.0e-03 | 1.4e-03 | 3.4e-01 | 4.7e-02 |
| Triglycerides (mmol/l) | thaw-buffer | 50 | 25 | 1.3e-03 | 1.6e-04 | 2.5e-03 | 2.5e-02 | 2.6e-02 |
| Triglycerides (mmol/l) | buffer-NMR | 50 | 25 | -2.7e-04 | -1.3e-03 | 7.2e-04 | 6.0e-01 | 2.6e-02 |
| *Medium LDL* | | | | | | | | |
| Particle concentration (mol/l) | thaw-buffer | 50 | 25 | 4.2e-09 | 1.6e-09 | 6.7e-09 | 1.3e-03 | 3.0e-08 |
| Particle concentration (mol/l) | buffer-NMR | 50 | 25 | -1.0e-09 | -2.6e-09 | 5.4e-10 | 2.0e-01 | 3.0e-08 |
| Total lipids (mmol/l) | thaw-buffer | 50 | 25 | 2.0e-02 | 7.1e-03 | 3.4e-02 | 2.6e-03 | 1.5e-01 |
| Total lipids (mmol/l) | buffer-NMR | 50 | 25 | -5.0e-03 | -1.3e-02 | 3.1e-03 | 2.3e-01 | 1.5e-01 |
| Phospholipids (mmol/l) | thaw-buffer | 50 | 25 | 2.1e-03 | -4.3e-04 | 4.7e-03 | 1.0e-01 | 3.4e-02 |
| Phospholipids (mmol/l) | buffer-NMR | 50 | 25 | -7.0e-04 | -2.4e-03 | 1.0e-03 | 4.3e-01 | 3.4e-02 |

| Metabolic traits | delay | N.obs | N.indiv | Beta | LCI | UCI | Pvalue | SD |
|---|---|---|---|---|---|---|---|---|
| Total cholesterol (mmol/l) | thaw-buffer | 50 | 25 | 1.7e-02 | 6.1e-03 | 2.7e-02 | 2.1e-03 | 1.1e-01 |
| Total cholesterol (mmol/l) | buffer-NMR | 50 | 25 | -3.8e-03 | -1.0e-02 | 2.7e-03 | 2.5e-01 | 1.1e-01 |
| Cholesterol esters (mmol/l) | thaw-buffer | 50 | 25 | 1.6e-02 | 7.2e-03 | 2.4e-02 | 3.1e-04 | 9.3e-02 |
| Cholesterol esters (mmol/l) | buffer-NMR | 50 | 25 | -3.3e-03 | -8.4e-03 | 1.9e-03 | 2.1e-01 | 9.3e-02 |
| Free cholesterol (mmol/l) | thaw-buffer | 50 | 25 | 9.3e-04 | -1.2e-03 | 3.0e-03 | 3.8e-01 | 2.2e-02 |
| Free cholesterol (mmol/l) | buffer-NMR | 50 | 25 | -5.3e-04 | -1.9e-03 | 8.7e-04 | 4.6e-01 | 2.2e-02 |
| Triglycerides (mmol/l) | thaw-buffer | 50 | 25 | 1.6e-03 | 8.3e-04 | 2.3e-03 | 3.8e-05 | 1.3e-02 |
| Triglycerides (mmol/l) | buffer-NMR | 50 | 25 | -5.7e-04 | -1.4e-03 | 3.0e-04 | 2.0e-01 | 1.3e-02 |
| *Small LDL* | | | | | | | | |
| Particle concentration (mol/l) | thaw-buffer | 50 | 25 | 4.9e-09 | 2.0e-09 | 7.8e-09 | 9.8e-04 | 3.4e-08 |
| Particle concentration (mol/l) | buffer-NMR | 50 | 25 | -1.4e-09 | -3.3e-09 | 4.8e-10 | 1.4e-01 | 3.4e-08 |
| Total lipids (mmol/l) | thaw-buffer | 50 | 25 | 1.3e-02 | 4.9e-03 | 2.2e-02 | 1.9e-03 | 9.6e-02 |
| Total lipids (mmol/l) | buffer-NMR | 50 | 25 | -3.8e-03 | -9.3e-03 | 1.6e-03 | 1.7e-01 | 9.6e-02 |
| Phospholipids (mmol/l) | thaw-buffer | 50 | 25 | 1.9e-03 | 8.9e-05 | 3.7e-03 | 4.0e-02 | 2.4e-02 |
| Phospholipids (mmol/l) | buffer-NMR | 50 | 25 | -9.0e-04 | -2.2e-03 | 4.3e-04 | 1.8e-01 | 2.4e-02 |
| Total cholesterol (mmol/l) | thaw-buffer | 50 | 25 | 1.1e-02 | 4.2e-03 | 1.7e-02 | 1.2e-03 | 7.1e-02 |
| Total cholesterol (mmol/l) | buffer-NMR | 50 | 25 | -2.4e-03 | -6.5e-03 | 1.7e-03 | 2.4e-01 | 7.1e-02 |
| Cholesterol esters (mmol/l) | thaw-buffer | 50 | 25 | 1.0e-02 | 5.2e-03 | 1.5e-02 | 8.0e-05 | 5.8e-02 |
| Cholesterol esters (mmol/l) | buffer-NMR | 50 | 25 | -2.0e-03 | -5.1e-03 | 1.1e-03 | 2.1e-01 | 5.8e-02 |
| Free cholesterol (mmol/l) | thaw-buffer | 50 | 25 | 4.0e-04 | -9.8e-04 | 1.8e-03 | 5.7e-01 | 1.4e-02 |
| Free cholesterol (mmol/l) | buffer-NMR | 50 | 25 | -4.5e-04 | -1.5e-03 | 5.5e-04 | 3.8e-01 | 1.4e-02 |
| Triglycerides (mmol/l) | thaw-buffer | 50 | 25 | 6.4e-04 | 2.3e-04 | 1.0e-03 | 2.2e-03 | 1.3e-02 |
| Triglycerides (mmol/l) | buffer-NMR | 50 | 25 | -4.8e-04 | -9.4e-04 | -1.8e-05 | 4.2e-02 | 1.3e-02 |
| *Very large HDL* | | | | | | | | |
| Particle concentration (mol/l) | thaw-buffer | 50 | 25 | -6.0e-08 | -7.3e-08 | -4.8e-08 | 0.0e+00 | 2.4e-07 |
| Particle concentration (mol/l) | buffer-NMR | 50 | 25 | -2.3e-08 | -3.2e-08 | -1.4e-08 | 1.4e-06 | 2.4e-07 |
| Total lipids (mmol/l) | thaw-buffer | 50 | 25 | -6.4e-02 | -7.7e-02 | -5.1e-02 | 0.0e+00 | 2.4e-01 |

| Metabolic traits | delay | N.obs | N.indiv | Beta | LCI | UCI | Pvalue | SD |
|---|---|---|---|---|---|---|---|---|
| Total lipids (mmol/l) | buffer-NMR | 50 | 25 | -2.3e-02 | -3.3e-02 | -1.3e-02 | 3.6e-06 | 2.4e-01 |
| Phospholipids (mmol/l) | thaw-buffer | 50 | 25 | -2.3e-02 | -2.8e-02 | -1.7e-02 | 2.2e-16 | 1.4e-01 |
| Phospholipids (mmol/l) | buffer-NMR | 50 | 25 | -1.3e-02 | -1.8e-02 | -8.7e-03 | 6.4e-09 | 1.4e-01 |
| Total cholesterol (mmol/l) | thaw-buffer | 50 | 25 | -4.1e-02 | -4.8e-02 | -3.3e-02 | 0.0e+00 | 1.0e-01 |
| Total cholesterol (mmol/l) | buffer-NMR | 50 | 25 | -9.2e-03 | -1.5e-02 | -3.5e-03 | 1.6e-03 | 1.0e-01 |
| Cholesterol esters (mmol/l) | thaw-buffer | 50 | 25 | -2.9e-02 | -3.5e-02 | -2.4e-02 | 0.0e+00 | 7.3e-02 |
| Cholesterol esters (mmol/l) | buffer-NMR | 50 | 25 | -6.7e-03 | -1.1e-02 | -2.6e-03 | 1.3e-03 | 7.3e-02 |
| Free cholesterol (mmol/l) | thaw-buffer | 50 | 25 | -1.1e-02 | -1.3e-02 | -9.2e-03 | 0.0e+00 | 3.2e-02 |
| Free cholesterol (mmol/l) | buffer-NMR | 50 | 25 | -2.5e-03 | -4.2e-03 | -8.4e-04 | 3.2e-03 | 3.2e-02 |
| Triglycerides (mmol/l) | thaw-buffer | 50 | 25 | -6.4e-04 | -1.7e-03 | 3.9e-04 | 2.2e-01 | 9.9e-03 |
| Triglycerides (mmol/l) | buffer-NMR | 50 | 25 | -6.0e-04 | -1.2e-03 | 3.4e-05 | 6.3e-02 | 9.9e-03 |
| *Large HDL* | | | | | | | | |
| Particle concentration (mol/l) | thaw-buffer | 50 | 25 | -2.2e-08 | -4.6e-08 | 3.2e-09 | 8.7e-02 | 6.4e-07 |
| Particle concentration (mol/l) | buffer-NMR | 50 | 25 | -6.2e-08 | -8.1e-08 | -4.3e-08 | 1.6e-10 | 6.4e-07 |
| Total lipids (mmol/l) | thaw-buffer | 50 | 25 | -1.5e-02 | -3.1e-02 | 1.3e-03 | 7.1e-02 | 4.1e-01 |
| Total lipids (mmol/l) | buffer-NMR | 50 | 25 | -4.0e-02 | -5.2e-02 | -2.7e-02 | 2.2e-10 | 4.1e-01 |
| Phospholipids (mmol/l) | thaw-buffer | 50 | 25 | -8.4e-03 | -1.7e-02 | 4.2e-05 | 5.1e-02 | 1.8e-01 |
| Phospholipids (mmol/l) | buffer-NMR | 50 | 25 | -1.9e-02 | -2.5e-02 | -1.3e-02 | 4.8e-09 | 1.8e-01 |
| Total cholesterol (mmol/l) | thaw-buffer | 50 | 25 | -8.3e-03 | -1.7e-02 | 1.8e-04 | 5.5e-02 | 2.2e-01 |
| Total cholesterol (mmol/l) | buffer-NMR | 50 | 25 | -2.0e-02 | -2.7e-02 | -1.3e-02 | 4.0e-09 | 2.2e-01 |
| Cholesterol esters (mmol/l) | thaw-buffer | 50 | 25 | -6.3e-03 | -1.3e-02 | 1.0e-04 | 5.4e-02 | 1.7e-01 |
| Cholesterol esters (mmol/l) | buffer-NMR | 50 | 25 | -1.5e-02 | -2.0e-02 | -1.0e-02 | 3.5e-09 | 1.7e-01 |
| Free cholesterol (mmol/l) | thaw-buffer | 50 | 25 | -2.0e-03 | -4.1e-03 | 1.0e-04 | 6.2e-02 | 5.4e-02 |
| Free cholesterol (mmol/l) | buffer-NMR | 50 | 25 | -5.0e-03 | -6.7e-03 | -3.3e-03 | 1.1e-08 | 5.4e-02 |
| Triglycerides (mmol/l) | thaw-buffer | 50 | 25 | 2.0e-03 | 1.3e-03 | 2.7e-03 | 5.9e-09 | 1.3e-02 |
| Triglycerides (mmol/l) | buffer-NMR | 50 | 25 | -6.9e-04 | -1.1e-03 | -3.2e-04 | 3.2e-04 | 1.3e-02 |
| *Medium HDL* | | | | | | | | |

| Metabolic traits | delay | N.obs | N.indiv | Beta | LCI | UCI | Pvalue | SD |
|---|---|---|---|---|---|---|---|---|
| Particle concentration (mol/l) | thaw-buffer | 50 | 25 | -2.3e-08 | -7.6e-08 | 3.0e-08 | 3.9e-01 | 3.7e-07 |
| Particle concentration (mol/l) | buffer-NMR | 50 | 25 | -3.5e-08 | -7.2e-08 | 7.0e-10 | 5.5e-02 | 3.7e-07 |
| Total lipids (mmol/l) | thaw-buffer | 50 | 25 | -9.4e-03 | -3.3e-02 | 1.4e-02 | 4.3e-01 | 1.6e-01 |
| Total lipids (mmol/l) | buffer-NMR | 50 | 25 | -1.5e-02 | -3.1e-02 | 3.5e-04 | 5.5e-02 | 1.6e-01 |
| Phospholipids (mmol/l) | thaw-buffer | 50 | 25 | -7.8e-03 | -1.8e-02 | 2.1e-03 | 1.2e-01 | 7.3e-02 |
| Phospholipids (mmol/l) | buffer-NMR | 50 | 25 | -6.9e-03 | -1.4e-02 | -2.3e-04 | 4.3e-02 | 7.3e-02 |
| Total cholesterol (mmol/l) | thaw-buffer | 50 | 25 | -6.0e-04 | -1.4e-02 | 1.3e-02 | 9.3e-01 | 8.9e-02 |
| Total cholesterol (mmol/l) | buffer-NMR | 50 | 25 | -7.9e-03 | -1.7e-02 | 7.7e-04 | 7.4e-02 | 8.9e-02 |
| Cholesterol esters (mmol/l) | thaw-buffer | 50 | 25 | 7.3e-04 | -9.8e-03 | 1.1e-02 | 8.9e-01 | 6.9e-02 |
| Cholesterol esters (mmol/l) | buffer-NMR | 50 | 25 | -6.2e-03 | -1.3e-02 | 7.7e-04 | 8.1e-02 | 6.9e-02 |
| Free cholesterol (mmol/l) | thaw-buffer | 50 | 25 | -1.3e-03 | -4.1e-03 | 1.4e-03 | 3.5e-01 | 2.0e-02 |
| Free cholesterol (mmol/l) | buffer-NMR | 50 | 25 | -1.7e-03 | -3.5e-03 | 5.5e-05 | 5.8e-02 | 2.0e-02 |
| Triglycerides (mmol/l) | thaw-buffer | 50 | 25 | -1.1e-03 | -1.7e-03 | -4.9e-04 | 3.4e-04 | 1.9e-02 |
| Triglycerides (mmol/l) | buffer-NMR | 50 | 25 | -4.1e-04 | -1.0e-03 | 2.2e-04 | 2.0e-01 | 1.9e-02 |
| *Small HDL* | | | | | | | | |
| Particle concentration (mol/l) | thaw-buffer | 50 | 25 | 5.7e-08 | -2.7e-08 | 1.4e-07 | 1.9e-01 | 4.7e-07 |
| Particle concentration (mol/l) | buffer-NMR | 50 | 25 | -1.1e-08 | -6.8e-08 | 4.5e-08 | 7.0e-01 | 4.7e-07 |
| Total lipids (mmol/l) | thaw-buffer | 50 | 25 | 1.3e-02 | -6.0e-03 | 3.2e-02 | 1.8e-01 | 1.0e-01 |
| Total lipids (mmol/l) | buffer-NMR | 50 | 25 | -2.9e-03 | -1.6e-02 | 9.9e-03 | 6.5e-01 | 1.0e-01 |
| Phospholipids (mmol/l) | thaw-buffer | 50 | 25 | -4.6e-03 | -1.2e-02 | 2.5e-03 | 2.1e-01 | 7.2e-02 |
| Phospholipids (mmol/l) | buffer-NMR | 50 | 25 | -6.2e-04 | -4.6e-03 | 3.3e-03 | 7.6e-01 | 7.2e-02 |
| Total cholesterol (mmol/l) | thaw-buffer | 50 | 25 | 1.8e-02 | 5.4e-03 | 3.0e-02 | 4.8e-03 | 6.1e-02 |
| Total cholesterol (mmol/l) | buffer-NMR | 50 | 25 | -2.7e-03 | -1.2e-02 | 7.1e-03 | 5.9e-01 | 6.1e-02 |
| Cholesterol esters (mmol/l) | thaw-buffer | 50 | 25 | 1.9e-02 | 8.3e-03 | 3.0e-02 | 5.5e-04 | 6.1e-02 |
| Cholesterol esters (mmol/l) | buffer-NMR | 50 | 25 | -2.3e-03 | -1.2e-02 | 7.1e-03 | 6.3e-01 | 6.1e-02 |
| Free cholesterol (mmol/l) | thaw-buffer | 50 | 25 | -1.6e-03 | -3.1e-03 | -1.3e-04 | 3.3e-02 | 1.2e-02 |
| Free cholesterol (mmol/l) | buffer-NMR | 50 | 25 | -4.0e-04 | -1.1e-03 | 3.0e-04 | 2.6e-01 | 1.2e-02 |

| Metabolic traits | delay | N.obs | N.indiv | Beta | LCI | UCI | Pvalue | SD |
|---|---|---|---|---|---|---|---|---|
| Triglycerides (mmol/l) | thaw-buffer | 50 | 25 | 4.9e-05 | -5.3e-04 | 6.3e-04 | 8.7e-01 | 2.2e-02 |
| Triglycerides (mmol/l) | buffer-NMR | 50 | 25 | 4.4e-04 | -1.9e-04 | 1.1e-03 | 1.7e-01 | 2.2e-02 |
| **Lipoprotein particle size** | | | | | | | | |
| VLDL particle size (nm) | thaw-buffer | 50 | 25 | 1.0e-01 | 4.9e-02 | 1.6e-01 | 2.0e-04 | 1.6e+00 |
| VLDL particle size (nm) | buffer-NMR | 50 | 25 | -3.8e-02 | -8.2e-02 | 5.4e-03 | 8.6e-02 | 1.6e+00 |
| LDL particle size (nm) | thaw-buffer | 50 | 25 | -5.2e-02 | -7.0e-02 | -3.4e-02 | 9.2e-09 | 7.7e-02 |
| LDL particle size (nm) | buffer-NMR | 50 | 25 | 6.0e-03 | -5.7e-03 | 1.8e-02 | 3.1e-01 | 7.7e-02 |
| HDL particle size (nm) | thaw-buffer | 50 | 25 | -3.6e-02 | -5.2e-02 | -2.0e-02 | 6.8e-06 | 2.8e-01 |
| HDL particle size (nm) | buffer-NMR | 50 | 25 | -2.2e-02 | -3.4e-02 | -9.2e-03 | 6.6e-04 | 2.8e-01 |
| **Cholesterol** | | | | | | | | |
| Total cholesterol (mmol/l) | thaw-buffer | 50 | 25 | -5.9e-02 | -1.2e-01 | 1.6e-03 | 5.6e-02 | 7.3e-01 |
| Total cholesterol (mmol/l) | buffer-NMR | 50 | 25 | -6.1e-02 | -1.0e-01 | -1.8e-02 | 5.9e-03 | 7.3e-01 |
| VLDL cholesterol (mmol/l) | thaw-buffer | 50 | 25 | -4.7e-02 | -6.3e-02 | -3.2e-02 | 2.4e-09 | 3.8e-01 |
| VLDL cholesterol (mmol/l) | buffer-NMR | 50 | 25 | -4.8e-03 | -1.5e-02 | 5.1e-03 | 3.4e-01 | 3.8e-01 |
| Remnant cholesterol (mmol/l) | thaw-buffer | 50 | 25 | -5.9e-02 | -9.0e-02 | -2.8e-02 | 1.9e-04 | 4.4e-01 |
| Remnant cholesterol (mmol/l) | buffer-NMR | 50 | 25 | -8.3e-03 | -2.8e-02 | 1.1e-02 | 4.1e-01 | 4.4e-01 |
| LDL cholesterol (mmol/l) | thaw-buffer | 50 | 25 | 3.2e-02 | -2.4e-03 | 6.7e-02 | 6.8e-02 | 3.6e-01 |
| LDL cholesterol (mmol/l) | buffer-NMR | 50 | 25 | -1.3e-02 | -3.5e-02 | 8.5e-03 | 2.3e-01 | 3.6e-01 |
| HDL cholesterol (mmol/l) | thaw-buffer | 50 | 25 | -3.2e-02 | -5.7e-02 | -7.2e-03 | 1.1e-02 | 4.0e-01 |
| HDL cholesterol (mmol/l) | buffer-NMR | 50 | 25 | -4.0e-02 | -5.7e-02 | -2.3e-02 | 5.9e-06 | 4.0e-01 |
| HDL2 cholesterol (mmol/l) | thaw-buffer | 50 | 25 | -2.4e-02 | -4.6e-02 | -8.7e-04 | 4.2e-02 | 3.6e-01 |
| HDL2 cholesterol (mmol/l) | buffer-NMR | 50 | 25 | -3.6e-02 | -5.2e-02 | -2.0e-02 | 6.2e-06 | 3.6e-01 |
| HDL3 cholesterol (mmol/l) | thaw-buffer | 50 | 25 | -8.4e-03 | -1.1e-02 | -5.5e-03 | 7.3e-09 | 3.5e-02 |
| HDL3 cholesterol (mmol/l) | buffer-NMR | 50 | 25 | -3.9e-03 | -6.1e-03 | -1.7e-03 | 4.6e-04 | 3.5e-02 |
| Esterified cholesterol (mmol/l) | thaw-buffer | 42 | 21 | 4.0e-02 | -1.1e-02 | 9.2e-02 | 1.2e-01 | 5.1e-01 |
| Esterified cholesterol (mmol/l) | buffer-NMR | 42 | 21 | -1.1e-03 | -4.2e-02 | 4.0e-02 | 9.6e-01 | 5.1e-01 |
| Free cholesterol (mmol/l) | thaw-buffer | 42 | 21 | -6.6e-02 | -9.2e-02 | -4.0e-02 | 9.1e-07 | 2.2e-01 |

| Metabolic traits | delay | N.obs | N.indiv | Beta | LCI | UCI | Pvalue | SD |
|---|---|---|---|---|---|---|---|---|
| Free cholesterol (mmol/l) | buffer-NMR | 42 | 21 | -6.3e-02 | -8.4e-02 | -4.2e-02 | 4.8e-09 | 2.2e-01 |
| **Glycerides and phospholipids** | | | | | | | | |
| Triglycerides (mmol/l) | thaw-buffer | 50 | 25 | -6.7e-04 | -2.1e-02 | 2.0e-02 | 9.5e-01 | 1.0e+00 |
| Triglycerides (mmol/l) | buffer-NMR | 50 | 25 | -1.0e-02 | -2.8e-02 | 7.3e-03 | 2.5e-01 | 1.0e+00 |
| VLDL triglycerides (mmol/l) | thaw-buffer | 50 | 25 | -7.3e-03 | -2.7e-02 | 1.3e-02 | 4.7e-01 | 9.0e-01 |
| VLDL triglycerides (mmol/l) | buffer-NMR | 50 | 25 | -8.2e-03 | -2.5e-02 | 8.6e-03 | 3.4e-01 | 9.0e-01 |
| LDL triglycerides (mmol/l) | thaw-buffer | 50 | 25 | 3.5e-03 | 1.4e-03 | 5.7e-03 | 1.1e-03 | 5.2e-02 |
| LDL triglycerides (mmol/l) | buffer-NMR | 50 | 25 | -1.3e-03 | -3.6e-03 | 9.1e-04 | 2.5e-01 | 5.2e-02 |
| HDL triglycerides (mmol/l) | thaw-buffer | 50 | 25 | 3.6e-04 | -1.2e-03 | 1.9e-03 | 6.4e-01 | 5.0e-02 |
| HDL triglycerides (mmol/l) | buffer-NMR | 50 | 25 | -1.3e-03 | -2.3e-03 | -2.1e-04 | 1.9e-02 | 5.0e-02 |
| Diacylglycerol (mmol/l) | thaw-buffer | 40 | 20 | -2.4e-03 | -8.1e-03 | 3.2e-03 | 4.0e-01 | 2.3e-02 |
| Diacylglycerol (mmol/l) | buffer-NMR | 40 | 20 | -4.7e-04 | -7.0e-03 | 6.1e-03 | 8.9e-01 | 2.3e-02 |
| Phosphoglycerides (mmol/l) | thaw-buffer | 42 | 21 | -9.3e-02 | -1.3e-01 | -5.1e-02 | 1.1e-05 | 3.7e-01 |
| Phosphoglycerides (mmol/l) | buffer-NMR | 42 | 21 | 7.3e-03 | -3.5e-02 | 4.9e-02 | 7.3e-01 | 3.7e-01 |
| Phosphatidylcholine + other cholines (mmol/l) | thaw-buffer | 42 | 21 | -1.1e-01 | -1.5e-01 | -7.0e-02 | 6.0e-08 | 3.5e-01 |
| Phosphatidylcholine + other cholines (mmol/l) | buffer-NMR | 42 | 21 | -4.7e-02 | -8.2e-02 | -1.2e-02 | 7.9e-03 | 3.5e-01 |
| Sphingomyelins (mmol/l) | thaw-buffer | 42 | 21 | -2.1e-02 | -4.0e-02 | -9.4e-04 | 4.0e-02 | 7.3e-02 |
| Sphingomyelins (mmol/l) | buffer-NMR | 42 | 21 | 2.2e-02 | 2.9e-03 | 4.1e-02 | 2.4e-02 | 7.3e-02 |
| Cholines (mmol/l) | thaw-buffer | 42 | 21 | -8.2e-02 | -1.4e-01 | -2.2e-02 | 7.5e-03 | 3.7e-01 |
| Cholines (mmol/l) | buffer-NMR | 42 | 21 | 5.6e-03 | -5.8e-02 | 6.9e-02 | 8.6e-01 | 3.7e-01 |
| **Apolipoproteins** | | | | | | | | |
| Apolipoprotein A-I (g/l) | thaw-buffer | 50 | 25 | -9.6e-03 | -2.3e-02 | 4.2e-03 | 1.7e-01 | 1.9e-01 |
| Apolipoprotein A-I (g/l) | buffer-NMR | 50 | 25 | -2.6e-02 | -3.7e-02 | -1.6e-02 | 1.0e-06 | 1.9e-01 |
| Apolipoprotein B (g/l) | thaw-buffer | 50 | 25 | -9.6e-04 | -1.6e-02 | 1.4e-02 | 9.0e-01 | 2.3e-01 |
| Apolipoprotein B (g/l) | buffer-NMR | 50 | 25 | -3.6e-03 | -1.3e-02 | 5.9e-03 | 4.6e-01 | 2.3e-01 |
| **Fatty acids** | | | | | | | | |

| Metabolic traits | delay | N.obs | N.indiv | Beta | LCI | UCI | Pvalue | SD |
|---|---|---|---|---|---|---|---|---|
| Total fatty acids (mmol/l) | thaw-buffer | 42 | 21 | -4.5e-02 | -2.9e-01 | 2.0e-01 | 7.2e-01 | 3.3e+00 |
| Total fatty acids (mmol/l) | buffer-NMR | 42 | 21 | -1.1e-01 | -2.9e-01 | 6.5e-02 | 2.2e-01 | 3.3e+00 |
| Fatty acid chain length | thaw-buffer | 42 | 21 | -1.0e-02 | -5.0e-02 | 2.9e-02 | 6.0e-01 | 2.8e-01 |
| Fatty acid chain length | buffer-NMR | 42 | 21 | 1.1e-01 | 6.8e-02 | 1.5e-01 | 7.8e-08 | 2.8e-01 |
| Degree of unsaturation | thaw-buffer | 42 | 21 | -1.9e-02 | -3.1e-02 | -8.4e-03 | 5.4e-04 | 8.0e-02 |
| Degree of unsaturation | buffer-NMR | 42 | 21 | -1.0e-02 | -1.6e-02 | -4.2e-03 | 6.6e-04 | 8.0e-02 |
| Docosahexaenoic acid (mmol/l) | thaw-buffer | 42 | 21 | -3.7e-03 | -7.9e-03 | 5.7e-04 | 9.0e-02 | 4.9e-02 |
| Docosahexaenoic acid (mmol/l) | buffer-NMR | 42 | 21 | -1.4e-03 | -4.8e-03 | 2.0e-03 | 4.1e-01 | 4.9e-02 |
| Linoleic acid (mmol/l) | thaw-buffer | 42 | 21 | -9.0e-03 | -6.1e-02 | 4.3e-02 | 7.3e-01 | 6.1e-01 |
| Linoleic acid (mmol/l) | buffer-NMR | 42 | 21 | -2.5e-02 | -6.4e-02 | 1.5e-02 | 2.2e-01 | 6.1e-01 |
| Conjugated linoleic acid (mmol/l) | thaw-buffer | 42 | 21 | -2.7e-04 | -6.6e-03 | 6.0e-03 | 9.3e-01 | 2.5e-02 |
| Conjugated linoleic acid (mmol/l) | buffer-NMR | 42 | 21 | 6.8e-03 | 2.3e-03 | 1.1e-02 | 2.8e-03 | 2.5e-02 |
| n-3 fatty acids (mmol/l) | thaw-buffer | 42 | 21 | 1.4e-03 | -1.2e-02 | 1.5e-02 | 8.4e-01 | 1.4e-01 |
| n-3 fatty acids (mmol/l) | buffer-NMR | 42 | 21 | -7.3e-03 | -1.6e-02 | 1.1e-03 | 8.8e-02 | 1.4e-01 |
| n-6 fatty acids (mmol/l) | thaw-buffer | 42 | 21 | -4.4e-02 | -1.1e-01 | 2.2e-02 | 1.9e-01 | 6.8e-01 |
| n-6 fatty acids (mmol/l) | buffer-NMR | 42 | 21 | -5.0e-02 | -1.0e-01 | -5.5e-04 | 4.8e-02 | 6.8e-01 |
| PUFA (mmol/l) | thaw-buffer | 42 | 21 | -4.3e-02 | -1.2e-01 | 3.3e-02 | 2.7e-01 | 8.0e-01 |
| PUFA (mmol/l) | buffer-NMR | 42 | 21 | -5.8e-02 | -1.1e-01 | -1.5e-03 | 4.4e-02 | 8.0e-01 |
| MUFA (mmol/l) | thaw-buffer | 42 | 21 | 1.0e-02 | -5.2e-02 | 7.3e-02 | 7.5e-01 | 1.4e+00 |
| MUFA (mmol/l) | buffer-NMR | 42 | 21 | 1.5e-02 | -4.0e-02 | 7.0e-02 | 6.0e-01 | 1.4e+00 |
| Saturated fatty acids (mmol/l) | thaw-buffer | 42 | 21 | -1.2e-02 | -1.5e-01 | 1.3e-01 | 8.7e-01 | 1.3e+00 |
| Saturated fatty acids (mmol/l) | buffer-NMR | 42 | 21 | -6.8e-02 | -1.5e-01 | 1.2e-02 | 9.4e-02 | 1.3e+00 |
| **Glycolysis related metabolites** | | | | | | | | |
| Glucose (mmol/l) | thaw-buffer | 50 | 25 | -7.4e-03 | -3.7e-02 | 2.3e-02 | 6.3e-01 | 4.9e-01 |
| Glucose (mmol/l) | buffer-NMR | 50 | 25 | -2.5e-02 | -5.0e-02 | 5.5e-04 | 5.5e-02 | 4.9e-01 |
| Lactate (mmol/l) | thaw-buffer | 50 | 25 | 2.0e-02 | 7.6e-04 | 3.9e-02 | 4.2e-02 | 3.9e-01 |

| Metabolic traits | delay | N.obs | N.indiv | Beta | LCI | UCI | Pvalue | SD |
|---|---|---|---|---|---|---|---|---|
| Lactate (mmol/l) | buffer-NMR | 50 | 25 | 9.5e-03 | -1.3e-02 | 3.2e-02 | 4.0e-01 | 3.9e-01 |
| Pyruvate (mmol/l) | thaw-buffer | 74 | 37 | -1.6e-03 | -3.2e-03 | -1.7e-05 | 4.8e-02 | 3.2e-02 |
| Pyruvate (mmol/l) | buffer-NMR | 74 | 37 | 5.7e-04 | -8.6e-04 | 2.0e-03 | 4.3e-01 | 3.2e-02 |
| Citrate (mmol/l) | thaw-buffer | 50 | 25 | -1.3e-03 | -1.2e-02 | 9.3e-03 | 8.1e-01 | 2.2e-02 |
| Citrate (mmol/l) | buffer-NMR | 50 | 25 | 2.3e-04 | -8.2e-03 | 8.7e-03 | 9.6e-01 | 2.2e-02 |
| Glycerol (mmol/l) | thaw-buffer | 74 | 37 | -1.4e-03 | -5.0e-03 | 2.3e-03 | 4.6e-01 | 2.1e-02 |
| Glycerol (mmol/l) | buffer-NMR | 74 | 37 | 3.6e-03 | 1.6e-03 | 5.5e-03 | 4.3e-04 | 2.1e-02 |
| **Amino acids** | | | | | | | | |
| Alanine (mmol/l) | thaw-buffer | 50 | 25 | -3.0e-03 | -6.1e-03 | -1.7e-05 | 4.9e-02 | 5.5e-02 |
| Alanine (mmol/l) | buffer-NMR | 50 | 25 | 5.6e-03 | 2.9e-03 | 8.4e-03 | 6.4e-05 | 5.5e-02 |
| Glutamine (mmol/l) | thaw-buffer | 50 | 25 | 1.1e-02 | 4.0e-03 | 1.8e-02 | 2.0e-03 | 5.6e-02 |
| Glutamine (mmol/l) | buffer-NMR | 50 | 25 | -9.4e-03 | -1.5e-02 | -4.0e-03 | 6.4e-04 | 5.6e-02 |
| Histidine (mmol/l) | thaw-buffer | 50 | 25 | -5.2e-03 | -7.9e-03 | -2.4e-03 | 2.2e-04 | 7.0e-03 |
| Histidine (mmol/l) | buffer-NMR | 50 | 25 | -9.4e-04 | -2.7e-03 | 7.7e-04 | 2.8e-01 | 7.0e-03 |
| Glycine (mmol/l) | thaw-buffer | 74 | 37 | 2.3e-02 | 1.8e-02 | 2.7e-02 | 0.0e+00 | 7.6e-02 |
| Glycine (mmol/l) | buffer-NMR | 74 | 37 | 2.5e-02 | 2.1e-02 | 2.8e-02 | 0.0e+00 | 7.6e-02 |
| *Branched-chain amino acids* | | | | | | | | |
| Isoleucine (mmol/l) | thaw-buffer | 50 | 25 | -1.6e-03 | -2.8e-03 | -3.4e-04 | 1.2e-02 | 2.5e-02 |
| Isoleucine (mmol/l) | buffer-NMR | 50 | 25 | -8.3e-04 | -2.4e-03 | 6.9e-04 | 2.9e-01 | 2.5e-02 |
| Leucine (mmol/l) | thaw-buffer | 50 | 25 | 1.5e-04 | -7.6e-04 | 1.1e-03 | 7.5e-01 | 2.1e-02 |
| Leucine (mmol/l) | buffer-NMR | 50 | 25 | 3.4e-03 | 2.5e-03 | 4.4e-03 | 1.4e-12 | 2.1e-02 |
| Valine (mmol/l) | thaw-buffer | 50 | 25 | 1.0e-03 | -2.0e-04 | 2.2e-03 | 1.0e-01 | 3.5e-02 |
| Valine (mmol/l) | buffer-NMR | 50 | 25 | 4.8e-03 | 3.3e-03 | 6.2e-03 | 2.1e-10 | 3.5e-02 |
| *Aromatic amino acids* | | | | | | | | |
| Phenylalanine (mmol/l) | thaw-buffer | 50 | 25 | 4.3e-03 | 2.3e-03 | 6.3e-03 | 2.1e-05 | 8.4e-03 |
| Phenylalanine (mmol/l) | buffer-NMR | 50 | 25 | 9.1e-03 | 7.9e-03 | 1.0e-02 | 0.0e+00 | 8.4e-03 |
| Tyrosine (mmol/l) | thaw-buffer | 50 | 25 | 1.4e-03 | 2.7e-04 | 2.5e-03 | 1.4e-02 | 1.2e-02 |

| Metabolic traits | delay | N.obs | N.indiv | Beta | LCI | UCI | Pvalue | SD |
|---|---|---|---|---|---|---|---|---|
| Tyrosine (mmol/l) | buffer-NMR | 50 | 25 | 4.3e-04 | -6.7e-04 | 1.5e-03 | 4.4e-01 | 1.2e-02 |
| **Ketone bodies** | | | | | | | | |
| Acetate (mmol/l) | thaw-buffer | 50 | 25 | 4.9e-03 | 4.0e-03 | 5.7e-03 | 0.0e+00 | 1.1e-02 |
| Acetate (mmol/l) | buffer-NMR | 50 | 25 | 2.4e-03 | 1.4e-03 | 3.3e-03 | 1.1e-06 | 1.1e-02 |
| Beta-hydroxybutyrate (mmol/l) | thaw-buffer | 50 | 25 | 3.1e-03 | 1.8e-03 | 4.5e-03 | 5.7e-06 | 1.6e-02 |
| Beta-hydroxybutyrate (mmol/l) | buffer-NMR | 50 | 25 | 1.5e-03 | 1.9e-04 | 2.8e-03 | 2.4e-02 | 1.6e-02 |
| **Fluid balance** | | | | | | | | |
| Creatinine (mmol/l) | thaw-buffer | 50 | 25 | 3.1e-03 | 1.8e-03 | 4.3e-03 | 1.3e-06 | 9.4e-03 |
| Creatinine (mmol/l) | buffer-NMR | 50 | 25 | 1.0e-03 | -3.5e-04 | 2.4e-03 | 1.4e-01 | 9.4e-03 |
| Albumin (signal area) | thaw-buffer | 50 | 25 | -1.7e-05 | -1.3e-03 | 1.3e-03 | 9.8e-01 | 4.6e-03 |
| Albumin (signal area) | buffer-NMR | 50 | 25 | -9.5e-04 | -1.5e-03 | -4.4e-04 | 2.9e-04 | 4.6e-03 |
| **Inflammation** | | | | | | | | |
| Glycoprotein acetyls (mmol/l) | thaw-buffer | 50 | 25 | -9.2e-03 | -1.8e-02 | -1.7e-04 | 4.6e-02 | 3.3e-01 |
| Glycoprotein acetyls (mmol/l) | buffer-NMR | 50 | 25 | -1.2e-02 | -2.3e-02 | -1.4e-03 | 2.8e-02 | 3.3e-01 |

*sTable 5. EDTA-plasma, post-storage handling effects*: *mean differences in metabolite concentrations (or trait value) comparing delays in sample preparation (i.e. thaw to buffer addition delay) and NMR profiling (buffer addition to NMR profiling delay) to the reference, for EDTA -plasma samples. Pyruvate, glycerol and glycine are not quantified in EDTA -plasma samples due to the interfering resonances of EDTA on their signals.*

*# Associations in sFigure3 and Figures 3 and 4 are presented in SD-units.* These SD point estimate can be obtained by dividing the point estimate (beta) in absolute (clinically meaningful) concentration by the metabolic trait standard deviation (SD), both provided in the below table.

***Abbreviations**: **C**=cholesterol; **IDL**=intermediate-density lipoprotein; **LCI**=lower confidence interval; **LDL**=low-density lipoprotein; **HDL**=high-density lipoprotein; **MUFA**=monounsaturated fatty acids; **N.obs**= number of observations (samples); **N.indiv**=number of individuals; **PUFA**=polyunsaturated fatty acids; **SD**=standard deviation; **UCI**= upper confidence interval; **VLDL**=very-low-density lipoprotein.*

| Metabolic traits | delay | N.obs | N.indiv | Beta | LCI | UCI | Pvalue | SD |
|---|---|---|---|---|---|---|---|---|
| **Lipoprotein subclasses** | | | | | | | | |
| *Extremely large VLDL* | | | | | | | | |
| Particle concentration (mol/l) | thaw-buffer | 50 | 25 | 1.0e-11 | -1.6e-12 | 2.2e-11 | 9.2e-02 | 2.1e-10 |
| Particle concentration (mol/l) | buffer-NMR | 50 | 25 | 9.2e-12 | -1.5e-12 | 2.0e-11 | 9.1e-02 | 2.1e-10 |
| Total lipids (mmol/l) | thaw-buffer | 50 | 25 | 2.1e-03 | -3.6e-04 | 4.6e-03 | 9.4e-02 | 4.6e-02 |
| Total lipids (mmol/l) | buffer-NMR | 50 | 25 | 1.9e-03 | -3.5e-04 | 4.2e-03 | 9.7e-02 | 4.6e-02 |
| Phospholipids (mmol/l) | thaw-buffer | 50 | 25 | 2.6e-04 | -3.9e-05 | 5.6e-04 | 8.9e-02 | 5.7e-03 |
| Phospholipids (mmol/l) | buffer-NMR | 50 | 25 | 2.0e-04 | -6.9e-05 | 4.8e-04 | 1.4e-01 | 5.7e-03 |
| Total cholesterol (mmol/l) | thaw-buffer | 50 | 25 | 3.6e-04 | -7.1e-05 | 7.8e-04 | 1.0e-01 | 8.5e-03 |
| Total cholesterol (mmol/l) | buffer-NMR | 50 | 25 | 2.4e-04 | -1.3e-04 | 6.1e-04 | 2.0e-01 | 8.5e-03 |
| Cholesterol esters (mmol/l) | thaw-buffer | 50 | 25 | 2.6e-04 | -8.8e-06 | 5.2e-04 | 5.8e-02 | 4.7e-03 |
| Cholesterol esters (mmol/l) | buffer-NMR | 50 | 25 | 1.4e-04 | -7.8e-05 | 3.6e-04 | 2.1e-01 | 4.7e-03 |
| Free cholesterol (mmol/l) | thaw-buffer | 50 | 25 | 1.0e-04 | -6.4e-05 | 2.7e-04 | 2.3e-01 | 3.7e-03 |
| Free cholesterol (mmol/l) | buffer-NMR | 50 | 25 | 9.9e-05 | -5.9e-05 | 2.6e-04 | 2.2e-01 | 3.7e-03 |
| Triglycerides (mmol/l) | thaw-buffer | 50 | 25 | 1.5e-03 | -2.6e-04 | 3.3e-03 | 9.4e-02 | 3.2e-02 |
| Triglycerides (mmol/l) | buffer-NMR | 50 | 25 | 1.5e-03 | -1.6e-04 | 3.1e-03 | 7.7e-02 | 3.2e-02 |
| *Very large VLDL* | | | | | | | | |
| Particle concentration (mol/l) | thaw-buffer | 50 | 25 | 1.1e-11 | -2.2e-11 | 4.4e-11 | 5.2e-01 | 1.3e-09 |
| Particle concentration (mol/l) | buffer-NMR | 50 | 25 | 2.9e-11 | -1.4e-11 | 7.2e-11 | 1.8e-01 | 1.3e-09 |
| Total lipids (mmol/l) | thaw-buffer | 50 | 25 | 1.1e-03 | -2.2e-03 | 4.5e-03 | 5.1e-01 | 1.2e-01 |

| Metabolic traits | delay | N.obs | N.indiv | Beta | LCI | UCI | Pvalue | SD |
|---|---|---|---|---|---|---|---|---|
| Total lipids (mmol/l) | buffer-NMR | 50 | 25 | 2.8e-03 | -1.3e-03 | 6.9e-03 | 1.8e-01 | 1.2e-01 |
| Phospholipids (mmol/l) | thaw-buffer | 50 | 25 | 1.1e-04 | -4.9e-04 | 7.0e-04 | 7.3e-01 | 2.0e-02 |
| Phospholipids (mmol/l) | buffer-NMR | 50 | 25 | 4.2e-04 | -2.0e-04 | 1.0e-03 | 1.9e-01 | 2.0e-02 |
| Total cholesterol (mmol/l) | thaw-buffer | 50 | 25 | 5.2e-04 | -5.0e-04 | 1.5e-03 | 3.2e-01 | 2.5e-02 |
| Total cholesterol (mmol/l) | buffer-NMR | 50 | 25 | 5.7e-04 | -3.6e-04 | 1.5e-03 | 2.3e-01 | 2.5e-02 |
| Cholesterol esters (mmol/l) | thaw-buffer | 50 | 25 | 4.2e-04 | -1.5e-04 | 9.9e-04 | 1.5e-01 | 1.4e-02 |
| Cholesterol esters (mmol/l) | buffer-NMR | 50 | 25 | 3.2e-04 | -1.9e-04 | 8.3e-04 | 2.2e-01 | 1.4e-02 |
| Free cholesterol (mmol/l) | thaw-buffer | 50 | 25 | 9.6e-05 | -3.5e-04 | 5.5e-04 | 6.7e-01 | 1.1e-02 |
| Free cholesterol (mmol/l) | buffer-NMR | 50 | 25 | 2.6e-04 | -1.7e-04 | 6.8e-04 | 2.4e-01 | 1.1e-02 |
| Triglycerides (mmol/l) | thaw-buffer | 50 | 25 | 5.0e-04 | -1.3e-03 | 2.3e-03 | 5.8e-01 | 7.8e-02 |
| Triglycerides (mmol/l) | buffer-NMR | 50 | 25 | 1.8e-03 | -9.2e-04 | 4.6e-03 | 1.9e-01 | 7.8e-02 |
| *Large VLDL* | | | | | | | | |
| Particle concentration (mol/l) | thaw-buffer | 50 | 25 | 2.3e-10 | 6.4e-11 | 3.9e-10 | 6.6e-03 | 7.1e-09 |
| Particle concentration (mol/l) | buffer-NMR | 50 | 25 | 1.2e-10 | -5.5e-11 | 3.0e-10 | 1.8e-01 | 7.1e-09 |
| Total lipids (mmol/l) | thaw-buffer | 50 | 25 | 1.3e-02 | 3.3e-03 | 2.2e-02 | 8.3e-03 | 4.1e-01 |
| Total lipids (mmol/l) | buffer-NMR | 50 | 25 | 7.2e-03 | -3.0e-03 | 1.7e-02 | 1.7e-01 | 4.1e-01 |
| Phospholipids (mmol/l) | thaw-buffer | 50 | 25 | 2.3e-03 | 5.7e-04 | 4.0e-03 | 8.9e-03 | 7.4e-02 |
| Phospholipids (mmol/l) | buffer-NMR | 50 | 25 | 1.3e-03 | -5.3e-04 | 3.1e-03 | 1.7e-01 | 7.4e-02 |
| Total cholesterol (mmol/l) | thaw-buffer | 50 | 25 | 1.8e-03 | -7.3e-04 | 4.3e-03 | 1.6e-01 | 9.6e-02 |
| Total cholesterol (mmol/l) | buffer-NMR | 50 | 25 | 2.2e-03 | -3.6e-04 | 4.7e-03 | 9.2e-02 | 9.6e-02 |
| Cholesterol esters (mmol/l) | thaw-buffer | 50 | 25 | 7.5e-04 | -6.5e-04 | 2.2e-03 | 2.9e-01 | 4.8e-02 |
| Cholesterol esters (mmol/l) | buffer-NMR | 50 | 25 | 1.4e-03 | 1.7e-05 | 2.7e-03 | 4.7e-02 | 4.8e-02 |
| Free cholesterol (mmol/l) | thaw-buffer | 50 | 25 | 1.0e-03 | -1.2e-04 | 2.2e-03 | 8.0e-02 | 4.8e-02 |
| Free cholesterol (mmol/l) | buffer-NMR | 50 | 25 | 8.2e-04 | -4.1e-04 | 2.1e-03 | 1.9e-01 | 4.8e-02 |
| Triglycerides (mmol/l) | thaw-buffer | 50 | 25 | 8.7e-03 | 2.9e-03 | 1.4e-02 | 3.2e-03 | 2.4e-01 |
| Triglycerides (mmol/l) | buffer-NMR | 50 | 25 | 3.7e-03 | -2.4e-03 | 9.7e-03 | 2.3e-01 | 2.4e-01 |
| *Medium VLDL* | | | | | | | | |

| Metabolic traits | delay | N.obs | N.indiv | Beta | LCI | UCI | Pvalue | SD |
|---|---|---|---|---|---|---|---|---|
| Particle concentration (mol/l) | thaw-buffer | 50 | 25 | 5.6e-10 | 1.7e-10 | 9.5e-10 | 4.6e-03 | 1.8e-08 |
| Particle concentration (mol/l) | buffer-NMR | 50 | 25 | 3.0e-10 | -8.4e-11 | 6.9e-10 | 1.3e-01 | 1.8e-08 |
| Total lipids (mmol/l) | thaw-buffer | 50 | 25 | 1.7e-02 | 4.4e-03 | 3.1e-02 | 8.6e-03 | 5.8e-01 |
| Total lipids (mmol/l) | buffer-NMR | 50 | 25 | 1.1e-02 | -2.5e-03 | 2.4e-02 | 1.1e-01 | 5.8e-01 |
| Phospholipids (mmol/l) | thaw-buffer | 50 | 25 | 3.4e-03 | 8.7e-04 | 5.9e-03 | 8.2e-03 | 1.1e-01 |
| Phospholipids (mmol/l) | buffer-NMR | 50 | 25 | 2.1e-03 | -3.6e-04 | 4.6e-03 | 9.4e-02 | 1.1e-01 |
| Total cholesterol (mmol/l) | thaw-buffer | 50 | 25 | 2.6e-04 | -3.6e-03 | 4.1e-03 | 8.9e-01 | 1.5e-01 |
| Total cholesterol (mmol/l) | buffer-NMR | 50 | 25 | 3.9e-03 | 3.3e-05 | 7.7e-03 | 4.8e-02 | 1.5e-01 |
| Cholesterol esters (mmol/l) | thaw-buffer | 50 | 25 | -2.2e-03 | -4.6e-03 | 1.3e-04 | 6.4e-02 | 7.5e-02 |
| Cholesterol esters (mmol/l) | buffer-NMR | 50 | 25 | 2.4e-03 | 1.7e-04 | 4.5e-03 | 3.4e-02 | 7.5e-02 |
| Free cholesterol (mmol/l) | thaw-buffer | 50 | 25 | 2.5e-03 | 8.5e-04 | 4.2e-03 | 3.0e-03 | 7.2e-02 |
| Free cholesterol (mmol/l) | buffer-NMR | 50 | 25 | 1.5e-03 | -2.7e-04 | 3.3e-03 | 9.6e-02 | 7.2e-02 |
| Triglycerides (mmol/l) | thaw-buffer | 50 | 25 | 1.4e-02 | 6.9e-03 | 2.1e-02 | 1.0e-04 | 3.2e-01 |
| Triglycerides (mmol/l) | buffer-NMR | 50 | 25 | 4.6e-03 | -2.4e-03 | 1.2e-02 | 2.0e-01 | 3.2e-01 |
| *Small VLDL* | | | | | | | | |
| Particle concentration (mol/l) | thaw-buffer | 50 | 25 | -1.1e-09 | -1.6e-09 | -6.4e-10 | 3.2e-06 | 1.7e-08 |
| Particle concentration (mol/l) | buffer-NMR | 50 | 25 | -1.5e-10 | -5.0e-10 | 2.1e-10 | 4.2e-01 | 1.7e-08 |
| Total lipids (mmol/l) | thaw-buffer | 50 | 25 | -2.5e-02 | -3.4e-02 | -1.6e-02 | 9.0e-08 | 3.1e-01 |
| Total lipids (mmol/l) | buffer-NMR | 50 | 25 | -3.2e-03 | -1.0e-02 | 4.0e-03 | 3.8e-01 | 3.1e-01 |
| Phospholipids (mmol/l) | thaw-buffer | 50 | 25 | -8.3e-03 | -1.1e-02 | -5.9e-03 | 1.3e-11 | 6.2e-02 |
| Phospholipids (mmol/l) | buffer-NMR | 50 | 25 | -2.3e-03 | -4.5e-03 | -1.7e-04 | 3.4e-02 | 6.2e-02 |
| Total cholesterol (mmol/l) | thaw-buffer | 50 | 25 | -1.7e-02 | -2.2e-02 | -1.3e-02 | 0.0e+00 | 8.7e-02 |
| Total cholesterol (mmol/l) | buffer-NMR | 50 | 25 | -7.8e-04 | -4.6e-03 | 3.1e-03 | 6.9e-01 | 8.7e-02 |
| Cholesterol esters (mmol/l) | thaw-buffer | 50 | 25 | -1.3e-02 | -1.6e-02 | -1.1e-02 | 0.0e+00 | 4.9e-02 |
| Cholesterol esters (mmol/l) | buffer-NMR | 50 | 25 | -1.8e-04 | -3.2e-03 | 2.8e-03 | 9.1e-01 | 4.9e-02 |
| Free cholesterol (mmol/l) | thaw-buffer | 50 | 25 | -3.9e-03 | -5.3e-03 | -2.5e-03 | 3.8e-08 | 4.0e-02 |
| Free cholesterol (mmol/l) | buffer-NMR | 50 | 25 | -5.9e-04 | -1.6e-03 | 4.3e-04 | 2.6e-01 | 4.0e-02 |

| Metabolic traits | delay | N.obs | N.indiv | Beta | LCI | UCI | Pvalue | SD |
|---|---|---|---|---|---|---|---|---|
| Triglycerides (mmol/l) | thaw-buffer | 50 | 25 | 6.3e-04 | -3.3e-03 | 4.5e-03 | 7.5e-01 | 1.7e-01 |
| Triglycerides (mmol/l) | buffer-NMR | 50 | 25 | -7.8e-05 | -3.0e-03 | 2.9e-03 | 9.6e-01 | 1.7e-01 |
| *Very Small VLDL* | | | | | | | | |
| Particle concentration (mol/l) | thaw-buffer | 50 | 25 | -2.0e-09 | -2.6e-09 | -1.5e-09 | 1.2e-13 | 9.2e-09 |
| Particle concentration (mol/l) | buffer-NMR | 50 | 25 | 3.1e-10 | -2.1e-10 | 8.3e-10 | 2.4e-01 | 9.2e-09 |
| Total lipids (mmol/l) | thaw-buffer | 50 | 25 | -2.7e-02 | -3.4e-02 | -2.0e-02 | 5.8e-14 | 1.1e-01 |
| Total lipids (mmol/l) | buffer-NMR | 50 | 25 | 5.1e-03 | -1.8e-03 | 1.2e-02 | 1.5e-01 | 1.1e-01 |
| Phospholipids (mmol/l) | thaw-buffer | 50 | 25 | -1.6e-03 | -3.9e-03 | 6.9e-04 | 1.7e-01 | 3.0e-02 |
| Phospholipids (mmol/l) | buffer-NMR | 50 | 25 | -2.7e-04 | -2.3e-03 | 1.8e-03 | 7.9e-01 | 3.0e-02 |
| Total cholesterol (mmol/l) | thaw-buffer | 50 | 25 | -2.3e-02 | -2.7e-02 | -1.9e-02 | 0.0e+00 | 4.6e-02 |
| Total cholesterol (mmol/l) | buffer-NMR | 50 | 25 | 6.6e-03 | 1.6e-03 | 1.2e-02 | 9.2e-03 | 4.6e-02 |
| Cholesterol esters (mmol/l) | thaw-buffer | 50 | 25 | -2.1e-02 | -2.5e-02 | -1.8e-02 | 0.0e+00 | 3.3e-02 |
| Cholesterol esters (mmol/l) | buffer-NMR | 50 | 25 | 5.7e-03 | 1.7e-03 | 9.6e-03 | 4.9e-03 | 3.3e-02 |
| Free cholesterol (mmol/l) | thaw-buffer | 50 | 25 | -1.7e-03 | -3.0e-03 | -4.8e-04 | 6.5e-03 | 1.4e-02 |
| Free cholesterol (mmol/l) | buffer-NMR | 50 | 25 | 9.4e-04 | -1.6e-04 | 2.0e-03 | 9.5e-02 | 1.4e-02 |
| Triglycerides (mmol/l) | thaw-buffer | 50 | 25 | -2.4e-03 | -4.2e-03 | -5.1e-04 | 1.2e-02 | 4.6e-02 |
| Triglycerides (mmol/l) | buffer-NMR | 50 | 25 | -1.3e-03 | -2.6e-03 | 1.3e-04 | 7.6e-02 | 4.6e-02 |
| *IDL* | | | | | | | | |
| Particle concentration (mol/l) | thaw-buffer | 50 | 25 | -4.5e-10 | -1.9e-09 | 1.0e-09 | 5.6e-01 | 2.0e-08 |
| Particle concentration (mol/l) | buffer-NMR | 50 | 25 | 2.0e-09 | 3.0e-10 | 3.7e-09 | 2.1e-02 | 2.0e-08 |
| Total lipids (mmol/l) | thaw-buffer | 50 | 25 | -3.2e-03 | -1.9e-02 | 1.3e-02 | 7.0e-01 | 2.0e-01 |
| Total lipids (mmol/l) | buffer-NMR | 50 | 25 | 2.2e-02 | 4.0e-03 | 4.1e-02 | 1.7e-02 | 2.0e-01 |
| Phospholipids (mmol/l) | thaw-buffer | 50 | 25 | -3.7e-04 | -3.8e-03 | 3.1e-03 | 8.3e-01 | 4.9e-02 |
| Phospholipids (mmol/l) | buffer-NMR | 50 | 25 | 3.2e-03 | -4.8e-04 | 6.9e-03 | 8.8e-02 | 4.9e-02 |
| Total cholesterol (mmol/l) | thaw-buffer | 50 | 25 | -1.7e-03 | -1.5e-02 | 1.1e-02 | 7.9e-01 | 1.3e-01 |
| Total cholesterol (mmol/l) | buffer-NMR | 50 | 25 | 2.1e-02 | 5.6e-03 | 3.6e-02 | 7.4e-03 | 1.3e-01 |
| Cholesterol esters (mmol/l) | thaw-buffer | 50 | 25 | -3.6e-03 | -1.4e-02 | 6.4e-03 | 4.8e-01 | 1.0e-01 |

| Metabolic traits | delay | N.obs | N.indiv | Beta | LCI | UCI | Pvalue | SD |
|---|---|---|---|---|---|---|---|---|
| Cholesterol esters (mmol/l) | buffer-NMR | 50 | 25 | 1.7e-02 | 4.9e-03 | 2.9e-02 | 5.6e-03 | 1.0e-01 |
| Free cholesterol (mmol/l) | thaw-buffer | 50 | 25 | 1.9e-03 | -1.3e-03 | 5.0e-03 | 2.4e-01 | 3.9e-02 |
| Free cholesterol (mmol/l) | buffer-NMR | 50 | 25 | 4.2e-03 | 6.3e-04 | 7.7e-03 | 2.1e-02 | 3.9e-02 |
| Triglycerides (mmol/l) | thaw-buffer | 50 | 25 | -1.1e-03 | -3.0e-03 | 8.1e-04 | 2.6e-01 | 3.1e-02 |
| Triglycerides (mmol/l) | buffer-NMR | 50 | 25 | -1.8e-03 | -3.3e-03 | -2.7e-04 | 2.1e-02 | 3.1e-02 |
| *Large LDL* | | | | | | | | |
| Particle concentration (mol/l) | thaw-buffer | 50 | 25 | 1.8e-09 | -5.5e-10 | 4.2e-09 | 1.3e-01 | 3.4e-08 |
| Particle concentration (mol/l) | buffer-NMR | 50 | 25 | 1.9e-09 | -5.4e-10 | 4.3e-09 | 1.3e-01 | 3.4e-08 |
| Total lipids (mmol/l) | thaw-buffer | 50 | 25 | 1.4e-02 | -2.9e-03 | 3.2e-02 | 1.0e-01 | 2.4e-01 |
| Total lipids (mmol/l) | buffer-NMR | 50 | 25 | 1.5e-02 | -2.6e-03 | 3.3e-02 | 9.5e-02 | 2.4e-01 |
| Phospholipids (mmol/l) | thaw-buffer | 50 | 25 | 9.2e-04 | -2.5e-03 | 4.4e-03 | 6.0e-01 | 5.1e-02 |
| Phospholipids (mmol/l) | buffer-NMR | 50 | 25 | 2.7e-03 | -9.4e-04 | 6.4e-03 | 1.5e-01 | 5.1e-02 |
| Total cholesterol (mmol/l) | thaw-buffer | 50 | 25 | 1.6e-02 | 2.2e-03 | 3.0e-02 | 2.3e-02 | 1.8e-01 |
| Total cholesterol (mmol/l) | buffer-NMR | 50 | 25 | 1.5e-02 | 2.0e-04 | 2.9e-02 | 4.7e-02 | 1.8e-01 |
| Cholesterol esters (mmol/l) | thaw-buffer | 50 | 25 | 1.4e-02 | 3.5e-03 | 2.4e-02 | 8.5e-03 | 1.4e-01 |
| Cholesterol esters (mmol/l) | buffer-NMR | 50 | 25 | 1.1e-02 | 4.0e-05 | 2.2e-02 | 4.9e-02 | 1.4e-01 |
| Free cholesterol (mmol/l) | thaw-buffer | 50 | 25 | 2.1e-03 | -1.3e-03 | 5.5e-03 | 2.2e-01 | 4.5e-02 |
| Free cholesterol (mmol/l) | buffer-NMR | 50 | 25 | 3.8e-03 | 2.2e-04 | 7.3e-03 | 3.7e-02 | 4.5e-02 |
| Triglycerides (mmol/l) | thaw-buffer | 50 | 25 | -2.5e-03 | -4.5e-03 | -5.3e-04 | 1.3e-02 | 2.5e-02 |
| Triglycerides (mmol/l) | buffer-NMR | 50 | 25 | -2.4e-03 | -4.1e-03 | -7.6e-04 | 4.2e-03 | 2.5e-02 |
| *Medium LDL* | | | | | | | | |
| Particle concentration (mol/l) | thaw-buffer | 50 | 25 | 4.9e-09 | 2.9e-09 | 6.9e-09 | 1.4e-06 | 2.9e-08 |
| Particle concentration (mol/l) | buffer-NMR | 50 | 25 | 6.7e-10 | -1.3e-09 | 2.6e-09 | 5.0e-01 | 2.9e-08 |
| Total lipids (mmol/l) | thaw-buffer | 50 | 25 | 2.5e-02 | 1.5e-02 | 3.6e-02 | 1.1e-06 | 1.5e-01 |
| Total lipids (mmol/l) | buffer-NMR | 50 | 25 | 4.6e-03 | -5.6e-03 | 1.5e-02 | 3.7e-01 | 1.5e-01 |
| Phospholipids (mmol/l) | thaw-buffer | 50 | 25 | 2.2e-03 | 2.7e-04 | 4.1e-03 | 2.6e-02 | 3.5e-02 |
| Phospholipids (mmol/l) | buffer-NMR | 50 | 25 | 1.2e-03 | -8.8e-04 | 3.3e-03 | 2.5e-01 | 3.5e-02 |

| Metabolic traits | delay | N.obs | N.indiv | Beta | LCI | UCI | Pvalue | SD |
|---|---|---|---|---|---|---|---|---|
| Total cholesterol (mmol/l) | thaw-buffer | 50 | 25 | 2.4e-02 | 1.6e-02 | 3.2e-02 | 6.3e-09 | 1.1e-01 |
| Total cholesterol (mmol/l) | buffer-NMR | 50 | 25 | 5.7e-03 | -2.5e-03 | 1.4e-02 | 1.7e-01 | 1.1e-01 |
| Cholesterol esters (mmol/l) | thaw-buffer | 50 | 25 | 2.2e-02 | 1.5e-02 | 2.9e-02 | 9.7e-11 | 9.0e-02 |
| Cholesterol esters (mmol/l) | buffer-NMR | 50 | 25 | 4.5e-03 | -2.1e-03 | 1.1e-02 | 1.8e-01 | 9.0e-02 |
| Free cholesterol (mmol/l) | thaw-buffer | 50 | 25 | 2.3e-03 | 7.3e-04 | 3.9e-03 | 4.0e-03 | 2.1e-02 |
| Free cholesterol (mmol/l) | buffer-NMR | 50 | 25 | 1.2e-03 | -4.6e-04 | 2.9e-03 | 1.5e-01 | 2.1e-02 |
| Triglycerides (mmol/l) | thaw-buffer | 50 | 25 | -9.7e-04 | -2.4e-03 | 4.5e-04 | 1.8e-01 | 1.3e-02 |
| Triglycerides (mmol/l) | buffer-NMR | 50 | 25 | -2.3e-03 | -3.7e-03 | -9.9e-04 | 6.7e-04 | 1.3e-02 |
| *Small LDL* | | | | | | | | |
| Particle concentration (mol/l) | thaw-buffer | 50 | 25 | 6.3e-09 | 4.1e-09 | 8.5e-09 | 1.7e-08 | 3.4e-08 |
| Particle concentration (mol/l) | buffer-NMR | 50 | 25 | 2.7e-10 | -2.0e-09 | 2.6e-09 | 8.2e-01 | 3.4e-08 |
| Total lipids (mmol/l) | thaw-buffer | 50 | 25 | 1.8e-02 | 1.2e-02 | 2.4e-02 | 1.7e-08 | 9.5e-02 |
| Total lipids (mmol/l) | buffer-NMR | 50 | 25 | 1.4e-03 | -5.1e-03 | 8.0e-03 | 6.7e-01 | 9.5e-02 |
| Phospholipids (mmol/l) | thaw-buffer | 50 | 25 | 2.6e-03 | 1.3e-03 | 3.9e-03 | 9.3e-05 | 2.4e-02 |
| Phospholipids (mmol/l) | buffer-NMR | 50 | 25 | 2.3e-05 | -1.6e-03 | 1.6e-03 | 9.8e-01 | 2.4e-02 |
| Total cholesterol (mmol/l) | thaw-buffer | 50 | 25 | 1.5e-02 | 1.1e-02 | 2.0e-02 | 3.1e-10 | 6.9e-02 |
| Total cholesterol (mmol/l) | buffer-NMR | 50 | 25 | 2.5e-03 | -2.5e-03 | 7.5e-03 | 3.2e-01 | 6.9e-02 |
| Cholesterol esters (mmol/l) | thaw-buffer | 50 | 25 | 1.4e-02 | 9.8e-03 | 1.8e-02 | 4.3e-12 | 5.6e-02 |
| Cholesterol esters (mmol/l) | buffer-NMR | 50 | 25 | 1.9e-03 | -2.0e-03 | 5.8e-03 | 3.3e-01 | 5.6e-02 |
| Free cholesterol (mmol/l) | thaw-buffer | 50 | 25 | 1.7e-03 | 7.8e-04 | 2.7e-03 | 4.1e-04 | 1.4e-02 |
| Free cholesterol (mmol/l) | buffer-NMR | 50 | 25 | 6.1e-04 | -5.1e-04 | 1.7e-03 | 2.9e-01 | 1.4e-02 |
| Triglycerides (mmol/l) | thaw-buffer | 50 | 25 | -1.0e-04 | -7.3e-04 | 5.3e-04 | 7.5e-01 | 1.2e-02 |
| Triglycerides (mmol/l) | buffer-NMR | 50 | 25 | -1.1e-03 | -1.7e-03 | -5.4e-04 | 1.5e-04 | 1.2e-02 |
| *Very large HDL* | | | | | | | | |
| Particle concentration (mol/l) | thaw-buffer | 50 | 25 | -4.7e-08 | -5.6e-08 | -3.7e-08 | 0.0e+00 | 2.4e-07 |
| Particle concentration (mol/l) | buffer-NMR | 50 | 25 | -4.3e-09 | -1.6e-08 | 7.3e-09 | 4.7e-01 | 2.4e-07 |
| Total lipids (mmol/l) | thaw-buffer | 50 | 25 | -5.1e-02 | -6.0e-02 | -4.1e-02 | 0.0e+00 | 2.4e-01 |

| Metabolic traits | delay | N.obs | N.indiv | Beta | LCI | UCI | Pvalue | SD |
|---|---|---|---|---|---|---|---|---|
| Total lipids (mmol/l) | buffer-NMR | 50 | 25 | -3.7e-03 | -1.6e-02 | 8.2e-03 | 5.4e-01 | 2.4e-01 |
| Phospholipids (mmol/l) | thaw-buffer | 50 | 25 | -2.1e-02 | -2.6e-02 | -1.6e-02 | 6.7e-16 | 1.4e-01 |
| Phospholipids (mmol/l) | buffer-NMR | 50 | 25 | -8.1e-03 | -1.3e-02 | -3.1e-03 | 1.5e-03 | 1.4e-01 |
| Total cholesterol (mmol/l) | thaw-buffer | 50 | 25 | -3.2e-02 | -3.7e-02 | -2.6e-02 | 0.0e+00 | 1.1e-01 |
| Total cholesterol (mmol/l) | buffer-NMR | 50 | 25 | 3.2e-03 | -4.3e-03 | 1.1e-02 | 4.0e-01 | 1.1e-01 |
| Cholesterol esters (mmol/l) | thaw-buffer | 50 | 25 | -2.2e-02 | -2.6e-02 | -1.8e-02 | 0.0e+00 | 7.4e-02 |
| Cholesterol esters (mmol/l) | buffer-NMR | 50 | 25 | 2.8e-03 | -2.8e-03 | 8.4e-03 | 3.3e-01 | 7.4e-02 |
| Free cholesterol (mmol/l) | thaw-buffer | 50 | 25 | -9.5e-03 | -1.1e-02 | -8.0e-03 | 0.0e+00 | 3.2e-02 |
| Free cholesterol (mmol/l) | buffer-NMR | 50 | 25 | 4.0e-04 | -1.6e-03 | 2.4e-03 | 6.9e-01 | 3.2e-02 |
| Triglycerides (mmol/l) | thaw-buffer | 50 | 25 | 1.6e-03 | 5.0e-04 | 2.7e-03 | 4.2e-03 | 1.0e-02 |
| Triglycerides (mmol/l) | buffer-NMR | 50 | 25 | 1.2e-03 | -6.4e-05 | 2.5e-03 | 6.2e-02 | 1.0e-02 |
| *Large HDL* | | | | | | | | |
| Particle concentration (mol/l) | thaw-buffer | 50 | 25 | 3.1e-08 | -6.4e-09 | 6.8e-08 | 1.1e-01 | 6.5e-07 |
| Particle concentration (mol/l) | buffer-NMR | 50 | 25 | -3.6e-08 | -7.3e-08 | 1.6e-09 | 6.1e-02 | 6.5e-07 |
| Total lipids (mmol/l) | thaw-buffer | 50 | 25 | 1.9e-02 | -3.6e-03 | 4.1e-02 | 1.0e-01 | 4.2e-01 |
| Total lipids (mmol/l) | buffer-NMR | 50 | 25 | -2.3e-02 | -4.6e-02 | -1.0e-03 | 4.0e-02 | 4.2e-01 |
| Phospholipids (mmol/l) | thaw-buffer | 50 | 25 | 5.4e-04 | -1.1e-02 | 1.3e-02 | 9.3e-01 | 1.9e-01 |
| Phospholipids (mmol/l) | buffer-NMR | 50 | 25 | -1.7e-02 | -2.9e-02 | -4.8e-03 | 6.1e-03 | 1.9e-01 |
| Total cholesterol (mmol/l) | thaw-buffer | 50 | 25 | 1.3e-02 | 3.2e-03 | 2.2e-02 | 8.4e-03 | 2.2e-01 |
| Total cholesterol (mmol/l) | buffer-NMR | 50 | 25 | -8.7e-03 | -1.9e-02 | 1.1e-03 | 8.1e-02 | 2.2e-01 |
| Cholesterol esters (mmol/l) | thaw-buffer | 50 | 25 | 1.1e-02 | 2.8e-03 | 1.9e-02 | 8.3e-03 | 1.7e-01 |
| Cholesterol esters (mmol/l) | buffer-NMR | 50 | 25 | -5.2e-03 | -1.4e-02 | 3.0e-03 | 2.1e-01 | 1.7e-01 |
| Free cholesterol (mmol/l) | thaw-buffer | 50 | 25 | 1.9e-03 | 1.3e-04 | 3.6e-03 | 3.6e-02 | 5.5e-02 |
| Free cholesterol (mmol/l) | buffer-NMR | 50 | 25 | -3.5e-03 | -5.2e-03 | -1.8e-03 | 8.2e-05 | 5.5e-02 |
| Triglycerides (mmol/l) | thaw-buffer | 50 | 25 | 5.3e-03 | 2.1e-03 | 8.6e-03 | 1.3e-03 | 1.4e-02 |
| Triglycerides (mmol/l) | buffer-NMR | 50 | 25 | 2.2e-03 | -1.1e-03 | 5.4e-03 | 1.9e-01 | 1.4e-02 |
| *Medium HDL* | | | | | | | | |

| Metabolic traits | delay | N.obs | N.indiv | Beta | LCI | UCI | Pvalue | SD |
|---|---|---|---|---|---|---|---|---|
| Particle concentration (mol/l) | thaw-buffer | 50 | 25 | -6.0e-08 | -1.1e-07 | -7.2e-09 | 2.6e-02 | 3.7e-07 |
| Particle concentration (mol/l) | buffer-NMR | 50 | 25 | -8.8e-08 | -1.5e-07 | -3.1e-08 | 2.4e-03 | 3.7e-07 |
| Total lipids (mmol/l) | thaw-buffer | 50 | 25 | -2.4e-02 | -4.7e-02 | -1.6e-03 | 3.6e-02 | 1.6e-01 |
| Total lipids (mmol/l) | buffer-NMR | 50 | 25 | -3.8e-02 | -6.3e-02 | -1.3e-02 | 2.5e-03 | 1.6e-01 |
| Phospholipids (mmol/l) | thaw-buffer | 50 | 25 | -1.7e-02 | -2.8e-02 | -7.0e-03 | 9.9e-04 | 7.4e-02 |
| Phospholipids (mmol/l) | buffer-NMR | 50 | 25 | -1.8e-02 | -2.9e-02 | -7.0e-03 | 1.4e-03 | 7.4e-02 |
| Total cholesterol (mmol/l) | thaw-buffer | 50 | 25 | -4.3e-03 | -1.7e-02 | 8.0e-03 | 4.9e-01 | 8.9e-02 |
| Total cholesterol (mmol/l) | buffer-NMR | 50 | 25 | -1.9e-02 | -3.2e-02 | -5.6e-03 | 5.3e-03 | 8.9e-02 |
| Cholesterol esters (mmol/l) | thaw-buffer | 50 | 25 | -1.3e-03 | -1.1e-02 | 8.4e-03 | 7.9e-01 | 6.9e-02 |
| Cholesterol esters (mmol/l) | buffer-NMR | 50 | 25 | -1.4e-02 | -2.5e-02 | -4.0e-03 | 6.6e-03 | 6.9e-02 |
| Free cholesterol (mmol/l) | thaw-buffer | 50 | 25 | -3.0e-03 | -5.7e-03 | -3.0e-04 | 3.0e-02 | 2.0e-02 |
| Free cholesterol (mmol/l) | buffer-NMR | 50 | 25 | -4.5e-03 | -7.4e-03 | -1.6e-03 | 2.5e-03 | 2.0e-02 |
| Triglycerides (mmol/l) | thaw-buffer | 50 | 25 | -2.6e-03 | -3.3e-03 | -2.0e-03 | 4.2e-14 | 1.7e-02 |
| Triglycerides (mmol/l) | buffer-NMR | 50 | 25 | -1.1e-03 | -1.8e-03 | -4.4e-04 | 1.2e-03 | 1.7e-02 |
| *Small HDL* | | | | | | | | |
| Particle concentration (mol/l) | thaw-buffer | 50 | 25 | -6.8e-08 | -1.6e-07 | 2.8e-08 | 1.6e-01 | 5.0e-07 |
| Particle concentration (mol/l) | buffer-NMR | 50 | 25 | -1.4e-07 | -2.5e-07 | -3.8e-08 | 7.7e-03 | 5.0e-07 |
| Total lipids (mmol/l) | thaw-buffer | 50 | 25 | -1.5e-02 | -3.7e-02 | 6.9e-03 | 1.8e-01 | 1.1e-01 |
| Total lipids (mmol/l) | buffer-NMR | 50 | 25 | -3.3e-02 | -5.7e-02 | -8.5e-03 | 8.1e-03 | 1.1e-01 |
| Phospholipids (mmol/l) | thaw-buffer | 50 | 25 | -6.1e-03 | -1.2e-02 | 7.7e-05 | 5.3e-02 | 7.2e-02 |
| Phospholipids (mmol/l) | buffer-NMR | 50 | 25 | -7.7e-03 | -1.4e-02 | -1.7e-03 | 1.1e-02 | 7.2e-02 |
| Total cholesterol (mmol/l) | thaw-buffer | 50 | 25 | -8.1e-03 | -2.5e-02 | 8.4e-03 | 3.4e-01 | 6.7e-02 |
| Total cholesterol (mmol/l) | buffer-NMR | 50 | 25 | -2.4e-02 | -4.3e-02 | -5.9e-03 | 9.8e-03 | 6.7e-02 |
| Cholesterol esters (mmol/l) | thaw-buffer | 50 | 25 | -6.8e-03 | -2.2e-02 | 8.6e-03 | 3.9e-01 | 6.5e-02 |
| Cholesterol esters (mmol/l) | buffer-NMR | 50 | 25 | -2.3e-02 | -4.0e-02 | -5.2e-03 | 1.1e-02 | 6.5e-02 |
| Free cholesterol (mmol/l) | thaw-buffer | 50 | 25 | -1.3e-03 | -2.6e-03 | 9.8e-05 | 6.9e-02 | 1.2e-02 |
| Free cholesterol (mmol/l) | buffer-NMR | 50 | 25 | -1.8e-03 | -3.0e-03 | -5.6e-04 | 4.5e-03 | 1.2e-02 |

| Metabolic traits | delay | N.obs | N.indiv | Beta | LCI | UCI | Pvalue | SD |
|---|---|---|---|---|---|---|---|---|
| Triglycerides (mmol/l) | thaw-buffer | 50 | 25 | -1.0e-03 | -1.8e-03 | -2.1e-04 | 1.4e-02 | 2.0e-02 |
| Triglycerides (mmol/l) | buffer-NMR | 50 | 25 | -6.0e-04 | -1.2e-03 | 5.2e-05 | 7.1e-02 | 2.0e-02 |
| **Lipoprotein particle size** | | | | | | | | |
| VLDL particle size (nm) | thaw-buffer | 50 | 25 | 3.0e-01 | 2.1e-01 | 3.9e-01 | 2.0e-10 | 1.6e+00 |
| VLDL particle size (nm) | buffer-NMR | 50 | 25 | 9.0e-02 | -1.2e-02 | 1.9e-01 | 8.3e-02 | 1.6e+00 |
| LDL particle size (nm) | thaw-buffer | 50 | 25 | -6.7e-02 | -8.7e-02 | -4.6e-02 | 2.0e-10 | 8.8e-02 |
| LDL particle size (nm) | buffer-NMR | 50 | 25 | 2.4e-02 | 5.2e-03 | 4.4e-02 | 1.3e-02 | 8.8e-02 |
| HDL particle size (nm) | thaw-buffer | 50 | 25 | -1.1e-02 | -2.4e-02 | 1.5e-03 | 8.3e-02 | 2.9e-01 |
| HDL particle size (nm) | buffer-NMR | 50 | 25 | 2.1e-03 | -1.4e-02 | 1.8e-02 | 8.0e-01 | 2.9e-01 |
| **Cholesterol** | | | | | | | | |
| Total cholesterol (mmol/l) | thaw-buffer | 50 | 25 | -1.6e-02 | -5.8e-02 | 2.5e-02 | 4.4e-01 | 7.3e-01 |
| Total cholesterol (mmol/l) | buffer-NMR | 50 | 25 | 6.4e-03 | -4.0e-02 | 5.3e-02 | 7.9e-01 | 7.3e-01 |
| VLDL cholesterol (mmol/l) | thaw-buffer | 50 | 25 | -3.7e-02 | -4.9e-02 | -2.6e-02 | 1.6e-10 | 3.9e-01 |
| VLDL cholesterol (mmol/l) | buffer-NMR | 50 | 25 | 1.3e-02 | 1.5e-03 | 2.4e-02 | 2.7e-02 | 3.9e-01 |
| Remnant cholesterol (mmol/l) | thaw-buffer | 50 | 25 | -3.9e-02 | -6.2e-02 | -1.6e-02 | 7.8e-04 | 4.6e-01 |
| Remnant cholesterol (mmol/l) | buffer-NMR | 50 | 25 | 3.4e-02 | 8.1e-03 | 5.9e-02 | 9.8e-03 | 4.6e-01 |
| LDL cholesterol (mmol/l) | thaw-buffer | 50 | 25 | 5.6e-02 | 3.0e-02 | 8.2e-02 | 3.1e-05 | 3.6e-01 |
| LDL cholesterol (mmol/l) | buffer-NMR | 50 | 25 | 2.3e-02 | -4.2e-03 | 5.1e-02 | 9.6e-02 | 3.6e-01 |
| HDL cholesterol (mmol/l) | thaw-buffer | 50 | 25 | -3.3e-02 | -5.8e-02 | -8.0e-03 | 9.7e-03 | 3.9e-01 |
| HDL cholesterol (mmol/l) | buffer-NMR | 50 | 25 | -5.0e-02 | -7.5e-02 | -2.6e-02 | 4.5e-05 | 3.9e-01 |
| HDL2 cholesterol (mmol/l) | thaw-buffer | 50 | 25 | -2.3e-02 | -4.5e-02 | -1.5e-03 | 3.6e-02 | 3.6e-01 |
| HDL2 cholesterol (mmol/l) | buffer-NMR | 50 | 25 | -4.4e-02 | -6.5e-02 | -2.3e-02 | 5.0e-05 | 3.6e-01 |
| HDL3 cholesterol (mmol/l) | thaw-buffer | 50 | 25 | -9.5e-03 | -1.4e-02 | -5.1e-03 | 2.1e-05 | 3.6e-02 |
| HDL3 cholesterol (mmol/l) | buffer-NMR | 50 | 25 | -6.6e-03 | -1.0e-02 | -2.9e-03 | 4.6e-04 | 3.6e-02 |
| Esterified cholesterol (mmol/l) | thaw-buffer | 42 | 21 | -2.4e-02 | -6.7e-02 | 1.9e-02 | 2.7e-01 | 5.2e-01 |
| Esterified cholesterol (mmol/l) | buffer-NMR | 42 | 21 | 1.3e-02 | -2.9e-02 | 5.5e-02 | 5.4e-01 | 5.2e-01 |
| Free cholesterol (mmol/l) | thaw-buffer | 42 | 21 | -7.5e-03 | -2.5e-02 | 1.0e-02 | 4.1e-01 | 2.2e-01 |

| Metabolic traits | delay | N.obs | N.indiv | Beta | LCI | UCI | Pvalue | SD |
|---|---|---|---|---|---|---|---|---|
| Free cholesterol (mmol/l) | buffer-NMR | 42 | 21 | -7.1e-03 | -2.8e-02 | 1.4e-02 | 5.1e-01 | 2.2e-01 |
| **Glycerides and phospholipids** | | | | | | | | |
| Triglycerides (mmol/l) | thaw-buffer | 50 | 25 | 1.9e-02 | 1.3e-03 | 3.6e-02 | 3.6e-02 | 9.8e-01 |
| Triglycerides (mmol/l) | buffer-NMR | 50 | 25 | 1.3e-03 | -1.5e-02 | 1.8e-02 | 8.8e-01 | 9.8e-01 |
| VLDL triglycerides (mmol/l) | thaw-buffer | 50 | 25 | 2.2e-02 | 4.6e-03 | 3.9e-02 | 1.3e-02 | 8.8e-01 |
| VLDL triglycerides (mmol/l) | buffer-NMR | 50 | 25 | 8.8e-03 | -8.3e-03 | 2.6e-02 | 3.1e-01 | 8.8e-01 |
| LDL triglycerides (mmol/l) | thaw-buffer | 50 | 25 | -3.6e-03 | -7.6e-03 | 4.0e-04 | 7.8e-02 | 4.9e-02 |
| LDL triglycerides (mmol/l) | buffer-NMR | 50 | 25 | -5.9e-03 | -9.4e-03 | -2.3e-03 | 1.2e-03 | 4.9e-02 |
| HDL triglycerides (mmol/l) | thaw-buffer | 50 | 25 | 1.7e-03 | -1.2e-05 | 3.4e-03 | 5.2e-02 | 4.8e-02 |
| HDL triglycerides (mmol/l) | buffer-NMR | 50 | 25 | 7.1e-05 | -1.8e-03 | 1.9e-03 | 9.4e-01 | 4.8e-02 |
| Diacylglycerol (mmol/l) | thaw-buffer | 40 | 20 | -9.9e-04 | -7.4e-03 | 5.4e-03 | 7.6e-01 | 2.6e-02 |
| Diacylglycerol (mmol/l) | buffer-NMR | 40 | 20 | 1.8e-03 | -2.5e-03 | 6.2e-03 | 4.1e-01 | 2.6e-02 |
| Phosphoglycerides (mmol/l) | thaw-buffer | 42 | 21 | -5.7e-02 | -1.0e-01 | -1.2e-02 | 1.3e-02 | 3.8e-01 |
| Phosphoglycerides (mmol/l) | buffer-NMR | 42 | 21 | -5.6e-02 | -1.2e-01 | 3.6e-03 | 6.5e-02 | 3.8e-01 |
| Phosphatidylcholine + other cholines (mmol/l) | thaw-buffer | 42 | 21 | -1.8e-02 | -6.1e-02 | 2.6e-02 | 4.3e-01 | 3.6e-01 |
| Phosphatidylcholine + other cholines (mmol/l) | buffer-NMR | 42 | 21 | -4.7e-02 | -1.0e-01 | 9.2e-03 | 1.0e-01 | 3.6e-01 |
| Sphingomyelins (mmol/l) | thaw-buffer | 42 | 21 | -3.2e-02 | -5.4e-02 | -1.0e-02 | 4.0e-03 | 8.1e-02 |
| Sphingomyelins (mmol/l) | buffer-NMR | 42 | 21 | -1.4e-02 | -3.8e-02 | 9.8e-03 | 2.5e-01 | 8.1e-02 |
| Cholines (mmol/l) | thaw-buffer | 42 | 21 | -1.4e-02 | -6.5e-02 | 3.6e-02 | 5.7e-01 | 3.8e-01 |
| Cholines (mmol/l) | buffer-NMR | 42 | 21 | -6.1e-02 | -1.2e-01 | 2.4e-03 | 5.9e-02 | 3.8e-01 |
| **Apolipoproteins** | | | | | | | | |
| Apolipoprotein A-I (g/l) | thaw-buffer | 50 | 25 | 2.8e-03 | -9.9e-03 | 1.6e-02 | 6.7e-01 | 1.9e-01 |
| Apolipoprotein A-I (g/l) | buffer-NMR | 50 | 25 | -2.3e-02 | -3.5e-02 | -1.1e-02 | 1.5e-04 | 1.9e-01 |
| Apolipoprotein B (g/l) | thaw-buffer | 50 | 25 | 1.3e-02 | -2.7e-04 | 2.7e-02 | 5.5e-02 | 2.4e-01 |
| Apolipoprotein B (g/l) | buffer-NMR | 50 | 25 | 1.9e-02 | 4.6e-03 | 3.3e-02 | 9.2e-03 | 2.4e-01 |
| **Fatty acids** | | | | | | | | |

| Metabolic traits | delay | N.obs | N.indiv | Beta | LCI | UCI | Pvalue | SD |
|---|---|---|---|---|---|---|---|---|
| Total fatty acids (mmol/l) | thaw-buffer | 42 | 21 | -1.0e-01 | -3.4e-01 | 1.4e-01 | 4.0e-01 | 3.5e+00 |
| Total fatty acids (mmol/l) | buffer-NMR | 42 | 21 | -2.4e-02 | -2.0e-01 | 1.5e-01 | 7.9e-01 | 3.5e+00 |
| Fatty acid chain length | thaw-buffer | 42 | 21 | 5.2e-02 | -2.0e-03 | 1.1e-01 | 5.9e-02 | 2.8e-01 |
| Fatty acid chain length | buffer-NMR | 42 | 21 | 1.1e-01 | 5.3e-03 | 2.1e-01 | 3.9e-02 | 2.8e-01 |
| Degree of unsaturation | thaw-buffer | 42 | 21 | 5.2e-04 | -7.9e-03 | 8.9e-03 | 9.0e-01 | 8.6e-02 |
| Degree of unsaturation | buffer-NMR | 42 | 21 | 4.3e-04 | -2.1e-02 | 2.2e-02 | 9.7e-01 | 8.6e-02 |
| Docosahexaenoic acid (mmol/l) | thaw-buffer | 42 | 21 | -3.1e-03 | -6.9e-03 | 7.8e-04 | 1.2e-01 | 4.9e-02 |
| Docosahexaenoic acid (mmol/l) | buffer-NMR | 42 | 21 | -5.7e-03 | -9.1e-03 | -2.2e-03 | 1.3e-03 | 4.9e-02 |
| Linoleic acid (mmol/l) | thaw-buffer | 42 | 21 | -2.6e-02 | -7.0e-02 | 1.7e-02 | 2.4e-01 | 6.4e-01 |
| Linoleic acid (mmol/l) | buffer-NMR | 42 | 21 | 2.7e-03 | -4.3e-02 | 4.8e-02 | 9.1e-01 | 6.4e-01 |
| Conjugated linoleic acid (mmol/l) | thaw-buffer | 42 | 21 | -4.2e-03 | -8.2e-03 | -2.3e-04 | 3.8e-02 | 2.4e-02 |
| Conjugated linoleic acid (mmol/l) | buffer-NMR | 42 | 21 | -1.8e-03 | -6.6e-03 | 3.1e-03 | 4.7e-01 | 2.4e-02 |
| n-3 fatty acids (mmol/l) | thaw-buffer | 42 | 21 | -1.5e-02 | -2.9e-02 | -1.5e-03 | 3.0e-02 | 1.5e-01 |
| n-3 fatty acids (mmol/l) | buffer-NMR | 42 | 21 | -4.1e-03 | -1.6e-02 | 7.5e-03 | 4.9e-01 | 1.5e-01 |
| n-6 fatty acids (mmol/l) | thaw-buffer | 42 | 21 | -6.9e-03 | -6.2e-02 | 4.8e-02 | 8.1e-01 | 7.2e-01 |
| n-6 fatty acids (mmol/l) | buffer-NMR | 42 | 21 | -2.0e-02 | -8.1e-02 | 4.1e-02 | 5.3e-01 | 7.2e-01 |
| PUFA (mmol/l) | thaw-buffer | 42 | 21 | -2.2e-02 | -8.2e-02 | 3.8e-02 | 4.7e-01 | 8.5e-01 |
| PUFA (mmol/l) | buffer-NMR | 42 | 21 | -2.4e-02 | -8.8e-02 | 4.0e-02 | 4.7e-01 | 8.5e-01 |
| MUFA (mmol/l) | thaw-buffer | 42 | 21 | 4.2e-02 | -2.7e-02 | 1.1e-01 | 2.3e-01 | 1.4e+00 |
| MUFA (mmol/l) | buffer-NMR | 42 | 21 | 9.5e-02 | -2.8e-02 | 2.2e-01 | 1.3e-01 | 1.4e+00 |
| Saturated fatty acids (mmol/l) | thaw-buffer | 42 | 21 | -1.2e-01 | -2.7e-01 | 2.1e-02 | 9.3e-02 | 1.3e+00 |
| Saturated fatty acids (mmol/l) | buffer-NMR | 42 | 21 | -9.5e-02 | -1.9e-01 | 2.9e-03 | 5.7e-02 | 1.3e+00 |
| **Glycolysis related metabolites** | | | | | | | | |
| Glucose (mmol/l) | thaw-buffer | 50 | 25 | 1.1e-02 | -9.7e-03 | 3.2e-02 | 2.9e-01 | 5.2e-01 |
| Glucose (mmol/l) | buffer-NMR | 50 | 25 | -1.0e-02 | -3.5e-02 | 1.5e-02 | 4.3e-01 | 5.2e-01 |
| Lactate (mmol/l) | thaw-buffer | 50 | 25 | -6.4e-03 | -2.1e-02 | 8.4e-03 | 4.0e-01 | 3.2e-01 |

| Metabolic traits | delay | N.obs | N.indiv | Beta | LCI | UCI | Pvalue | SD |
|---|---|---|---|---|---|---|---|---|
| Lactate (mmol/l) | buffer-NMR | 50 | 25 | -1.2e-02 | -2.7e-02 | 4.2e-03 | 1.5e-01 | 3.2e-01 |
| Pyruvate (mmol/l) | thaw-buffer | | | NA | NA | NA | NA | NA |
| Pyruvate (mmol/l) | buffer-NMR | | | NA | NA | NA | NA | NA |
| Citrate (mmol/l) | thaw-buffer | 50 | 25 | 8.8e-03 | 1.7e-03 | 1.6e-02 | 1.5e-02 | 3.4e-02 |
| Citrate (mmol/l) | buffer-NMR | 50 | 25 | 9.4e-03 | 1.6e-03 | 1.7e-02 | 1.9e-02 | 3.4e-02 |
| Glycerol (mmol/l) | thaw-buffer | | | NA | NA | NA | NA | NA |
| Glycerol (mmol/l) | buffer-NMR | | | NA | NA | NA | NA | NA |
| **Amino acids** | | | | | | | | |
| Alanine (mmol/l) | thaw-buffer | 50 | 25 | -5.5e-03 | -9.5e-03 | -1.5e-03 | 7.7e-03 | 5.2e-02 |
| Alanine (mmol/l) | buffer-NMR | 50 | 25 | -8.0e-04 | -4.7e-03 | 3.1e-03 | 6.9e-01 | 5.2e-02 |
| Glutamine (mmol/l) | thaw-buffer | 50 | 25 | -1.4e-02 | -2.6e-02 | -2.7e-03 | 1.5e-02 | 6.0e-02 |
| Glutamine (mmol/l) | buffer-NMR | 50 | 25 | -7.0e-03 | -1.4e-02 | -1.6e-04 | 4.5e-02 | 6.0e-02 |
| Histidine (mmol/l) | thaw-buffer | 50 | 25 | -1.3e-02 | -1.6e-02 | -8.9e-03 | 4.8e-11 | 9.3e-03 |
| Histidine (mmol/l) | buffer-NMR | 50 | 25 | -4.4e-03 | -7.2e-03 | -1.7e-03 | 1.4e-03 | 9.3e-03 |
| Glycine (mmol/l) | thaw-buffer | | | NA | NA | NA | NA | NA |
| Glycine (mmol/l) | buffer-NMR | | | NA | NA | NA | NA | NA |
| *Branched-chain amino acids* | | | | | | | | |
| Isoleucine (mmol/l) | thaw-buffer | 50 | 25 | -4.2e-03 | -6.1e-03 | -2.3e-03 | 1.3e-05 | 2.5e-02 |
| Isoleucine (mmol/l) | buffer-NMR | 50 | 25 | -3.5e-03 | -5.0e-03 | -1.9e-03 | 1.0e-05 | 2.5e-02 |
| Leucine (mmol/l) | thaw-buffer | 50 | 25 | -2.8e-03 | -3.6e-03 | -2.0e-03 | 1.9e-11 | 2.2e-02 |
| Leucine (mmol/l) | buffer-NMR | 50 | 25 | -4.6e-04 | -1.1e-03 | 2.2e-04 | 1.8e-01 | 2.2e-02 |
| Valine (mmol/l) | thaw-buffer | 50 | 25 | -2.3e-03 | -4.0e-03 | -5.2e-04 | 1.1e-02 | 3.6e-02 |
| Valine (mmol/l) | buffer-NMR | 50 | 25 | 4.7e-04 | -7.9e-04 | 1.7e-03 | 4.6e-01 | 3.6e-02 |
| *Aromatic amino acids* | | | | | | | | |
| Phenylalanine (mmol/l) | thaw-buffer | 50 | 25 | -2.9e-03 | -4.3e-03 | -1.4e-03 | 1.5e-04 | 6.7e-03 |
| Phenylalanine (mmol/l) | buffer-NMR | 50 | 25 | -1.1e-03 | -2.7e-03 | 5.5e-04 | 2.0e-01 | 6.7e-03 |
| Tyrosine (mmol/l) | thaw-buffer | 50 | 25 | 2.9e-04 | -1.0e-03 | 1.6e-03 | 6.7e-01 | 1.2e-02 |

| Metabolic traits | delay | N.obs | N.indiv | Beta | LCI | UCI | Pvalue | SD |
|---|---|---|---|---|---|---|---|---|
| Tyrosine (mmol/l) | buffer-NMR | 50 | 25 | 3.4e-04 | -1.1e-03 | 1.8e-03 | 6.6e-01 | 1.2e-02 |
| **Ketone bodies** | | | | | | | | |
| Acetate (mmol/l) | thaw-buffer | 50 | 25 | 2.7e-03 | 1.5e-03 | 3.9e-03 | 8.5e-06 | 9.7e-03 |
| Acetate (mmol/l) | buffer-NMR | 50 | 25 | -1.3e-03 | -2.4e-03 | -2.7e-04 | 1.3e-02 | 9.7e-03 |
| Beta-hydroxybutyrate (mmol/l) | thaw-buffer | 50 | 25 | 5.4e-03 | 3.7e-03 | 7.1e-03 | 2.3e-10 | 1.5e-02 |
| Beta-hydroxybutyrate (mmol/l) | buffer-NMR | 50 | 25 | 2.9e-03 | 1.0e-03 | 4.9e-03 | 3.0e-03 | 1.5e-02 |
| **Fluid balance** | | | | | | | | |
| Creatinine (mmol/l) | thaw-buffer | 50 | 25 | 2.8e-03 | 2.0e-03 | 3.6e-03 | 2.3e-12 | 8.9e-03 |
| Creatinine (mmol/l) | buffer-NMR | 50 | 25 | -9.2e-04 | -2.2e-03 | 3.5e-04 | 1.6e-01 | 8.9e-03 |
| Albumin (signal area) | thaw-buffer | 50 | 25 | 8.9e-04 | 2.6e-04 | 1.5e-03 | 5.5e-03 | 3.5e-03 |
| Albumin (signal area) | buffer-NMR | 50 | 25 | -4.6e-04 | -1.1e-03 | 2.2e-04 | 1.9e-01 | 3.5e-03 |
| **Inflammation** | | | | | | | | |
| Glycoprotein acetyls (mmol/l) | thaw-buffer | 50 | 25 | 6.6e-03 | -2.4e-03 | 1.6e-02 | 1.5e-01 | 3.2e-01 |
| Glycoprotein acetyls (mmol/l) | buffer-NMR | 50 | 25 | -2.8e-03 | -9.1e-03 | 3.6e-03 | 4.0e-01 | 3.2e-01 |

*sTable 6. Spearman correlation: serum, pre-storage handling effects. Spearman rank correlation coefficients between metabolic concentrations (or values) in reference conditions samples (4°C, 1.5h) and samples incubated at (i) 4°C, 24h; (ii) 4°C, 48h; (iii) 21°C, 24h; (iv) 21°C, 48h, before centrifugation.*

| Metabolic traits | 4°C,24h | 4°C,48h | 21°C,24h | 21°C,48h |
|---|---|---|---|---|
| **Lipoprotein subclasses** | | | | |
| *Extremely large VLDL* | | | | |
| Particle concentration (mol/l) | 9.2e-01 | 9.1e-01 | 8.9e-01 | 7.7e-01 |
| Total lipids (mmol/l) | 9.2e-01 | 9.1e-01 | 9.0e-01 | 7.7e-01 |
| Phospholipids (mmol/l) | 8.9e-01 | 9.2e-01 | 9.1e-01 | 7.9e-01 |
| Total cholesterol (mmol/l) | 9.5e-01 | 9.5e-01 | 9.4e-01 | 8.5e-01 |
| Cholesterol esters (mmol/l) | 9.5e-01 | 9.5e-01 | 9.3e-01 | 8.1e-01 |
| Free cholesterol (mmol/l) | 9.1e-01 | 9.3e-01 | 9.3e-01 | 8.3e-01 |
| Triglycerides (mmol/l) | 9.1e-01 | 9.1e-01 | 8.8e-01 | 7.6e-01 |
| *Very large VLDL* | | | | |
| Particle concentration (mol/l) | 9.6e-01 | 9.6e-01 | 9.5e-01 | 8.3e-01 |
| Total lipids (mmol/l) | 9.6e-01 | 9.6e-01 | 9.5e-01 | 8.3e-01 |
| Phospholipids (mmol/l) | 9.7e-01 | 9.7e-01 | 9.7e-01 | 8.5e-01 |
| Total cholesterol (mmol/l) | 9.7e-01 | 9.7e-01 | 9.7e-01 | 8.6e-01 |
| Cholesterol esters (mmol/l) | 9.7e-01 | 9.7e-01 | 9.7e-01 | 8.6e-01 |
| Free cholesterol (mmol/l) | 9.7e-01 | 9.7e-01 | 9.7e-01 | 8.5e-01 |
| Triglycerides (mmol/l) | 9.6e-01 | 9.6e-01 | 9.5e-01 | 8.3e-01 |
| *Large VLDL* | | | | |
| Particle concentration (mol/l) | 9.9e-01 | 9.8e-01 | 9.6e-01 | 9.0e-01 |
| Total lipids (mmol/l) | 9.8e-01 | 9.8e-01 | 9.6e-01 | 9.0e-01 |
| Phospholipids (mmol/l) | 9.8e-01 | 9.8e-01 | 9.6e-01 | 8.9e-01 |
| Total cholesterol (mmol/l) | 9.8e-01 | 9.8e-01 | 9.7e-01 | 8.8e-01 |
| Cholesterol esters (mmol/l) | 9.7e-01 | 9.7e-01 | 9.6e-01 | 8.2e-01 |
| Free cholesterol (mmol/l) | 9.8e-01 | 9.9e-01 | 9.8e-01 | 9.1e-01 |
| Triglycerides (mmol/l) | 9.8e-01 | 9.9e-01 | 9.6e-01 | 9.0e-01 |

| Metabolic traits | 4°C,24h | 4°C,48h | 21°C,24h | 21°C,48h |
|---|---|---|---|---|
| *Medium VLDL* | | | | |
| Particle concentration (mol/l) | 9.7e-01 | 9.8e-01 | 9.7e-01 | 9.0e-01 |
| Total lipids (mmol/l) | 9.7e-01 | 9.9e-01 | 9.6e-01 | 8.9e-01 |
| Phospholipids (mmol/l) | 9.8e-01 | 9.8e-01 | 9.6e-01 | 8.9e-01 |
| Total cholesterol (mmol/l) | 9.7e-01 | 9.7e-01 | 9.8e-01 | 8.8e-01 |
| Cholesterol esters (mmol/l) | 9.7e-01 | 9.7e-01 | 9.4e-01 | 8.6e-01 |
| Free cholesterol (mmol/l) | 9.9e-01 | 9.9e-01 | 9.7e-01 | 9.1e-01 |
| Triglycerides (mmol/l) | 9.8e-01 | 9.9e-01 | 9.7e-01 | 9.2e-01 |
| *Small VLDL* | | | | |
| Particle concentration (mol/l) | 9.8e-01 | 9.8e-01 | 9.6e-01 | 9.2e-01 |
| Total lipids (mmol/l) | 9.7e-01 | 9.8e-01 | 9.7e-01 | 9.3e-01 |
| Phospholipids (mmol/l) | 9.7e-01 | 9.7e-01 | 9.9e-01 | 9.5e-01 |
| Total cholesterol (mmol/l) | 9.8e-01 | 9.7e-01 | 9.8e-01 | 9.2e-01 |
| Cholesterol esters (mmol/l) | 9.8e-01 | 9.7e-01 | 9.6e-01 | 9.0e-01 |
| Free cholesterol (mmol/l) | 9.9e-01 | 9.9e-01 | 9.9e-01 | 9.5e-01 |
| Triglycerides (mmol/l) | 9.7e-01 | 9.6e-01 | 9.6e-01 | 9.0e-01 |
| *Very Small VLDL* | | | | |
| Particle concentration (mol/l) | 1.0e+00 | 9.9e-01 | 9.7e-01 | 9.2e-01 |
| Total lipids (mmol/l) | 9.9e-01 | 9.8e-01 | 9.7e-01 | 9.2e-01 |
| Phospholipids (mmol/l) | 9.8e-01 | 9.6e-01 | 9.5e-01 | 9.0e-01 |
| Total cholesterol (mmol/l) | 9.7e-01 | 9.4e-01 | 9.4e-01 | 8.2e-01 |
| Cholesterol esters (mmol/l) | 9.7e-01 | 9.5e-01 | 9.5e-01 | 7.7e-01 |
| Free cholesterol (mmol/l) | 9.7e-01 | 9.3e-01 | 9.4e-01 | 9.1e-01 |
| Triglycerides (mmol/l) | 9.9e-01 | 9.8e-01 | 9.9e-01 | 9.4e-01 |
| *IDL* | | | | |
| Particle concentration (mol/l) | 9.7e-01 | 9.7e-01 | 9.5e-01 | 9.2e-01 |
| Total lipids (mmol/l) | 9.7e-01 | 9.7e-01 | 9.3e-01 | 9.0e-01 |
| Phospholipids (mmol/l) | 9.5e-01 | 9.4e-01 | 8.9e-01 | 8.8e-01 |
| Total cholesterol (mmol/l) | 9.3e-01 | 9.3e-01 | 9.3e-01 | 8.8e-01 |

| Metabolic traits | 4°C,24h | 4°C,48h | 21°C,24h | 21°C,48h |
|---|---|---|---|---|
| Cholesterol esters (mmol/l) | 9.7e-01 | 9.5e-01 | 9.5e-01 | 9.0e-01 |
| Free cholesterol (mmol/l) | 9.3e-01 | 9.2e-01 | 8.4e-01 | 8.0e-01 |
| Triglycerides (mmol/l) | 1.0e+00 | 9.9e-01 | 9.7e-01 | 9.5e-01 |
| *Large LDL* | | | | |
| Particle concentration (mol/l) | 9.5e-01 | 9.3e-01 | 9.2e-01 | 9.0e-01 |
| Total lipids (mmol/l) | 9.5e-01 | 9.1e-01 | 9.1e-01 | 8.7e-01 |
| Phospholipids (mmol/l) | 9.4e-01 | 9.3e-01 | 9.1e-01 | 8.8e-01 |
| Total cholesterol (mmol/l) | 9.7e-01 | 9.3e-01 | 9.1e-01 | 8.8e-01 |
| Cholesterol esters (mmol/l) | 9.4e-01 | 9.3e-01 | 8.8e-01 | 9.0e-01 |
| Free cholesterol (mmol/l) | 9.5e-01 | 9.0e-01 | 8.2e-01 | 7.7e-01 |
| Triglycerides (mmol/l) | 9.9e-01 | 9.9e-01 | 9.6e-01 | 9.6e-01 |
| *Medium LDL* | | | | |
| Particle concentration (mol/l) | 9.3e-01 | 9.2e-01 | 8.8e-01 | 8.9e-01 |
| Total lipids (mmol/l) | 9.4e-01 | 9.3e-01 | 8.7e-01 | 9.1e-01 |
| Phospholipids (mmol/l) | 9.7e-01 | 9.5e-01 | 9.4e-01 | 9.1e-01 |
| Total cholesterol (mmol/l) | 9.5e-01 | 9.2e-01 | 8.5e-01 | 8.7e-01 |
| Cholesterol esters (mmol/l) | 9.6e-01 | 9.3e-01 | 8.6e-01 | 8.7e-01 |
| Free cholesterol (mmol/l) | 9.4e-01 | 9.1e-01 | 8.7e-01 | 8.8e-01 |
| Triglycerides (mmol/l) | 9.8e-01 | 9.8e-01 | 9.5e-01 | 9.4e-01 |
| *Small LDL* | | | | |
| Particle concentration (mol/l) | 9.5e-01 | 9.3e-01 | 9.1e-01 | 9.2e-01 |
| Total lipids (mmol/l) | 9.5e-01 | 9.2e-01 | 8.9e-01 | 9.3e-01 |
| Phospholipids (mmol/l) | 9.9e-01 | 9.7e-01 | 9.6e-01 | 9.4e-01 |
| Total cholesterol (mmol/l) | 9.4e-01 | 9.1e-01 | 8.4e-01 | 8.2e-01 |
| Cholesterol esters (mmol/l) | 9.6e-01 | 9.0e-01 | 8.3e-01 | 7.9e-01 |
| Free cholesterol (mmol/l) | 9.5e-01 | 9.2e-01 | 8.8e-01 | 9.1e-01 |
| Triglycerides (mmol/l) | 9.9e-01 | 9.9e-01 | 9.9e-01 | 9.7e-01 |
| *Very large HDL* | | | | |
| Particle concentration (mol/l) | 9.8e-01 | 9.8e-01 | 9.8e-01 | 9.0e-01 |

| Metabolic traits | 4°C,24h | 4°C,48h | 21°C,24h | 21°C,48h |
|---|---|---|---|---|
| Total lipids (mmol/l) | 9.8e-01 | 9.8e-01 | 9.8e-01 | 9.0e-01 |
| Phospholipids (mmol/l) | 9.9e-01 | 9.9e-01 | 9.8e-01 | 9.6e-01 |
| Total cholesterol (mmol/l) | 9.7e-01 | 9.8e-01 | 9.8e-01 | 8.5e-01 |
| Cholesterol esters (mmol/l) | 9.7e-01 | 9.8e-01 | 9.8e-01 | 8.4e-01 |
| Free cholesterol (mmol/l) | 9.8e-01 | 9.8e-01 | 9.7e-01 | 8.9e-01 |
| Triglycerides (mmol/l) | 9.9e-01 | 9.8e-01 | 9.8e-01 | 9.1e-01 |
| *Large HDL* | | | | |
| Particle concentration (mol/l) | 9.8e-01 | 9.8e-01 | 9.5e-01 | 9.4e-01 |
| Total lipids (mmol/l) | 9.8e-01 | 9.9e-01 | 9.5e-01 | 9.4e-01 |
| Phospholipids (mmol/l) | 9.8e-01 | 9.9e-01 | 9.4e-01 | 9.1e-01 |
| Total cholesterol (mmol/l) | 9.9e-01 | 1.0e+00 | 9.6e-01 | 9.5e-01 |
| Cholesterol esters (mmol/l) | 9.9e-01 | 9.9e-01 | 9.6e-01 | 9.5e-01 |
| Free cholesterol (mmol/l) | 9.9e-01 | 9.9e-01 | 9.6e-01 | 9.5e-01 |
| Triglycerides (mmol/l) | 8.9e-01 | 8.8e-01 | 8.5e-01 | 7.9e-01 |
| *Medium HDL* | | | | |
| Particle concentration (mol/l) | 9.8e-01 | 9.7e-01 | 9.4e-01 | 8.0e-01 |
| Total lipids (mmol/l) | 9.9e-01 | 9.7e-01 | 9.5e-01 | 8.0e-01 |
| Phospholipids (mmol/l) | 9.9e-01 | 9.7e-01 | 9.5e-01 | 8.1e-01 |
| Total cholesterol (mmol/l) | 9.9e-01 | 9.8e-01 | 9.7e-01 | 7.6e-01 |
| Cholesterol esters (mmol/l) | 9.8e-01 | 9.9e-01 | 9.6e-01 | 7.8e-01 |
| Free cholesterol (mmol/l) | 9.9e-01 | 9.7e-01 | 9.7e-01 | 7.6e-01 |
| Triglycerides (mmol/l) | 9.8e-01 | 9.7e-01 | 9.7e-01 | 9.3e-01 |
| *Small HDL* | | | | |
| Particle concentration (mol/l) | 9.5e-01 | 9.5e-01 | 9.5e-01 | 6.5e-01 |
| Total lipids (mmol/l) | 9.6e-01 | 9.5e-01 | 9.3e-01 | 7.0e-01 |
| Phospholipids (mmol/l) | 9.7e-01 | 9.6e-01 | 9.2e-01 | 6.1e-01 |
| Total cholesterol (mmol/l) | 8.9e-01 | 8.4e-01 | 8.5e-01 | 6.2e-01 |
| Cholesterol esters (mmol/l) | 9.0e-01 | 9.0e-01 | 8.3e-01 | 6.7e-01 |
| Free cholesterol (mmol/l) | 9.5e-01 | 9.8e-01 | 9.1e-01 | 6.0e-01 |

| Metabolic traits | 4°C,24h | 4°C,48h | 21°C,24h | 21°C,48h |
|---|---|---|---|---|
| Triglycerides (mmol/l) | 9.9e-01 | 9.8e-01 | 9.8e-01 | 9.7e-01 |
| **Lipoprotein particle size** | | | | |
| VLDL particle size (nm) | 9.8e-01 | 9.7e-01 | 9.5e-01 | 9.1e-01 |
| LDL particle size (nm) | 9.5e-01 | 9.4e-01 | 9.4e-01 | 7.0e-01 |
| HDL particle size (nm) | 9.9e-01 | 9.9e-01 | 9.9e-01 | 9.7e-01 |
| **Cholesterol** | | | | |
| Total cholesterol (mmol/l) | 9.7e-01 | 9.5e-01 | 9.3e-01 | 9.3e-01 |
| VLDL cholesterol (mmol/l) | 9.8e-01 | 9.8e-01 | 9.7e-01 | 8.9e-01 |
| Remnant cholesterol (mmol/l) | 9.8e-01 | 9.7e-01 | 9.7e-01 | 9.2e-01 |
| LDL cholesterol (mmol/l) | 9.6e-01 | 9.3e-01 | 8.6e-01 | 9.0e-01 |
| HDL cholesterol (mmol/l) | 9.9e-01 | 9.8e-01 | 9.5e-01 | 8.7e-01 |
| HDL2 cholesterol (mmol/l) | 9.9e-01 | 9.8e-01 | 9.5e-01 | 9.0e-01 |
| HDL3 cholesterol (mmol/l) | 9.8e-01 | 9.5e-01 | 8.7e-01 | 4.7e-01 |
| Esterified cholesterol (mmol/l) | 9.5e-01 | 9.2e-01 | 9.1e-01 | 8.6e-01 |
| Free cholesterol (mmol/l) | 9.2e-01 | 9.8e-01 | 9.5e-01 | 9.6e-01 |
| **Glycerides and phospholipids** | | | | |
| Triglycerides (mmol/l) | 9.9e-01 | 9.9e-01 | 9.8e-01 | 9.2e-01 |
| VLDL triglycerides (mmol/l) | 9.9e-01 | 9.8e-01 | 9.6e-01 | 8.8e-01 |
| LDL triglycerides (mmol/l) | 1.0e+00 | 9.9e-01 | 9.7e-01 | 9.5e-01 |
| HDL triglycerides (mmol/l) | 1.0e+00 | 1.0e+00 | 9.9e-01 | 9.8e-01 |
| Diacylglycerol (mmol/l) | 3.2e-01 | 1.9e-01 | 2.8e-01 | 5.3e-01 |
| Phosphoglycerides (mmol/l) | 9.5e-01 | 9.5e-01 | 9.1e-01 | 8.9e-01 |
| Phosphatidylcholine + other cholines (mmol/l) | 9.5e-01 | 9.5e-01 | 9.4e-01 | 8.9e-01 |
| Sphingomyelins (mmol/l) | 6.9e-01 | 6.9e-01 | 6.6e-01 | 6.5e-01 |
| Cholines (mmol/l) | 8.5e-01 | 8.3e-01 | 7.8e-01 | 6.7e-01 |
| **Apolipoproteins** | | | | |
| Apolipoprotein A-I (g/l) | 9.8e-01 | 9.8e-01 | 9.7e-01 | 8.5e-01 |
| Apolipoprotein B (g/l) | 9.9e-01 | 9.8e-01 | 9.7e-01 | 9.5e-01 |
| **Fatty acids** | | | | |

| Metabolic traits | 4°C,24h | 4°C,48h | 21°C,24h | 21°C,48h |
|---|---|---|---|---|
| Total fatty acids (mmol/l) | 9.5e-01 | 9.7e-01 | 9.6e-01 | 9.6e-01 |
| Fatty acid chain length | 7.6e-01 | 7.9e-01 | 8.8e-01 | 8.5e-01 |
| Degree of unsaturation | 9.2e-01 | 9.5e-01 | 9.0e-01 | 9.1e-01 |
| Docosahexaenoic acid (mmol/l) | 9.1e-01 | 9.1e-01 | 9.5e-01 | 9.4e-01 |
| Linoleic acid (mmol/l) | 9.7e-01 | 9.7e-01 | 9.7e-01 | 9.5e-01 |
| Conjugated linoleic acid (mmol/l) | 5.4e-01 | 5.5e-01 | 7.1e-01 | 6.3e-01 |
| n-3 fatty acids (mmol/l) | 8.8e-01 | 9.1e-01 | 9.7e-01 | 9.5e-01 |
| n-6 fatty acids (mmol/l) | 9.4e-01 | 9.5e-01 | 9.6e-01 | 9.5e-01 |
| PUFA (mmol/l) | 9.5e-01 | 9.7e-01 | 9.8e-01 | 9.6e-01 |
| MUFA (mmol/l) | 9.8e-01 | 9.9e-01 | 9.7e-01 | 9.7e-01 |
| Saturated fatty acids (mmol/l) | 9.7e-01 | 9.8e-01 | 9.5e-01 | 9.7e-01 |
| **Glycolysis related metabolites** | | | | |
| Glucose (mmol/l) | 9.7e-01 | 9.1e-01 | 7.0e-01 | 4.8e-01 |
| Lactate (mmol/l) | 7.3e-01 | 5.5e-01 | 3.3e-01 | 5.2e-01 |
| Pyruvate (mmol/l) | 4.4e-01 | 2.8e-01 | -6.5e-02 | -6.6e-02 |
| Citrate (mmol/l) | 8.6e-01 | 8.2e-01 | 8.0e-01 | 8.7e-01 |
| Glycerol (mmol/l) | 9.0e-01 | 4.9e-01 | 6.7e-01 | 5.1e-01 |
| **Amino acids** | | | | |
| Alanine (mmol/l) | 9.5e-01 | 9.2e-01 | 8.6e-01 | 5.6e-01 |
| Glutamine (mmol/l) | 9.6e-01 | 9.4e-01 | 9.2e-01 | 8.1e-01 |
| Histidine (mmol/l) | 3.9e-01 | 5.6e-01 | 5.0e-01 | 4.8e-01 |
| Glycine (mmol/l) | 8.9e-01 | 8.0e-01 | 8.9e-01 | 7.7e-01 |
| *Branched-chain amino acids* | | | | |
| Isoleucine (mmol/l) | 9.8e-01 | 9.8e-01 | 9.2e-01 | 9.4e-01 |
| Leucine (mmol/l) | 9.8e-01 | 9.7e-01 | 9.1e-01 | 9.2e-01 |
| Valine (mmol/l) | 9.8e-01 | 9.8e-01 | 9.6e-01 | 9.5e-01 |
| *Aromatic amino acids* | | | | |
| Phenylalanine (mmol/l) | 7.4e-01 | 7.3e-01 | 7.7e-01 | 6.5e-01 |
| Tyrosine (mmol/l) | 8.9e-01 | 8.9e-01 | 8.9e-01 | 8.6e-01 |

| Metabolic traits | 4°C,24h | 4°C,48h | 21°C,24h | 21°C,48h |
|---|---|---|---|---|
| **Ketone bodies** | | | | |
| Acetate (mmol/l) | 8.3e-01 | 7.0e-01 | 4.5e-01 | 6.0e-01 |
| Beta-hydroxybutyrate (mmol/l) | 8.7e-01 | 7.3e-01 | 8.6e-01 | 7.3e-01 |
| **Fluid balance** | | | | |
| Creatinine (mmol/l) | 9.3e-01 | 9.6e-01 | 9.4e-01 | 9.5e-01 |
| Albumin (signal area) | 9.6e-01 | 9.1e-01 | 8.5e-01 | 9.0e-01 |
| **Inflammation** | | | | |
| Glycoprotein acetyls (mmol/l) | 9.9e-01 | 9.9e-01 | 9.9e-01 | 9.7e-01 |

*sTable 7. Spearman correlation: EDTA-plasma, pre-storage handling effects. Spearman rank correlation coefficients between metabolic concentrations (or values) in reference conditions samples (4°C, 1.5h) and samples incubated at (i) 4°C, 24h; (ii) 4°C, 48h; (iii) 21°C, 24h; (iv) 21°C, 48h, before centrifugation. Pyruvate, glycerol and glycine are not quantified in EDTA - plasma samples due to the interfering resonances of EDTA on their signals.*

| Metabolic traits | 4°C,24h | 4°C,48h | 21°C,24h | 21°C,48h |
|---|---|---|---|---|
| **Lipoprotein subclasses** | | | | |
| *Extremely large VLDL* | | | | |
| Particle concentration (mol/l) | 8.5e-01 | 9.5e-01 | 9.7e-01 | 7.3e-01 |
| Total lipids (mmol/l) | 8.5e-01 | 9.5e-01 | 9.7e-01 | 7.3e-01 |
| Phospholipids (mmol/l) | 8.9e-01 | 9.7e-01 | 9.8e-01 | 7.9e-01 |
| Total cholesterol (mmol/l) | 8.5e-01 | 9.5e-01 | 9.7e-01 | 7.5e-01 |
| Cholesterol esters (mmol/l) | 8.3e-01 | 9.4e-01 | 9.5e-01 | 7.4e-01 |
| Free cholesterol (mmol/l) | 9.0e-01 | 9.7e-01 | 9.8e-01 | 8.0e-01 |
| Triglycerides (mmol/l) | 8.5e-01 | 9.6e-01 | 9.7e-01 | 7.2e-01 |
| *Very large VLDL* | | | | |
| Particle concentration (mol/l) | 9.3e-01 | 9.6e-01 | 9.9e-01 | 8.6e-01 |
| Total lipids (mmol/l) | 9.3e-01 | 9.6e-01 | 9.9e-01 | 8.7e-01 |
| Phospholipids (mmol/l) | 9.2e-01 | 9.6e-01 | 9.9e-01 | 8.6e-01 |
| Total cholesterol (mmol/l) | 9.0e-01 | 9.5e-01 | 9.9e-01 | 8.1e-01 |
| Cholesterol esters (mmol/l) | 8.8e-01 | 9.5e-01 | 9.9e-01 | 8.0e-01 |
| Free cholesterol (mmol/l) | 9.2e-01 | 9.5e-01 | 9.9e-01 | 8.6e-01 |
| Triglycerides (mmol/l) | 9.3e-01 | 9.7e-01 | 9.9e-01 | 8.7e-01 |
| *Large VLDL* | | | | |
| Particle concentration (mol/l) | 9.7e-01 | 9.7e-01 | 9.6e-01 | 8.6e-01 |
| Total lipids (mmol/l) | 9.6e-01 | 9.7e-01 | 9.7e-01 | 8.7e-01 |
| Phospholipids (mmol/l) | 9.5e-01 | 9.8e-01 | 9.6e-01 | 8.8e-01 |
| Total cholesterol (mmol/l) | 9.5e-01 | 9.7e-01 | 9.8e-01 | 8.8e-01 |
| Cholesterol esters (mmol/l) | 9.2e-01 | 9.5e-01 | 9.7e-01 | 8.5e-01 |
| Free cholesterol (mmol/l) | 9.5e-01 | 9.7e-01 | 9.9e-01 | 8.8e-01 |

| Metabolic traits | 4°C,24h | 4°C,48h | 21°C,24h | 21°C,48h |
|---|---|---|---|---|
| Triglycerides (mmol/l) | 9.8e-01 | 9.7e-01 | 9.4e-01 | 8.6e-01 |
| *Medium VLDL* | | | | |
| Particle concentration (mol/l) | 9.2e-01 | 9.8e-01 | 9.4e-01 | 8.4e-01 |
| Total lipids (mmol/l) | 9.3e-01 | 9.9e-01 | 9.4e-01 | 8.3e-01 |
| Phospholipids (mmol/l) | 9.2e-01 | 9.9e-01 | 9.3e-01 | 8.3e-01 |
| Total cholesterol (mmol/l) | 9.1e-01 | 9.5e-01 | 9.7e-01 | 8.8e-01 |
| Cholesterol esters (mmol/l) | 9.3e-01 | 9.4e-01 | 9.5e-01 | 8.6e-01 |
| Free cholesterol (mmol/l) | 9.6e-01 | 9.7e-01 | 9.6e-01 | 8.7e-01 |
| Triglycerides (mmol/l) | 9.6e-01 | 9.9e-01 | 9.6e-01 | 9.0e-01 |
| *Small VLDL* | | | | |
| Particle concentration (mol/l) | 9.5e-01 | 9.8e-01 | 9.8e-01 | 9.1e-01 |
| Total lipids (mmol/l) | 9.4e-01 | 9.8e-01 | 9.7e-01 | 9.1e-01 |
| Phospholipids (mmol/l) | 9.6e-01 | 9.8e-01 | 9.6e-01 | 9.2e-01 |
| Total cholesterol (mmol/l) | 9.4e-01 | 9.8e-01 | 9.8e-01 | 9.7e-01 |
| Cholesterol esters (mmol/l) | 9.4e-01 | 9.6e-01 | 9.7e-01 | 9.5e-01 |
| Free cholesterol (mmol/l) | 9.8e-01 | 9.8e-01 | 9.8e-01 | 9.5e-01 |
| Triglycerides (mmol/l) | 9.6e-01 | 9.9e-01 | 9.7e-01 | 8.9e-01 |
| *Very Small VLDL* | | | | |
| Particle concentration (mol/l) | 9.6e-01 | 9.7e-01 | 9.7e-01 | 9.3e-01 |
| Total lipids (mmol/l) | 9.6e-01 | 9.8e-01 | 9.7e-01 | 9.4e-01 |
| Phospholipids (mmol/l) | 9.2e-01 | 9.8e-01 | 9.5e-01 | 9.4e-01 |
| Total cholesterol (mmol/l) | 8.8e-01 | 9.5e-01 | 9.1e-01 | 8.7e-01 |
| Cholesterol esters (mmol/l) | 8.3e-01 | 9.3e-01 | 8.8e-01 | 8.1e-01 |
| Free cholesterol (mmol/l) | 9.2e-01 | 9.9e-01 | 9.8e-01 | 9.7e-01 |
| Triglycerides (mmol/l) | 9.5e-01 | 9.6e-01 | 9.4e-01 | 9.3e-01 |
| *IDL* | | | | |
| Particle concentration (mol/l) | 9.5e-01 | 9.9e-01 | 9.6e-01 | 9.4e-01 |
| Total lipids (mmol/l) | 9.4e-01 | 9.8e-01 | 9.5e-01 | 9.3e-01 |
| Phospholipids (mmol/l) | 9.4e-01 | 9.7e-01 | 9.2e-01 | 8.8e-01 |

| Metabolic traits | 4°C,24h | 4°C,48h | 21°C,24h | 21°C,48h |
|---|---|---|---|---|
| Total cholesterol (mmol/l) | 9.0e-01 | 9.5e-01 | 9.1e-01 | 9.0e-01 |
| Cholesterol esters (mmol/l) | 9.0e-01 | 9.4e-01 | 9.0e-01 | 8.9e-01 |
| Free cholesterol (mmol/l) | 9.3e-01 | 9.5e-01 | 8.9e-01 | 8.5e-01 |
| Triglycerides (mmol/l) | 9.7e-01 | 9.8e-01 | 9.6e-01 | 9.5e-01 |
| *Large LDL* | | | | |
| Particle concentration (mol/l) | 9.6e-01 | 9.7e-01 | 9.4e-01 | 9.3e-01 |
| Total lipids (mmol/l) | 9.5e-01 | 9.6e-01 | 9.3e-01 | 9.1e-01 |
| Phospholipids (mmol/l) | 9.4e-01 | 9.6e-01 | 9.4e-01 | 9.3e-01 |
| Total cholesterol (mmol/l) | 9.4e-01 | 9.6e-01 | 9.4e-01 | 9.2e-01 |
| Cholesterol esters (mmol/l) | 9.4e-01 | 9.6e-01 | 9.5e-01 | 9.2e-01 |
| Free cholesterol (mmol/l) | 9.3e-01 | 9.5e-01 | 9.0e-01 | 8.0e-01 |
| Triglycerides (mmol/l) | 9.8e-01 | 9.8e-01 | 9.7e-01 | 9.7e-01 |
| *Medium LDL* | | | | |
| Particle concentration (mol/l) | 9.6e-01 | 9.7e-01 | 9.6e-01 | 9.3e-01 |
| Total lipids (mmol/l) | 9.6e-01 | 9.6e-01 | 9.5e-01 | 9.1e-01 |
| Phospholipids (mmol/l) | 9.4e-01 | 9.7e-01 | 9.7e-01 | 9.5e-01 |
| Total cholesterol (mmol/l) | 9.5e-01 | 9.7e-01 | 9.3e-01 | 9.1e-01 |
| Cholesterol esters (mmol/l) | 9.6e-01 | 9.7e-01 | 9.4e-01 | 9.1e-01 |
| Free cholesterol (mmol/l) | 9.5e-01 | 9.7e-01 | 9.4e-01 | 8.8e-01 |
| Triglycerides (mmol/l) | 9.4e-01 | 9.8e-01 | 9.6e-01 | 9.4e-01 |
| *Small LDL* | | | | |
| Particle concentration (mol/l) | 9.6e-01 | 9.8e-01 | 9.5e-01 | 9.2e-01 |
| Total lipids (mmol/l) | 9.6e-01 | 9.8e-01 | 9.5e-01 | 9.1e-01 |
| Phospholipids (mmol/l) | 9.3e-01 | 9.7e-01 | 9.6e-01 | 9.1e-01 |
| Total cholesterol (mmol/l) | 9.4e-01 | 9.8e-01 | 9.2e-01 | 8.9e-01 |
| Cholesterol esters (mmol/l) | 9.7e-01 | 9.9e-01 | 9.5e-01 | 8.9e-01 |
| Free cholesterol (mmol/l) | 9.5e-01 | 9.8e-01 | 9.3e-01 | 8.3e-01 |
| Triglycerides (mmol/l) | 9.7e-01 | 9.9e-01 | 9.7e-01 | 9.6e-01 |
| *Very large HDL* | | | | |

| Metabolic traits | 4°C,24h | 4°C,48h | 21°C,24h | 21°C,48h |
|---|---|---|---|---|
| Particle concentration (mol/l) | 9.9e-01 | 9.9e-01 | 9.8e-01 | 9.8e-01 |
| Total lipids (mmol/l) | 9.9e-01 | 9.8e-01 | 9.8e-01 | 9.7e-01 |
| Phospholipids (mmol/l) | 1.0e+00 | 9.9e-01 | 9.9e-01 | 9.9e-01 |
| Total cholesterol (mmol/l) | 9.7e-01 | 9.6e-01 | 9.7e-01 | 9.6e-01 |
| Cholesterol esters (mmol/l) | 9.6e-01 | 9.7e-01 | 9.6e-01 | 9.7e-01 |
| Free cholesterol (mmol/l) | 1.0e+00 | 9.8e-01 | 9.7e-01 | 9.7e-01 |
| Triglycerides (mmol/l) | 9.2e-01 | 9.5e-01 | 8.8e-01 | 8.9e-01 |
| *Large HDL* | | | | |
| Particle concentration (mol/l) | 9.6e-01 | 9.7e-01 | 9.8e-01 | 9.5e-01 |
| Total lipids (mmol/l) | 9.6e-01 | 9.7e-01 | 9.8e-01 | 9.6e-01 |
| Phospholipids (mmol/l) | 9.2e-01 | 9.7e-01 | 9.8e-01 | 9.4e-01 |
| Total cholesterol (mmol/l) | 9.8e-01 | 9.8e-01 | 9.8e-01 | 9.6e-01 |
| Cholesterol esters (mmol/l) | 9.8e-01 | 9.8e-01 | 9.9e-01 | 9.7e-01 |
| Free cholesterol (mmol/l) | 9.6e-01 | 9.8e-01 | 9.9e-01 | 9.7e-01 |
| Triglycerides (mmol/l) | 8.7e-01 | 8.7e-01 | 8.3e-01 | 9.5e-01 |
| *Medium HDL* | | | | |
| Particle concentration (mol/l) | 8.1e-01 | 9.3e-01 | 9.1e-01 | 7.0e-01 |
| Total lipids (mmol/l) | 8.1e-01 | 9.3e-01 | 9.1e-01 | 7.0e-01 |
| Phospholipids (mmol/l) | 8.4e-01 | 9.4e-01 | 9.2e-01 | 7.2e-01 |
| Total cholesterol (mmol/l) | 8.0e-01 | 9.3e-01 | 9.0e-01 | 7.4e-01 |
| Cholesterol esters (mmol/l) | 7.8e-01 | 9.3e-01 | 9.0e-01 | 7.4e-01 |
| Free cholesterol (mmol/l) | 8.3e-01 | 9.4e-01 | 8.9e-01 | 7.1e-01 |
| Triglycerides (mmol/l) | 9.6e-01 | 9.8e-01 | 9.5e-01 | 9.0e-01 |
| *Small HDL* | | | | |
| Particle concentration (mol/l) | 7.9e-01 | 8.8e-01 | 8.6e-01 | 4.9e-01 |
| Total lipids (mmol/l) | 8.0e-01 | 8.7e-01 | 8.7e-01 | 4.7e-01 |
| Phospholipids (mmol/l) | 8.6e-01 | 9.7e-01 | 9.6e-01 | 6.1e-01 |
| Total cholesterol (mmol/l) | 7.4e-01 | 8.8e-01 | 8.2e-01 | 4.8e-01 |
| Cholesterol esters (mmol/l) | 7.1e-01 | 9.0e-01 | 7.9e-01 | 5.8e-01 |

| Metabolic traits | 4°C,24h | 4°C,48h | 21°C,24h | 21°C,48h |
|---|---|---|---|---|
| Free cholesterol (mmol/l) | 7.9e-01 | 9.7e-01 | 9.4e-01 | 5.7e-01 |
| Triglycerides (mmol/l) | 9.5e-01 | 9.7e-01 | 9.8e-01 | 9.8e-01 |
| **Lipoprotein particle size** | | | | |
| VLDL particle size (nm) | 9.7e-01 | 9.9e-01 | 9.8e-01 | 8.7e-01 |
| LDL particle size (nm) | 7.6e-01 | 8.4e-01 | 8.0e-01 | 3.4e-01 |
| HDL particle size (nm) | 9.8e-01 | 9.8e-01 | 9.9e-01 | 9.9e-01 |
| **Cholesterol** | | | | |
| Total cholesterol (mmol/l) | 9.6e-01 | 9.8e-01 | 9.8e-01 | 9.6e-01 |
| VLDL cholesterol (mmol/l) | 9.1e-01 | 9.3e-01 | 9.6e-01 | 9.2e-01 |
| Remnant cholesterol (mmol/l) | 9.3e-01 | 9.7e-01 | 9.3e-01 | 9.0e-01 |
| LDL cholesterol (mmol/l) | 9.5e-01 | 9.6e-01 | 9.4e-01 | 9.1e-01 |
| HDL cholesterol (mmol/l) | 9.3e-01 | 9.8e-01 | 9.5e-01 | 9.2e-01 |
| HDL2 cholesterol (mmol/l) | 9.3e-01 | 9.8e-01 | 9.8e-01 | 9.4e-01 |
| HDL3 cholesterol (mmol/l) | 9.2e-01 | 9.7e-01 | 8.2e-01 | 6.8e-01 |
| Esterified cholesterol (mmol/l) | 9.6e-01 | 9.7e-01 | 9.5e-01 | 9.4e-01 |
| Free cholesterol (mmol/l) | 9.1e-01 | 9.6e-01 | 9.5e-01 | 9.5e-01 |
| **Glycerides and phospholipids** | | | | |
| Triglycerides (mmol/l) | 9.7e-01 | 1.0e+00 | 9.6e-01 | 8.9e-01 |
| VLDL triglycerides (mmol/l) | 9.6e-01 | 9.9e-01 | 9.4e-01 | 8.8e-01 |
| LDL triglycerides (mmol/l) | 9.6e-01 | 9.8e-01 | 9.7e-01 | 9.6e-01 |
| HDL triglycerides (mmol/l) | 9.9e-01 | 9.9e-01 | 9.9e-01 | 9.5e-01 |
| Diacylglycerol (mmol/l) | 5.9e-01 | 4.4e-01 | 1.1e-01 | 3.6e-01 |
| Phosphoglycerides (mmol/l) | 9.1e-01 | 9.3e-01 | 9.7e-01 | 8.9e-01 |
| Phosphatidylcholine + other cholines (mmol/l) | 8.7e-01 | 9.2e-01 | 9.7e-01 | 9.1e-01 |
| Sphingomyelins (mmol/l) | 8.3e-01 | 6.7e-01 | 5.4e-01 | 8.6e-01 |
| Cholines (mmol/l) | 9.1e-01 | 9.2e-01 | 9.5e-01 | 8.8e-01 |
| **Apolipoproteins** | | | | |
| Apolipoprotein A-I (g/l) | 9.3e-01 | 9.8e-01 | 9.8e-01 | 8.7e-01 |
| Apolipoprotein B (g/l) | 9.1e-01 | 9.4e-01 | 9.3e-01 | 9.0e-01 |

| Metabolic traits | 4°C,24h | 4°C,48h | 21°C,24h | 21°C,48h |
|---|---|---|---|---|
| **Fatty acids** | | | | |
| Total fatty acids (mmol/l) | 9.4e-01 | 9.8e-01 | 9.7e-01 | 9.6e-01 |
| Fatty acid chain length | 7.2e-01 | 7.3e-01 | 7.3e-01 | 7.1e-01 |
| Degree of unsaturation | 9.1e-01 | 9.6e-01 | 8.3e-01 | 7.3e-01 |
| Docosahexaenoic acid (mmol/l) | 9.8e-01 | 9.7e-01 | 9.8e-01 | 9.2e-01 |
| Linoleic acid (mmol/l) | 9.4e-01 | 9.6e-01 | 9.5e-01 | 9.5e-01 |
| Conjugated linoleic acid (mmol/l) | 8.6e-01 | 5.6e-01 | 9.1e-01 | 5.8e-01 |
| n-3 fatty acids (mmol/l) | 9.6e-01 | 9.5e-01 | 9.8e-01 | 9.2e-01 |
| n-6 fatty acids (mmol/l) | 9.6e-01 | 9.6e-01 | 9.6e-01 | 9.4e-01 |
| PUFA (mmol/l) | 9.5e-01 | 9.8e-01 | 9.6e-01 | 9.3e-01 |
| MUFA (mmol/l) | 9.8e-01 | 9.9e-01 | 9.9e-01 | 9.9e-01 |
| Saturated fatty acids (mmol/l) | 9.1e-01 | 9.6e-01 | 9.5e-01 | 9.4e-01 |
| **Glycolysis related metabolites** | | | | |
| Glucose (mmol/l) | 9.1e-01 | 7.2e-01 | 7.7e-01 | 4.8e-01 |
| Lactate (mmol/l) | 7.5e-01 | 3.3e-02 | 2.6e-01 | 5.2e-01 |
| Pyruvate (mmol/l) | NA | NA | NA | NA |
| Citrate (mmol/l) | 8.5e-01 | 8.8e-01 | 9.0e-01 | 7.8e-01 |
| Glycerol (mmol/l) | NA | NA | NA | NA |
| **Amino acids** | | | | |
| Alanine (mmol/l) | 9.5e-01 | 9.7e-01 | 8.0e-01 | 4.4e-01 |
| Glutamine (mmol/l) | 9.5e-01 | 9.0e-01 | 9.7e-01 | 8.5e-01 |
| Histidine (mmol/l) | 7.9e-01 | 3.0e-01 | 6.5e-01 | 1.6e-01 |
| Glycine (mmol/l) | NA | NA | NA | NA |
| *Branched-chain amino acids* | | | | |
| Isoleucine (mmol/l) | 9.1e-01 | 9.4e-01 | 9.2e-01 | 8.6e-01 |
| Leucine (mmol/l) | 9.8e-01 | 9.7e-01 | 9.5e-01 | 9.0e-01 |
| Valine (mmol/l) | 9.7e-01 | 9.8e-01 | 9.8e-01 | 9.8e-01 |
| *Aromatic amino acids* | | | | |
| Phenylalanine (mmol/l) | 6.2e-01 | 5.4e-01 | 6.8e-01 | 3.2e-01 |

| Metabolic traits | 4°C,24h | 4°C,48h | 21°C,24h | 21°C,48h |
|---|---|---|---|---|
| Tyrosine (mmol/l) | 9.2e-01 | 9.1e-01 | 9.4e-01 | 8.9e-01 |
| **Ketone bodies** | | | | |
| Acetate (mmol/l) | 4.9e-01 | 2.8e-01 | 2.8e-01 | 4.1e-01 |
| Beta-hydroxybutyrate (mmol/l) | 9.7e-01 | 8.9e-01 | 8.8e-01 | 6.7e-01 |
| **Fluid balance** | | | | |
| Creatinine (mmol/l) | 9.4e-01 | 9.3e-01 | 9.3e-01 | 9.0e-01 |
| Albumin (signal area) | 8.6e-01 | 9.1e-01 | 8.7e-01 | 7.2e-01 |
| **Inflammation** | | | | |
| Glycoprotein acetyls (mmol/l) | 9.8e-01 | 9.9e-01 | 9.8e-01 | 9.6e-01 |

*sTable 8. Spearman correlation: serum, post-storage handling effects. Spearman rank correlation coefficients between metabolic concentrations (or values) in reference conditions samples (i.e. no sample preparation or NMR analysis delays) and samples (i) left for 24h before addition of sodium buffer followed by immediate NMR analysis (i.e. thaw to buffer addition delay); (ii) thawed overnight, addition of sodium buffer, then left for 24h before NMR analysis (buffer addition to NMR profiling delay).*

| Metabolic traits | thaw-buffer delay | buffer-nmr delay |
|---|---|---|
| **Lipoprotein subclasses** | | |
| *Extremely large VLDL* | | |
| Particle concentration (mol/l) | 8.9e-01 | 9.7e-01 |
| Total lipids (mmol/l) | 8.9e-01 | 9.7e-01 |
| Phospholipids (mmol/l) | 9.0e-01 | 9.7e-01 |
| Total cholesterol (mmol/l) | 8.9e-01 | 9.7e-01 |
| Cholesterol esters (mmol/l) | 8.6e-01 | 9.2e-01 |
| Free cholesterol (mmol/l) | 9.1e-01 | 9.7e-01 |
| Triglycerides (mmol/l) | 9.0e-01 | 9.7e-01 |
| *Very large VLDL* | | |
| Particle concentration (mol/l) | 9.1e-01 | 9.8e-01 |
| Total lipids (mmol/l) | 9.1e-01 | 9.8e-01 |
| Phospholipids (mmol/l) | 9.2e-01 | 9.7e-01 |
| Total cholesterol (mmol/l) | 9.3e-01 | 9.8e-01 |
| Cholesterol esters (mmol/l) | 9.4e-01 | 9.7e-01 |
| Free cholesterol (mmol/l) | 9.3e-01 | 9.8e-01 |
| Triglycerides (mmol/l) | 9.0e-01 | 9.8e-01 |
| *Large VLDL* | | |
| Particle concentration (mol/l) | 9.9e-01 | 9.9e-01 |
| Total lipids (mmol/l) | 9.9e-01 | 9.9e-01 |
| Phospholipids (mmol/l) | 9.8e-01 | 9.9e-01 |
| Total cholesterol (mmol/l) | 9.8e-01 | 9.9e-01 |
| Cholesterol esters (mmol/l) | 9.7e-01 | 9.9e-01 |

| Metabolic traits | thaw-buffer delay | buffer-nmr delay |
|---|---|---|
| Free cholesterol (mmol/l) | 9.8e-01 | 1.0e+00 |
| Triglycerides (mmol/l) | 9.7e-01 | 9.9e-01 |
| *Medium VLDL* | | |
| Particle concentration (mol/l) | 1.0e+00 | 1.0e+00 |
| Total lipids (mmol/l) | 9.9e-01 | 9.9e-01 |
| Phospholipids (mmol/l) | 9.9e-01 | 9.9e-01 |
| Total cholesterol (mmol/l) | 9.6e-01 | 9.9e-01 |
| Cholesterol esters (mmol/l) | 9.6e-01 | 9.8e-01 |
| Free cholesterol (mmol/l) | 9.9e-01 | 1.0e+00 |
| Triglycerides (mmol/l) | 9.9e-01 | 1.0e+00 |
| *Small VLDL* | | |
| Particle concentration (mol/l) | 9.8e-01 | 9.9e-01 |
| Total lipids (mmol/l) | 9.8e-01 | 9.9e-01 |
| Phospholipids (mmol/l) | 9.9e-01 | 9.9e-01 |
| Total cholesterol (mmol/l) | 9.6e-01 | 9.8e-01 |
| Cholesterol esters (mmol/l) | 9.7e-01 | 9.7e-01 |
| Free cholesterol (mmol/l) | 9.9e-01 | 9.9e-01 |
| Triglycerides (mmol/l) | 9.9e-01 | 9.9e-01 |
| *Very Small VLDL* | | |
| Particle concentration (mol/l) | 9.5e-01 | 9.9e-01 |
| Total lipids (mmol/l) | 9.4e-01 | 9.9e-01 |
| Phospholipids (mmol/l) | 9.6e-01 | 9.9e-01 |
| Total cholesterol (mmol/l) | 8.6e-01 | 9.3e-01 |
| Cholesterol esters (mmol/l) | 8.6e-01 | 9.1e-01 |
| Free cholesterol (mmol/l) | 9.4e-01 | 9.8e-01 |
| Triglycerides (mmol/l) | 9.9e-01 | 9.9e-01 |
| *IDL* | | |
| Particle concentration (mol/l) | 9.7e-01 | 9.9e-01 |
| Total lipids (mmol/l) | 9.7e-01 | 9.8e-01 |

| Metabolic traits | thaw-buffer delay | buffer-nmr delay |
|---|---|---|
| Phospholipids (mmol/l) | 9.8e-01 | 9.9e-01 |
| Total cholesterol (mmol/l) | 9.6e-01 | 9.7e-01 |
| Cholesterol esters (mmol/l) | 9.4e-01 | 9.8e-01 |
| Free cholesterol (mmol/l) | 9.6e-01 | 9.8e-01 |
| Triglycerides (mmol/l) | 9.9e-01 | 9.9e-01 |
| *Large LDL* | | |
| Particle concentration (mol/l) | 9.8e-01 | 9.9e-01 |
| Total lipids (mmol/l) | 9.7e-01 | 9.9e-01 |
| Phospholipids (mmol/l) | 9.7e-01 | 9.8e-01 |
| Total cholesterol (mmol/l) | 9.7e-01 | 9.9e-01 |
| Cholesterol esters (mmol/l) | 9.7e-01 | 9.9e-01 |
| Free cholesterol (mmol/l) | 9.8e-01 | 9.8e-01 |
| Triglycerides (mmol/l) | 9.9e-01 | 9.9e-01 |
| *Medium LDL* | | |
| Particle concentration (mol/l) | 9.8e-01 | 9.7e-01 |
| Total lipids (mmol/l) | 9.7e-01 | 9.9e-01 |
| Phospholipids (mmol/l) | 9.7e-01 | 9.8e-01 |
| Total cholesterol (mmol/l) | 9.7e-01 | 9.9e-01 |
| Cholesterol esters (mmol/l) | 9.8e-01 | 9.8e-01 |
| Free cholesterol (mmol/l) | 9.8e-01 | 9.8e-01 |
| Triglycerides (mmol/l) | 9.8e-01 | 9.8e-01 |
| *Small LDL* | | |
| Particle concentration (mol/l) | 9.8e-01 | 9.8e-01 |
| Total lipids (mmol/l) | 9.7e-01 | 9.8e-01 |
| Phospholipids (mmol/l) | 9.8e-01 | 9.8e-01 |
| Total cholesterol (mmol/l) | 9.8e-01 | 9.8e-01 |
| Cholesterol esters (mmol/l) | 9.4e-01 | 9.9e-01 |
| Free cholesterol (mmol/l) | 9.8e-01 | 9.8e-01 |
| Triglycerides (mmol/l) | 9.9e-01 | 9.9e-01 |

| Metabolic traits | thaw-buffer delay | buffer-nmr delay |
|---|---|---|
| *Very large HDL* | | |
| Particle concentration (mol/l) | 9.8e-01 | 9.9e-01 |
| Total lipids (mmol/l) | 9.8e-01 | 9.9e-01 |
| Phospholipids (mmol/l) | 9.9e-01 | 1.0e+00 |
| Total cholesterol (mmol/l) | 9.7e-01 | 9.8e-01 |
| Cholesterol esters (mmol/l) | 9.7e-01 | 9.9e-01 |
| Free cholesterol (mmol/l) | 9.8e-01 | 9.9e-01 |
| Triglycerides (mmol/l) | 9.5e-01 | 9.9e-01 |
| *Large HDL* | | |
| Particle concentration (mol/l) | 9.9e-01 | 9.9e-01 |
| Total lipids (mmol/l) | 9.9e-01 | 9.9e-01 |
| Phospholipids (mmol/l) | 9.9e-01 | 9.9e-01 |
| Total cholesterol (mmol/l) | 1.0e+00 | 1.0e+00 |
| Cholesterol esters (mmol/l) | 9.9e-01 | 1.0e+00 |
| Free cholesterol (mmol/l) | 9.9e-01 | 9.9e-01 |
| Triglycerides (mmol/l) | 9.8e-01 | 9.9e-01 |
| *Medium HDL* | | |
| Particle concentration (mol/l) | 8.9e-01 | 9.7e-01 |
| Total lipids (mmol/l) | 8.9e-01 | 9.7e-01 |
| Phospholipids (mmol/l) | 9.3e-01 | 9.7e-01 |
| Total cholesterol (mmol/l) | 9.1e-01 | 9.6e-01 |
| Cholesterol esters (mmol/l) | 9.0e-01 | 9.5e-01 |
| Free cholesterol (mmol/l) | 9.2e-01 | 9.6e-01 |
| Triglycerides (mmol/l) | 9.8e-01 | 9.8e-01 |
| *Small HDL* | | |
| Particle concentration (mol/l) | 8.8e-01 | 9.3e-01 |
| Total lipids (mmol/l) | 8.6e-01 | 9.3e-01 |
| Phospholipids (mmol/l) | 9.8e-01 | 9.8e-01 |
| Total cholesterol (mmol/l) | 7.4e-01 | 7.8e-01 |

| Metabolic traits | thaw-buffer delay | buffer-nmr delay |
|---|---|---|
| Cholesterol esters (mmol/l) | 8.7e-01 | 8.7e-01 |
| Free cholesterol (mmol/l) | 9.7e-01 | 9.8e-01 |
| Triglycerides (mmol/l) | 9.8e-01 | 9.6e-01 |
| **Lipoprotein particle size** | | |
| VLDL particle size (nm) | 9.9e-01 | 9.9e-01 |
| LDL particle size (nm) | 8.2e-01 | 9.3e-01 |
| HDL particle size (nm) | 9.9e-01 | 9.9e-01 |
| **Cholesterol** | | |
| Total cholesterol (mmol/l) | 9.9e-01 | 9.9e-01 |
| VLDL cholesterol (mmol/l) | 9.4e-01 | 9.9e-01 |
| Remnant cholesterol (mmol/l) | 9.5e-01 | 9.9e-01 |
| LDL cholesterol (mmol/l) | 9.7e-01 | 9.9e-01 |
| HDL cholesterol (mmol/l) | 9.9e-01 | 9.9e-01 |
| HDL2 cholesterol (mmol/l) | 9.9e-01 | 9.8e-01 |
| HDL3 cholesterol (mmol/l) | 9.8e-01 | 9.8e-01 |
| Esterified cholesterol (mmol/l) | 9.8e-01 | 9.9e-01 |
| Free cholesterol (mmol/l) | 9.7e-01 | 9.7e-01 |
| **Glycerides and phospholipids** | | |
| Triglycerides (mmol/l) | 9.9e-01 | 1.0e+00 |
| VLDL triglycerides (mmol/l) | 9.9e-01 | 9.9e-01 |
| LDL triglycerides (mmol/l) | 1.0e+00 | 9.9e-01 |
| HDL triglycerides (mmol/l) | 9.9e-01 | 9.9e-01 |
| Diacylglycerol (mmol/l) | 4.1e-01 | 1.9e-01 |
| Phosphoglycerides (mmol/l) | 9.2e-01 | 9.2e-01 |
| Phosphatidylcholine + other cholines (mmol/l) | 9.2e-01 | 9.3e-01 |
| Sphingomyelins (mmol/l) | 8.6e-01 | 8.6e-01 |
| Cholines (mmol/l) | 8.8e-01 | 8.7e-01 |
| **Apolipoproteins** | | |
| Apolipoprotein A-I (g/l) | 9.8e-01 | 9.8e-01 |

| Metabolic traits | thaw-buffer delay | buffer-nmr delay |
|---|---|---|
| Apolipoprotein B (g/l) | 9.4e-01 | 9.9e-01 |
| **Fatty acids** | | |
| Total fatty acids (mmol/l) | 9.4e-01 | 9.8e-01 |
| Fatty acid chain length | 9.3e-01 | 9.0e-01 |
| Degree of unsaturation | 8.8e-01 | 9.6e-01 |
| Docosahexaenoic acid (mmol/l) | 9.7e-01 | 9.8e-01 |
| Linoleic acid (mmol/l) | 9.6e-01 | 9.6e-01 |
| Conjugated linoleic acid (mmol/l) | 5.8e-01 | 7.9e-01 |
| n-3 fatty acids (mmol/l) | 9.6e-01 | 9.8e-01 |
| n-6 fatty acids (mmol/l) | 9.6e-01 | 9.6e-01 |
| PUFA (mmol/l) | 9.7e-01 | 9.6e-01 |
| MUFA (mmol/l) | 1.0e+00 | 9.9e-01 |
| Saturated fatty acids (mmol/l) | 9.3e-01 | 9.8e-01 |
| **Glycolysis related metabolites** | | |
| Glucose (mmol/l) | 9.7e-01 | 9.8e-01 |
| Lactate (mmol/l) | 9.8e-01 | 9.9e-01 |
| Pyruvate (mmol/l) | 9.8e-01 | 9.9e-01 |
| Citrate (mmol/l) | 7.7e-01 | 8.9e-01 |
| Glycerol (mmol/l) | 8.9e-01 | 9.2e-01 |
| **Amino acids** | | |
| Alanine (mmol/l) | 9.7e-01 | 9.8e-01 |
| Glutamine (mmol/l) | 9.6e-01 | 9.5e-01 |
| Histidine (mmol/l) | 4.8e-01 | 7.7e-01 |
| Glycine (mmol/l) | 9.7e-01 | 9.6e-01 |
| *Branched-chain amino acids* | | |
| Isoleucine (mmol/l) | 9.6e-01 | 9.6e-01 |
| Leucine (mmol/l) | 9.8e-01 | 9.9e-01 |
| Valine (mmol/l) | 9.9e-01 | 9.9e-01 |
| *Aromatic amino acids* | | |

| Metabolic traits | thaw-buffer delay | buffer-nmr delay |
|---|---|---|
| Phenylalanine (mmol/l) | 7.7e-01 | 9.0e-01 |
| Tyrosine (mmol/l) | 9.6e-01 | 9.5e-01 |
| **Ketone bodies** | | |
| Acetate (mmol/l) | 9.5e-01 | 9.7e-01 |
| Beta-hydroxybutyrate (mmol/l) | 9.6e-01 | 9.7e-01 |
| **Fluid balance** | | |
| Creatinine (mmol/l) | 9.2e-01 | 9.2e-01 |
| Albumin (signal area) | 9.2e-01 | 9.5e-01 |
| **Inflammation** | | |
| Glycoprotein acetyls (mmol/l) | 9.8e-01 | 9.8e-01 |

*sTable 9. Spearman correlation: EDTA-plasma, post-storage handling effects. Spearman rank correlation coefficients between metabolic concentrations (or values) in reference conditions samples (i.e. no sample preparation or NMR analysis delays) and samples (i) left for 24h before addition of sodium buffer followed by immediate NMR analysis (i.e. thaw to buffer addition delay); (ii) thawed overnight, addition of sodium buffer, then left for 24h before NMR analysis (buffer addition to NMR profiling delay). Pyruvate, glycerol and glycine are not quantified in EDTA -plasma samples due to the interfering resonances of EDTA on their signals.*

| Metabolic traits | thaw-buffer delay | buffer-nmr delay |
|---|---|---|
| **Lipoprotein subclasses** | | |
| *Extremely large VLDL* | | |
| Particle concentration (mol/l) | 9.6e-01 | 9.1e-01 |
| Total lipids (mmol/l) | 9.6e-01 | 9.2e-01 |
| Phospholipids (mmol/l) | 9.6e-01 | 9.3e-01 |
| Total cholesterol (mmol/l) | 9.6e-01 | 9.4e-01 |
| Cholesterol esters (mmol/l) | 9.6e-01 | 9.3e-01 |
| Free cholesterol (mmol/l) | 9.7e-01 | 9.4e-01 |
| Triglycerides (mmol/l) | 9.6e-01 | 9.1e-01 |
| *Very large VLDL* | | |
| Particle concentration (mol/l) | 9.8e-01 | 9.3e-01 |
| Total lipids (mmol/l) | 9.8e-01 | 9.4e-01 |
| Phospholipids (mmol/l) | 9.8e-01 | 9.4e-01 |
| Total cholesterol (mmol/l) | 9.7e-01 | 9.6e-01 |
| Cholesterol esters (mmol/l) | 9.8e-01 | 9.6e-01 |
| Free cholesterol (mmol/l) | 9.8e-01 | 9.6e-01 |
| Triglycerides (mmol/l) | 9.8e-01 | 9.3e-01 |
| *Large VLDL* | | |
| Particle concentration (mol/l) | 9.7e-01 | 9.8e-01 |
| Total lipids (mmol/l) | 9.7e-01 | 9.8e-01 |
| Phospholipids (mmol/l) | 9.8e-01 | 9.8e-01 |
| Total cholesterol (mmol/l) | 9.7e-01 | 9.7e-01 |

| Metabolic traits | thaw-buffer delay | buffer-nmr delay |
|---|---|---|
| Cholesterol esters (mmol/l) | 9.5e-01 | 9.4e-01 |
| Free cholesterol (mmol/l) | 9.7e-01 | 9.7e-01 |
| Triglycerides (mmol/l) | 9.8e-01 | 9.9e-01 |
| *Medium VLDL* | | |
| Particle concentration (mol/l) | 9.7e-01 | 9.8e-01 |
| Total lipids (mmol/l) | 9.7e-01 | 9.8e-01 |
| Phospholipids (mmol/l) | 9.6e-01 | 9.8e-01 |
| Total cholesterol (mmol/l) | 9.4e-01 | 9.7e-01 |
| Cholesterol esters (mmol/l) | 9.7e-01 | 9.8e-01 |
| Free cholesterol (mmol/l) | 9.8e-01 | 9.8e-01 |
| Triglycerides (mmol/l) | 9.9e-01 | 9.9e-01 |
| *Small VLDL* | | |
| Particle concentration (mol/l) | 9.9e-01 | 9.8e-01 |
| Total lipids (mmol/l) | 9.8e-01 | 9.8e-01 |
| Phospholipids (mmol/l) | 9.8e-01 | 9.8e-01 |
| Total cholesterol (mmol/l) | 9.8e-01 | 9.7e-01 |
| Cholesterol esters (mmol/l) | 9.6e-01 | 9.6e-01 |
| Free cholesterol (mmol/l) | 9.8e-01 | 9.7e-01 |
| Triglycerides (mmol/l) | 9.9e-01 | 9.9e-01 |
| *Very Small VLDL* | | |
| Particle concentration (mol/l) | 9.6e-01 | 9.6e-01 |
| Total lipids (mmol/l) | 9.6e-01 | 9.5e-01 |
| Phospholipids (mmol/l) | 9.5e-01 | 9.8e-01 |
| Total cholesterol (mmol/l) | 9.3e-01 | 9.1e-01 |
| Cholesterol esters (mmol/l) | 9.1e-01 | 8.9e-01 |
| Free cholesterol (mmol/l) | 9.6e-01 | 9.7e-01 |
| Triglycerides (mmol/l) | 9.7e-01 | 9.7e-01 |
| *IDL* | | |
| Particle concentration (mol/l) | 9.8e-01 | 9.8e-01 |

| Metabolic traits | thaw-buffer delay | buffer-nmr delay |
|---|---|---|
| Total lipids (mmol/l) | 9.8e-01 | 9.7e-01 |
| Phospholipids (mmol/l) | 9.8e-01 | 9.7e-01 |
| Total cholesterol (mmol/l) | 9.7e-01 | 9.6e-01 |
| Cholesterol esters (mmol/l) | 9.7e-01 | 9.4e-01 |
| Free cholesterol (mmol/l) | 9.7e-01 | 9.7e-01 |
| Triglycerides (mmol/l) | 9.8e-01 | 9.8e-01 |
| *Large LDL* | | |
| Particle concentration (mol/l) | 9.9e-01 | 9.8e-01 |
| Total lipids (mmol/l) | 9.8e-01 | 9.8e-01 |
| Phospholipids (mmol/l) | 9.8e-01 | 9.8e-01 |
| Total cholesterol (mmol/l) | 9.9e-01 | 9.7e-01 |
| Cholesterol esters (mmol/l) | 9.7e-01 | 9.7e-01 |
| Free cholesterol (mmol/l) | 9.8e-01 | 9.6e-01 |
| Triglycerides (mmol/l) | 9.7e-01 | 9.7e-01 |
| *Medium LDL* | | |
| Particle concentration (mol/l) | 9.8e-01 | 9.8e-01 |
| Total lipids (mmol/l) | 9.8e-01 | 9.8e-01 |
| Phospholipids (mmol/l) | 9.9e-01 | 9.9e-01 |
| Total cholesterol (mmol/l) | 9.8e-01 | 9.7e-01 |
| Cholesterol esters (mmol/l) | 9.8e-01 | 9.8e-01 |
| Free cholesterol (mmol/l) | 9.9e-01 | 9.7e-01 |
| Triglycerides (mmol/l) | 9.4e-01 | 9.5e-01 |
| *Small LDL* | | |
| Particle concentration (mol/l) | 9.8e-01 | 9.8e-01 |
| Total lipids (mmol/l) | 9.7e-01 | 9.8e-01 |
| Phospholipids (mmol/l) | 9.9e-01 | 9.9e-01 |
| Total cholesterol (mmol/l) | 9.8e-01 | 9.7e-01 |
| Cholesterol esters (mmol/l) | 9.6e-01 | 9.7e-01 |
| Free cholesterol (mmol/l) | 9.7e-01 | 9.9e-01 |

| Metabolic traits | thaw-buffer delay | buffer-nmr delay |
|---|---|---|
| Triglycerides (mmol/l) | 9.8e-01 | 9.8e-01 |
| *Very large HDL* | | |
| Particle concentration (mol/l) | 9.9e-01 | 9.9e-01 |
| Total lipids (mmol/l) | 9.9e-01 | 9.9e-01 |
| Phospholipids (mmol/l) | 9.9e-01 | 9.9e-01 |
| Total cholesterol (mmol/l) | 9.8e-01 | 9.7e-01 |
| Cholesterol esters (mmol/l) | 9.8e-01 | 9.8e-01 |
| Free cholesterol (mmol/l) | 9.9e-01 | 9.9e-01 |
| Triglycerides (mmol/l) | 9.5e-01 | 9.4e-01 |
| *Large HDL* | | |
| Particle concentration (mol/l) | 9.9e-01 | 9.9e-01 |
| Total lipids (mmol/l) | 9.9e-01 | 9.9e-01 |
| Phospholipids (mmol/l) | 9.9e-01 | 9.9e-01 |
| Total cholesterol (mmol/l) | 9.9e-01 | 9.9e-01 |
| Cholesterol esters (mmol/l) | 1.0e+00 | 1.0e+00 |
| Free cholesterol (mmol/l) | 9.9e-01 | 9.9e-01 |
| Triglycerides (mmol/l) | 8.8e-01 | 8.8e-01 |
| *Medium HDL* | | |
| Particle concentration (mol/l) | 8.6e-01 | 8.8e-01 |
| Total lipids (mmol/l) | 8.6e-01 | 8.6e-01 |
| Phospholipids (mmol/l) | 9.0e-01 | 9.0e-01 |
| Total cholesterol (mmol/l) | 8.8e-01 | 8.5e-01 |
| Cholesterol esters (mmol/l) | 8.9e-01 | 8.4e-01 |
| Free cholesterol (mmol/l) | 8.6e-01 | 8.6e-01 |
| Triglycerides (mmol/l) | 9.5e-01 | 9.6e-01 |
| *Small HDL* | | |
| Particle concentration (mol/l) | 8.2e-01 | 8.3e-01 |
| Total lipids (mmol/l) | 8.5e-01 | 8.5e-01 |
| Phospholipids (mmol/l) | 9.8e-01 | 9.8e-01 |

| Metabolic traits | thaw-buffer delay | buffer-nmr delay |
|---|---|---|
| Total cholesterol (mmol/l) | 7.3e-01 | 7.0e-01 |
| Cholesterol esters (mmol/l) | 8.3e-01 | 7.3e-01 |
| Free cholesterol (mmol/l) | 9.4e-01 | 9.5e-01 |
| Triglycerides (mmol/l) | 9.7e-01 | 9.6e-01 |
| **Lipoprotein particle size** | | |
| VLDL particle size (nm) | 9.9e-01 | 9.7e-01 |
| LDL particle size (nm) | 7.8e-01 | 7.8e-01 |
| HDL particle size (nm) | 9.9e-01 | 9.8e-01 |
| **Cholesterol** | | |
| Total cholesterol (mmol/l) | 9.8e-01 | 9.8e-01 |
| VLDL cholesterol (mmol/l) | 9.5e-01 | 9.5e-01 |
| Remnant cholesterol (mmol/l) | 9.7e-01 | 9.6e-01 |
| LDL cholesterol (mmol/l) | 9.9e-01 | 9.6e-01 |
| HDL cholesterol (mmol/l) | 9.8e-01 | 9.7e-01 |
| HDL2 cholesterol (mmol/l) | 9.7e-01 | 9.7e-01 |
| HDL3 cholesterol (mmol/l) | 9.5e-01 | 9.5e-01 |
| Esterified cholesterol (mmol/l) | 9.8e-01 | 9.9e-01 |
| Free cholesterol (mmol/l) | 9.7e-01 | 9.6e-01 |
| **Glycerides and phospholipids** | | |
| Triglycerides (mmol/l) | 9.9e-01 | 9.9e-01 |
| VLDL triglycerides (mmol/l) | 9.8e-01 | 9.8e-01 |
| LDL triglycerides (mmol/l) | 9.7e-01 | 9.7e-01 |
| HDL triglycerides (mmol/l) | 9.9e-01 | 9.8e-01 |
| Diacylglycerol (mmol/l) | 7.2e-01 | 7.7e-01 |
| Phosphoglycerides (mmol/l) | 9.3e-01 | 9.4e-01 |
| Phosphatidylcholine + other cholines (mmol/l) | 9.1e-01 | 9.2e-01 |
| Sphingomyelins (mmol/l) | 7.7e-01 | 8.1e-01 |
| Cholines (mmol/l) | 8.3e-01 | 8.6e-01 |
| **Apolipoproteins** | | |

| Metabolic traits | thaw-buffer delay | buffer-nmr delay |
|---|---|---|
| Apolipoprotein A-I (g/l) | 9.7e-01 | 9.8e-01 |
| Apolipoprotein B (g/l) | 9.7e-01 | 9.5e-01 |
| **Fatty acids** | | |
| Total fatty acids (mmol/l) | 9.6e-01 | 9.7e-01 |
| Fatty acid chain length | 8.8e-01 | 5.3e-01 |
| Degree of unsaturation | 9.4e-01 | 8.3e-01 |
| Docosahexaenoic acid (mmol/l) | 9.8e-01 | 9.8e-01 |
| Linoleic acid (mmol/l) | 9.8e-01 | 9.5e-01 |
| Conjugated linoleic acid (mmol/l) | 7.3e-01 | 7.8e-01 |
| n-3 fatty acids (mmol/l) | 9.7e-01 | 9.8e-01 |
| n-6 fatty acids (mmol/l) | 9.6e-01 | 9.6e-01 |
| PUFA (mmol/l) | 9.7e-01 | 9.7e-01 |
| MUFA (mmol/l) | 9.8e-01 | 9.6e-01 |
| Saturated fatty acids (mmol/l) | 9.8e-01 | 9.7e-01 |
| **Glycolysis related metabolites** | | |
| Glucose (mmol/l) | 9.7e-01 | 9.7e-01 |
| Lactate (mmol/l) | 9.9e-01 | 9.9e-01 |
| Pyruvate (mmol/l) | NA | NA |
| Citrate (mmol/l) | 8.8e-01 | 8.5e-01 |
| Glycerol (mmol/l) | NA | NA |
| **Amino acids** | | |
| Alanine (mmol/l) | 9.7e-01 | 9.6e-01 |
| Glutamine (mmol/l) | 8.8e-01 | 9.6e-01 |
| Histidine (mmol/l) | 3.6e-01 | 5.7e-01 |
| Glycine (mmol/l) | NA | NA |
| *Branched-chain amino acids* | | |
| Isoleucine (mmol/l) | 9.0e-01 | 9.2e-01 |
| Leucine (mmol/l) | 9.9e-01 | 9.8e-01 |
| Valine (mmol/l) | 9.8e-01 | 9.9e-01 |

| Metabolic traits | thaw-buffer delay | buffer-nmr delay |
|---|---|---|
| *Aromatic amino acids* | | |
| Phenylalanine (mmol/l) | 7.5e-01 | 7.4e-01 |
| Tyrosine (mmol/l) | 9.1e-01 | 9.6e-01 |
| **Ketone bodies** | | |
| Acetate (mmol/l) | 8.3e-01 | 9.0e-01 |
| Beta-hydroxybutyrate (mmol/l) | 9.8e-01 | 9.1e-01 |
| **Fluid balance** | | |
| Creatinine (mmol/l) | 9.7e-01 | 9.3e-01 |
| Albumin (signal area) | 9.0e-01 | 8.9e-01 |
| **Inflammation** | | |
| Glycoprotein acetyls (mmol/l) | 9.7e-01 | 9.9e-01 |

*sTable 10. Literature table: summary of previous studies assessing the effects of pre and post-storage handling conditions on serum or plasma metabolic traits also measured by Nightingale® NMR platform.*

*__Note 1__: **Untargeted** metabolomics (U), aims to measure as many metabolites as possible in a biological sample, without any requirement that the metabolites be annotated beforehand. Quantification is given as relative concentrations. **Targeted** metabolomics (T), provides absolute concentration of a selected panel of annotated metabolites. **Semi-targeted** metabolomics (S) provides absolute concentration using one calibration model to quantify multiple metabolites, instead of one quantification model per metabolite. Quantification is therefore less accurate than in targeted metabolomics.*
*__Note 2__: Studies using NMR have very similar metabolite coverage as the NMR platform used here. Untargeted studies due to its nature, i.e. the type of data (e.g. full spectra) and the statistical methods used (e.g. partial least squares- discriminate analysis), are more biased towards reporting non-robust metabolites. Therefore, for all untargeted NMR studies cited in sTable 6, it is assumed that non-reported metabolites are stable. For example, Bernini et al. 2011[9], uses an untargeted NMR platform, that like the one used in this paper will detect isoleucine, since the authors did not report a change in this amino-acid, it is assumed that it is stable under the conditions they tested. However, in the table below we did not list all metabolic traits detected by the platform used in our study alongside the references, that use NMR untargeted platform, which report stability of a metabolite by omission.*
*__Abbreviations__: **CPMG**=Carr-Purcell-Meiboom-Gill; **$^{13}$C**= Carbon-13; **EDTA**=Ethylenediaminetetraacetic acid; **ESI**= Electrospray ionization; **FIA**= Flow injection analysis; **GC**= Gas Chromatography; **($^1$H) NMR**= Proton nuclear magnetic resonance; **HDL**= High density lipoprotein; **HSQC**=Heteronuclear Single Quantum Coherence; **J-res**=J-resolved; **LC**=Liquid Chromatography; **LDL**= Low density lipoprotein; **MS**= Mass Spectrometry; **N**= number of individuals; **NMR**= Nuclear Magnetic Resonance; **NOESY**=Nuclear Overhauser Effect Spectroscopy; **RT**=Room Temperature; **STOCSY**= Statistical Total Correlation Spectroscopy; **SST**= Serum-Separating Tube; **1D**=one dimension.*
**__Details of the metabolomics platforms__: **Biocrates®**= Biocrates® Life Sciences AG (Innsbruck, Austria), commercial suppliers of the AbsoluteIQD™ commercial kits; **ICL NPC**= Imperial College London National Phenome Centre, laboratory of Jeremy Nicolson, John Lindon and colleagues; **Nightingale Health®**=commercial laboratory formerly known as Brainshake Inc. The same platform is also used by the Mika Ala-Korpela lab at the Biocentre Oulu platform (Finland).*
**‡ studies testing the effects of pre-analytical variation on a different pre-analytical *phase* e.g. studies testing *post*-centrifugation conditions during pre-storage (i.e. long-term freezing) sample handling, whilst we tested *pre*-centrifugation conditions during pre-storage handling.**

| **Metabolic trait** | **Reference** | **Sample handling phase: Pre/post-storage (i.e. long-term freezing); Pre/post-centrifugation;** | **Sample type** | **Incubation temperature** | **Incubation duration** | **Analytical Platform* [Targeted (T), Untargeted (U), Semi-targeted (S)]** | ***N*** | **Message** |
|---|---|---|---|---|---|---|---|---|
| **Cholesterol** | | | | | | | | |
| **Total cholesterol** | Key et al. 1996 [5] | Pre-storage: pre-centrifugation; | Serum, sodium citrate-plasma; Fasting status not specified; Serum reference: 20ºC, 2h Plasma reference: 20ºC (20 min) and after 4ºC, 2h | 4ºC | 2, 6, 24h | Clinical Chemistry / (T) | 28 | **Stable** (decrease in mean percentage change up to 3%) |

| | | | | | | | | |
|---|---|---|---|---|---|---|---|---|
| | Clark et al. 2003[6] | Pre-storage: pre-centrifugation; | Potassium EDTA-plasma; Non-fasting; Reference: centrifuged immediately | 4°C, 21°C | 0, 1-4, 7 days | Clinical Chemistry / (T) | 12 | **Stable** (mean percentage change less than 0.5% per day) |
| | Boyanton et al. 2002[7] | Pre-storage: pre and ǂpost-centrifugation; | Serum, lithium heparin-plasma; Non-fasting; Reference: 0.5h, 25°C | 25°C | 0.5, 4, 8, 16, 24, 32, 40, 48, 56h | Clinical Chemistry / (T) | 10 | Pre-storage/pre-centrifugation: **increase**, more pronounced in plasma than serum. Pre-storage/ǂpost-centrifugation: **stable**. |
| | Oddoze et al. 2012[8] | Pre-storage: pre-centrifugation; | Serum, lithium heparin and fluoride plasma; Fasting status not specified; Serum reference: 0.5h; Plasma reference: centrifuged immediately | 4°C, 25°C | 0, 2, 4, 6, 24h | Clinical Chemistry / (T) | 10 | **Stable**. |
| | *Current report* | *Pre-storage: pre-centrifugation; Post-storage: post-centrifugation;* | *Serum, potassium EDTA-plasma; Non-fasting; Reference, pre-storage: 1.5h, 4°C; Reference, post-storage: no sample or NMR analysis delay* | *Pre-storage/ pre-centrifugation: 4°C, 21°C Post-storage/ post-centrifugation: 4°C* | *Pre-storage/ pre-centrifugation: 1.5, 24, 48h Post-storage/ post-centrifugation: 0, 24h* | *($^1$H) NMR, Nightingale Health* / (T)* | *Pre: 23 Post:25* | *Pre-storage/pre-centrifugation: **stable** at 4°C; mean increase up to 0.2SD at 21°C Post-storage/post-centrifugation: **stable**.* |
| **LDL cholesterol** | Key et al. 1996[5] | Pre-storage: pre-centrifugation; | Serum, sodium citrate-plasma; Fasting status not specified; Serum reference: 20°C, 2h Plasma reference: 20°C (20 min) and after 4°C, 2h | 4°C | 2, 6, 24h | Clinical Chemistry / (T) | 28 | **Stable** (mean percentage change up to 3.4%) |
| | Clark et al. 2003 [6] | Pre-storage: pre-centrifugation; | Potassium EDTA-plasma; Non-fasting; Reference: centrifuged immediately | 4°C, 21°C | 0, 1-4, 7 days | Clinical Chemistry / (T) | 12 | **Stable** (mean percentage change less than 1% per day) |
| | Oddoze et al. 2012[8] | Pre-storage: pre-centrifugation; | Serum, lithium heparin and fluoride plasma; Fasting status not specified; Serum reference: 0.5h; Plasma reference: centrifuged immediately | 4°C, 25°C | 0, 2, 4, 6, 24h | Clinical Chemistry / (T) | 10 | **Stable**. |
| | *Current report* | *Pre-storage: pre-centrifugation; Post-storage: post-centrifugation;* | *Serum, potassium EDTA-plasma; Non-fasting; Reference, pre-storage: 1.5h, 4°C;* | *Pre-storage/ pre-centrifugation: 4°C, 21°C* | *Pre-storage/ pre-centrifugation: 1.5, 24, 48h Post-storage/ post-* | *($^1$H) NMR, Nightingale Health* / (T)* | *Pre: 23 Post:25* | *Pre-storage/pre-centrifugation: **stable** at 4°C; mean increase up to 0.3SD at 21°C* |

| | | | | | | | | |
|---|---|---|---|---|---|---|---|---|
| | | | *Reference, post-storage: no sample or NMR analysis delay* | *Post-storage/ post-centrifugation: 4°C* | *centrifugation: 0, 24h* | | | *Post-storage/post-centrifugation: **stable**.* |
| **HDL cholesterol** | Key et al. 1996[5] | Pre-storage: pre-centrifugation; | Serum, sodium citrate-plasma; Fasting status not specified; Serum reference: 20°C, 2h Plasma reference: 20°C (20 min) and after 4°C, 2h | 4°C | 2, 6, 24h | Clinical Chemistry / (T) | 28 | **Stable** (decrease in mean percentage change up to 4.4%) |
| | Clark et al. 2003[6] | Pre-storage: pre-centrifugation; | Potassium EDTA-plasma; Non-fasting; Reference: centrifuged immediately | 4°C, 21°C | 0, 1-4, 7 days | Clinical Chemistry / (T) | 12 | **Stable** (mean percentage change less than 1% per day) |
| | Oddoze et al. 2012[8] | Pre-storage: pre-centrifugation; | Serum, lithium heparin and fluoride plasma; Fasting status not specified; Serum reference: 0.5h; Plasma reference: centrifuged immediately | 4°C, 25°C | 0, 2, 4, 6, 24h | Clinical Chemistry / (T) | 10 | **Stable**. |
| | *Current report* | *Pre-storage: pre-centrifugation; Post-storage: post-centrifugation;* | *Serum, potassium EDTA-plasma; Non-fasting; Reference, pre-storage: 1.5h, 4°C; Reference, post-storage: no sample or NMR analysis delay* | *Pre-storage/ pre-centrifugation: 4°C, 21°C Post-storage/ post-centrifugation: 4°C* | *Pre-storage/ pre-centrifugation: 1.5, 24, 48h Post-storage/ post-centrifugation: 0, 24h* | *$(^1H)$ NMR, Nightingale Health* / (T)* | *Pre: 23 Post:25* | *Pre-storage/pre-centrifugation: **stable**; Post-storage/post-centrifugation: **stable**.* |
| **Glycerides and phospholipids** | | | | | | | | |
| **Triglycerides** | Key et al. 1996[5] | Pre-storage: pre-centrifugation; | Serum, sodium citrate-plasma; Fasting status not specified; Serum reference: 20°C, 2h Plasma reference: 20°C (20 min) and after 4°C, 2h | 4°C | 2, 6, 24h | Clinical Chemistry / (T) | 28 | **Stable** (increase in mean percentage change up to 5.9% but spearman rank correlation between reference and variant conditions is equal or above 0.95) |
| | Boyanton et al. 2002[7] | Pre-storage: pre and ǂpost-centrifugation; | Serum, lithium heparin-plasma; Non-fasting; Reference: 0.5, 25°C | 25°C | 0.5, 4, 8, 16, 24, 32, 40, 48, 56h | Clinical Chemistry / (T) | 10 | Pre-storage: pre and ǂpost-centrifugation: **stable**. |
| | Clark et al. 2003[6] | Pre-storage: pre-centrifugation; | Potassium EDTA-plasma; Non-fasting; | 4°C, 21°C | 0, 1-4, 7 days | Clinical Chemistry / (T) | 12 | **Stable** (mean percentage change |

| | | | | | | | | |
|---|---|---|---|---|---|---|---|---|
| | | | Reference: centrifuged immediately | | | | | less than 0.5% per day) |
| | Oddoze et al. 2012[8] | Pre-storage: pre-centrifugation; | Serum, lithium heparin and fluoride plasma; Fasting status not specified; Serum reference: 0.5h; Plasma reference: centrifuged immediately | 4ºC, 25ºC | 0, 2, 4, 6, 24h | Clinical Chemistry / (T) | 10 | **Stable**. |
| | Bernini et al. 2011[9] | Pre-storage: pre and ǂpost-centrifugation; | Serum (SST), potassium EDTA-plasma; Fasting status not specified; Reference: centrifuged immediately (for serum after 30 min) and frozen immediately | Pre-centrifugation: 4ºC, 25ºC Post-centrifugation: 25ºC | Pre-centrifugation: 0-4h Post-centrifugation: 0,6,12, 24h | ($^1$H) NMR (1D, NOESY CPMG) / (U) | Pre:6 Post:5 | Pre-storage/pre-centrifugation: **stable;** Pre-storage/ǂpost-centrifugation: **decrease**. |
| | *Current report* | *Pre-storage: pre-centrifugation; Post-storage: post-centrifugation;* | *Serum, potassium EDTA-plasma; Non-fasting; Reference, pre-storage: 1.5h, 4ºC; Reference, post-storage: no sample or NMR analysis delay* | *Pre-storage/ pre-centrifugation: 4ºC, 21ºC Post-storage/ post-centrifugation: 4ºC* | *Pre-storage/ pre-centrifugation: 1.5, 24, 48h Post-storage/ post-centrifugation: 0, 24h* | *($^1$H) NMR, Nightingale Health* / (T)* | *Pre: 23 Post:25* | *Pre-storage/pre-centrifugation: **stable**; Post-storage/post-centrifugation: **stable**.* |
| **Phosphatidylcholine** | ǂAnton et al. 2015[10] | Pre-storage: ǂpost-centrifugation; | Serum; Fasting; Reference:max 5h, on ice | Dry ice, wet ice, RT (22-24ºC) | 0,12, 24, 36h | FIA-ESI-MS/MS, Biocrates*/ (S) | 19 (males) | **Decrease** (at RT i.e. 22-24ºC) |
| | ǂPinto et al 2014[11] | Pre-storage: ǂpost-centrifugation; | Heparin or EDTA-plasma (not clear); Fasting status not specified; | RT | 1-21h | ($^1$H) NMR (1D, NOESY, CPMG, STOCSY) / (U) | 3 (pregnant) | **Increase** |
| | *Current report* | *Pre-storage: pre-centrifugation; Post-storage: post-centrifugation;* | *Serum, potassium EDTA-plasma; Non-fasting; Reference, pre-storage: 1.5h, 4ºC; Reference, post-storage: no sample or NMR analysis delay* | *Pre-storage/ pre-centrifugation: 4ºC, 21ºC Post-storage/ post-centrifugation: 4ºC* | *Pre-storage/ pre-centrifugation: 1.5, 24, 48h Post-storage/ post-centrifugation: 0, 24h* | *($^1$H) NMR, Nightingale Health* / (T)* | *Pre: 20 Post:21* | *Pre-storage/pre-centrifugation: **stable** at 4ºC; mean decrease up to 0.2SD at 21ºC. Post-storage/post-centrifugation: mean decrease up to 0.3SD.* |
| **Sphingomyelins** | ǂAnton et al. 2015[10] | Pre-storage: ǂpost-centrifugation; | Serum; Fasting; Reference: max 5h, on ice | Dry ice, wet ice, RT (22-24ºC) | 0,12, 24, 36h | FIA-ESI-MS/MS, Biocrates* / (S) | 19 (males) | **Stable** |
| | ǂPinto et al 2014[11] | Pre-storage: ǂpost-centrifugation; | Heparin or EDTA-plasma (not clear); Fasting status not specified; | RT | 1-21h | ($^1$H) NMR (1D, NOESY, CPMG, STOCSY) / (U) | 3 (pregnant) | **Increase** |

| | | | | | | | | |
|---|---|---|---|---|---|---|---|---|
| | Breier et al. 2014[12] | Pre-storage: pre-centrifugation; | Serum, potassium EDTA-plasma; Fasting; Reference, EDTA-plasma: centrifuged immediately; Reference, serum: 0.5h, 21°C. | Serum and EDTA-plasma: 4°C; EDTA-plasma: 21°C | Serum and EDTA-plasma: 0, 3, 6, 24h EDTA-plasma: 24h | ESI-LC-MS/MS, MS/MS, Biocrates*/ (S) | 22 | **Stable** |
| | *Current report* | *Pre-storage: pre-centrifugation; Post-storage: post-centrifugation;* | *Serum, potassium EDTA-plasma; Non-fasting; Reference, pre-storage: 1.5h, 4°C; Reference, post-storage: no sample or NMR analysis delay* | *Pre-storage/ pre-centrifugation: 4°C, 21°C Post-storage/ post-centrifugation: 4°C* | *Pre-storage/ pre-centrifugation: 1.5, 24, 48h Post-storage/ post-centrifugation: 0, 24h* | *($^1$H) NMR, Nightingale Health* / (T)* | *Pre: 20 Post:21* | *Pre-storage/pre-centrifugation: **stable**. Post-storage/post-centrifugation: mean changes up to 0.3SD.* |
| **Cholines** | Jobard et al. 2016[13] | Pre-storage: pre and ǂpost-centrifugation; | Serum, heparin-plasma; Fasting; Reference (pre-centrifugation): 1h,22°C; Reference (post-centrifugation): 15min | Pre-centrifugation: 4°C, 22°C Post-centrifugation: 22°C | Pre-centrifugation: 1h (4°C), 6h (4°C, 22°C); Post-centrifugation: 15min, 1h | ($^1$H) NMR (1D, CPMG, NOESY, ($^1$H-$^{13}$C) HSQC, STOCSY, J-res) / (U) | 96 | Pre-storage/pre-centrifugation: **Increase** at 22°C, 6h; **stable** at 4°C. Pre-storage/ǂpost-centrifugation: **stable** |
| | ǂPinto et al 2014[11] | Pre-storage: ǂpost-centrifugation; | Heparin or EDTA-plasma (not clear); Fasting status not specified; | RT | 1-21h | ($^1$H) NMR (1D, NOESY, CPMG, STOCSY) / (U) | 3 (pregnant) | **Increase** |
| | Bernini et al. 2011[9] | Pre-storage: pre and ǂpost-centrifugation; | Serum (SST), potassium EDTA-plasma; Fasting status not specified; Reference: centrifuged immediately (for serum after 30 min) and frozen immediately | Pre-centrifugation: 4°C, 25°C Post-centrifugation: 25°C | Pre-centrifugation: 0-4h Post-centrifugation: 0,6,12, 24h | ($^1$H) NMR (1D, NOESY CPMG) / (U) | Pre: 6 Post: 5 | Pre-storage/pre-centrifugation: **stable;** Pre-storage/ǂpost-centrifugation: **decrease**. |
| | *Current report* | *Pre-storage: pre-centrifugation; Post-storage: post-centrifugation;* | *Serum, potassium EDTA-plasma; Non-fasting; Reference, pre-storage: 1.5h, 4°C; Reference, post-storage: no sample or NMR analysis delay* | *Pre-storage/ pre-centrifugation: 4°C, 21°C Post-storage/ post-centrifugation: 4°C* | *Pre-storage/ pre-centrifugation: 1.5, 24, 48h Post-storage/ post-centrifugation: 0, 24h* | *($^1$H) NMR, Nightingale Health* / (T)* | *Pre: 20 Post:21* | *Pre-storage/pre-centrifugation: **stable**. Post-storage/post-centrifugation: **stable**.* |

| Apolipoproteins | | | | | | | | |
|---|---|---|---|---|---|---|---|---|
| **Apolipoprotein A-I** | Clark et al. 2003[6] | Pre-storage: pre-centrifugation; | Potassium EDTA-plasma; Non-fasting; Reference: centrifuged immediately | 4°C, 21°C | 0,1-4, 7 days | Clinical Chemistry / (T) | 12 | **Stable** (mean percentage change less than 0.5% per day) |
| | Oddoze et al. 2012[8] | Pre-storage: pre-centrifugation; | Serum (SST), lithium heparin and fluoride plasma; Fasting status not specified; Serum reference: 0.5h; Plasma reference: centrifuged immediately | 4°C, 25°C | 0, 2, 4, 6, 24h | Clinical Chemistry / (T) | 10 | **Stable**. |
| | *Current report* | *Pre-storage: pre-centrifugation; Post-storage: post-centrifugation;* | *Serum, potassium EDTA-plasma; Non-fasting; Reference, pre-storage: 1.5h, 4°C; Reference, post-storage: no sample or NMR analysis delay* | *Pre-storage/ pre-centrifugation: 4°C, 21°C Post-storage/ post-centrifugation: 4°C* | *Pre-storage/ pre-centrifugation: 1.5, 24, 48h Post-storage/ post-centrifugation: 0, 24h* | *($^1$H) NMR, Nightingale Health* / (T)* | *Pre: 23 Post:25* | *Pre-storage/pre-centrifugation: **stable**. Post-storage/post-centrifugation: mean **decrease** up to 0.1SD.* |
| **Apolipoprotein B** | Clark et al. 2003[6] | Pre-storage: pre-centrifugation; | Potassium EDTA-plasma; Non-fasting; Reference: centrifuged immediately | 4°C, 21°C | 0,1-4, 7 days | Clinical Chemistry | 12 | **Stable** (mean percentage change less than 0.5% per day) |
| | Oddoze et al. 2012[8] | Pre-storage: pre-centrifugation; | Serum, lithium heparin and fluoride plasma; Fasting status not specified; Serum reference: 0.5h; Plasma reference: centrifuged immediately | 4°C, 25°C | 0, 2, 4, 6, 24h | Clinical Chemistry / (T) | 10 | **Stable**. |
| | *Current report* | *Pre-storage: pre-centrifugation; Post-storage: post-centrifugation;* | *Serum, potassium EDTA-plasma; Non-fasting; Reference, pre-storage: 1.5h, 4°C; Reference, post-storage: no sample or NMR analysis delay* | *Pre-storage/ pre-centrifugation: 4°C, 21°C Post-storage/ post-centrifugation: 4°C* | *Pre-storage/ pre-centrifugation: 1.5, 24, 48h Post-storage/ post-centrifugation: 0, 24h* | *($^1$H) NMR, Nightingale Health* / (T)* | *Pre: 23 Post:25* | *Pre-storage/pre-centrifugation: **stable** at 4°C; mean increase up to 0.1SD at 21°C. Post-storage/post-centrifugation: **stable**.* |

| | | | | | | | | |
|---|---|---|---|---|---|---|---|---|
| **Fatty Acids** | | | | | | | | |
| **Total fatty acids** | Jobard et al. 2016[13] | Pre-storage: pre and ǂpost-centrifugation; | Serum, heparin-plasma; Fasting; Reference (pre-centrifugation): 1h,22°C; Reference (post-centrifugation): 15min | Pre-centrifugation: 4°C, 22°C Post-centrifugation: 22°C | Pre-centrifugation: 1h (4°C), 6h (4°C, 22°C); Post-centrifugation: 15min, 1h | ($^{1}H$) NMR (1D, CPMG, NOESY, ($^{1}H$-$^{13}C$) HSQC, STOCSY, J-res) / (U) | 96 | Pre-storage/pre-centrifugation: **Increase** at 22°C, 6h; **stable** at 4°C, 1 and 6h. Pre-storage/ǂpost-centrifugation: **stable** |
| | Bernini et al. 2011[9] | Pre-storage: pre and ǂpost-centrifugation; | Serum (SST), potassium EDTA-plasma; Fasting status not specified; Reference: centrifuged immediately (for serum after 30 min) and frozen immediately | Pre-centrifugation: 4°C, 25°C Post-centrifugation: 25°C | Pre-centrifugation: 0-4h Post-centrifugation: 0,6,12, 24h | ($^{1}H$) NMR (1D, NOESY CPMG) / (U) | Pre:6 Post:5 | Pre-storage/pre-centrifugation: **stable;** Pre-storage/ǂpost-centrifugation: **decrease**. |
| | *Current report* | *Pre-storage: pre-centrifugation; Post-storage: post-centrifugation;* | *Serum, potassium EDTA-plasma; Non-fasting; Reference, pre-storage: 1.5h, 4°C; Reference, post-storage: no sample or NMR analysis delay* | *Pre-storage/ pre-centrifugation: 4°C, 21°C Post-storage/ post-centrifugation: 4°C* | *Pre-storage/ pre-centrifugation: 1.5, 24, 48h Post-storage/ post-centrifugation: 0, 24h* | *($^{1}H$) NMR, Nightingale Health* / (T)* | *Pre: 20 Post:21* | *Pre-storage/pre-centrifugation: **stable**. Post-storage/post-centrifugation: **stable**.* |
| **Glycolysis related metabolites** | | | | | | | | |
| **Glucose** | Boyanton et al. 2002[7] | Pre-storage: pre and ǂpost-centrifugation; | Serum, lithium heparin-plasma; Non-fasting; Reference: 0.5, 25°C | 25°C | 0.5, 4, 8, 16, 24, 32, 40, 48, 56h | Clinical Chemistry / (T) | 10 | Pre-storage/pre-centrifugation: **decrease** rapidly in 24h, afterwards more slowly. More pronounced in plasma than serum. Pre-storage/ǂpost centrifugation: **stable**. |
| | Oddoze et al. 2012[8] | Pre-storage: pre-centrifugation; | Serum, lithium heparin and fluoride plasma; Fasting status not specified; Serum reference: 0.5h; Plasma reference: centrifuged immediately | 4°C, 25°C | 0, 2, 4, 6, 24h | Clinical Chemistry / (T) | 10 | **Decrease**, greater at 25°C (-10%) than 4°C (-4%). Stable, at 4°C and 25°C, if in sodium fluorite tubes. |
| | Bernini et al. 2011[9] | Pre-storage: pre and ǂpost-centrifugation; | Serum (SST), potassium EDTA-plasma; Fasting status not specified; Reference: centrifuged immediately (for serum | Pre-centrifugation: 4°C, 25°C Post-centrifugation: 25°C | Pre-centrifugation: 0-4h Post-centrifugation: 0,6,12, 24h | ($^{1}H$) NMR (1D, NOESY CPMG) / (U) | Pre:6 Post:5 | Pre-storage/pre-centrifugation: **decrease** (greater in serum); Pre-storage/ǂpost-centrifugation: **stable**. |

| | | | | | | | | |
|---|---|---|---|---|---|---|---|---|
| | | | after 30 min) and frozen immediately | | | | | |
| | Jobard et al. 2016[13] | Pre-storage: pre and ǂpost-centrifugation; | Serum, heparin-plasma; Fasting; Reference (pre-centrifugation): 1h,22°C; Reference (post-centrifugation): 15min | Pre-centrifugation: 4°C, 22°C Post-centrifugation: 22°C | Pre-centrifugation: 1h (4°C), 6h (4°C, 22°C); Post-centrifugation: 15min, 1h | ($^1$H) NMR (1D, CPMG, NOESY, ($^1$H-$^{13}$C) HSQC, STOCSY, J-res) / (U) | 96 | Pre-storage/pre-centrifugation: **decrease** at 22°C, 6h; **stable** at 4°C, 1 and 6h. Pre-storage/ǂpost-centrifugation: **stable.** |
| | Kamlage et al. 2014 [14] | Pre-storage: pre and ǂpost-centrifugation; | Potassium EDTA -plasma; Fasting status not specified; Pooled plasma | Pre-centrifugation: wet ice, RT (19-22°C) Post-centrifugation: 4°C, 12°C, RT (19-22°C) | Pre-centrifugation: 2, 6h Post-centrifugation: 0, 0.5, 2, 5, 16h | GC-MS, LC-MS/MS, MxP® Broad profiling, MxP®Lipids, Catecholamines, Eicosanoids / (U and T) | 20 | Pre-storage/pre-centrifugation: **decrease** Pre-storage/ǂpost-centrifugation: **stable.** |
| | Fliniaux et al. 2011[15] | Pre-storage: pre-centrifugation; | Serum (SST); Fasting status not specified; Reference: 4h, RT | 4°C, RT | 4, 24h | ($^1$H) NMR (1D and CPMG, ($^1$H-$^{13}$C) HSQC, J-res) / (U) | 7 | **Decrease** at RT. **Stable** at 4°C. |
| | Bervoets et al 2015[16] | Pre-storage: pre-centrifugation; | LiHe plasma; Fasting; Reference: 0.5, ice | 4°C | 0.5, 3, 8h | ($^1$H) NMR (1D, CPMG) / (U) | 20 | **Decrease** |
| | *Current report* | *Pre-storage: pre-centrifugation; Post-storage: post-centrifugation;* | *Serum, potassium EDTA-plasma; Non-fasting; Reference, pre-storage: 1.5h, 4°C; Reference, post-storage: no sample or NMR analysis delay* | *Pre-storage/ pre-centrifugation: 4°C, 21°C Post-storage/ post-centrifugation: 4°C* | *Pre-storage/ pre-centrifugation: 1.5, 24, 48h Post-storage/ post-centrifugation: 0, 24h* | *($^1$H) NMR, Nightingale Health* / (T)* | *Pre: 23 Post:25* | *Pre-storage/pre-centrifugation: mean **decrease** up to 1.4SD (more pronounced at 21°C). Post-storage/post-centrifugation: **stable**.* |
| **Lactate** | Boyanton et al. 2002 [7] | Pre-storage: pre and ǂpost-centrifugation; | Serum, lithium heparin-plasma; Non-fasting; Reference: 0.5, 25°C | 25°C | 0.5, 4, 8, 16, 24, 32, 40, 48, 56h | Clinical Chemistry / (T) | 10 | Pre-storage/pre-centrifugation: **increase**. Pre-storage/ǂpost centrifugation: **stable**. |
| | Oddoze et al. 2012 [8] | Pre-storage: pre-centrifugation; | Serum, lithium heparin and fluoride plasma; Fasting status not specified; Serum reference: 0.5h; Plasma reference: centrifuged immediately | 4°C, 25°C | 0, 2, 4, 6, 24h | Clinical Chemistry / (T) | 10 | **Increase.** Stable, at 4°C and 25°C, if in sodium fluorite tubes. |
| | Bernini et al. 2011 [9] | Pre-storage: pre and ǂpost centrifugation; | Serum (SST), potassium EDTA-plasma; Fasting status not specified; Reference: centrifuged immediately | Pre-centrifugation: 4°C, 25°C Post-centrifugation: 25°C | Pre-centrifugation: 0-4h Post-centrifugation: 0,6,12, 24h | ($^1$H) NMR (1D, NOESY CPMG) / (U) | Pre:6 Post:5 | Pre-storage/pre-centrifugation: **increase**; Pre-storage/ǂpost-centrifugation: **stable**. |

| | | | | | | | | |
|---|---|---|---|---|---|---|---|---|
| | | | (for serum after 30 min) and frozen immediately | | | | | |
| | Jobard et al. 2016 [13] | Pre-storage: pre and ǂpost-centrifugation; | Serum, heparin-plasma; Fasting; Reference (pre-centrifugation): 1h,22ºC; Reference (post-centrifugation): 15min | Pre-centrifugation: 4ºC, 22ºC Post-centrifugation: 22ºC | Pre-centrifugation: 1h (4ºC), 6h (4ºC, 22ºC); Post-centrifugation: 15min, 1h | ($^1$H) NMR (1D, CPMG, NOESY, ($^1$H-$^{13}$C) HSQC, STOCSY, J-res) / (U) | 96 | Pre-storage/pre-centrifugation: **decrease** at 22ºC, 6h; **stable** at 4ºC, 1 and 6h. Pre-storage/ǂpost-centrifugation: **stable** |
| | Kamlage et al. 2014 [14] | Pre-storage: pre and ǂpost-centrifugation; | Potassium EDTA -plasma; Fasting status not specified; Pooled plasma | Pre-centrifugation: wet ice, RT (19-22ºC) Post-centrifugation: 4ºC, 12ºC, RT (19-22ºC) | Pre-centrifugation: 2, 6h Post-centrifugation: 0, 0.5, 2, 5, 16h | GC-MS, LC-MS/MS, MxP® Broad profiling, MxP®Lipids, Catecholamines, Eicosanoids / (U and T) | 20 | Pre-storage/pre-centrifugation: **increase** Pre-storage/ǂpost-centrifugation: **stable.** |
| | Fliniaux et al. 2011 [15] | Pre-storage: pre-centrifugation; | Serum (SST); Fasting status not specified; Reference: 4h, RT | 4ºC, RT | 4, 24h | ($^1$H) NMR (1D and CPMG, ($^1$H-$^{13}$C) HSQC, J-res) / (U) | 7 | **Increase** at RT. **Stable** at 4ºC. |
| | Trezzi et al 2016[17] | Pre-storage: pre-centrifugation; | Potassium EDTA-plasma; Fasting status not specified; | 4ºC, RT (18-23ºC) | U: 10, 30 and 60 min T: 0.5, 3, 23h | GC-MS, clinical chemistry/ (U and T) | U: 3 T: 10 | **Increase** |
| | Bervoets et al 2015[16] | Pre-storage: pre-centrifugation; | LiHe plasma; Fasting; Reference: 0.5, ice | 4ºC | 0.5, 3, 8h | ($^1$H) NMR (1D, CPMG) / (U) | 20 | **Increase** |
| | *Current report* | *Pre-storage: pre-centrifugation; Post-storage: post-centrifugation;* | *Serum, potassium EDTA-plasma; Non-fasting; Reference, pre-storage: 1.5h, 4ºC; Reference, post-storage: no sample or NMR analysis delay* | *Pre-storage/ pre-centrifugation: 4ºC, 21ºC Post-storage/ post-centrifugation: 4ºC* | *Pre-storage/ pre-centrifugation: 1.5, 24, 48h Post-storage/ post-centrifugation: 0, 24h* | *($^1$H) NMR, Nightingale Health* / (T)* | *Pre: 23 Post:25* | *Pre-storage/pre-centrifugation: mean **increase** up to 1.4SD (more pronounced at 21ºC). Post-storage/post-centrifugation: **stable**.* |
| **Pyruvate** | Bernini et al. 2011 [9] | Pre-storage: pre and ǂpost-centrifugation; | Serum (SST), potassium EDTA-plasma; Fasting status not specified; Reference: centrifuged immediately (for serum after 30 min) and frozen immediately | Pre-centrifugation: 4ºC, 25ºC Post-centrifugation: 25ºC | Pre-centrifugation: 0-4h Post-centrifugation: 0,6,12, 24h | ($^1$H) NMR (1D, NOESY CPMG) / (U) | Pre:6 Post:5 | Pre-storage/pre-centrifugation: **plasma**: slightly **decrease/stable** at 4ºC; **increase** at 25ºC; **serum**: **decreases** at 4ºC, **stable** at 25ºC Pre-storage/ǂpost-centrifugation: **stable**. |
| | Nishiumi et al 2017[18] | Pre-storage: pre-centrifugation; | Sodium EDTA-plasma; Fasting status not specified; | Cold temperature, room temperature | Cold temperature: 1, 4, 8h Room temperature: 0, 15, 30 min | GC-MS, LC-MS/MS/(T) | 1 | Room temperature: **increase**; Cold temperature: **decrease**. |

| | | | | | | | | |
|---|---|---|---|---|---|---|---|---|
| | Kamlage et al. 2014 [14] | Pre-storage: pre and ‡post-centrifugation; | Potassium EDTA -plasma; Fasting status not specified; Pooled plasma | Pre-centrifugation: wet ice, RT (19-22°C)<br>Post-centrifugation: 4°C, 12°C, RT (19-22°C) | Pre-centrifugation: 2, 6h<br>Post-centrifugation: 0, 0.5, 2, 5, 16h | GC-MS, LC-MS/MS, MxP® Broad profiling, MxP®Lipids, Catecholamines, Eicosanoids / (U and T) | 20 | Pre-storage/pre-centrifugation: **decrease.**<br>Pre-storage/‡post-centrifugation: **decrease.** |
| | Bervoets et al 2015 [16] | Pre-storage: pre-centrifugation; | LiHe plasma; Fasting; Reference: 0.5, ice | 4°C | 0.5, 3, 8h | ($^1$H) NMR (1D, CPMG) / (U) | 20 | **Decrease** |
| | *Current report* | *Pre-storage: pre-centrifugation; Post-storage: post-centrifugation;* | *Serum; Non-fasting; Reference, pre-storage: 1.5h, 4°C; Reference, post-storage: no sample or NMR analysis delay* | *Pre-storage/ pre-centrifugation: 4°C, 21°C*<br>*Post-storage/ post-centrifugation: 4°C* | *Pre-storage/ pre-centrifugation: 1.5, 24, 48h*<br>*Post-storage/ post-centrifugation: 0, 24h* | *($^1$H) NMR, Nightingale Health* / (T)* | *Pre:20*<br>*Post:37* | *Pre-storage/pre-centrifugation: slight **decrease/stable** at 4°C; mean increase of 1.2SD at 21°C. Post-storage/post-centrifugation: **stable**.* |
| **Citrate** | Jobard et al. 2016 [13] | Pre-storage: pre and ‡post-centrifugation; | Serum, heparin-plasma; Fasting; Reference (pre-centrifugation): 1h,22°C; Reference (post-centrifugation): 15min | Pre-centrifugation: 4°C, 22°C<br>Post-centrifugation: 22°C | Pre-centrifugation: 1h (4°C), 6h (4°C, 22°C);<br>Post-centrifugation: 15min, 1h | ($^1$H) NMR (1D, CPMG, NOESY, ($^1$H-$^{13}$C) HSQC, STOCSY, J-res) / (U) | 96 | Pre-storage/pre-centrifugation: **Increase** at 22°C, 6h (but Variable Importance in Projection <1); **stable** at 4°C, 1 and 6h.<br>Pre-storage/‡post-centrifugation: **stable** |
| | Bernini et al. 2011 [9] | Pre-storage: pre and ‡post-centrifugation; | Serum (SST), potassium EDTA-plasma; Fasting status not specified; Reference: centrifuged immediately (for serum after 30 min) and frozen immediately | Pre-centrifugation: 4°C, 25°C<br>Post-centrifugation: 25°C | Pre-centrifugation: 0-4h<br>Post-centrifugation: 0,6,12, 24h | ($^1$H) NMR (1D, NOESY CPMG) / (U) | Pre:6<br>Post:5 | Pre-storage/pre-centrifugation: **stable;**<br>Pre-storage/‡post-centrifugation: **increase**. |
| | *Current report* | *Pre-storage: pre-centrifugation; Post-storage: post-centrifugation;* | *Serum, potassium EDTA-plasma; Non-fasting; Reference, pre-storage: 1.5h, 4°C; Reference, post-storage: no sample or NMR analysis delay* | *Pre-storage/ pre-centrifugation: 4°C, 21°C*<br>*Post-storage/ post-centrifugation: 4°C* | *Pre-storage/ pre-centrifugation: 1.5, 24, 48h*<br>*Post-storage/ post-centrifugation: 0, 24h* | *($^1$H) NMR, Nightingale Health* / (T)* | *Pre:23*<br>*Post:25* | *Pre-storage/pre-centrifugation: changes up to 0.2SD. Post-storage/post-centrifugation: serum **stable**; EDTA-plasma, **increase** up to 0.3SD.* |

| | | | | | | | | |
|---|---|---|---|---|---|---|---|---|
| **Glycerol** | Jobard et al. 2016 [13] | Pre-storage: pre and ǂpost-centrifugation; | Serum, heparin-plasma; Fasting; Reference (pre-centrifugation): 1h,22°C; Reference (post-centrifugation): 15min | Pre-centrifugation: 4°C, 22°C Post-centrifugation: 22°C | Pre-centrifugation: 1h (4°C), 6h (4°C, 22°C); Post-centrifugation: 15min, 1h | ($^1$H) NMR (1D, CPMG, NOESY, ($^1$H-$^{13}$C) HSQC, STOCSY, J-res) / (U) | 96 | Pre-storage/pre-centrifugation: **Increase** at 22°C, 6h (but Variable Importance in Projection <1); **stable** at 4°C, 1 and 6h. Pre-storage/ǂpost-centrifugation: **stable** |
| | Kamlage et al. 2014 [14] | Pre-storage: pre and ǂpost-centrifugation; | Potassium EDTA -plasma; Fasting status not specified; Pooled plasma | Pre-centrifugation: wet ice, RT (19-22°C) Post-centrifugation: 4°C, 12°C, RT (19-22°C) | Pre-centrifugation: 2, 6h Post-centrifugation: 0, 0.5, 2, 5, 16h | GC-MS, LC-MS/MS, MxP® Broad profiling, MxP®Lipids, Catecholamines, Eicosanoids / (U and T) | 20 | Pre-storage/pre-centrifugation: **decrease.** Pre-storage/ǂpost-centrifugation: **increase.** |
| | *Current report* | *Pre-storage: pre-centrifugation; Post-storage: post-centrifugation;* | *Serum, potassium EDTA-plasma; Non-fasting; Reference, pre-storage: 1.5h, 4°C; Reference, post-storage: no sample or NMR analysis delay* | *Pre-storage/ pre-centrifugation: 4°C, 21°C Post-storage/ post-centrifugation: 4°C* | *Pre-storage/ pre-centrifugation: 1.5, 24, 48h Post-storage/ post-centrifugation: 0, 24h* | *($^1$H) NMR, Nightingale Health* / (T)* | *Pre:23 Post:37* | *Pre-storage/pre-centrifugation: **stable** at 21°C; mean increase of 0.5SD at 4°C. Post-storage/post-centrifugation: **stable**.* |
| **Amino Acids** | | | | | | | | |
| **Alanine** | Jobard et al. 2016 [13] | Pre-storage: pre and ǂpost-centrifugation; | Serum, heparin-plasma; Fasting; Reference (pre-centrifugation): 1h,22°C; Reference (post-centrifugation): 15min | Pre-centrifugation: 4°C, 22°C Post-centrifugation: 22°C | Pre-centrifugation: 1h (4°C), 6h (4°C, 22°C); Post-centrifugation: 15min, 1h | ($^1$H) NMR (1D, CPMG, NOESY, ($^1$H-$^{13}$C) HSQC, STOCSY, J-res) / (U) | 96 | Pre-storage/pre-centrifugation: **increase** at 22°C, 6h (but for serum Variable Importance in Projection <1); **stable** at 4°C, 1 and 6h. Pre-storage/ǂpost-centrifugation: **stable** |
| | Breier et al. 2014[12] | Pre-storage: pre-centrifugation; | Serum, potassium EDTA-plasma; Fasting; Reference, EDTA-plasma: centrifuged immediately; Reference, serum: 0.5h, 21°C. | Serum and EDTA-plasma: 4°C; EDTA-plasma: 21°C | Serum and EDTA-plasma: 0, 3, 6, 24h EDTA-plasma: 24h | ESI-LC-MS/MS, MS/MS, Biocrates*/ (S) | 22 | EDTA-plasma: **increase**; |
| | *Current report* | *Pre-storage: pre-centrifugation; Post-storage: post-centrifugation;* | *Serum, potassium EDTA-plasma; Non-fasting; Reference, pre-storage: 1.5h, 4°C; Reference, post-storage: no sample or NMR analysis delay* | *Pre-storage/ pre-centrifugation: 4°C, 21°C Post-storage/ post-centrifugation: 4°C* | *Pre-storage/ pre-centrifugation: 1.5, 24, 48h Post-storage/ post-centrifugation: 0, 24h* | *($^1$H) NMR, Nightingale Health* / (T)* | *Pre:23 Post:25* | *Pre-storage/pre-centrifugation: mean **increase** up to 1.2SD (more pronounced at 21°C). Post-storage/post-centrifugation: **stable**.* |

| | | | | | | | | |
|---|---|---|---|---|---|---|---|---|
| **Glutamine** | Jobard et al. 2016 [13] | Pre-storage: pre and ǂpost-centrifugation; | Serum, heparin-plasma; Fasting; Reference (pre-centrifugation): 1h,22°C; Reference (post-centrifugation): 15min | Pre-centrifugation: 4°C, 22°C Post-centrifugation: 22°C | Pre-centrifugation: 1h (4°C), 6h (4°C, 22°C); Post-centrifugation: 15min, 1h | $(^1H)$ NMR (1D, CPMG, NOESY, $(^1H$-$^{13}C)$ HSQC, STOCSY, J-res) / (U) | 96 | Pre-storage/pre-centrifugation: **decrease** at 22°C, 6h (but Variable Importance in Projection <1); **stable** at 4°C, 1 and 6h.<br>Pre-storage/ǂpost-centrifugation: **stable** |
| | ǂAnton et al. 2015[10] | Pre-storage: ǂpost-centrifugation; | Serum; Fasting; Reference: max 5h, on ice | Dry ice, wet ice, RT (22-24°C) | 0,12, 24, 36h | FIA-ESI-MS/MS, Biocrates*/ (S) | 19 (males) | **Decrease** (22-24°C and wet ice) |
| | *Current report* | *Pre-storage: pre-centrifugation; Post-storage: post-centrifugation;* | *Serum, potassium EDTA-plasma; Non-fasting; Reference, pre-storage: 1.5h, 4°C; Reference, post-storage: no sample or NMR analysis delay* | *Pre-storage/ pre-centrifugation: 4°C, 21°C Post-storage/ post-centrifugation: 4°C* | *Pre-storage/ pre-centrifugation: 1.5, 24, 48h Post-storage/ post-centrifugation: 0, 24h* | *$(^1H)$ NMR, Nightingale Health* / (T)* | *Pre: 23 Post:25* | *Pre-storage/pre-centrifugation: mean de**crease** up to 0.9SD (more pronounced at 21°C). Post-storage/post-centrifugation: changes up to 0.2SD.* |
| **Histidine** | Jobard et al. 2016 [13] | Pre-storage: pre and ǂpost-centrifugation; | Serum, heparin-plasma; Fasting; Reference (pre-centrifugation): 1h,22°C; Reference (post-centrifugation): 15min | Pre-centrifugation: 4°C, 22°C Post-centrifugation: 22°C | Pre-centrifugation: 1h (4°C), 6h (4°C, 22°C); Post-centrifugation: 15min, 1h | $(^1H)$ NMR (1D, CPMG, NOESY, $(^1H$-$^{13}C)$ HSQC, STOCSY, J-res) / (U) | 96 | Pre-storage/pre-centrifugation: **Increase** at 22°C, 6h (but Variable Importance in Projection <1); **stable** at 4°C, 1 and 6h.<br>Pre-storage/ǂpost-centrifugation: **stable** |
| | Bernini et al. 2011[9] | Pre-storage: pre and ǂpost-centrifugation; | Serum (SST), potassium EDTA-plasma; Fasting status not specified; Reference: centrifuged immediately (for serum after 30 min) and frozen immediately | Pre-centrifugation: 4°C, 25°C Post-centrifugation: 25°C | Pre-centrifugation: 0-4h Post-centrifugation: 0,6,12, 24h | $(^1H)$ NMR (1D, NOESY CPMG) / (U) | Pre:6 Post:5 | Pre-storage/pre-centrifugation: **stable;**<br>Pre-storage/ǂpost-centrifugation: **decrease**. |
| | Breier et al. 2014[12] | Pre-storage: pre-centrifugation; | Serum, potassium EDTA-plasma; Fasting; Reference, EDTA-plasma: centrifuged immediately; Reference, serum: 0.5h, 21°C. | Serum and EDTA-plasma: 4°C; EDTA-plasma: 21°C | Serum and EDTA-plasma: 0, 3, 6, 24h EDTA-plasma: 24h | ESI-LC-MS/MS, MS/MS, Biocrates*/ (S) | 22 | EDTA-plasma: **increase** only at 21°C. |
| | *Current report* | *Pre-storage: pre-centrifugation; Post-storage: post-centrifugation;* | *Serum, potassium EDTA-plasma; Non-fasting; Reference, pre-storage: 1.5h, 4°C; Reference, post-storage: no sample or* | *Pre-storage/ pre-centrifugation: 4°C, 21°C Post-storage/ post-centrifugation: 4°C* | *Pre-storage/ pre-centrifugation: 1.5, 24, 48h Post-storage/ post-centrifugation: 0, 24h* | *$(^1H)$ NMR, Nightingale Health* / (T)* | *Pre:23 Post:25* | *Pre-storage/pre-centrifugation: serum, mean **increase** up to 1.2SD; EDTA-plasma, mean **decrease** up to 0.8SD. Post-storage/post-centrifugation:* |

| | | | | | | | | |
|---|---|---|---|---|---|---|---|---|
| | | | *NMR analysis delay* | | | | | ***decrease** up to 1.4SD.* |
| **Glycine** | Jobard et al. 2016 [13] | Pre-storage: pre and ‡post-centrifugation; | Serum, heparin-plasma; Fasting; Reference (pre-centrifugation): 1h,22°C; Reference (post-centrifugation): 15min | Pre-centrifugation: 4°C, 22°C Post-centrifugation: 22°C | Pre-centrifugation: 1h (4°C), 6h (4°C, 22°C); Post-centrifugation: 15min, 1h | ($^{1}$H) NMR (1D, CPMG, NOESY, ($^{1}$H-$^{13}$C) HSQC, STOCSY, J-res) / (U) | 96 | Pre-storage/pre-centrifugation: **Increase** at 22°C, 6h (but Variable Importance in Projection <1); **stable** at 4°C, 1 and 6h. Pre-storage/‡post-centrifugation: **stable** |
| | ‡Anton et al. 2015[10] | Pre-storage: ‡post-centrifugation; | Serum; Fasting; Reference:max 5h, on ice | Dry ice, wet ice, RT (22-24°C) | 0,12, 24, 36h | FIA-ESI-MS/MS, Biocrates*/ (S) | 19 (males) | **Increase** (at RT i.e. 22-24°C) |
| | Breier et al. 2014[12] | Pre-storage: pre-centrifugation; | Serum, potassium EDTA-plasma; Fasting; Reference, EDTA-plasma: centrifuged immediately; Reference, serum: 0.5h, 21°C. | Serum and EDTA-plasma: 4°C; EDTA-plasma: 21°C | Serum and EDTA-plasma: 0, 3, 6, 24h EDTA-plasma: 24h | ESI-LC-MS/MS, MS/MS, Biocrates*/ (S) | 22 | Serum: **increase.** |
| | *Current report* | *Pre-storage: pre-centrifugation; Post-storage: post-centrifugation;* | *Serum; Non-fasting; Reference, pre-storage: 1.5h, 4°C; Reference, post-storage: no sample or NMR analysis delay* | *Pre-storage/ pre-centrifugation: 4°C, 21°C Post-storage/ post-centrifugation: 4°C* | *Pre-storage/ pre-centrifugation: 1.5, 24, 48h Post-storage/ post-centrifugation: 0, 24h* | *($^{1}$H) NMR, Nightingale Health* / (T)* | *Pre:23 Post:37* | *Pre-storage/pre-centrifugation: mean **increase** up to 1SD. Post-storage/post-centrifugation: mean **increase** up to 0.3SD.* |
| ***Branched-chain Amino Acids*** | | | | | | | | |
| **Isoleucine** | Jobard et al. 2016 [13] | Pre-storage: pre and ‡post-centrifugation; | Serum, heparin-plasma; Fasting; Reference (pre-centrifugation): 1h,22°C; Reference (post-centrifugation): 15min | Pre-centrifugation: 4°C, 22°C Post-centrifugation: 22°C | Pre-centrifugation: 1h (4°C), 6h (4°C, 22°C); Post-centrifugation: 15min, 1h | ($^{1}$H) NMR (1D, CPMG, NOESY, ($^{1}$H-$^{13}$C) HSQC, STOCSY, J-res) / (U) | 96 | Pre-storage/pre-centrifugation: **Increase** at 22°C, 6h (but Variable Importance in Projection <1); **stable** at 4°C, 1 and 6h. Pre-storage/‡post-centrifugation: **stable** |
| | ‡Anton et al. 2015[10] | Pre-storage: ‡post-centrifugation; | Serum; Fasting; Reference:max 5h, on ice | Dry ice, wet ice, RT (22-24°C) | 0,12, 24, 36h | FIA-ESI-MS/MS, Biocrates*/ (S) | 19 (males) | **Increase** (at RT i.e. 22-24°C) |
| | Breier et al. 2014[12] | Pre-storage: pre-centrifugation; | Serum, potassium EDTA-plasma; Fasting; Reference, EDTA-plasma: centrifuged immediately; Reference, serum: 0.5h, 21°C. | Serum and EDTA-plasma: 4°C; EDTA-plasma: 21°C | Serum and EDTA-plasma: 0, 3, 6, 24h EDTA-plasma: 24h | ESI-LC-MS/MS, MS/MS, Biocrates*/ (S) | 22 | EDTA-plasma: **increase** only at 21°C. |
| | *Current report* | *Pre-storage: pre-centrifugation; Post-storage: post-centrifugation;* | *Serum, potassium EDTA-plasma; Non-fasting;* | *Pre-storage/ pre-centrifugation: 4°C, 21°C* | *Pre-storage/ pre-centrifugation: 1.5, 24, 48h* | *($^{1}$H) NMR, Nightingale Health* / (T)* | *Pre:23 Post:25* | *Pre-storage/pre-centrifugation: EDTA-plasma, stable; serum, mean* |

| | | | | | | | | |
|---|---|---|---|---|---|---|---|---|
| | | | *Reference, pre-storage: 1.5h, 4°C; Reference, post-storage: no sample or NMR analysis delay* | *Post-storage/ post-centrifugation: 4°C* | *Post-storage/ post-centrifugation: 0, 24h* | | | ***increase** up to 0.4SD. Post-storage/post-centrifugation: stable.* |
| **Leucine** | Jobard et al. 2016 [13] | Pre-storage: pre and ‡post-centrifugation; | Serum, heparin-plasma; Fasting; Reference (pre-centrifugation): 1h,22°C; Reference (post-centrifugation): 15min | Pre-centrifugation: 4°C, 22°C Post-centrifugation: 22°C | Pre-centrifugation: 1h (4°C), 6h (4°C, 22°C); Post-centrifugation: 15min, 1h | ($^{1}$H) NMR (1D, CPMG, NOESY, ($^{1}$H-$^{13}$C) HSQC, STOCSY, J-res) / (U) | 96 | Pre-storage/pre-centrifugation: **Increase** at 22°C, 6h (but Variable Importance in Projection <1); **stable** at 4°C, 1 and 6h. Pre-storage/‡post-centrifugation: **stable.** |
| | Breier et al. 2014[12] | Pre-storage: pre-centrifugation; | Serum, potassium EDTA-plasma; Fasting; Reference, EDTA-plasma: centrifuged immediately; Reference, serum: 0.5h, 21°C. | Serum and EDTA-plasma: 4°C; EDTA-plasma: 21°C | Serum and EDTA-plasma: 0, 3, 6, 24h EDTA-plasma: 24h | ESI-LC-MS/MS, MS/MS, Biocrates*/ (S) | 22 | **Increase**. |
| | *Current report* | *Pre-storage: pre-centrifugation; Post-storage: post-centrifugation;* | *Serum, potassium EDTA-plasma; Non-fasting; Reference, pre-storage: 1.5h, 4°C; Reference, post-storage: no sample or NMR analysis delay* | *Pre-storage/ pre-centrifugation: 4°C, 21°C Post-storage/ post-centrifugation: 4°C* | *Pre-storage/ pre-centrifugation: 1.5, 24, 48h Post-storage/ post-centrifugation: 0, 24h* | *($^{1}$H) NMR, Nightingale Health* / (T)* | *Pre:23 Post:25* | *Pre-storage/pre-centrifugation: mean **increase** up to 0.8SD. Post-storage/post-centrifugation: **stable**.* |
| **Valine** | Jobard et al. 2016 [13] | Pre-storage: pre and ‡post-centrifugation; | Serum, heparin-plasma; Fasting; Reference (pre-centrifugation): 1h,22°C; Reference (post-centrifugation): 15min | Pre-centrifugation: 4°C, 22°C Post-centrifugation: 22°C | Pre-centrifugation: 1h (4°C), 6h (4°C, 22°C); Post-centrifugation: 15min, 1h | ($^{1}$H) NMR (1D, CPMG, NOESY, ($^{1}$H-$^{13}$C) HSQC, STOCSY, J-res) / (U) | 96 | Pre-storage/pre-centrifugation: **Increase** at 22°C, 6h (but Variable Importance in Projection <1); **stable** at 4°C, 1 and 6h. Pre-storage/‡post-centrifugation: **stable.** |
| | *Current report* | *Pre-storage: pre-centrifugation; Post-storage: post-centrifugation;* | *Serum, potassium EDTA-plasma; Non-fasting; Reference, pre-storage: 1.5h, 4°C; Reference, post-storage: no sample or NMR analysis delay* | *Pre-storage/ pre-centrifugation: 4°C, 21°C Post-storage/ post-centrifugation: 4°C* | *Pre-storage/ pre-centrifugation: 1.5, 24, 48h Post-storage/ post-centrifugation: 0, 24h* | *($^{1}$H) NMR, Nightingale Health* / (T)* | *Pre:23 Post:25* | *Pre-storage/pre-centrifugation: mean **increase** up to 0.8SD. Post-storage/post-centrifugation: **stable**.* |
| ***Aromatic Amino Acids*** | | | | | | | | |
| **Phenylalanine** | Jobard et al. 2016 [13] | Pre-storage: pre and ‡post-centrifugation; | Serum, heparin-plasma; Fasting; Reference (pre-centrifugation): 1h,22°C; Reference | Pre-centrifugation: 4°C, 22°C Post-centrifugation: 22°C | Pre-centrifugation: 1h (4°C), 6h (4°C, 22°C); Post-centrifugation: 15min, 1h | ($^{1}$H) NMR (1D, CPMG, NOESY, ($^{1}$H-$^{13}$C) HSQC, STOCSY, J-res) / (U) | 96 | Pre-storage/pre-centrifugation: **Increase** at 22°C, 6h (but Variable Importance in Projection <1); **stable** at 4°C, 1 and 6h. |

| | | | | | | | | |
|---|---|---|---|---|---|---|---|---|
| | | | (post-centrifugation): 15min | | | | | Pre-storage/‡post-centrifugation: **stable.** |
| | ‡Anton et al. 2015[10] | Pre-storage: ‡post-centrifugation; | Serum; Fasting; Reference:max 5h, on ice | Dry ice, wet ice, RT (22-24°C) | 0,12, 24, 36h | FIA-ESI-MS/MS, Biocrates*/ (S) | 19 (males) | **Increase** (at RT i.e. 22-24°C) |
| | Breier et al. 2014[12] | Pre-storage: pre-centrifugation; | Serum, potassium EDTA-plasma; Fasting; Reference, EDTA-plasma: centrifuged immediately; Reference, serum: 0.5h, 21°C. | Serum and EDTA-plasma: 4°C; EDTA-plasma: 21°C | Serum and EDTA-plasma: 0, 3, 6, 24h EDTA-plasma: 24h | ESI-LC-MS/MS, MS/MS, Biocrates*/ (S) | 22 | EDTA-plasma: **increase** only at 21°C. Serum: **increase.** |
| | *Current report* | *Pre-storage: pre-centrifugation; Post-storage: post-centrifugation;* | *Serum, potassium EDTA-plasma; Non-fasting; Reference, pre-storage: 1.5h, 4°C; Reference, post-storage: no sample or NMR analysis delay* | *Pre-storage/ pre-centrifugation: 4°C, 21°C Post-storage/ post-centrifugation: 4°C* | *Pre-storage/ pre-centrifugation: 1.5, 24, 48h Post-storage/ post-centrifugation: 0, 24h* | *$(^{1}H)$ NMR, Nightingale Health* / (T)* | *Pre:23 Post:25* | *Pre-storage/pre-centrifugation: mean **increase** up to 1.2SD; Post-storage/post-centrifugation: serum, mean **increase** up to 1.1SD EDTA-plasma, **decrease** up to 0.4SD.* |
| **Tyrosine** | Jobard et al. 2016 [13] | Pre-storage: pre and ‡post-centrifugation; | Serum, heparin-plasma; Fasting; Reference (pre-centrifugation): 1h,22°C; Reference (post-centrifugation): 15min | Pre-centrifugation: 4°C, 22°C Post-centrifugation: 22°C | Pre-centrifugation: 1h (4°C), 6h (4°C, 22°C); Post-centrifugation: 15min, 1h | $(^{1}H)$ NMR (1D, CPMG, NOESY, $(^{1}H-^{13}C)$ HSQC, STOCSY, J-res) / (U) | 96 | Pre-storage/pre-centrifugation: **Increase** at 22°C, 6h (but Variable Importance in Projection <1); **stable** at 4°C, 1 and 6h. Pre-storage/‡post-centrifugation: **stable.** |
| | Breier et al. 2014[12] | Pre-storage: pre-centrifugation; | Serum, potassium EDTA-plasma; Fasting; Reference, EDTA-plasma: centrifuged immediately; Reference, serum: 0.5h, 21°C. | Serum and EDTA-plasma: 4°C; EDTA-plasma: 21°C | Serum and EDTA-plasma: 0, 3, 6, 24h EDTA-plasma: 24h | ESI-LC-MS/MS, MS/MS, Biocrates*/ (S) | 22 | EDTA-plasma: **increase** only at 21°C. Serum: **increase.** |
| | *Current report* | *Pre-storage: pre-centrifugation; Post-storage: post-centrifugation;* | *Serum, potassium EDTA-plasma; Non-fasting; Reference, pre-storage: 1.5h, 4°C; Reference, post-storage: no sample or NMR analysis delay* | *Pre-storage/ pre-centrifugation: 4°C, 21°C Post-storage/ post-centrifugation: 4°C* | *Pre-storage/ pre-centrifugation: 1.5, 24, 48h Post-storage/ post-centrifugation: 0, 24h* | *$(^{1}H)$ NMR, Nightingale Health* / (T)* | *Pre:23 Post:25* | *Pre-storage/pre-centrifugation: mean **increase** up to 0.5SD; Post-storage/post-centrifugation: **stable**.* |
| **Ketone bodies** | | | | | | | | |
| **Acetate** | Jobard et al. 2016 [13] | Pre-storage: pre and ‡post-centrifugation; | Serum, heparin-plasma; Fasting; Reference (pre-centrifugation): | Pre-centrifugation: 4°C, 22°C Post-centrifugation: 22°C | Pre-centrifugation: 1h (4°C), 6h (4°C, 22°C); | $(^{1}H)$ NMR (1D, CPMG, NOESY, $(^{1}H-^{13}C)$ HSQC, STOCSY, J-res) / (U) | 96 | Pre-storage/pre-centrifugation: **decrease** at 22°C, 6h (but Variable Importance in Projection <1); |

| | | | | | | | | |
|---|---|---|---|---|---|---|---|---|
| | | | 1h,22°C; Reference (post-centrifugation): 15min | | Post-centrifugation: 15min, 1h | | | **stable** at 4°C, 1 and 6h. Pre-storage/‡post-centrifugation: **stable.** |
| | *Current report* | *Pre-storage: pre-centrifugation; Post-storage: post-centrifugation;* | *Serum, potassium EDTA-plasma; Non-fasting; Reference, pre-storage: 1.5h, 4°C; Reference, post-storage: no sample or NMR analysis delay* | *Pre-storage/ pre-centrifugation: 4°C, 21°C Post-storage/ post-centrifugation: 4°C* | *Pre-storage/ pre-centrifugation: 1.5, 24, 48h Post-storage/ post-centrifugation: 0, 24h* | *($^1$H) NMR, Nightingale Health* / (T)* | *Pre:23 Post:25* | *Pre-storage/pre-centrifugation: serum, mean* ***increase*** *up to 0.9SD; EDTA-plasma, mean* ***decrease*** *up to 0.9SD. Post-storage/post-centrifugation: mean* ***increase*** *up to 0.5SD.* |
| **Beta-hydroxybutyrate** | Jobard et al. 2016 [13] | Pre-storage: pre and ‡post-centrifugation; | Serum, heparin-plasma; Fasting; Reference (pre-centrifugation): 1h,22°C; Reference (post-centrifugation): 15min | Pre-centrifugation: 4°C, 22°C Post-centrifugation: 22°C | Pre-centrifugation: 1h (4°C), 6h (4°C, 22°C); Post-centrifugation: 15min, 1h | ($^1$H) NMR (1D, CPMG, NOESY, ($^1$H-$^{13}$C) HSQC, STOCSY, J-res) / (U) | 96 | Pre-storage/pre-centrifugation: **decrease** at 22°C, 6h (but Variable Importance in Projection <1); **stable** at 4°C, 1 and 6h. Pre-storage/‡post-centrifugation: **stable.** |
| | *Current report* | *Pre-storage: pre-centrifugation; Post-storage: post-centrifugation;* | *Serum, potassium EDTA-plasma; Non-fasting; Reference, pre-storage: 1.5h, 4°C; Reference, post-storage: no sample or NMR analysis delay* | *Pre-storage/ pre-centrifugation: 4°C, 21°C Post-storage/ post-centrifugation: 4°C* | *Pre-storage/ pre-centrifugation: 1.5, 24, 48h Post-storage/ post-centrifugation: 0, 24h* | *($^1$H) NMR, Nightingale Health* / (T)* | *Pre:23 Post:25* | *Pre-storage/pre-centrifugation:* ***stable****. Post-storage/post-centrifugation: mean* ***increase*** *up to 0.4SD.* |
| **Fluid Balance** | | | | | | | | |
| **Creatinine** | Clark et al. 2003 [6] | Pre-storage: pre-centrifugation; | Potassium EDTA-plasma; Non-fasting; Reference: centrifuged immediately | 4°C, 21°C | 0, 1-4, 7 days | Clinical Chemistry / (T) | 12 | **Stable at 4°C** (mean percentage change less than 0.5% per day); **Increase** more than 5% per day at **21°C**. |
| | Boyanton et al. 2002 [7] | Pre-storage: pre and ‡post-centrifugation; | Serum, lithium heparin-plasma; Non-fasting; Reference: 0.5, 25°C | 25°C | 0.5, 4, 8, 16, 24, 32, 40, 48, 56h | Clinical Chemistry / (T) | 10 | Pre-storage/pre-centrifugation: **Increase** by 110% in plasma and 60% in serum after 24h (possibly due to interference of pseudo-creatinines in the assay). Pre-storage/‡post-centrifugation: **stable**. |
| | Oddoze et al. 2012 [8] | Pre-storage: pre-centrifugation; | Serum, lithium heparin and fluoride plasma; Fasting status not specified; | 4°C, 25°C | 0, 2, 4, 6, 24h | Clinical Chemistry / (T) | 10 | **Stable**. |

| | | | | | | | | |
|---|---|---|---|---|---|---|---|---|
| | | | Serum reference: 0.5h; Plasma reference: centrifuged immediately | | | | | |
| | Jobard et al. 2016 [13] | Pre-storage: pre and ǂpost-centrifugation; | Serum, heparin-plasma; Fasting; Reference (pre-centrifugation): 1h,22°C; Reference (post-centrifugation): 15min | Pre-centrifugation: 4°C, 22°C Post-centrifugation: 22°C | Pre-centrifugation: 1h (4°C), 6h (4°C, 22°C); Post-centrifugation: 15min, 1h | ($^1$H) NMR (1D, CPMG, NOESY, ($^1$H-$^{13}$C) HSQC, STOCSY, J-res) / (U) | 96 | Pre-storage/pre-centrifugation: **decrease** at 22°C, 6h (but Variable Importance in Projection <1); **stable** at 4°C, 1 and 6h. Pre-storage/ǂpost-centrifugation: **stable.** |
| | *Current report* | *Pre-storage: pre-centrifugation; Post-storage: post-centrifugation;* | *Serum, potassium EDTA-plasma; Non-fasting; Reference, pre-storage: 1.5h, 4°C; Reference, post-storage: no sample or NMR analysis delay* | *Pre-storage/ pre-centrifugation: 4°C, 21°C Post-storage/ post-centrifugation: 4°C* | *Pre-storage/ pre-centrifugation: 1.5, 24, 48h Post-storage/ post-centrifugation: 0, 24h* | *($^1$H) NMR, Nightingale Health* / (T)* | *Pre:23 Post:25* | *Pre-storage/pre-centrifugation: **stable**. Post-storage/post-centrifugation: mean **increase** up to 0.4SD.* |
| **Albumin** | Clark et al. 2003 [6] | Pre-storage: pre-centrifugation; | Potassium EDTA-plasma; Non-fasting; Reference: centrifuged immediately | 4°C, 21°C | 0, 1-4, 7 days | Clinical Chemistry / (T) | 12 | **Stable** (mean percentage change less than 0.5% per day) |
| | Boyanton et al. 2002 [7] | Pre-storage: pre and ǂpost centrifugation; | Serum, lithium heparin-plasma; Non-fasting; Reference: 0.5, 25°C | 25°C | 0.5, 4, 8, 16, 24, 32, 40, 48, 56h | Clinical Chemistry / (T) | 10 | Pre-storage/pre-centrifugation: **Increase** after 24h of 7%. Pre-storage/ǂpost-centrifugation: **stable**. |
| | Oddoze et al. 2012 [8] | Pre-storage: pre-centrifugation; | Serum, lithium heparin and fluoride plasma; Fasting status not specified; Serum reference: 0.5h; Plasma reference: centrifuged immediately | 4°C, 25°C | 0, 2, 4, 6, 24h | Clinical Chemistry / (T) | 10 | **Stable**. |
| | Bernini et al. 2011[9] | Pre-storage: pre and ǂpost-centrifugation; | Serum (SST), potassium EDTA-plasma; Fasting status not specified; Reference: centrifuged immediately (for serum after 30 min) and frozen immediately | Pre-centrifugation: 4°C, 25°C Post-centrifugation: 25°C | Pre-centrifugation: 0-4h Post-centrifugation: 0,6,12, 24h | ($^1$H) NMR (1D, NOESY CPMG) / (U) | Pre:6 Post:5 | Pre-storage/pre-centrifugation: **stable;** Pre-storage/ǂpost-centrifugation: **decrease**. |
| | *Current report* | *Pre-storage: pre-centrifugation; Post-storage: post-centrifugation;* | *Serum, potassium EDTA-plasma; Non-fasting;* | *Pre-storage/ pre-centrifugation: 4°C, 21°C* | *Pre-storage/ pre-centrifugation: 1.5, 24, 48h* | *($^1$H) NMR, Nightingale Health* / (T)* | *Pre:23 Post:25* | *Pre-storage/pre-centrifugation: **stable** at 4°C; mean increase up to 0.6SD at 21°C;* |

| | | | | | | | | |
|---|---|---|---|---|---|---|---|---|
| | | | *Reference, pre-storage: 1.5h, 4ºC; Reference, post-storage: no sample or NMR analysis delay* | *Post-storage/ post-centrifugation: 4ºC* | *Post-storage/ post-centrifugation: 0, 24h* | | | *Post-storage/post-centrifugation:* ***stable****.* |
| **Other untargeted NMR studies** | | | | | | | | |
| Not applicable | ǂBarton et al. 2008[19] | Post-storage: ǂpost-centrifugation; | Serum; Fasting status not specified; | 4ºC | 0, 24, 36h | ($^1$H) NMR (1D, NOESY), ICL NPC* / (U) | 40 | Alterations in proteins and protein fragments. |
| Not applicable | Teahan et al. 2006[20] | Pre-storage: pre and ǂpost-centrifugation; | Serum (pre-centrifugation only), heparin-plasma; Non-fasting; Reference: 0.5h | pre-centrifugation: Ice, RT post-centrifugation: RT | 0.5,1,2,3h | ($^1$H) NMR (1D, NOESY, CPMG), ICL NPC* / (U) | 4 | Pre-storage/pre-centrifugation: **stable** overall on ice. Pre-storage/ǂpost-centrifugation: **stable** overall. Changes in some lipids and possibly some low-molecular-weight metabolites. |

10. Anton G, Wilson R, Yu ZH, Prehn C, Zukunft S, Adamski J, Heier M, Meisinger C, Romisch-Margl W, Wang-Sattler R, Hveem K, Wolfenbuttel B, Peters A, Kastenmuller G and Waldenberger M.

Pre-analytical sample quality: metabolite ratios as an intrinsic marker for prolonged room temperature exposure of serum samples. *PloS one*. 2015;10:e0121495.
11. Pinto J, Domingues MR, Galhano E, Pita C, Almeida Mdo C, Carreira IM and Gil AM. Human plasma stability during handling and storage: impact on NMR metabolomics. *Analyst*. 2014;139:1168-77.
12. Breier M, Wahl S, Prehn C, Fugmann M, Ferrari U, Weise M, Banning F, Seissler J, Grallert H, Adamski J and Lechner A. Targeted Metabolomics Identifies Reliable and Stable Metabolites in Human Serum and Plasma Samples. *PloS one*. 2014;9.
13. Jobard E, Tredan O, Postoly D, Andre F, Martin AL, Elena-Herrmann B and Boyault S. A Systematic Evaluation of Blood Serum and Plasma Pre-Analytics for Metabolomics Cohort Studies. *Int J Mol Sci*. 2016;17.
14. Kamlage B, Maldonado SG, Bethan B, Peter E, Schmitz O, Liebenberg V and Schatz P. Quality markers addressing preanalytical variations of blood and plasma processing identified by broad and targeted metabolite profiling. *Clin Chem*. 2014;60:399-412.
15. Fliniaux O, Gaillard G, Lion A, Cailleu D, Mesnard F and Betsou F. Influence of common preanalytical variations on the metabolic profile of serum samples in biobanks. *Journal of Biomolecular Nmr*. 2011;51:457-465.
16. Bervoets L, Louis E, Reekmans G, Mesotten L, Thomeer M, Adriaensens P and Linsen L. Influence of preanalytical sampling conditions on the H-1 NMR metabolic profile of human blood plasma and introduction of the Standard PREanalytical Code used in biobanking. *Metabolomics*. 2015;11:1197-1207.
17. Trezzi JP, Bulla A, Bellora C, Rose M, Lescuyer P, Kiehntopf M, Hiller K and Betsou F. LacaScore: a novel plasma sample quality control tool based on ascorbic acid and lactic acid levels. *Metabolomics*. 2016;12:96.
18. Nishiumi S, Suzuki M, Kobayashi T and Yoshida M. Differences in metabolite profiles caused by pre-analytical blood processing procedures. *J Biosci Bioeng*. 2018;125:613-618.
19. Barton RH, Nicholson JK, Elliott P and Holmes E. High-throughput 1H NMR-based metabolic analysis of human serum and urine for large-scale epidemiological studies: validation study. *Int J Epidemiol*. 2008;37 Suppl 1:i31-40.
20. Teahan O, Gamble S, Holmes E, Waxman J, Nicholson JK, Bevan C and Keun HC. Impact of analytical bias in metabonomic studies of human blood serum and plasma. *Anal Chem*. 2006;78:4307-18.
